# Structural genomics and interactomics of 2019 Wuhan novel coronavirus, 2019-nCoV, indicate evolutionary conserved functional regions of viral proteins

**DOI:** 10.1101/2020.02.10.942136

**Authors:** Hongzhu Cui, Ziyang Gao, Ming Liu, Senbao Lu, Sun Mo, Winnie Mkandawire, Oleksandr Narykov, Suhas Srinivasan, Dmitry Korkin

## Abstract

During its first month, the recently emerged 2019 Wuhan novel coronavirus (2019-nCoV) has already infected many thousands of people in mainland China and worldwide and took hundreds of lives. However, the swiftly spreading virus also caused an unprecedentedly rapid response from the research community facing the unknown health challenge of potentially enormous proportions. Unfortunately, the experimental research to understand the molecular mechanisms behind the viral infection and to design a vaccine or antivirals is costly and takes months to develop. To expedite the advancement of our knowledge we leverage the data about the related coronaviruses that is readily available in public databases, and integrate these data into a single computational pipeline. As a result, we provide a comprehensive structural genomics and interactomics road-maps of 2019-nCoV and use these information to infer the possible functional differences and similarities with the related SARS coronavirus. All data are made publicly available to the research community at http://korkinlab.org/wuhan

## Introduction

Within month and a half, since it was first discovered in Wuhan, China, the novel deadly Coronavirus, 2019-nCov, has infected more than 37,000, with the death toll already surpassing that of the 2003 SARS epidemic [1-3]. In spite of the world-wide efforts by scientific community to swiftly address this health crisis, the vaccines may be months and even years away [4, 5]. For instance, a Phase I trial of a vaccine for a closely related 2019-nCov, severe acute respiratory syndrome (SARS) virus, was announced in December 2004, after two years of the disease outbreak [6]. And a vaccine for Middle East respiratory syndrome (MERS), another coronavirus that emerged in 2012, was patented in 2019, with the Phase I trials introduced in the same year [7, 8]. Nevertheless, in the past two decades, a massive amount of work has been done to understand the molecular basis of the coronavirus infection and evolution, develop effective treatment in forms of both vaccines and antiviral drugs, and propose efficient measures for viral detection and prevention [9-18]. Structures of many individual proteins of SARS, MERS, and related coronaviruses, as well as their biological interactions with other viral and host proteins have been explored along with the experimental testing of anti-viral properties of small compounds [16, 19-31].

However, experimental data of the same scale for 2019-nCov may take years to obtain by the research community. Can it be facilitated? The answer lies in the use of the modern bioinformatics methods that can drastically streamline knowledge discovery by quickly providing the important insights about possible molecular mechanisms behind the infection, pinpointing the likely protein targets for the anti-viral treatments, and predicting the efficacy of the existing antivirals developed for other coronaviruses. By leveraging the previously known information on genome sequences as well as protein structure and function, bioinformaticians have been successfully helping the virologists by structurally characterizing proteins of novel viruses, determining the evolutionary trajectories, identifying interactions with host proteins, and providing other important biological insights. In particular, a plethora of results has been achieved through comparative, or homology, modeling principles [32, 33]. In addition to the global structural genomics initiatives focusing on determining the 3D structures of proteins on a genome scale [34] and specific efforts on rapid structural characterization of proteins in emerging viruses [35-38], multiple works have used comparative modeling to predict the structures of protein-protein interaction complexes [39-41], facilitating structure-based drug discovery [33, 42, 43], inferring protein functions [44], determining the macromolecular interaction network [45-47], and providing molecular insights into viral evolution [48-50].

Here, using an integrated bioinformatics approach, we provide a comprehensive structural genomics and interactomics analysis of the Wuhan 2019-nCov virus. The structural information on the individual 2019-nCov proteins and their interactions with each other and with the human proteins allows to accurately determine the putative functional sites. These functional sites, combined with the evolutionary sequence analysis of 2019-nCov as well as the closely related human SARS and bat coronavirus proteomes, provide us with a structure-based view of the evolutionary diversity of 2019-nCov, allow to estimate how similar the functioning of 2019-nCov virus is when compared with SARS, and forecast how likely the antibodies and candidate ligands that are efficient in inhibiting the SARS functions will be efficient in doing the same for 2019-nCov.

## Results

The recently sequenced genomes of 2019-nCoV strains combined with the comparative analysis of the SARS genome organization and transcription allowed to determine the tentative list of gene products [51]. It has been suggested that 2019-nCoV has 16 predicted non-structural proteins (that are referred to as wNsp1-wNsp16 here) constituting a polyprotein (wORF1ab), followed by (at least) 13 downstream ORFs: <u>S</u>urface, ORF3a, ORF3b, <u>E</u>nvelope, <u>M</u>embrane, ORF6, ORF7a, ORF7b, ORF8, <u>N</u>ucleocapsid, ORF9a, ORF9b, and ORF10 that we refer in this work as wS, wORF3a, wORF3b, wE, wM, wORF6, wORF7a, wORF7b, wORF8, wN, wORF9a, wORF9b, and wORF10, correspondingly. The three viral species whose proteins shared the highest similarity were consistently the same: human SARS (SARS-Cov), bat coronavirus (BtCoV), as well as another bat betacoronavirus (BtRf-BetaCoV).

### Comparative analysis of 2019-nCoV proteins with the evolutionary related coronavirus proteins reveals unevenly distributed large genomic insertions

Searching against UniProt database [52] resulted in matches for the polyprotein (wORF1ab), all four structural proteins (wS, wE, wM, and wN), and six ORFs (wORF3a, wORF6, wORF7a, wORF7b, wORF8, and wORF10) also referred to as accessory proteins. The closest protein matches from UniProt shared sequence identity with the related 2019-nCoV proteins as high as 91% (with wORF1ab and wN) and as low as 57% (withwORF8) (Supl. Table S1). The majority of differences were the single-residue substitutions spread across the protein sequence (see multiple sequence alignment files for all protein in Suppl. Materials).

Perhaps the most profound differences lie in the sequences of the multi-domain protein wNsp3 and surface protein wS: our analysis revealed that compared to the related coronavirus proteins (multiple sequence alignments for both proteins can be found in Suppl. Materials) the two proteins had large sequence inserts. In particular, wNsp3 had a novel large (25 - 41 res., depending on the alignment method) insert between its two putative functional domains homologous to N-terminal domain and adenosine diphosphate ribose 1″ phosphatase (ADRP) of SARS [53, 54] (Suppl. Materials). Interestingly, the closest matching peptide, apart from the virus itself, was found in C-Jun-amino-terminal kinase-interacting protein 4 (Seq. id is 46%) of *Labrus bergylta*, a species of marine ray finned fish. Being significantly more diverse than the other three structural proteins, wS was found to have 4 inserts (4-6 res.) that seemed unique to 2019-nCoV and two additional inserts shared with human SARS proteins (Suppl. Materials).

### Three recent strains of bat SARS-like coronavirus from 2013, 2015, and 2017 share extremely high proteome similarity with 2019-nCoV

The unusually low conservation of ORF6, ORF8, and surface proteins between 2019-nCoV and human SARS, bat coronavirus, as well as another bat betacoronavirus BtRf-BetaCoV, prompted us to perform an expanded search for ORF8 homologs using NCBI BLAST’s blatp tool against a large non-redundant protein sequences repository (nr) [55]. Our search resulted in three new homologs of ORF8 from three different isolates of bat SARS-like coronavirus: bat-SL-CoVZC45 (GenBank ID: MG772933, collected in 2017), bat-SL-CoVZXC21 (GenBank ID: MG772934, collected in 2015), and RaTG13 (GenBank ID: MN996532, collected in 2013) that shared a striking similarity with wORF8, unseen in other strains before: the sequence identities between each of these three homologs and wORF8 ranged between the 94 and 95%. Further analysis showed that the proteomes of these isolates shared even higher sequence identity with the other proteins of 2019-nCoV: from 88.4% to 100% for 2015 and 2017 isolates, and even higher 97.4%-100% for the 2013 isolate (Supl. Table S1).

In spite of the significant similarity of the three isolates to 2019-nCoV, important differences were observed. First, similar to other viruses, the 2017 and 2015 isolates did not have the four sequence inserts that were found in wS. Second, neither of the two isolate had the large insert between the tow domain of wNsp3 of wORF1ab described above. On the contrary, the 2013 isolate had both, the four sequence inserts in its surface protein, matching those in wS and the large insert in Nsp3 although the sequence is different from that one in wNsp3. The main difference between 2019-nCoV and 2013 isolate is the lack of Orf10 in the latter: while genomic sequences of both 2015 and 2017 isolates can be translated into a protein product that shares 97.4% of sequence identity (Supl. Table S1), because of a single nucleotide deletion in that region one cannot translate a full-length ORF10 in RaTG13: the resulting frameshift causes a premature stop-codon when translating the sequence.

### Structural genomics and interactomics analysis of 2019-nCoV

Next, as a result of a comprehensive comparative modeling effort, we were able to structurally characterize 17 individual proteins, including 13 non-structural proteins of wORF1ab (wNsp1, wNsp3, wNsp4, wNsp5, wNsp7, wNsp8, wNsp9, wNsp10, wNsp12, wNsp14, wNsp15, wNsp16), three structural proteins (wE, wN, and wS), as well as one ORF (wORF7a). For two proteins, wNsp3 and wN, multiple individual domains were modeled (Fig.1, Fig.2,). The templates for the majority models were homologous protein structures from other coronaviruses, with a high target-to-template sequence similarity (seq. ids: 75-96%, except for ORF8). For two proteins, wN and wNsp3, multiple domains were modeled. N-terminal and C-terminal domains of wN correspond to N-terminal RNA-binding domain and C-terminal dimerization domain of SARS, respectively. The modeled domains 1-6 of wNsp1 correspond to (1) N-terminal domain; (2) adenosine diphosphate ribose 1″ phosphatase domain; (3) SUD domain (SARS-specific unique domain) containing two macrodomains; (4) SUD domain C; (5) the papain-like protease PLPro domain; and (6) Y domain of SARS (Fig. 1). A previously identified transmembrane domain of SARS Nsp3 was mapped to the sequence of wNsp3, but could not be modeled due to the lack of a template structure.

**Figure 1.**
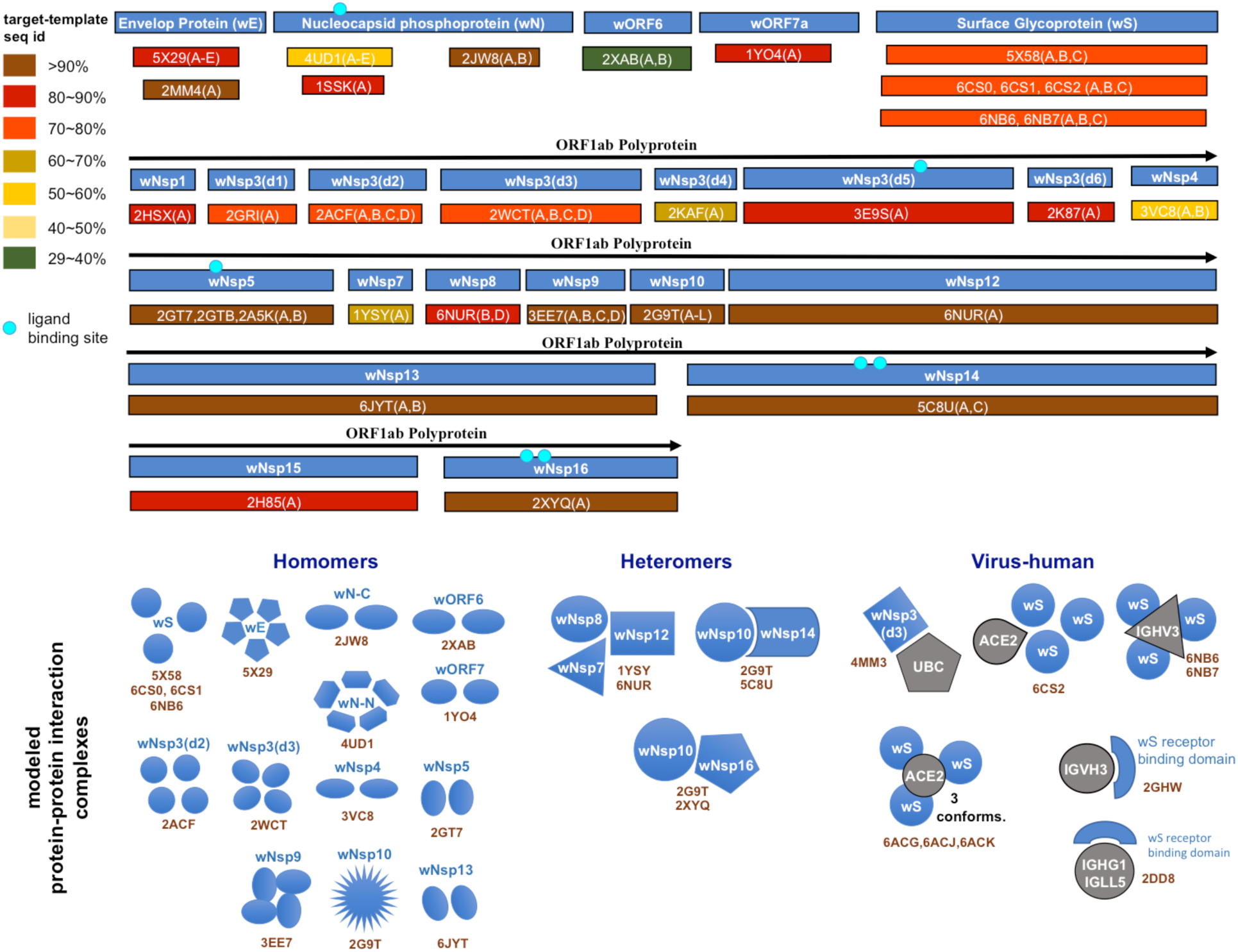
Structural genomics and interactomics road map. Shown are the individual proteins and protein complexes that were targeted for structural characterization together with PDB ID of their templates from Protein Data Bank (PDB).

**Figure 2.**
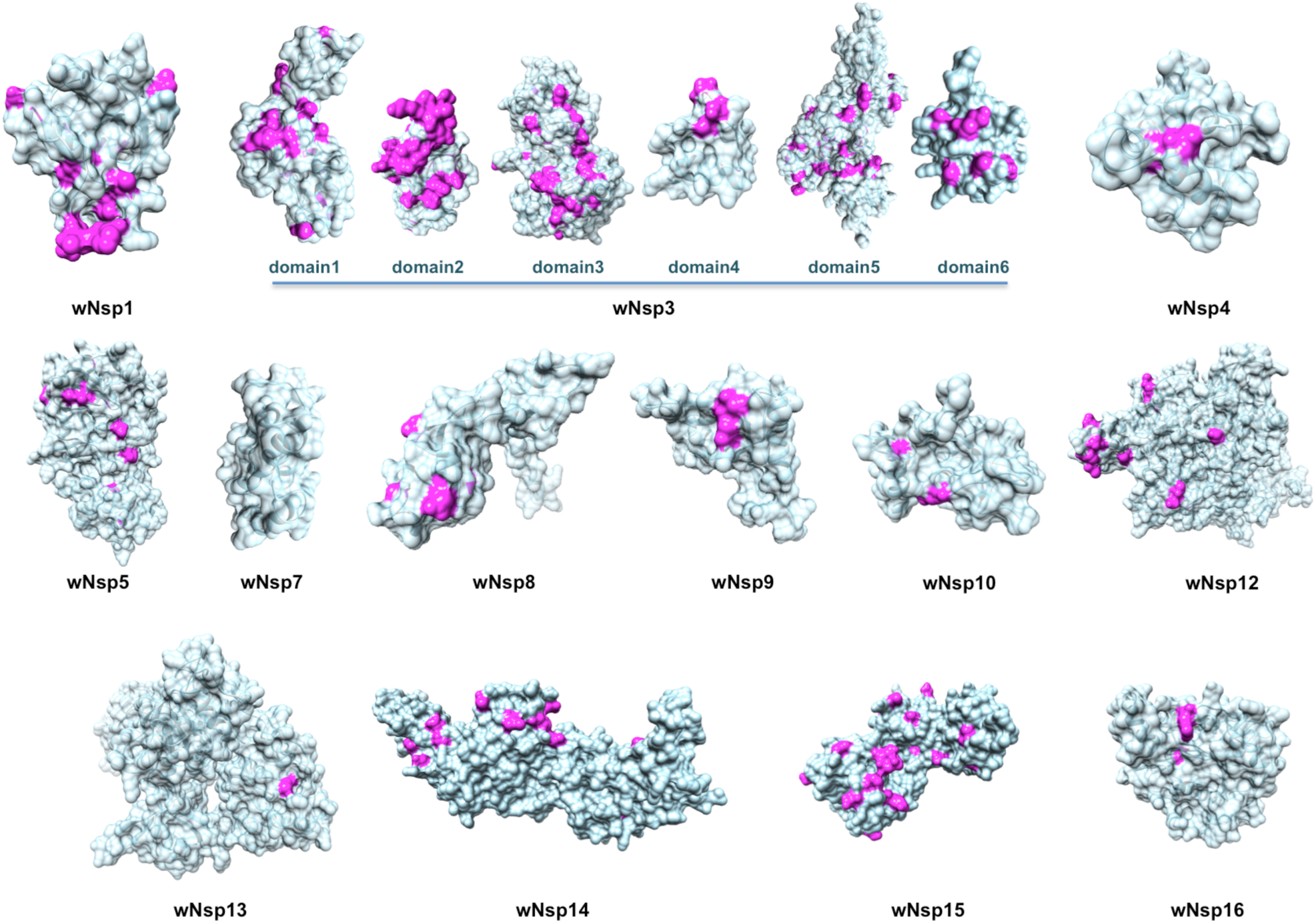
Structurally characterized non-structural proteins of 2019-nCoV. Highlighted in pink are mutations found when aligning the proteins against their homologs from the closest related coronaviruses: 2019-nCoV and human SARS, bat coronavirus, and another bat betacoronavirus BtRf-BetaCoV. The structurally resolved part of wNsp7 is sequentially identical to its homolog.

The structural analysis of the modeled proteins combined with the sequence conservation analysis revealed several findings. First, we found that the mutated residues tend to locate on the protein’s surface, supporting the previous observations in other families of RNA viruses that the core residues of viral proteins are more conserved than the surface residues [48, 56, 57]. Furthermore, in a substantial number of proteins, distributions of mutated positions exhibited spatial patterns, with groups of mutations found to form clusters on the protein surfaces (Fig.2). These obtained models were then used as reference structures to map and analyze the protein-binding and ligand-binding sites.

Next, using comparative modeling we structurally characterized protein interaction complexes, for both intra-viral (homo- and hetero-oligomers) and host-viral interactions where host proteins were exclusively human. In total, we obtained structural models for 16 homo-oligomeric complexes, three hetero-oligomeric complexes, and eight human-virus interaction complexes (Fig. 3). The intra-viral hetero-oligomeric complexes included exclusively the interactions between the non-structural proteins (wNsp7, wNsp8, wNsp10, wNsp12, wNsp14, and wNsp16). The modeled host-viral interaction complexes included three types of interactions: nonstructural protein wNsp3 (papain-like protease, PLpro domain) interacting with human ubiquitin-aldehyde, surface protein wS (in its trimeric form) interacting the human receptor ACE3 in different conformations, as well as the same protein wS interacting with several neutralizing antibodies. Based on the obtained models, the protein interaction binding sites were extracted and analyzed with respect to their evolutionary conservation.

**Figure 3.**
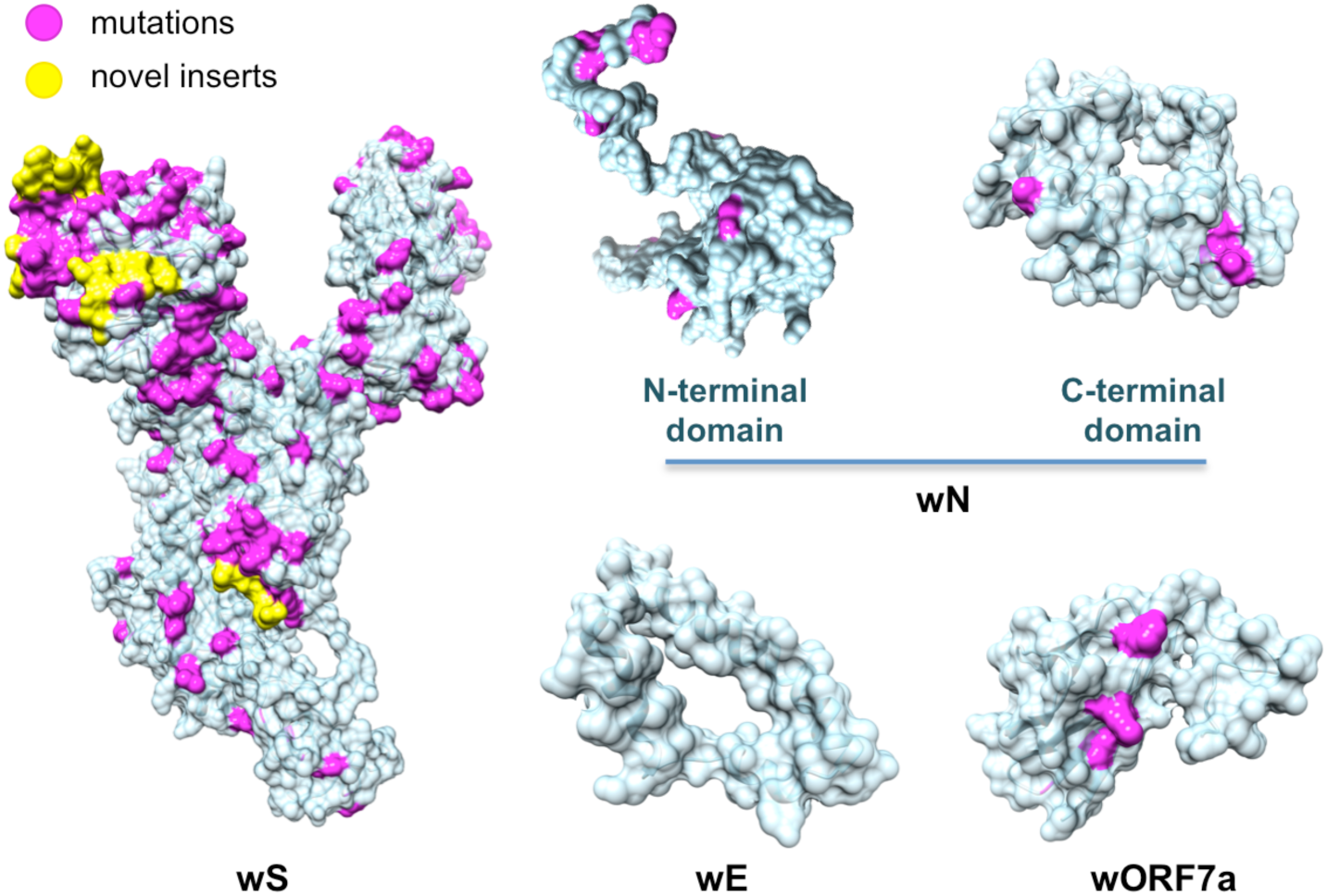
Structurally characterized structural proteins and an ORF of 2019-nCoV. Highlighted in pink are mutations found when aligning the proteins against their homologs from the closest related coronaviruses: 2019-nCoV and human SARS, bat coronavirus, and another bat betacoronavirus BtRf-BetaCoV. Highlighted in yellow are novel protein inserts found in wS.

### Evolutionary conservation and divergence of functional regions of 2019-nCoV

The analysis of the evolutionary conservation of protein binding sites revealed several patterns. First, we found that all protein binding sites of non-structural proteins involving in the intra-viral heteromeric complexes, wNSP7-wNsp8-wNsp12, wNsp10-wNsp14, and wNsp10-wNsp16, are either fully conserved or allow at most one mutation on the periphery of the binding region, in spite of the fact that each proteins had multiple mutations on their surfaces (Fig. 4A, see Suppl. Materials for the alignments of individual proteins and annotation with the protein binding sites). Furthermore, we observed the same behavior when analyzing the interaction between papain-like protease PLpro domain of wNsp3 and with human ubiquitin-aldehyde (Fig. 4B, see Suppl. Materials): the only two mutated residues were located on a border of the binding region and thus were unlikely to disrupt the protein-protein interaction.

**Figure 4.**
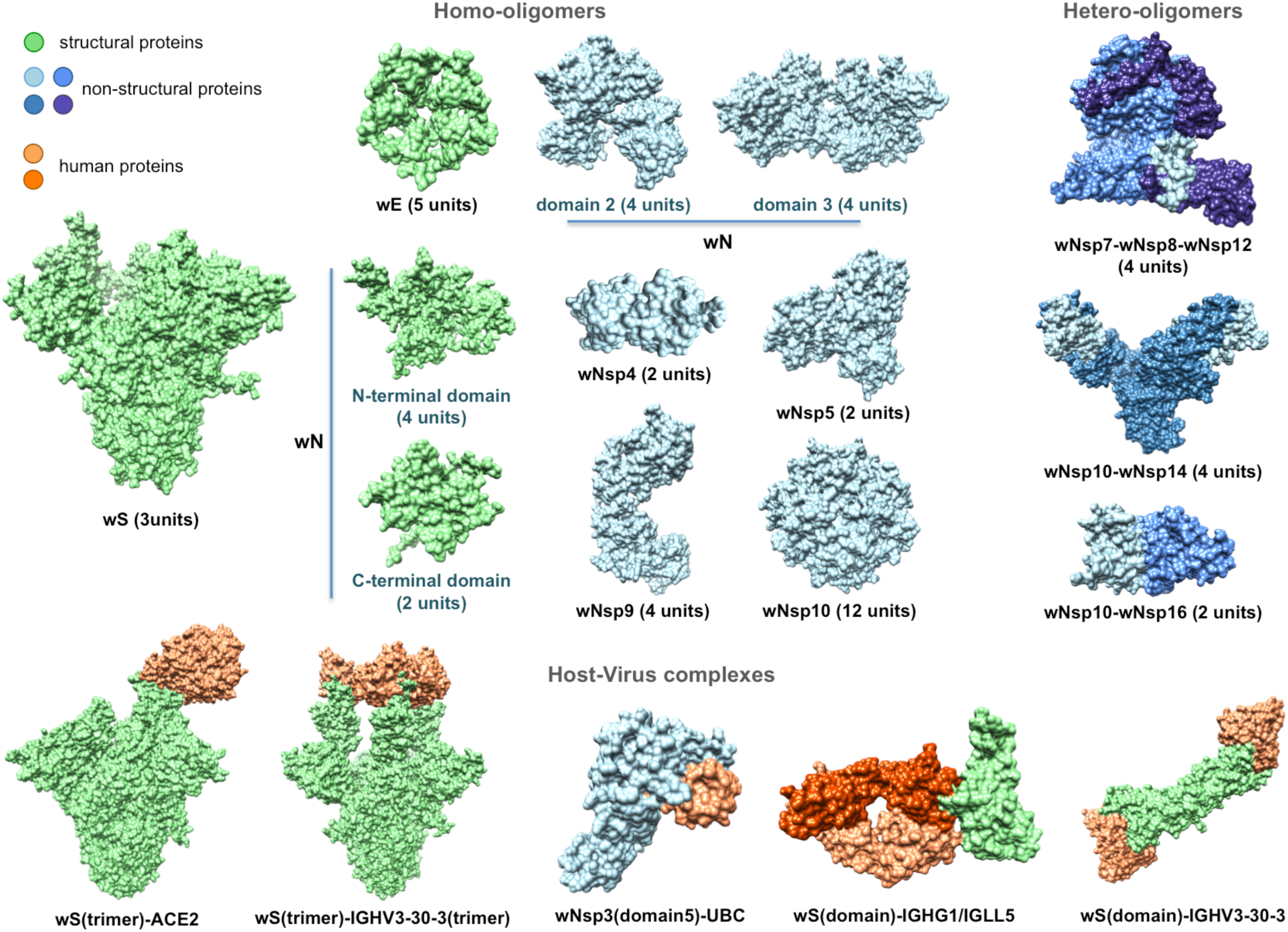
Structurally characterized intra-viral and host-viral protein-protein interaction complexes of 2019-nCoV. Human proteins (colored in orange) are identified through their gene names. For each intra-viral structure, the number of subunits involved in the interaction is specified.

The surface of wS presents a striking contrast to the majority of 2019-nCoV proteins due to its heavily mutated surface (Fig. 3; see Suppl. Materials for the alignment of wS with the related proteins). First, the four novel sequence inserts and two inserts shed with the closest strains were expected to affect the protein’s function. Interestingly, three of the novel inserts are located in the first NTD domain, while the fourth one is located immediately before the S2 cleavage site and inside the homo-trimerization interaction interface. While the RBD domain of wS was not affected by those inserts it was the most heavily mutated region of wS with the likely disruptive functional effects on the interactions with the human ACE2 receptor and monoclonal antibodies mAb 396 [58] and mAb 80R [59] (Fig. 4B). The NTD domain has been considered a target of another antibody Ab G2 previously shown to work in MERS [60]. However, based on the structural superposition of the NTD domain, it is likely that the expected interaction will be disrupted by the novel sequence inserts to wS.

Finally, the analysis of seven ligand binding sites for multiple candidate inhibitors previously identified for four proteins in SARS and MERS showed that many of the LBS were intact in the corresponding 2019-nCoV proteins (Fig. D, see Suppl. Materials for the alignments of individual proteins and annotation with the ligand binding sites), such as wN, wNsp3, wNsp5, and wNsp16. For wNsp14, the ligand binding site for several inhibitors was mutated while a co-localized binding site for another ligand was intact.

### Joint intra-viral and human-virus protein-protein interaction network for SARS CoV indicates potential system-wide roles of 2019-nCoV proteins

After constructing the individual networks of intra-viral interactions and virus-host interactions, they were merged to form a unified network of SARS-CoV interactions, with the hypothesis that most interactions could be conserved in 2019-nCoV (Fig. 6). The network analysis of all three networks showed that unifying the intra-viral and virus-host networks reduces the number of components (islands) in the SARS-CoV-Host interactome (Table 2). This suggested that the virus-host interactome map missed some viral interactions that increased the number of components. The clustering coefficient and average node degree were both higher in the unified interactome compared to the virus-host interactome.

**Table 1.** The list of 2019-nCoV proteins analyzed and structurally characterized in this work.

| protein | accession | length | template<br>PDB id | Trgt-tmplt<br>Seq ID | organism |
| --- | --- | --- | --- | --- | --- |
| wS, surface glycoprotein | YP_009724390 | 1273 | 6ACK | 75% | SARS |
| wE, envelope protein | YP_009724392 | 75 | 5X29 | 89% | SARS |
| wORF7a | YP_009724395 | 121 | 1YO4 | 90% | SARS |
| wN, nucleocapsid<br>phosphoprotein | YP_009724397 | 419 | 2JW8 | 96% | SARS |
|  |  |  | 1SSK | 83% | SARS |
|  |  |  | 4UD1 | 51% | MERS |
| wNsp1 | YP_009725297 | 115 | 2HSX | 86% | SARS |
| wNsp3-domain1 | YP_009725299 | 107 | 2GRI | 79% | SARS |
| wNsp3-domain2 | YP_009725299 | 175 | 2ACF | 72% | SARS |
| wNsp3-domain3 | YP_009725299 | 264 | 2WCT | 76% | SARS |
| wNsp3-domain4 | YP_009725299 | 67 | 2KAF | 70% | SARS |
| wNsp3-domain5 | YP_009725299 | 315 | 3E9S | 82% | SARS |
| wNsp3-domain6 | YP_009725299 | 113 | 2K87 | 82% | SARS |
| wNsp4 | YP_009725300 | 91 | 3VC8 | 60% | MHV |
| wNsp5 | YP_009725301 | 306 | 2GT7 | 96% | SARS |
| wNsp7 | YP_009725302 | 83 | 1YSY | 67% | SARS |
| wNsp8 | YP_009725304 | 114 | 6NUR | 85% | SARS |
| wNsp9 | YP_009725305 | 110 | 3EE7 | 99% | SARS |
| wNsp10 | YP_009725306 | 121 | 2G9T | 98% | SARS |
| wNsp12 | YP_009725307 | 770 | 6NUR | 97% | SARS |
| wNsp13 | YP_009725308 | 596 | 6JYT | 100% | SARS |
| wNsp14 | YP_009725309 | 526 | 5C8U | 95% | SARS |
| wNsp15 | YP_009725310 | 346 | 2H85 | 86% | SARS |
| wNsp16 | YP_009725311 | 288 | 2XYQ | 94% | SARS |

**Table 2.** Network Parameters of the SARS-CoV intraviral, virus-host and unified networks. The table shows topological statistics for the three networks. Among the many computed statistics, the shown parameters include the number of nodes and edges in the networks, the average degree, number of components (independent networks), diameter (maximum shortest path) and clustering coefficient.

| **Network Parameters** | **SARS-CoV**<br>**Intraviral**<br>**Interactome** | **SARS-CoV-Host**<br>**Interactome** | **Unified**<br>**Interactome** |
| --- | --- | --- | --- |
| No. of nodes | 31 | 118 | 125 |
| No. of edges | 86 | 114 | 206 |
| No. of components | 1 | 8 | 2 |
| Diameter | 4 | 14 | 7 |
| Average degree | 4.710 | 1.95 | 3.04 |
| Clustering coefficient | 0.448 | 0.0 | 0.068 |

**Figure 5.**
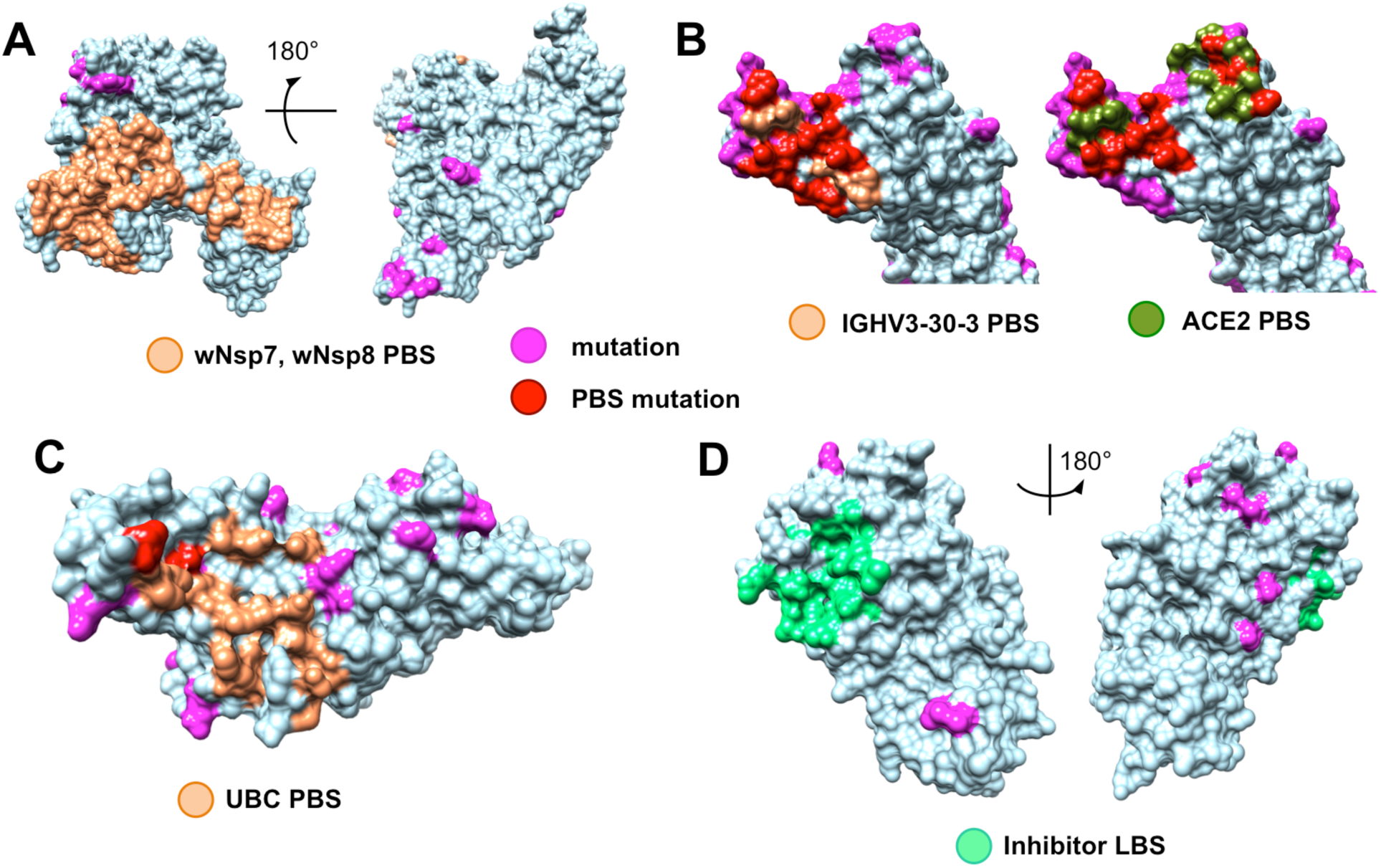
Evolutionary conservation of functional sites in 2019-nCoV proteins. A. Fully conserved protein binding sites (PBS, light orange) of wNsp12 in its interaction with wNsp7 and wNsp8 while other parts of the protein surface shows mutations (magenta); B. Both major monoclonal antibody binding site (light orange) and ACE2 receptor binding site (dark green) of wS are heavily mutated (binding site mutations are shown in red) compared to the same binding sites in other coronaviruses; mutations not located on the two binding sites are shown in magenta; C. Nearly intact protein binding site (light orange) of wNsp (papain-like protease PLpro domain) for its putative interaction with human ubiquitin-aldehyde (binding site mutations for the only two residues are shown in red, non-binding site mutations are shown in magenta); D. Fully conserved inhibitor ligand binding site (LBS, green) for wNsp5; non-binding site mutations are shown in magenta.

**Figure 6.**
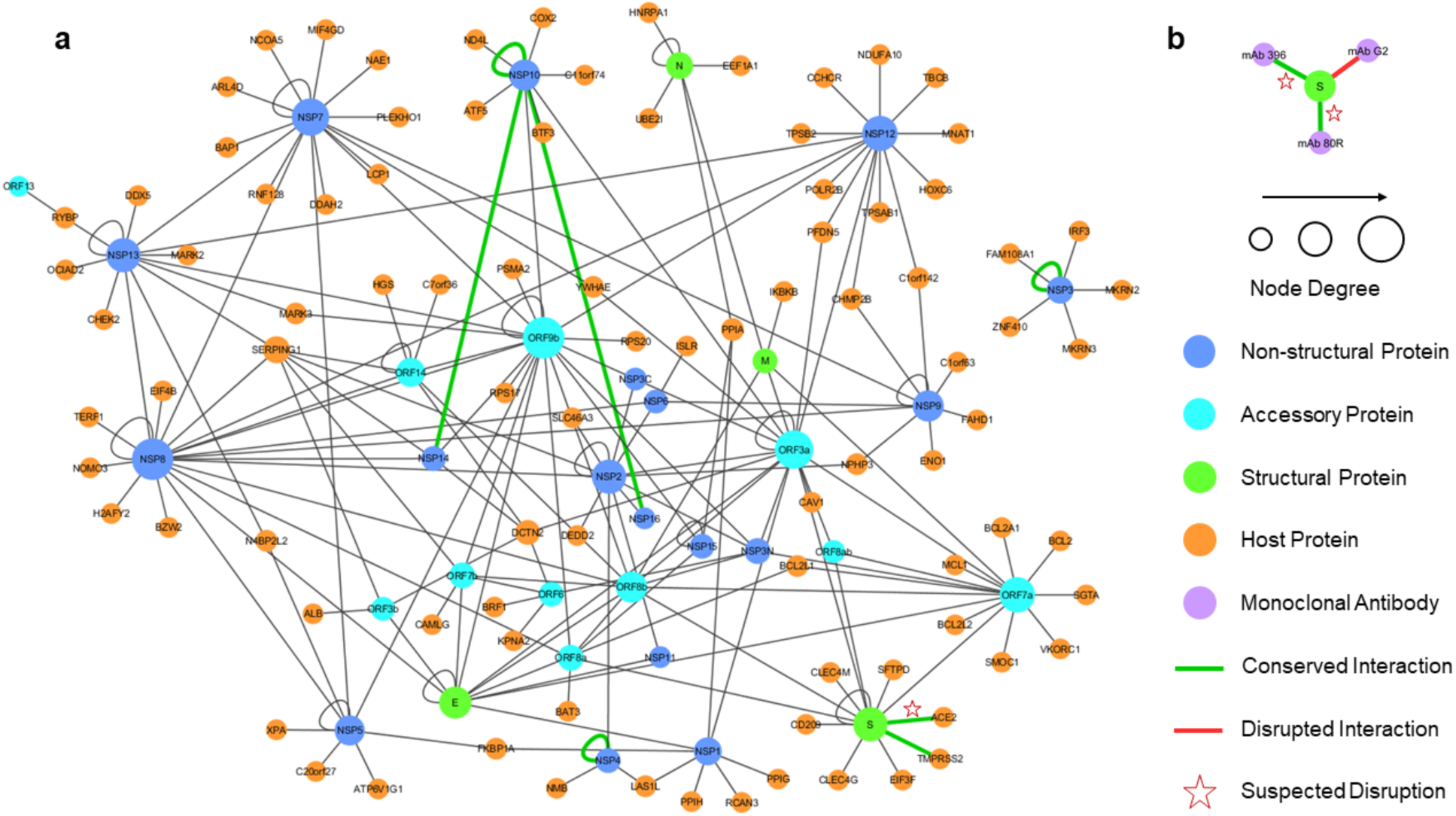
The unified interactome with 2019-nCoV interactions inferred through homology. (a) The SARS-CoV intra-viral and virus-host interactions are merged to create a unified interaction network indicating the types of proteins; the size of a node reflects the node degree. The 2019-nCoV interactions that are inferred from SARS-CoV are represented by green edges, with stars indicating the suspected disruption of the interaction based on the evolutionary conservation of the binding sites. (b) Structural modeling prediction of interaction between 2019-nCoV S protein and three monoclonal antibodies.

The network topology of the unified interactome indicated the presence of several viral hubs for structural, non-structural, as well as accessory proteins (Fig. 6). The viral proteins form all the hubs and there are a few of specific interest. ORF9b is one of the largest hubs in the network thought to play a secondary role only in intra-viral interactions and not necessary for replication [61], but a recent study has shown that ORF9b hinders immunity by targeting the mitochondria and limits host cell responses [62]. NSP8 is the other major hub and interacts with other replicase proteins, including two other hubs, i.e. NSP7 and NSP 12, which together play a crucial role in replication [63].

Another crucial non-structural protein is NSP1, also present in the 2019-nCoV, which is known to be a primary disruptor of innate immunity. The host interaction partners of NSP1 modulate the Calcineurin/NFAT pathway that plays an important role in immune cell activation [64]. Overexpression of NSP1 is associated with immunopathogenecity and long-term cytokine dysregulation as observed in severe SARS cases. And inhibition of cyclophilins/immunopilins (host interaction partners) by cyclosporine A (CspA) blocks the replication of CoVs of all genera.

There is an important interaction between the SARS spike protein (S) and the host angiotensin-converting enzyme 2 (ACE2), as it is associated with cross-species and human-to-human transmissions. The same interaction is also inferred from structural modeling in 2019-nCoV between wS and ACE2, but might be disrupted due to a substantial number of mutations in the receptor binding site of wS while another recent study proposed that the interaction will be preserved [65]. Similar to SARS-CoV [66], wS protein in 2019-nCoV is expected to interact with type II transmembrane protease (TMPRSS2) [67] and is likely to be involved in inhibition of antibody-mediated neutralization. Thus, wS remains the important target for vaccines and drugs previously evaluated in SARS and MERS while a neutralizing antibody targeting the wS protein could provide passive immunity [68]. In addition, there are 7 interactions from SARS-CoV determined from structural characterization of the protein complexes that are predicted to be either conserved or potentially disrupted in 2019-nCoV (green edges in Fig. 1). An important target for vaccines and drugs is the surface (S) protein which has been evaluated in SARS-CoV and MERS-CoV with an idea that a neutralizing antibody targeting wS protein could provide passive immunity for 2019-nCoV [68]. We also structurally modeled interactions between the 2019-nCoV wS protein and three human monoclonal antibodies that were previously studied in SARS-CoV for immunotherapy and mapped the information about the evolutionary conserved and diverse surface residues. We find that two interactions, with mAb 396 [58] and mAb 80R [59], are likely to be disrupted due to the heavily mutated binding sites while another one, Ab G2 [60], is likely to be disrupted due to a novel insert into the sequence of wS. These findings may provide guidelines in the search of potential antibody candidates for treatment.

## Discussion

This work provides an initial large-scale structural genomics and interactomics effort towards understanding the structure, function, and evolution of the 2019-nCoV virus. The goal of this computational work is two-fold. First, by making the structural road map and the related findings fully available to the research community, we aim to facilitate the process of structure-guided research, where accurate structural models of proteins and their interaction complexes already exist. Second, by providing a comparative analysis between the new virus and its closest relatives from the perspective of protein- and ligand-binding, we hope to help experimental scientists in their understanding of the molecular mechanisms implicated in the infection by the new coronavirus as well as in the process of vaccine development and antiviral drug discovery.

Through integrating the information on structure, function, and evolution in a comparative study of 2019-nCoV and the closely related coronaviruses, one can make several preliminary conclusions. First, the extended peptide sequence newly introduced to wNsp3 between two structurally, and possibly functionally, independent domains of this protein, might act as a long inter-domain linker, thus extending the conformational flexibility of this multi-domain protein. Second, the presence of the four novel inserts and one highly variable region of the surface protein wS and the analysis of this large-scale sequence changes with respect to intra-viral and viral-host interaction leads us to conclude that these inserts might have structural impact on the homo-trimeric form of the protein as well as the impact on the functions carried out by NTD domain. Third, the structurally modelable repertoire of 2019-nCoV proteome also pinpoints to the interesting targets for structural biology and hybrid methods. For instance, the whole structures of the multi-domain proteins wN and wNsp3 could be resolved by integrating individual models with the lower-resolution but whole-protein covering techniques such as cryogenic electron microscopy (CryoEM). The structure of wNsp3 is especially interesting because of the presence of the novel peptide introduced between the two structural domains of the protein.

The evolutionary analysis of the protein and ligand-binding sites mapped on the surfaces of 2019-nCoV may provide new insights into the virus functioning and its future treatment. The 100% or near 100% evolutionary conservation of the protein binding sites on the surfaces of nonstructural proteins wNsp7, wNsp8, wNsp10, wNsp12, wNsp14, and wNsp16 that correspond to the intra-viral interactions for three complexes is consistent with our previous observations that the intra-viral interactions are significantly more conserved than viral-host interactions [48, 49, 69]. However the near-perfect conservation of the human ubiquitin-aldehyde protein binding site on the surface of wNsp3 is rather intriguing suggesting the critical role of this interaction in the functioning of SARS-like coronaviruses. Lastly, the conservation of the ligand bind sites for many putative inhibitors previously developed for SARS and MERS suggests the development of antiviral drugs as a promising direction for addressing this new global health threat.

## Materials & Methods

The goal of this work is to identify the evolutionary differences between 2019-nCoV and the closest coronavirus species, human SARS coronavirus and bat SARS-like coronavirus, and to predict their possible functional implications. To do so, we structurally characterize individual proteins as well as intra-viral and human-virus protein complexes, extract the information on their interaction interfaces and ligand binding, and superpose the evolutionary difference and conservation information with the binding information. Specifically, our integrative computational pipeline will include the following five steps.

First, for each of the candidate 2019-nCoV proteins a set of sequentially similar coronavirus proteins is determined and aligned. Second, structural models of 2019-nCoV proteins are obtained using template-based, or homology, modeling. Third, the protein-protein interaction complex structures of 2019-nCoV proteins interacting with each other and/or with the human proteins are determined using a multi-chain comparative modeling protocol. Fourth, the protein-binding sites will be extracted from the obtained models of protein-protein interaction complexes, and protein-ligand binding sites will be extracted from the evolutionary close coronavirus protein-ligand templates and mapped to the relevant structural models of 2019-nCoV through structural alignment. Fifth, the information on the evolutionary differences and conservations observed between the protein sequences of 2019-nCoV and the related coronaviruses and extracted from the above protein sequence alignments will mapped onto the structural models of the 2019-nCoV proteins to determine if the protein- and ligand-binding site are functionally conserved. Lastly, a joint human-virus and virus-virus interactome is predicted through homology and analyzed. All the obtained models and sequence alignments have been made publically available to the research community.

### Protein sequence data collection and analysis

Available sequences for protein candidates wS, wORF3a, wE, wM, wORF6, wORF7a, wORF7b, wORF8, wN, and wORF10 have been extracted from NCBI Virus repository [70] (collected on January 29, 2019) and consequently used further in the sequence analysis and structural modeling (Suppl. Materials). The Uniprot BLAST-based search was performed for each of the proteins using default parameters. From the results of each search, the final selection was done based on the pairwise sequence identity (>60%) as well as the relationship (*Coronaviridae* family). Each of 2019-nCoV proteins was then aligned with the found related coronavirus proteins using a multiple sequence alignment method Clustal Omega [71].

### Structural characterization of protein and protein complexes

The structure of each protein has been determined using a single-template comparative modeling protocols with MODELLER software package [72]. Frist, the template for each protein sequence has been identified using a PSI-BLAST search in Protein Data Bank (PDB) [73]. In general, a structural template with the highest sequence identity has been selected that covers at least 50 residues of the target sequence has at least 30% sequence identity. The polyprotein wORF1ab was first split into 16 putative proteins based on its alignment with the human SARS polyprotein, with each protein independently searched against PDB. In total, structural templates for 17 proteins have been determined (Table 1, Figs. 1-3). In some cases, several independent templates, each covering an individual protein domain, of a large target 2019-nCoV protein were determined. The obtained template was then used in the comparative modeling protocol, generating five models. Each model was then assessed using DOPE statistical potential [74] and the best-scoring model was selected as a final prediction.

To model a protein-protein interaction complex, a multi-chain modeling protocol is used [39]. Specifically, we align the corresponding pairs of homologous proteins and combine them into a single alignment where the individual chains are separated by “/” symbol. The alignment is used as an input together with the multi-chain structural template of a homologous complex. In case of viral-human interaction (these are the only virus-host structural templates found), the human protein remains the same in the alignment. In total, 18 structural templates of protein complexes involving homologs of 14 2019-nCoV proteins were retrieved (Fig. 3). Similar to the single-protein protocol, five candidate models are generated for each complex conformation, and each model is assessed using DOPE statistical potential following selection of the best-scoring model as a final prediction.

### Mapping of functional regions and evolutionary conservation

We next extract protein- and ligand-binding sites and map it onto the models of 2019-nCoV proteins. For protein binding sites, the obtained modeled structures of protein complexes that involve 2019-nCoV proteins are considered. For each 2019-nCoV protein in a protein-protein interaction complex, we identify all binding residues that constitute its protein-binding site. Given an interaction between two proteins, a residue on one protein is defined as a protein binding site residue if there is at least one pair of atoms, one from this residue and another from a residue in the second protein, with VDW surfaces not farther than 1.0 Å. Using this definition, the binding sites are identified with UCSF Chimera [75]. The ligand binding sites are identified and mapped using a different protocol –we rely on the default definition of protein-ligand binding site residues from PDB 3D Ligand View, since this standard is widely accepted. Once all ligand binding sites residues are identified for a related coronavirus, the residues are then mapped onto the surface of the related 2019-nCoV protein using a structural alignment between the two proteins, which puts into a one-to-one correspondence residues from both proteins.

### Inferring intra-viral and virus-host protein-protein interaction network

Next, we leverage the obtained protein homology information to predict and map all possible intra-viral and virus-host protein interactions at a systems level. Since the 2019-nCoV genome exhibits substantial similarity to the 2002 SARS-CoV genome [76] and proteome [77], we hypothesize that many of the interactions observed in the SARS proteome are expected to be preserved in the 2019-nCoV proteome as well, unless the corresponding binding sites were affected. The information on the interactome will help us understanding the global mechanistic processes of the viral molecular machinery during viral infection, survival within the host, and replication. With this knowledge, we can discern the protein interactions that are crucial for transmission and replication—such interactions can be potential candidates for inhibitory drugs [78]. Furthermore, using the systems approach, we can identify hubs and bottlenecks new to 2019-nCoV that could again be targeted by the antiviral drugs.

For this purpose, we create a comprehensive integrated SARS-CoV interactome that consists of both intra-viral and virus-host interactions. The SARS intra-viral interactome was created using published data, where the SARS-CoV ORFeome was cloned and a genome-wide analysis of viral protein interactions through yeast-two-hybrid (Y2H) matrix screens was performed [61]. The Y2H matrix screen was summarized by combining interactions in one direction, both directions and self-interaction. We also included intra-viral interactions gathered from the literature review [61]. The aggregated intra-viral interaction network consists of 31 proteins and 86 unique interactions.

We then construct the SARS-CoV-Host interactome through published interaction data, where first the SARS-CoV ORFeome was cloned for Y2H screens and used as a bait [64]. Specifically, a cDNA library encoding 5,000 different human genes were used that acted as prey molecules. We also include virus-host interactions mined from literature survey of 5,000 abstracts [64]. The curated virus-host interaction network consists of 118 proteins, including 93 host proteins, and 114 unique virus-host interactions. Next, to create an integrated network, the two individual networks were imported in Cytoscape [79] and merged to form a unified interactome representing both intra-viral and virus-host interactions. Finally, we include our predictions from structural modeling of 2019-nCoV intra-viral and virus-host interactions, surveyed recent literature on predicted 2019-nCoV interactions and annotated them accordingly in the unified interactome. The unified interactome consists of 125 proteins (94 host proteins) and 200 unique interactions. In addition, network analysis is performed to compute a set of topological statistics (degree distribution, clustering coefficient, and other important characteristics) to characterize parameters of the three networks, before which the networks are pruned for duplicate edges. Finally, the 200 putative interactions were annotated based on the extent to which the protein binding sites of the 2019-nCoV proteins were altered, compared to their SARS homologs. Based on the evolutionary conservation analysis of the putative protein binding sites extracted from the modeled complexes, some interactions were annotated as potentially disrupted.

## Supporting information

Ligand binding site mapping for wN

Ligand binding site mapping for wNsp3

Ligand binding site mapping for wNsp5 (Non-SARS Group)

Ligand binding site mapping for wNsp5 (SARS Group 1)

Ligand binding site mapping for wNsp5 (SARS Group 2)

Ligand binding site mapping for wNsp14

Ligand binding site mapping for wNsp16

Protein binding site mapping for wNsp7, wNsp8, wNsp10, wNsp 12, wNsp14, wNsp16

Protein binding site mapping for wNsp3

Protein binding site mapping for wS

viral protein sequence alignments

## Availability

All data generated in this work is freely available to the research community via Korkin Lab web-server (korkinlab.org/wuhan).

