## Supplemental Table S1 for "Structural genomics and interactomics of 2019 Wuhan novel coronavirus, 2019-nCoV, indicate evolutionary conserved functional regions of viral proteins"

### Supplementary Table S1. The closest UniProt blast search hits for 2019-nCoV proteins and their comparison with the three novel isolates of BatCoV (2013, 2015, and 2017)

| 2019-nCov protein | UniProt ID | Virus | Seq ID, % | 2017-BatCoV Seq ID, % | 2013 BatCoV Seq ID, % | NCBI ID 2017 | NCBI ID 2015 | NCBI ID 2013 |
|---|---|---|---|---|---|---|---|---|
| wPRF1ab | A0A166ZL34 | Bat coronavirus, BtCoV | 91.3 | 95.7 | 98.6 | AVP78030.1 | AVP78041.1 | QHR63299.1 |
| | P0C6X7 | Human SARS-CoV | 86.2 | | | | | |
| | A0A0U1WHI4 | BtRf-BetaCoV | 85.8 | | | | | |
| wN | R9QTB4 | Bat coronavirus, BtCoV | 91.0 | 94.3 | 99.1 | AVP78038.1 | AVP78049.1 | QHR63308.1 |
| | P59595 | Human SARS-CoV | 90.5 | | | | | |
| | A0A0U1WHI6 | BtRf-BetaCoV | 89.8 | | | | | |
| wORF3a | Q0Q474 | Bat coronavirus, BtCoV | 74.5 | 91.0 | 97.8 | AVP78032.1 | AVP78043.1 | QHR63301.1 |
| | P59632 | Human SARS-CoV | 72.7 | | | | | |
| | A0A0U1UZ48 | BtRf-BetaCoV | 70.9 | | | | | |
| wE | Q3I5J3 | Bat coronavirus, BtCoV | 94.7 | 100.0 | 100.0 | AVP78033.1 | AVP78044.1 | QHR63302.1 |
| | P59637 | Human SARS-CoV | 94.7 | | | | | |
| | A0A0U1WJY0 | BtRf-BetaCoV | 92.1 | | | | | |
| wM | Q0Q472 | Bat coronavirus, BtCoV | 91.7 | 98.7 | 99.6 | AVP78034.1 | | QHR63303.1 |
| | P59596 | Human SARS-CoV | 90.5 | | | | | |
| | A0A0U1WHH9 | BtRf-BetaCoV | 90.4 | | | | | |
| wORF6 | Q3I5J1 | Bat coronavirus, BtCoV | 68.9 | 93.4 | 100.0 | AVP78035.1 | AVP78046.1 | QHR63304.1 |
| | P59634 | Human SARS-CoV | 68.9 | | | | | |
| | A0A0U1WHI0 | BtRf-BetaCoV | 68.9 | | | | | |
| wORF7a | Q3I5J0 | Bat coronavirus, BtCoV | 88.5 | 88.4 | 97.5 | AVP78036.1 | AVP78047.1 | QHR63305.1 |
| | A0A0U1UZE3 | BtRf-BetaCoV | 86.1 | | | | | |
| | P59635 | Human SARS-CoV | 85.2 | | | | | |
| wORF7b | Q3I5I9 | Bat coronavirus, BtCoV | 85.7 | N/A | 97.7 | | | QHR63306.1 |
| | A0A0U1WHI9 | BtRf-BetaCoV | 85.7 | | | | | |
| | Q7TFA1 | Human SARS-CoV | 85.4 | | | | | |
| wORF8 | Q0Q469 | Bat coronavirus | 57.0 | 94.2 | 95.0 | AVP78037.1 | AVP78048.1 | QHR63307.1 |
| wS | Q3LZX1 | Bat coronavirus, BtCoV | 76.0 | 81.0 | 97.4 | AVP78031.1 | AVP78042.1 | QHR63300.1 |
| | P59594 | Human SARS-CoV | 76.0 | | | | | |
| | A0A0U1WHI6 | BtRf-BetaCoV | 73.7 | | | | | |
| wORF10 | FJ882928.1 | Human SARS-CoV | 84.2 | 97.4 | N/A | MG772933.1 (translated) | | |
| | DQ648857.1 | Bat coronavirus, BtCoV | 81.6 | | | | | |
