## Supplementary material for "Structural genomics and interactomics of 2019 Wuhan novel coronavirus, 2019-nCoV, indicate evolutionary conserved functional regions of viral proteins": Ligand binding site mapping for wN

Ligand BS:

- N~2~,N~2~-DIMETHYL-N~1~-(6-OXO-5,6-DIHYDROPHENANTHRIDIN-2-YL)GLYCINAMIDE (4KXJ-P34, OC43)
- ADENOSINE MONOPHOSPHATE (4LI4-AMP, OC43)
- URIDINE-5'-MONOPHOSPHATE (4LM7-U5P, OC43)
- GUANOSINE-5'-MONOPHOSPHATE (4LM9-5GP, OC43)
- CYTIDINE-5'-MONOPHOSPHATE (4LMC-C5P, OC43)
- 6-chloranyl-7-(2-morpholin-4-ylethylamino)quinoline-5,8-dione (4LMT-CQD,OC43)

```
CLUSTAL O(1.2.4) multiple sequence alignment

YP_009724397                        MSDNGPQ-NQRNAPRITFGGPSDSTGSNQNGERSGARSKQRRPQGLPNNTASWFTALTQH 59
SP|P59595|NCAP_CVHSA                MSDNGPQSNQRSAPRITFGGPTDSTDNNQNGGRNGARPKQRRPQGLPNNTASWFTALTQH 60
TR|Q6UZF2|Q6UZF2_CVHSA              MSDNGPQSNQRSAPRITFGGPTDSTDNNQNGGRNGARPKQRRPQGLPNNTASWFTALTQH 60
TR|Q6JH41|Q6JH41_CVHSA              MSDNGPQSNQRSAPRITFGGPTDSTDNNQNGGRNGARPKQRRPQGLPNNTASWFTALTQH 60
TR|Q6UZE8|Q6UZE8_CVHSA              MSDNGPQSNQRSAPRITFGGPTDSTDNNQNGGRNGARPKQRRPQGLPNNTASWFTALTQH 60
TR|Q692D7|Q692D7_CVHSA              MSDNGPQSNQRSAPRITFGGPTDSTDNNQNGGRNGARPKQRRPQGLPNNTASWFTALTQH 60
TR|R9QTB4|R9QTB4_CVHSA              MSDNGPQ-NQRSAPRITFGGPSDSTDNNQDGGRSGARPKQRRPQGLPNNTASWFTALTQH 59
SP|Q3I5I7|NCAP_BCRP3                MSDNGPQ-NQRSAPRITFGGPTDSTDNNQDGGRSGARPKQRRPQGLPNNTASWFTALTQH 59
TR|A0A0K1YZZ7|A0A0K1YZZ7_CVHSA MSDNGPH-NQRSASRITFGGPTDSTDNNQNGGRNGARPKQRRPQGLPNNTASWFTALTQH 59
TR|R9QTA2|R9QTA2_CVHSA              MSDNGPQQNQRSAPRITFGGPTDSADNNQDGGRSGARPKQRRPQGLPNNTASWFTALTQH 60
SP|Q3LZX4|NCAP_BCHK3                MSDNGPQ-SQRSAPRITFGGPADSNDNNQDGGRSGARPKQRRPQGLPNNTASWFTALTQH 59
TR|A0A0U1WHJ1|A0A0U1WHJ1_CVHSA MSDNGPQ-NQRSAPRITFGGPSDSTDNNQDGGRSGVRPKQRRPQGLPNNTASWFTALTQH 59
SP|Q0Q468|NCAP_BC279                MSDNGPQ-NQRSAPRITFGGPSDSTDNNQDGGRSGARPKQRRPQGLPNNTASWFTALTQH 59
TR|A0A166ZLE5|A0A166ZLE5_9NIDO MSDNGTQ-NQRSASRITFGGPSDSTDNNQDGGRSGARPKQRRPQGLPNNTASWFTALTQH 59
TR|A0A0U1UZD6|A0A0U1UZD6_CVHSA MSDNGTQ-NQRSASRITFGGPSDSTDNNQDGGRSGARPKQRRPQGLPNNTASWFTALTQH 59
                                    ***** : .**.* *******:** ..**:* *.*.* **********************

YP_009724397                        GKEDLKFPRGQGVPINTNSSPDDQIGYYRRATRRIRGGDGKMKDLSPRWYFYYLGTGPEA 119
SP|P59595|NCAP_CVHSA                GKEELRFPRGQGVPINTNSGPDDQIGYYRRATRRVRGGDGKMKELSPRWYFYYLGTGPEA 120
TR|Q6UZF2|Q6UZF2_CVHSA              GKEELRFPRGQGVPINTNSGPDDQIGYYRRATRRVRGGDGKMKELSPRWYFYYLGTGPEA 120
TR|Q6JH41|Q6JH41_CVHSA              GKEELRFPRGQGVPINTNSGPDDQIGYYRRATRRVRGGDGKMKELSPRWYFYYLGTGPEA 120
TR|Q6UZE8|Q6UZE8_CVHSA              GKEELRFPRGQGVPINTNSGPDDQIGYYRRATRRVRGGDGKMKELSPRWYFYYLGTGPEA 120
TR|Q692D7|Q692D7_CVHSA              GKEELRFPRGQGVPINTNSGPDDQIGFYRRATRRVRGGDGKMKELSPRWYFYYLGTGPEA 120
TR|R9QTB4|R9QTB4_CVHSA              GKEELRFPRGQGVPINTNSGKDDQIGYYRRATRRVRGGDGKMKELSPRWYFYYLGTGPEA 119
SP|Q3I5I7|NCAP_BCRP3                GKEELRFPRGQGVPINTNSGKDDQIGYYRRATRRVRGGDGKMKELSPRWYFYYLGTGPEA 119
TR|A0A0K1YZZ7|A0A0K1YZZ7_CVHSA GKEELRFPQGQGVPINTNSGPDDQIGYYRRATRRVRGGDGKMKELSPRWYFYYLGTGPEA 119
TR|R9QTA2|R9QTA2_CVHSA              GKEELRFPRGQGVPINTNSGKDDQIGYYRRATRRVRGGDGKMKELSPRWYFYYLGTGPEA 120
SP|Q3LZX4|NCAP_BCHK3                GKEELRFPRGQGVPINTNSGKDDQIGYYRRATRRVRGGDGKMKELSPRWYFYYLGTGPEA 119
TR|A0A0U1WHJ1|A0A0U1WHJ1_CVHSA GKEGLKFPQGQGVPINTNSGRDDQIGYYRRATRRVRGGDGKMKELSPRWYFYYLGTGPEA 119
SP|Q0Q468|NCAP_BC279                GKEELRFPRGQGVPINTNSGKDDQIGYYRRATRRVRGGDGKMKKLSPRWYFYYLGTGPEA 119
TR|A0A166ZLE5|A0A166ZLE5_9NIDO GKEGLKFPQGQGVPINTNSGTDDQIGYYRRATRRVRGGDGKMKELSPRWYFYYLGTGPEA 119
TR|A0A0U1UZD6|A0A0U1UZD6_CVHSA GKEGLKFPQGQGVPINTNSGRDDQIGYYRRATRRVRGGDGKMKELSPRWYFYYLGTGPEA 119
                                    *** *:**:**********. *****:*******:********.****************
```

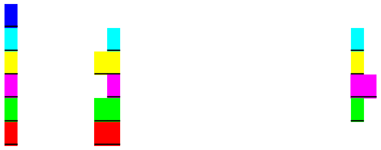

```
YP_009724397                          GLPYGANKDGIIWVATEGALNTPKDHIGTRNPANNAAIVLQLPQGTTLPKGFYAEGSRGG 179
SP|P59595|NCAP_CVHSA                  SLPYGANKEGIVWVATEGALNTPKDHIGTRNPNNNAATVLQLPQGTTLPKGFYAEGSRGG 180
TR|Q6UZF2|Q6UZF2_CVHSA                SLPYGANKEGIVWVATEGALNTPKDHIGTRNPNNNAATVLQLPQGTTLPKGFYAEGSRGG 180
TR|Q6JH41|Q6JH41_CVHSA                SLPYGANKEGIVWVATEGALNTPKDHIGTRNPNNNAATVLQLPQGTTLPKGFYAEGSRGG 180
TR|Q6UZE8|Q6UZE8_CVHSA                SLPYGANKEGIVWVATEGALNTPKDHIGTRNPNNNAATVLQLPQGTTLPKGFYAEGSRGG 180
TR|Q692D7|Q692D7_CVHSA                SLPYGANKEGIVWVATEGALNTPKDHIGTRNPNNNAATVLQLPQGTTLPKGFYAEGSRGG 180
TR|R9QTB4|R9QTB4_CVHSA                SLPYGANKEGIIWVATEGALNTPKDHIGTRNPNNNAAIVLQLPQGTTLPKGFYAEGSRGG 179
SP|Q3I5I7|NCAP_BCRP3                  SLPYGANKEGIVWVATEGALNTPKDHIGTRNPNNNAAIVLQLPQGTTLPKGFYAEGSRGG 179
TR|A0A0K1YZZ7|A0A0K1YZZ7_CVHSA SLPYGANKEGIIWVATEGALNTPKDHIGTRNPNNNAATVLQLPQGTTLPKGFYAEGSRGG 179
TR|R9QTA2|R9QTA2_CVHSA                SLPYGANKEGIVWVAIEGALNTPKDHIGTRNPNNNAAIVLQLPQGTTLPKGFYAEGSRGG 180
SP|Q3LZX4|NCAP_BCHK3                  SLPYGANKEGIVWVATEGALNTPKDHIGTRNPNNNAAIVLQLPQGTTLPKGFYAEGSRGG 179
TR|A0A0U1WHJ1|A0A0U1WHJ1_CVHSA SLPYGANKEGIVWVATEGALNTPKDHIGTRNPNNNAAIVLQLPQGTTLPKGFYAEGSRNG 179
SP|Q0Q468|NCAP_BC279                  SLPYGANKEGIVWVATEGALNTPKDHIGTRNPNNNAAIVLQLPQGTTLPKGFYAEGSRGG 179
TR|A0A166ZLE5|A0A166ZLE5_9NIDO SLPYGANKEGIVWVATEGALNTPKDHIGTRNPNNNAAIVLQLPQGTTLPKGFYAEGSRSG 179
TR|A0A0U1UZD6|A0A0U1UZD6_CVHSA SLPYGANKEGIVWVATEGALNTPKDHIGTRNPNNNAAIVLQLPQGTTLPKGFYAEGSRSG 179
                                      .*******:**:*** **************** **** ********************.*

YP_009724397                          SQASSRSSSRSRNSSRNSTPGSSRGTSPARMAGNGGDAALALLLLDRLNQLESKMSGKGQ 239
SP|P59595|NCAP_CVHSA                  SQASSRSSSRSRGNSRNSTPGSSRGNSPARMASGGGETALALLLLDRLNQLESKVSGKGQ 240
TR|Q6UZF2|Q6UZF2_CVHSA                SQASSRSSSRSRGNSRNSTPGSSRGNSPARMASGGGETALALLLLDRLNQLESKVSGKGQ 240
TR|Q6JH41|Q6JH41_CVHSA                SQASSRSSSRSRGNSRNSTPGSSRGNSPARMASGGGETALALLLLDRLNQLESKVSGKGQ 240
TR|Q6UZE8|Q6UZE8_CVHSA                SQASSRSSSRSRGNSRNSTPGSSRGNSPARMASGGGETALALLLLDRLNQLESKVSGKGQ 240
TR|Q692D7|Q692D7_CVHSA                SQASSRSSSRSRGNSRNSTPGSSRGNSPARMASGGGETALALLLLDRLNQLESKVSGKGQ 240
TR|R9QTB4|R9QTB4_CVHSA                SQASSRSSSRSRGNSRNSTPGSSRGNSPARMASGGGETALALLLLDRLNQLESKVSGKGQ 239
SP|Q3I5I7|NCAP_BCRP3                  SQASSRSSSRSRGNSRNSTPGSSRGNSPARMASGGGETALALLLLDRLNQLESKVSGRSQ 239
TR|A0A0K1YZZ7|A0A0K1YZZ7_CVHSA SQASSRSSSRSRGNSRNSTPGSSRGTSPARIASGGGETALALLLLDRLNQLESKVSGKGQ 239
TR|R9QTA2|R9QTA2_CVHSA                SQASSRSSSRSRGNSRNSTPGSSRGNSPARMASGGGETALALLLLDRLNQLESKVSGKGQ 240
SP|Q3LZX4|NCAP_BCHK3                  SQSSSRSSSRSRGNSRNSTPGSSRGSSPARLASGGGETALALLLLDRLNQLESKVSGKGQ 239
TR|A0A0U1WHJ1|A0A0U1WHJ1_CVHSA SQASSRSSSRSRGNSRTSTPGSSRGNSPARVASGGGETALALLLLDRLNQLESKVSGKGQ 239
SP|Q0Q468|NCAP_BC279                  SQASSRSSSRSRGNSRNSTPGSSRGNSPARMASGSGETALALLLLDRLNQLESKVSGKGQ 239
TR|A0A166ZLE5|A0A166ZLE5_9NIDO SQASSRSSSRSRGNSRTSTPGSSRGNSPARVASGGGETALALLLLDRLNQLESKVSGKGQ 239
TR|A0A0U1UZD6|A0A0U1UZD6_CVHSA SQASSRSSSRSRGNSRISTPGSSRGNSPARVASGGGETALALLLLDRLNQLESKVSGKGQ 239
                                      **:*********..** ********.****:*...*::*****************:**:.*

YP_009724397                          QQQGQTVTKKSAAEASKKPRQKRTATKAYNVTQAFGRRGPEQTQGNFGDQELIRQGTDYK 299
SP|P59595|NCAP_CVHSA                  QQQGQTVTKKSAAEASKKPRQKRTATKQYNVTQAFGRRGPEQTQGNFGDQDLIRQGTDYK 300
TR|Q6UZF2|Q6UZF2_CVHSA                QQQGQTVTKKSAAEASKKPRQKRTATKQYNVTQAFGRRGPEQTQGNFGDQDLIRQGTDYK 300
TR|Q6JH41|Q6JH41_CVHSA                QQQGQTVTKKSAAEASKKPRQKRTATKQYNVTQAFGRRGPEQTQGNFGDQDLIRQGTDYK 300
TR|Q6UZE8|Q6UZE8_CVHSA                QQQGQTVTKKSAAEASKKPRQKRTATKQYNVTQAFGRRGPEQTQGNFGDQDLIRQGTDYK 300
TR|Q692D7|Q692D7_CVHSA                QQQGQTVTKKSAAEASKKPRQKRTATKQYNVTQAFGRRGPEQTQGNFGDQDLIRQGTDYK 300
TR|R9QTB4|R9QTB4_CVHSA                QQQGQTVTKKSAAEASKKPRQKRTATKSYNVTQAFGRRGPEQTQGNFGDQDLIRQGTDYK 299
SP|Q3I5I7|NCAP_BCRP3                  QQQGQTVTKKSAAEASKKPRQKRTATKQYNVTQAFGRRGPEQTQGNFGDQELIRQGTDYK 299
TR|A0A0K1YZZ7|A0A0K1YZZ7_CVHSA QQQGQTVTKKSAAEASKKPRQKRTATKQYNVTQAFGRRGPDQTQGNFGDQDLIRQGTDYK 299
TR|R9QTA2|R9QTA2_CVHSA                QQQGQTVTKKSAAEASKKPRQKRTATKQYNVTQAFGRRGPEQTQGNFGDQELIRQGTDYK 300
SP|Q3LZX4|NCAP_BCHK3                  QQPGQTVTKKSAAEASKKPRQKRTATKQYNVTQAFGRRGPEQTQGNFGDQELIRQGIDYK 299
TR|A0A0U1WHJ1|A0A0U1WHJ1_CVHSA QQQGQTVTKKSASEASKKPRQKRTATKQYNVTQAFGRRGPDQTQGNFGDQELIRQGTDYK 299
SP|Q0Q468|NCAP_BC279                  QQQGQTVTKKSAAEASKKPRQKRTATKSYNVTQAFGRRGPEQTQGNFGDQDLIRQGTDYK 299
TR|A0A166ZLE5|A0A166ZLE5_9NIDO QQQGQTVTKKSASEASKKPRQKRTATKQYNVTQAFGRRGPDQTQGNFGDQELIRQGTDYK 299
TR|A0A0U1UZD6|A0A0U1UZD6_CVHSA QQQGQTVTKKSASEASKKPRQKRTATKQYNVTQAFGRRGPDQTQGNFGDQELIRQGTDYK 299
                                      ** *********:************** ************:*********:***** ***

YP_009724397                          HWPQIAQFAPSASAFFGMSRIGMEVTPSGTWLTYTGAIKLDDKDPNFKDQVILLNKHIDA 359
SP|P59595|NCAP_CVHSA                  HWPQIAQFAPSASAFFGMSRIGMEVTPSGTWLTYHGAIKLDDKDPQFKDNVILLNKHIDA 360
TR|Q6UZF2|Q6UZF2_CVHSA                HWPQIAQFAPSASAFFGMSRIGMEVTPSGTWLTYHGAIKLDDKDPQFKDNVILLNKHIDA 360
TR|Q6JH41|Q6JH41_CVHSA                HWPQIAQFAPSASAFFGMSRIGMEVTPSGTWLTYHGAIKLDDKDPQFKDNVILLNKHIDA 360
TR|Q6UZE8|Q6UZE8_CVHSA                HWPQIAQFAPSASAFFGMSRIGMEVTPSGTWLTYHGAIKLDDKDPQFKDNVILLNKHIDA 360
TR|Q692D7|Q692D7_CVHSA                HWPQIAQFAPSASAFFGMSRIGMEVTPSGTWLTYHGAIKLDDKDPQFKDNVILLNKHIDA 360
TR|R9QTB4|R9QTB4_CVHSA                HWPQIAQFAPSASAFFGMSRIGMEVTPSGTWLTYHGAIKLDDKDPQFKDNVILLNKHIDA 359
SP|Q3I5I7|NCAP_BCRP3                  HWPQIAQFAPSASAFFGMSRIGMEVTPSGTWLTYHGAIKLDDKDPQFKDNVILLNKHIDA 359
TR|A0A0K1YZZ7|A0A0K1YZZ7_CVHSA HWPQIAQFAPSASAFFGMSRIGMEVTPSGTWLTYHGAIKLDDKDPQFKDNVILLNKHIDA 359
TR|R9QTA2|R9QTA2_CVHSA                HWPQIAQFAPSASAFFGMSRIGMEVTPSGTWLTYHGAIKLDDKDPQFKDNVILLNKHIDA 360
SP|Q3LZX4|NCAP_BCHK3                  HWPQIAQFAPSASAFFGMSRIGMEVTPSGTWLTYHGAIKLDDKDPQFKDNVILLNKHIDA 359
TR|A0A0U1WHJ1|A0A0U1WHJ1_CVHSA HWPQIAQFAPSASAFFGMSRIGMEVTPSGTWLTYHGAIKLDDKDPQFKDNVILLNKHIDA 359
SP|Q0Q468|NCAP_BC279                  YWPQIAQFAPSASAFFGMSRIGMEVTPLGTWLTYHGAIKLDDKDPQFKDNVILLNKHIDA 359
TR|A0A166ZLE5|A0A166ZLE5_9NIDO HWPQIAQFAPSASAFFGMSRIGMEVTPSGTWLTYHGAIKLDDKDPQFKDNVILLNKHIDA 359
TR|A0A0U1UZD6|A0A0U1UZD6_CVHSA HWPQIAQFAPSASAFFGMSRIGMEVTPSGTWLTYHGAIKLDDKDPQFKDNVILLNKHIDA 359
                                      :************************ ****** *********:***:**********
```

```
YP_009724397                       YKTFPPTEPKKDKKKKADETQALPQRQKKQQTVTLLPAADLDDFSKQLQQSMSSADSTQA 419
SP|P59595|NCAP_CVHSA               YKTFPPTEPKKDKKKKTDEAQPLPQRQKKQPTVTLLPAADMDDFSRQLQNSMSGASADST 420
TR|Q6UZF2|Q6UZF2_CVHSA             YKTFPPTEPKKDKKKKTDEAQPLPQRQKKQPTVTLLPAADMDDFSRQLQNSMSGASADST 420
TR|Q6JH41|Q6JH41_CVHSA             YKTFPPTEPKKDKKKKTDEAQPLPQRQKKQPTVTLLPAADMDDFSRQLQNSMSGASADST 420
TR|Q6UZE8|Q6UZE8_CVHSA             YKTFPPTEPKKDKKKKTDEAQPLPQRQKKQPTVTLLPAADMDDFSRQLQNSMSGASADST 420
TR|Q692D7|Q692D7_CVHSA             YKTFPPTEPKKDKKKKTDEAQPLPQRQKKQPTVTLLPAADMDDFSRQLQNSMSGASADST 420
TR|R9QTB4|R9QTB4_CVHSA             YKTFPPTEPKKDKKKKTDEAQPLPQRK-KQPTVTLLPAADMDDFSRQLQNSMSGASADST 418
SP|Q3I5I7|NCAP_BCRP3               YKIFPPTEPKKDKKKKTDEAQPLPQRQKKQPTVTLLPAADMDDFSRQLQNSMSGASADST 419
TR|A0A0K1YZZ7|A0A0K1YZZ7_CVHSA     YKTFPPTEPKKDKKKKTDEAQPLPQRQKKQPTVTLLPAADMDDFSRQLQNSMSGASADST 419
TR|R9QTA2|R9QTA2_CVHSA             YKTFPPTEPKKDKKKKTDEAQPLPQRQKKQPTVTLLPAADMDDFSRQLQNSMSGASADST 420
SP|Q3LZX4|NCAP_BCHK3               YKTFPPTEPKKDKKKKTDEAQPLPQRQKKQPTVTLLPAADMDDFSRQLQHSMSGASADST 419
TR|A0A0U1WHJ1|A0A0U1WHJ1_CVHSA     YKTFPPTEPKKDKKKKTDEAQPLPQRQKKQPTVTLLPAADMDDFSRQLQNSMSGASADST 419
SP|Q0Q468|NCAP_BC279               YKAFPPTEPKKDKKKKTDEAQPLPQRK-KQPTVTLLPAADMDDFSRQLQNSMSGASADST 418
TR|A0A166ZLE5|A0A166ZLE5_9NIDO     YKTFPPTEPKKDKKKKTDEAQPLPQRK-KQPTVTLLPAADMDDFSRQLQNSMSGASADST 418
TR|A0A0U1UZD6|A0A0U1UZD6_CVHSA     YKTFPPTEPKKDKKKKTDEAQPLPQRK-KQPTVTLLPAADMDDFSRQLQNSMSGASADST 418
                                   ** *************:**:* ****: ** *********:****:***:***.*.: .:

YP_009724397                       --
SP|P59595|NCAP_CVHSA               QA 422
TR|Q6UZF2|Q6UZF2_CVHSA             QA 422
TR|Q6JH41|Q6JH41_CVHSA             QA 422
TR|Q6UZE8|Q6UZE8_CVHSA             QA 422
TR|Q692D7|Q692D7_CVHSA             QA 422
TR|R9QTB4|R9QTB4_CVHSA             QA 420
SP|Q3I5I7|NCAP_BCRP3               QA 421
TR|A0A0K1YZZ7|A0A0K1YZZ7_CVHSA     QA 421
TR|R9QTA2|R9QTA2_CVHSA             QA 422
SP|Q3LZX4|NCAP_BCHK3               QA 421
TR|A0A0U1WHJ1|A0A0U1WHJ1_CVHSA     QA 421
SP|Q0Q468|NCAP_BC279               QA 420
TR|A0A166ZLE5|A0A166ZLE5_9NIDO     QA 420
TR|A0A0U1UZD6|A0A0U1UZD6_CVHSA     QA 420
```
