## Supplementary material for "Structural genomics and interactomics of 2019 Wuhan novel coronavirus, 2019-nCoV, indicate evolutionary conserved functional regions of viral proteins": Ligand binding site mapping for wNsp3

5-amino-2-methyl-N-[(1R)-1-naphthalen-1-ylethyl]benzamide (3E9S-TTT, SARS)

N-[(4-fluorophenyl)methyl]-1-[(1R)-1-naphthalen-1-ylethyl]piperidine-4-carboxamide (4OVZ-P85, SARS)

N-[(3-fluorophenyl)methyl]-1-[(1R)-1-naphthalen-1-ylethyl]piperidine-4-carboxamide (4OW0-S88, SARS)

```
QHN73794                      GEVITFDNLKTLLSLREVRTIKVFTTVDNINLHTQVVDMSMTYGQQFGPTYLDGADVTKI 1607
SP|P0C6X7|R1AB_CVHSA          GEVLSLDKLKSLLSLREVKTIKVFTTVDNTNLHTQLVDMSMTYGQQFGPTYLDGADVTKI 1584
TR|Q6UZF5|Q6UZF5_CVHSA        GEVLSLDKLKSLLSLREVKTIKVFTTVDNTNLHTQLVDMSMTYGQQFGPTYLDGADVTKI 1584
TR|Q6UZF1|Q6UZF1_CVHSA        GEVLSLDKLKSLLSLREVKTIKVFTTVDNTNLHTQLVDMSMTYGQQFGPTYLDGADVTKI 1584
TR|Q6JH48|Q6JH48_CVHSA        GEVLSLDKLKSLLSLREVKTIKVFTTVDNTNLHTQLVDMSMTYGQQFGPTYLDGADVTKI 1584
TR|Q692E6|Q692E6_CVHSA        GEVLSLDKLKSLLSLREVKTIKVFTTVDNTNLHTQLVDMSMTYGQQFGPTYLDGADVTKI 1584
TR|A0A0K1YZY7|A0A0K1YZY7_CVHSA GEVQPLDKLKSLLSLREVKTIKVFTTVDNTNLHTQIVDMSMTYGQQFGPTYLDGADVTKI 1584
SP|P0C6W2|R1AB_BCHK3          GEVLPLDKLKSLLSLREVKTIKVFTTVDNTNLHTHIVDMSMTYGQQFGPTYLDGADVTKI 1578
SP|P0C6W6|R1AB_BCRP3          GEVLPLDKLKSLLSLREVKTIKVFTTVDNTNLHTQLVDMSMTYGQQLGPTYLEGADVTKI 1582
SP|P0C6V9|R1AB_BC279          GEVLLLDKLKSLLSLREVRTIKVFTTVDNTNLHTQIVDMSMTYGQQFGPTYLDGADVTKI 1590
TR|A0A0U1WHI4|A0A0U1WHI4_CVHSA GEVLPLDKLKSLLSLREVKTIKVFTTVDNTNLHTQLVDMSVTYGQHFGPTYLDGADVTKV 1579
TR|A0A0U1WHG0|A0A0U1WHG0_CVHSA GEVLPLDKLKSLLSLREVKTIKVFTTVDNTNLHTQVVDMSMTYGQHFGSTYLDGADVTKV 1579
TR|A0A166ZL34|A0A166ZL34_9NIDO GEVLPLDKLKSLLSLREVKTIKVFTTVDNTNLHTQVVDMSMTYGQHFGSTYLDGADVTKV 1579
TR|R9QTB2|R9QTB2_CVHSA        GEVLPLDKLKSLLSLREVKTIKVFTTVDNTNLHTQIVDMSMTYGQQFGSTYLDGADVTKI 1576
TR|R9QTH2|R9QTH2_CVHSA        GEVLPLDKLKSLLSLREVKTIKVFTTVDNTNLHTQLVDMSMTYGQQFGPTYLDGADVTKI 1585
SP|P0C6U8|R1A_CVHSA           GEVLSLDKLKSLLSLREVKTIKVFTTVDNTNLHTQLVDMSMTYGQQFGPTYLDGADVTKI 1584
TR|Q6JH47|Q6JH47_CVHSA        GEVLSLDKLKSLLSLREVKTIKVFTTVDNTNLHTQLVDMSMTYGQQFGPTYLDGADVTKI 1584
TR|Q692E5|Q692E5_CVHSA        GEVLSLDKLKSLLSLREVKTIKVFTTVDNTNLHTQLVDMSMTYGQQFGPTYLDGADVTKI 1584
SP|P0C6F8|R1A_BCHK3           GEVLPLDKLKSLLSLREVKTIKVFTTVDNTNLHTHIVDMSMTYGQQFGPTYLDGADVTKI 1578
TR|A0A0K1Z0N1|A0A0K1Z0N1_CVHSA GEVQPLDKLKSLLSLREVKTIKVFTTVDNTNLHTQIVDMSMTYGQQFGPTYLDGADVTKI 1584
SP|P0C6F5|R1A_BC279           GEVLLLDKLKSLLSLREVRTIKVFTTVDNTNLHTQIVDMSMTYGQQFGPTYLDGADVTKI 1590
SP|P0C6T7|R1A_BCRP3           GEVLPLDKLKSLLSLREVKTIKVFTTVDNTNLHTQLVDMSMTYGQQLGPTYLEGADVTKI 1582
                              ***  :*:**:*******:********** ****::****:****::* ***:******:


QHN73794                      KPHNSHEGKTFYVLPNDDTLRVEAFEYYHTTDPSFLGRYMSALNHTKKWKYPQVNGLTSI 1667
SP|P0C6X7|R1AB_CVHSA          KPHVNHEGKTFFVLPSDDTLRSEAFEYYHTLDESFLGRYMSALNHTKKWKFPQVGGLTSI 1644
TR|Q6UZF5|Q6UZF5_CVHSA        KPHVNHEGKTFFVLPSDDTLRSEAFEYYHTLDESFLGRYMSALNHTKKWKFPQVGGLTSI 1644
TR|Q6UZF1|Q6UZF1_CVHSA        KPHVNHEGKTFFVLPSDDTLRSEAFEYYHTLDESFLGRYMSALNHTKKWKFPQVGGLTSI 1644
TR|Q6JH48|Q6JH48_CVHSA        KPHVNHEGKTFFVLPSDDTLRSEAFEYYHTLDESFLGRYMSALNHTKKWKFPQVGGLTSI 1644
TR|Q692E6|Q692E6_CVHSA        KPHVNHEGKTFFVLPSDDTLRSEAFEYYHTLDESFLGRYMSALNHTKKWKFPQVGGLTSI 1644
TR|A0A0K1YZY7|A0A0K1YZY7_CVHSA KPHVNHEGKTFYVLPSDDTLRSEAFEYYHTLDESFLGRYMSALNHTKKWKFPQVGGLTSI 1644
SP|P0C6W2|R1AB_BCHK3          KPHVNHEGKTFFVLPSDDTLRSEAFEYYHTIDESFLGRYMSALNHTKKWKFPQVGGLTSI 1638
SP|P0C6W6|R1AB_BCRP3          KPHVNHEGKTFFVLPSDDTLRSEAFEYYHTLDESFLGRYMSALNHTKKWKFPQVGGLTSI 1642
SP|P0C6V9|R1AB_BC279          KPHAKHEGKTFFVLPSDDTLRSEAFEYYHTLDESFLGRYMSALNHTKKWKFPQIGGLTSI 1650
TR|A0A0U1WHI4|A0A0U1WHI4_CVHSA KPHVNHEGKTFFVLPSDDTLRSEAFEYYHTLDESFLGRYMSALSHTKKWKFPQVGGLTSI 1639
TR|A0A0U1WHG0|A0A0U1WHG0_CVHSA KPHVNHEGKTFFVLPSDDTLRSEAFEYYHTLDESFLGRYMSALNHTKKWKFPQVGGLTSI 1639
TR|A0A166ZL34|A0A166ZL34_9NIDO KPHVNHEGKTFFVLPSDDTLRSEAFGYYHTLDESFLGRYMSALNHTKKWKFPQVGGLTSI 1639
TR|R9QTB2|R9QTB2_CVHSA        KPHVNHEGKTFFVLPSDDTLRSEAFEYYHTLDESFLGRYMSALSHTKKWKFPQVGGLTSI 1636
TR|R9QTH2|R9QTH2_CVHSA        KPHVNHEGKTFFVLPSDDTLRSEAFEYYHTLDESFLGRYMSALNHTKKWKFPQVGGLTSI 1645
SP|P0C6U8|R1A_CVHSA           KPHVNHEGKTFFVLPSDDTLRSEAFEYYHTLDESFLGRYMSALNHTKKWKFPQVGGLTSI 1644
TR|Q6JH47|Q6JH47_CVHSA        KPHVNHEGKTFFVLPSDDTLRSEAFEYYHTLDESFLGRYMSALNHTKKWKFPQVGGLTSI 1644
TR|Q692E5|Q692E5_CVHSA        KPHVNHEGKTFFVLPSDDTLRSEAFEYYHTLDESFLGRYMSALNHTKKWKFPQVGGLTSI 1644
SP|P0C6F8|R1A_BCHK3           KPHVNHEGKTFFVLPSDDTLRSEAFEYYHTIDESFLGRYMSALNHTKKWKFPQVGGLTSI 1638
TR|A0A0K1Z0N1|A0A0K1Z0N1_CVHSA KPHVNHEGKTFYVLPSDDTLRSEAFEYYHTLDESFLGRYMSALNHTKKWKFPQVGGLTSI 1644
SP|P0C6F5|R1A_BC279           KPHAKHEGKTFFVLPSDDTLRSEAFEYYHTLDESFLGRYMSALNHTKKWKFPQIGGLTSI 1650
SP|P0C6T7|R1A_BCRP3           KPHVNHEGKTFFVLPSDDTLRSEAFEYYHTLDESFLGRYMSALNHTKKWKFPQVGGLTSI 1642
                              *** .******:***.***** *** **** * **********.******:**:.*****


QHN73794                      KWADNNCYLATALLTLQQIELKFNPPALQDAYYRARAGEAANFCALILAYCNKTVGELGD 1727
SP|P0C6X7|R1AB_CVHSA          KWADNNCYLSSVLLALQQLEVKFNAPALQEAYYRARAGDAANFCALILAYSNKTVGELGD 1704
TR|Q6UZF5|Q6UZF5_CVHSA        KWADNNCYLSSVLLALQQLEVKFNAPALQEAYYRARAGDAANFCALILAYSNKTVGELGD 1704
TR|Q6UZF1|Q6UZF1_CVHSA        KWADNNCYLSSVLLALQQLEVKFNAPALQEAYYRARAGDAANFCALILAYSNKTVGELGD 1704
TR|Q6JH48|Q6JH48_CVHSA        KWADNNCYLSSVLLALQQLEVKFNAPALQEAYYRARAGDAANFCALILAYSNKTVGELGD 1704
TR|Q692E6|Q692E6_CVHSA        KWADNNCYLSSVLLALQQLEVKFNAPALQEAYYRARAGDAANFCALILAYSNKTVGELGD 1704
TR|A0A0K1YZY7|A0A0K1YZY7_CVHSA KWADNNCYLSSVLLALQQIEVKFNAPALQEAYYRARAGDAANFCALILAYSNKTVGELGD 1704
SP|P0C6W2|R1AB_BCHK3          KWADNNCYLSSVLLALQQVEVKFNAPALQEAYYRARAGDAANFCALILAYSNKTVGELGD 1698
SP|P0C6W6|R1AB_BCRP3          KWADNNCYLSSVLLALQQIEVKFNAPALQEAYYRARAGDAANFCALILAYSNKTVGELGD 1702
SP|P0C6V9|R1AB_BC279          KWADNNCYLSSVLLALQQIEVKFNAPALQEAYYRARAGDAANFCALILAYSNRTVGELGD 1710
TR|A0A0U1WHI4|A0A0U1WHI4_CVHSA KWADNNCYLSSVLLALQQIEVKFNAPALQEAYYRARAGDAANFCALILAYSNKTVGELGD 1699
TR|A0A0U1WHG0|A0A0U1WHG0_CVHSA KWADNNCYLSSVLLALQQIEVKFNAPALQEAYYRARAGEAANFCALILAYSNKTVGELGD 1699
TR|A0A166ZL34|A0A166ZL34_9NIDO KWADNNCYLSSVLLALQQIEVKFNAPALQEAYYRARAGDAANFCALTLAYSNKTVGDLGD 1699
TR|R9QTB2|R9QTB2_CVHSA        KWADNNCYLSSVLLALQQIEVKFNAPALQEAYYRARAGDAANFCALILAYSYKTVGELGD 1696
TR|R9QTH2|R9QTH2_CVHSA        KWADNNCYLSSVLLALQQIEVKFNAPALQEAYYRARAGDAANFCALILAYSNKTVGELGD 1705
SP|P0C6U8|R1A_CVHSA           KWADNNCYLSSVLLALQQLEVKFNAPALQEAYYRARAGDAANFCALILAYSNKTVGELGD 1704
TR|Q6JH47|Q6JH47_CVHSA        KWADNNCYLSSVLLALQQLEVKFNAPALQEAYYRARAGDAANFCALILAYSNKTVGELGD 1704
TR|Q692E5|Q692E5_CVHSA        KWADNNCYLSSVLLALQQLEVKFNAPALQEAYYRARAGDAANFCALILAYSNKTVGELGD 1704
SP|P0C6F8|R1A_BCHK3           KWADNNCYLSSVLLALQQVEVKFNAPALQEAYYRARAGDAANFCALILAYSNKTVGELGD 1698
TR|A0A0K1Z0N1|A0A0K1Z0N1_CVHSA KWADNNCYLSSVLLALQQIEVKFNAPALQEAYYRARAGDAANFCALILAYSNKTVGELGD 1704
SP|P0C6F5|R1A_BC279           KWADNNCYLSSVLLALQQIEVKFNAPALQEAYYRARAGDAANFCALILAYSNRTVGELGD 1710
SP|P0C6T7|R1A_BCRP3           KWADNNCYLSSVLLALQQIEVKFNAPALQEAYYRARAGDAANFCALILAYSNKTVGELGD 1702
                              *********::.**:***:*:*** ****:********:******* ***. :***:***
```

```
QHN73794                           VRETMSYLFQHANLDSCKRVLNVVCKTCGQQQTTLKGVEAVMYMGTLSYEQFKKGVQIPC 1787
SP|P0C6X7|R1AB_CVHSA               VRETMTHLLQHANLESAKRVLNVVCKHCGQKTTTLTGVEAVMYMGTLSYDNLKTGVSIPC 1764
TR|Q6UZF5|Q6UZF5_CVHSA             VRETMTHLLQHANLESAKRVLNVVCKHCGQKTTTLTGVEAVMYMGTLSYDNLKTGVSIPC 1764
TR|Q6UZF1|Q6UZF1_CVHSA             VRETMTHLLQHANLESAKRVLNVVCKHCGQKTTTLTGVEAVMYMGTLSYDNLKTGVSIPC 1764
TR|Q6JH48|Q6JH48_CVHSA             VRETMTHLLQHANLESAKRVLNVVCKHCGQKTTTLTGVEAVMYMGTLSYDNLKTGVSIPC 1764
TR|Q692E6|Q692E6_CVHSA             VRETMTHLLQHANLESAKRVLNVVCKHCGQKTTTLTGVEAVMYMGTLSYDNLKTGVSIPC 1764
TR|A0A0K1YZY7|A0A0K1YZY7_CVHSA     VRETMTHLLQHANLESAKRVLNVVCKHCGQKTTTLTGVEAVMYMGTLSYDNLKTGVSIPC 1764
SP|P0C6W2|R1AB_BCHK3               VRETMTHLLQHANLESAKRVLNVVCKHCGQKTTTLKGVEAVMYMGTLSYDELKTGVSIPC 1758
SP|P0C6W6|R1AB_BCRP3               VRETMTHLLQHANLESAKRVLNVVCKHCGQKTTTLTGVEAVMYMGTLSYDNLKMGVSIPC 1762
SP|P0C6V9|R1AB_BC279               VRETMTHLLQHANLESAKRVLNVVCKTCGQKSTTLTGVEAVMYMGTLSYEELKTGVTIPC 1770
TR|A0A0U1WHI4|A0A0U1WHI4_CVHSA     VRETMTHLLQHANLEFAKRVLNLVCKHCGQKTTTLTGVEAVMYMGTLSYDELKTGVSIPC 1759
TR|A0A0U1WHG0|A0A0U1WHG0_CVHSA     VRETMTHLLQHANLEFAKRVLNLVCKHCGQKTTTLTGVEAVMYMGTLSYDELKTGVSIPC 1759
TR|A0A166ZL34|A0A166ZL34_9NIDO     VRETMTHLLQHANLEFAKRVLNLVCKHCGQKTTTLTGVEAVMYMGTLSYDELKTGVSIPC 1759
TR|R9QTB2|R9QTB2_CVHSA             VRETMTHLLQHANLESAKRVLNVVCKHCGQKTTTLTGVEAVMYMGTLSYDELKTGVSIPC 1756
TR|R9QTH2|R9QTH2_CVHSA             VRETMAHLLQHANLESAKRVLNVVCKHCGQKTTTLTGVEAVMYMGTLSYDNLKTGVSVPC 1765
SP|P0C6U8|R1A_CVHSA                VRETMTHLLQHANLESAKRVLNVVCKHCGQKTTTLTGVEAVMYMGTLSYDNLKTGVSIPC 1764
TR|Q6JH47|Q6JH47_CVHSA             VRETMTHLLQHANLESAKRVLNVVCKHCGQKTTTLTGVEAVMYMGTLSYDNLKTGVSIPC 1764
TR|Q692E5|Q692E5_CVHSA             VRETMTHLLQHANLESAKRVLNVVCKHCGQKTTTLTGVEAVMYMGTLSYDNLKTGVSIPC 1764
SP|P0C6F8|R1A_BCHK3                VRETMTHLLQHANLESAKRVLNVVCKHCGQKTTTLKGVEAVMYMGTLSYDELKTGVSIPC 1758
TR|A0A0K1Z0N1|A0A0K1Z0N1_CVHSA     VRETMTHLLQHANLESAKRVLNVVCKHCGQKTTTLTGVEAVMYMGTLSYDNLKTGVSIPC 1764
SP|P0C6F5|R1A_BC279                VRETMTHLLQHANLESAKRVLNVVCKTCGQKSTTLTGVEAVMYMGTLSYEELKTGVTIPC 1770
SP|P0C6T7|R1A_BCRP3                VRETMTHLLQHANLESAKRVLNVVCKHCGQKTTTLTGVEAVMYMGTLSYDNLKMGVSIPC 1762
                                   *****::*:*****: .*****:*** ***: ***.**************:::* ** :**

QHN73794                           TCGKQATKYLVQQESPFVMMSAPPAQYELKHGTFTCASEYTGNYQCGHYKHITSKETLYC 1847
SP|P0C6X7|R1AB_CVHSA               VCGRDATQYLVQQESSFVMMSAPPAEYKLQQGTFLCANEYTGNYQCGHYTHITAKETLYR 1824
TR|Q6UZF5|Q6UZF5_CVHSA             VCGRDATQYLVQQESSFVMMSAPPAEYKLQQGTFLCANEYTGNYQCGHYTHITAKETLYR 1824
TR|Q6UZF1|Q6UZF1_CVHSA             VCGRDATQYLVQQESSFVMMSAPPAEYKLQQGTFLCANEYTGNYQCGHYTHITAKETLYR 1824
TR|Q6JH48|Q6JH48_CVHSA             VCGRDATQYLVQQESSFVMMSAPPAEYKLQQGTFLCANEYTGNYQCGHYTHITAKETLYR 1824
TR|Q692E6|Q692E6_CVHSA             VCGRDATQYLVQQESSFVMMSAPPAEYKLQQGTFLCANEYTGNYQCGHYTHITAKETLYR 1824
TR|A0A0K1YZY7|A0A0K1YZY7_CVHSA     VCGRDATQYLIQQESSFVMMSAPPAEYKLQQGTFLCANEYTGNYQCGHYTHVTAKETLYR 1824
SP|P0C6W2|R1AB_BCHK3               VCGRNATQYLVQQESSFVMMSAPPAEYKLQQGAFLCANEYTGNYQCGHYTHITAKETLYR 1818
SP|P0C6W6|R1AB_BCRP3               VCGRDATQYLVQQESSFVMMSAPPAEYKLQQGTFLCANEYTGNYQCGHYTHITAKETLYR 1822
SP|P0C6V9|R1AB_BC279               ICGRDATQYLVQQESSFVMMSAPPSEYTLQQGAFLCANEYTGSYQCGHYTHVTVKETLYR 1830
TR|A0A0U1WHI4|A0A0U1WHI4_CVHSA     VCGRGATQYLVQQESSFVMMSAPPAEYKLQQGTFLCANEYTGNYQCGHYTHITAKETLYR 1819
TR|A0A0U1WHG0|A0A0U1WHG0_CVHSA     VCGRDATQYLVQQESSFVMMSAPPAEYKLQQSTFLCANEYTGNYQCGHYTHITAKETLYR 1819
TR|A0A166ZL34|A0A166ZL34_9NIDO     VCGRDATQYLVQQESSFVMMSAPPAEYKLQQGTFLCANEYTGNYQCGHYTHITAKETLYH 1819
TR|R9QTB2|R9QTB2_CVHSA             VCGRNATQYLVQQESSFVMMSAPPAEYKLQQGTFLCANEYTGNYQCGHYTHITAKETLYR 1816
TR|R9QTH2|R9QTH2_CVHSA             VCGRDATQYLVQQESSFVMMSAPPAEYKLQQGTFLCANEYTGNYQCGHYTHITAKETLYR 1825
SP|P0C6U8|R1A_CVHSA                VCGRDATQYLVQQESSFVMMSAPPAEYKLQQGTFLCANEYTGNYQCGHYTHITAKETLYR 1824
TR|Q6JH47|Q6JH47_CVHSA             VCGRDATQYLVQQESSFVMMSAPPAEYKLQQGTFLCANEYTGNYQCGHYTHITAKETLYR 1824
TR|Q692E5|Q692E5_CVHSA             VCGRDATQYLVQQESSFVMMSAPPAEYKLQQGTFLCANEYTGNYQCGHYTHITAKETLYR 1824
SP|P0C6F8|R1A_BCHK3                VCGRNATQYLVQQESSFVMMSAPPAEYKLQQGAFLCANEYTGNYQCGHYTHITAKETLYR 1818
TR|A0A0K1Z0N1|A0A0K1Z0N1_CVHSA     VCGRDATQYLIQQESSFVMMSAPPAEYKLQQGTFLCANEYTGNYQCGHYTHVTAKETLYR 1824
SP|P0C6F5|R1A_BC279                ICGRDATQYLVQQESSFVMMSAPPSEYTLQQGAFLCANEYTGSYQCGHYTHVTVKETLYR 1830
SP|P0C6T7|R1A_BCRP3                VCGRDATQYLVQQESSFVMMSAPPAEYKLQQGTFLCANEYTGNYQCGHYTHITAKETLYR 1822
                                    **: **:**:**** ********::* *::.:* **.****.******.*:* *****

QHN73794                           IDGALLTKSSEYKGPITDVFYKENSYTTTIKPVTYKLDGVVCTEIDPKLDNYYKKDNSYF 1907
SP|P0C6X7|R1AB_CVHSA               IDGAHLTKMSEYKGPVTDVFYKETSYTTTIKPVSYKLDGVTYTEIEPKLDGYYKKDNAYY 1884
TR|Q6UZF5|Q6UZF5_CVHSA             IDGAHLTKMSEYKGPVTDVFYKETSYTTTIKPVSYKLDGVTYTEIEPKLDGYYKKDNAYY 1884
TR|Q6UZF1|Q6UZF1_CVHSA             IDGAHLTKMSEYKGPVTDVFYKETSYTTTIKPVSYKLDGVTYTEIEPKLDGYYKKDNAYY 1884
TR|Q6JH48|Q6JH48_CVHSA             IDGAHLTKMSEYKGPVTDVFYKETSYTTTIKPVSYKLDGVTYTEIEPKLDGYYKKDNAYY 1884
TR|Q692E6|Q692E6_CVHSA             IDGAHLTKMSEYKGPVTDVFYKETSYTTTIKPVSYKLDGVTYTEIEPKLDGYYKKDNAYY 1884
TR|A0A0K1YZY7|A0A0K1YZY7_CVHSA     IDGAHLTKMSEYKGPVTDVFYKETSYTTTIKPVSYKLDGVTYTEIEPKLDGYYKKDNAYY 1884
SP|P0C6W2|R1AB_BCHK3               VDGAHLTKMSEYKGPVTDVFYKETSYTTAIKPVSYKLDGVTYTEIEPKLDGYYKKGNAYY 1878
SP|P0C6W6|R1AB_BCRP3               IDGAHLTKMSEYKGPVTDVFYKETSYTTTIKPVSYKLDGVTYTEIEPKLDGYYKKDNAYY 1882
SP|P0C6V9|R1AB_BC279               IDGAYLTKMSEYKGPVTDVFYKEISYTTTIKPVSYKLDGVIYTEIQPKLDEYYKKDNAYY 1890
TR|A0A0U1WHI4|A0A0U1WHI4_CVHSA     IDGAHLTKMSEYKGPVTDVFYKETSYTTTIKPVSYKLDGVTYTEIEPKLDGYYKKDNAYY 1879
TR|A0A0U1WHG0|A0A0U1WHG0_CVHSA     IDGAHLTKMSEYKGPVTDVFYKETSYTTTIKPVSYKLDGVTYTEIEPKLDGYYKKDNAYY 1879
TR|A0A166ZL34|A0A166ZL34_9NIDO     IDGAHLTKMSEYKGPVTDVFYKETSYTTTIKPVSYKLDGVTYTEIEPKLDGYYKKDNAY- 1878
TR|R9QTB2|R9QTB2_CVHSA             IDGAHLTKMSEYKGPVTDVFYKETSYTTTIKPVSYKLDGVTYTEIEPKLDGYYKKDNAYY 1876
TR|R9QTH2|R9QTH2_CVHSA             IDGAHLTKMSEYKGPVTDVFYKETSYTTTIKPVSYKLDGVTYTEIEPKLDGYYKKDNAYY 1885
SP|P0C6U8|R1A_CVHSA                IDGAHLTKMSEYKGPVTDVFYKETSYTTTIKPVSYKLDGVTYTEIEPKLDGYYKKDNAYY 1884
TR|Q6JH47|Q6JH47_CVHSA             IDGAHLTKMSEYKGPVTDVFYKETSYTTTIKPVSYKLDGVTYTEIEPKLDGYYKKDNAYY 1884
TR|Q692E5|Q692E5_CVHSA             IDGAHLTKMSEYKGPVTDVFYKETSYTTTIKPVSYKLDGVTYTEIEPKLDGYYKKDNAYY 1884
SP|P0C6F8|R1A_BCHK3                VDGAHLTKMSEYKGPVTDVFYKETSYTTAIKPVSYKLDGVTYTEIEPKLDGYYKKGNAYY 1878
TR|A0A0K1Z0N1|A0A0K1Z0N1_CVHSA     IDGAHLTKMSEYKGPVTDVFYKETSYTTTIKPVSYKLDGVTYTEIEPKLDGYYKKDNAYY 1884
SP|P0C6F5|R1A_BC279                IDGAYLTKMSEYKGPVTDVFYKEISYTTTIKPVSYKLDGVIYTEIQPKLDEYYKKDNAYY 1890
SP|P0C6T7|R1A_BCRP3                IDGAHLTKMSEYKGPVTDVFYKETSYTTTIKPVSYKLDGVTYTEIEPKLDGYYKKDNAYY 1882
                                   :*** *** ******:******* ****:****:******  ***:**** ****.*:*
```
