## Supplementary material for "Structural genomics and interactomics of 2019 Wuhan novel coronavirus, 2019-nCoV, indicate evolutionary conserved functional regions of viral proteins": Ligand binding site mapping for wNsp5 (Non-SARS Group)

Ligand BS:

N-{4-[(1H-benzotriazol-1-ylacetyl)(thiophen-3-ylmethyl)amino]phenyl}thiophene-2-carboxamide (4YOI-4F4, HKU4)

N-[4-(acetylamino)phenyl]-2-(1H-benzotriazol-1-yl)-N-[(1R)-2-(tert-butylamino)-2-oxo-1-(thiophen-3-yl)ethyl]acetamide (4YOG-4F5, HKU4)

ADENOSINE-5-DIPHOSPHORIBOSE (4MEA-APR, HKU4)

N-{4-[(1H-benzotriazol-1-ylacetyl)(thiophen-3-ylmethyl)amino]phenyl}benzamide (4YOJ-RFM, HKU4)

(1R,2S)-2-{[N-({[4-benzyl-1-(tert-butoxycarbonyl)piperidin-4-yl]oxy}carbonyl)-L-leucyl]amino}-1-hydroxy-3-[(3S)-2-oxopyrrolidin-3-yl]propane-1-sulfonic acid (5WKL-AVY, MERS)

(1S,2S)-2-[(N-{[2-(3-chlorophenyl)ethoxy]carbonyl}-L-leucyl)amino]-1-hydroxy-3-[(3S)-2-oxopyrrolidin-3-yl]propane-1-sulfonic acid (5WKK-AW4, MERS)

(1R,2S)-2-({N-[(benzyloxy)carbonyl]-L-leucyl}amino)-1-hydroxy-3-[(3S)-2-oxopyrrolidin-3-yl]propane-1-sulfonic acid (5WKJ-B1S, MERS)

(1R,2S)-2-[(N-{[2-(3-chlorophenyl)ethoxy]carbonyl}-L-leucyl)amino]-1-hydroxy-3-[(3S)-2-oxopyrrolidin-3-yl]propane-1-sulfonic acid (5WKK-B3G, MERS)

(1S,2S)-2-{[N-({[4-benzyl-1-(tert-butoxycarbonyl)piperidin-4-yl]oxy}carbonyl)-L-leucyl]amino}-1-hydroxy-3-[(3S)-2-oxopyrrolidin-3-yl]propane-1-sulfonic acid (5WKL-B3J, MERS)

(1R,2S)-2-{[N-({[1-(tert-butoxycarbonyl)-4-ethylpiperidin-4-yl]oxy}carbonyl)-L-leucyl]amino}-1-hydroxy-3-[(3S)-2-oxopyrrolidin-3-yl]propane-1-sulfonic acid (5WKM-B6Y, MERS)

(1S,2S)-2-({N-[(benzyloxy)carbonyl]-L-leucyl}amino)-1-hydroxy-3-[(3S)-2-oxopyrrolidin-3-yl]propane-1-sulfonic acid (5WKJ-K36, MERS)

(1S,2S)-2-{[N-({[1-(tert-butoxycarbonyl)-4-ethylpiperidin-4-yl]oxy}carbonyl)-L-leucyl]amino}-1-hydroxy-3-[(3S)-2-oxopyrrolidin-3-yl]propane-1-sulfonic acid (5WKM-N02, MERS)

(S)-N-benzyl-3-((S)-2-cinnamamido-3-phenylpropanamido)-2-oxo-4-((S)-2-oxopyrrolidin-3-yl)butanamide (6FV2-D03, NL63)

(2~{S})-4-methyl-~{N}-[(2~{S},3~{R})-3-oxidanyl-4-oxidanylidene-1-[(3~{S})-2-oxidanylidenepyrrolidin-3-yl]-4-[(phenylmethyl)amino]butan-2-yl]-2-[[(~{E})-3-phenylprop-2-enoyl]amino]pentanamide (6FV1-E8E, NL63)

```
QHN73794                        EAACCHLAKALNDFSNSGSDVLYQPPQTSITSAVLQSGFRKMAFPSGKVEGCMVQVTCGT 3287
SP|P0C6X7|R1AB_CVHSA            EAACCHLAKALNDFSNSGADVLYQPPQTSITSAVLQSGFRKMAFPSGKVEGCMVQVTCGT 3264
TR|Q6UZF5|Q6UZF5_CVHSA          EAACCHLAKALNDFSNSGADVLYQPPQTSITSAVLQSGFRKMAFPSGKVEGCMVQVTCGT 3264
TR|Q6UZF1|Q6UZF1_CVHSA          EAACCHLAKALNDFSNSGADVLYQPPQTSITSAVLQSGFRKMAFPSGKVEGCMVQVTCGT 3264
TR|Q6JH48|Q6JH48_CVHSA          EAACCHLAKALNDFSNSGADVLYQPPQTSITSAVLQSGFRKMAFPSGKVEGCMVQVTCGT 3264
TR|Q692E6|Q692E6_CVHSA          EAACCHLAKALNDFSNSGADVLYQPPQTSITSAVLQSGFRKMAFPSGKVEGCMVQVTCGT 3264
TR|A0A0K1YZY7|A0A0K1YZY7_CVHSA  EAACCHLAKALNDFSNSGSDVLYQPPQTSITSAVLQSGFRKMAFPSGKVEGCMVQVTCGT 3264
SP|P0C6W2|R1AB_BCHK3            EAACCHLAKALNDFSNSGADVLYQPPQTSITSAVLQSGFRKMAFPSGKVEGCMVQVTCGT 3258
SP|P0C6W6|R1AB_BCRP3            EAACCHLAKALNDFSNSGADVLYQPPQTSITSAVLQSGFRKMAFPSGKVEGCMVQVTCGT 3262
SP|P0C6V9|R1AB_BC279            EAACCHLAKALNDFSNSGADVLYQPPQTSITSAVLQSGFRKMAFPSGKVEGCMVQVTCGT 3270
TR|A0A0U1WHI4|A0A0U1WHI4_CVHSA  EAACCHLAKALNDFSNSGADVLYQPPQTSITSAVLQSGFRRMAFPSGKVEGCMVQVTCGT 3259
TR|A0A0U1WHG0|A0A0U1WHG0_CVHSA  EAACCHLAKALNDFSNSGADVLYQPPQTSITSAVLQSGFKKVAFPSGKVEGCMVQVTCGT 3259
TR|A0A166ZL34|A0A166ZL34_9NIDO  EAACCHLAKALNDFSNSGADVLYQPPQTSITSAVLQSGFKKMAFPSGKVEGCMVQVTCGT 3066
TR|R9QTB2|R9QTB2_CVHSA          EAACCHLAKALNDFSNSGADVLYQPPQTSITSAVLQSGFRKMAFPSGKVEGCMVQVTCGT 3256
TR|R9QTH2|R9QTH2_CVHSA          EAACCHLAKALNDFSNSGSDVLYQPPQTSITSAVLQSGFRKMAFPSGKVEGCMVQVTCGT 3265
SP|P0C6U8|R1A_CVHSA             EAACCHLAKALNDFSNSGADVLYQPPQTSITSAVLQSGFRKMAFPSGKVEGCMVQVTCGT 3264
TR|Q6JH47|Q6JH47_CVHSA          EAACCHLAKALNDFSNSGADVLYQPPQTSITSAVLQSGFRKMAFPSGKVEGCMVQVTCGT 3264
TR|Q692E5|Q692E5_CVHSA          EAACCHLAKALNDFSNSGADVLYQPPQTSITSAVLQSGFRKMAFPSGKVEGCMVQVTCGT 3264
SP|P0C6F8|R1A_BCHK3             EAACCHLAKALNDFSNSGADVLYQPPQTSITSAVLQSGFRKMAFPSGKVEGCMVQVTCGT 3258
TR|A0A0K1Z0N1|A0A0K1Z0N1_CVHSA  EAACCHLAKALNDFSNSGSDVLYQPPQTSITSAVLQSGFRKMAFPSGKVEGCMVQVTCGT 3264
SP|P0C6F5|R1A_BC279             EAACCHLAKALNDFSNSGADVLYQPPQTSITSAVLQSGFRKMAFPSGKVEGCMVQVTCGT 3270
SP|P0C6T7|R1A_BCRP3             EAACCHLAKALNDFSNSGADVLYQPPQTSITSAVLQSGFRKMAFPSGKVEGCMVQVTCGT 3262
                                ******************:********************:::******************
```

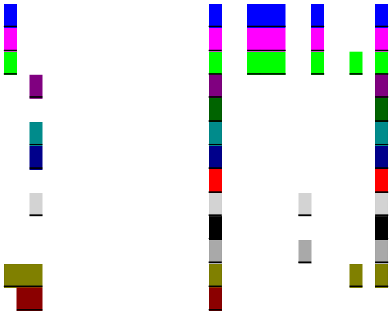

```
QHN73794                          TTLNGLWLDDVVYCPRHVICTSEDMLNPNYEDLLIRKSNHNFLVQAGNVQLRVIGHSMQN 3347
SP|P0C6X7|R1AB_CVHSA              TTLNGLWLDDTVYCPRHVICTAEDMLNPNYEDLLIRKSNHSFLVQAGNVQLRVIGHSMQN 3324
TR|Q6UZF5|Q6UZF5_CVHSA            TTLNGLWLDDTVYCPRHVICTAEDMLNPNYEDLLIRKSNHSFLVQAGNVQLRVIGHSMQN 3324
TR|Q6UZF1|Q6UZF1_CVHSA            TTLNGLWLDDTVYCPRHVICTAEDMLNPNYEDLLIRKSNHSFLVQAGNVQLRVIGHSMQN 3324
TR|Q6JH48|Q6JH48_CVHSA            TTLNGLWLDDTVYCPRHVICTAEDMLNPNYEDLLIRKANHSFLVQAGNVQLRVIGHSMQN 3324
TR|Q692E6|Q692E6_CVHSA            TTLNGLWLDDTVYCPRHVICTAEDMLNPNYEDLLIRKSNHSFLVQAGNVQLRVIGHSMQN 3324
TR|A0A0K1YZY7|A0A0K1YZY7_CVHSA    TTLNGLWLDDTVYCPRHVICTAEDMLNPNYEDLLIRKSNHSFLVQAGNVQLRVIGHSMQN 3324
SP|P0C6W2|R1AB_BCHK3              TTLNGLWLDDTVYCPRHVVCTAEDMLNPNYDDLLIRKSNHSFLVQAGNVQLRVIGHSMQN 3318
SP|P0C6W6|R1AB_BCRP3              TTLNGLWLDDTVYCPRHVICTAEDMLNPNYEDLLIRKSNHSFLVQAGNVQLRVIGHSMQN 3322
SP|P0C6V9|R1AB_BC279              TTLNGLWLDDTVYCPRHVICTAEDMLNPNYEDLLIRKSNHSFLVQAGNVQLRVIGHSMQN 3330
TR|A0A0U1WHI4|A0A0U1WHI4_CVHSA    TTLNGLWLDDTVYCPRHVICTAEDMLNPNYEDLLIRKSNHSFLVQAGNVQLRVIGHSMQN 3319
TR|A0A0U1WHG0|A0A0U1WHG0_CVHSA    TTLNGLWLDDTVYCPRHVVCTVEDMLNPNYEDLLIRKSNHSFLVQTGNVQLRVIGHSMQN 3319
TR|A0A166ZL34|A0A166ZL34_9NIDO    TTLNGLWLDDTVYCPRHVVCTVEDMLNPNYEDLLIRKSNHSFLVQTGNVQLRVIGHSMQN 3126
TR|R9QTB2|R9QTB2_CVHSA            TTLNGLWLDDTVYCPRHVICTAEDMLNPNYEDLLIRKSNHSFLVQAGNVQLRVIGHSMQN 3316
TR|R9QTH2|R9QTH2_CVHSA            TTLNGLWLDDTVYCPRHVICTAEDMLNPNYEDLLIRKSNHSFLVQAGNVQLRVIGHSMQN 3325
SP|P0C6U8|R1A_CVHSA               TTLNGLWLDDTVYCPRHVICTAEDMLNPNYEDLLIRKSNHSFLVQAGNVQLRVIGHSMQN 3324
TR|Q6JH47|Q6JH47_CVHSA            TTLNGLWLDDTVYCPRHVICTAEDMLNPNYEDLLIRKANHSFLVQAGNVQLRVIGHSMQN 3324
TR|Q692E5|Q692E5_CVHSA            TTLNGLWLDDTVYCPRHVICTAEDMLNPNYEDLLIRKSNHSFLVQAGNVQLRVIGHSMQN 3324
SP|P0C6F8|R1A_BCHK3               TTLNGLWLDDTVYCPRHVVCTAEDMLNPNYDDLLIRKSNHSFLVQAGNVQLRVIGHSMQN 3318
TR|A0A0K1Z0N1|A0A0K1Z0N1_CVHSA    TTLNGLWLDDTVYCPRHVICTAEDMLNPNYEDLLIRKSNHSFLVQAGNVQLRVIGHSMQN 3324
SP|P0C6F5|R1A_BC279               TTLNGLWLDDTVYCPRHVICTAEDMLNPNYEDLLIRKSNHSFLVQAGNVQLRVIGHSMQN 3330
SP|P0C6T7|R1A_BCRP3               TTLNGLWLDDTVYCPRHVICTAEDMLNPNYEDLLIRKSNHSFLVQAGNVQLRVIGHSMQN 3322
                                  **********.*******:** *******:******:**.****:**************
```

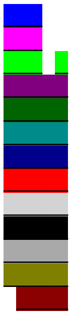

```
QHN73794                          CVLKLKVDTANPKTPKYKFVRIQPGQTFSVLACYNGSPSGVYQCAMRPNFTIKGSFLNGS 3407
SP|P0C6X7|R1AB_CVHSA              CLLRLKVDTSNPKTPKYKFVRIQPGQTFSVLACYNGSPSGVYQCAMRPNHTIKGSFLNGS 3384
TR|Q6UZF5|Q6UZF5_CVHSA            CLLRLKVDTSNPKTPKYKFVRIQPGQTFSVLACYNGSPSGVYQCAMRPNHTIKGSFLNGS 3384
TR|Q6UZF1|Q6UZF1_CVHSA            CLLRLKVDTSNPKTPKYKFVRIQPGQTFSVLACYNGSPSGVYQCAMRPNHTIKGSFLNGS 3384
TR|Q6JH48|Q6JH48_CVHSA            CLLRLKVDTSNPKTPKYKFVRIQPGQTFSVLACYNGSPSGVYQCAMRPNHTIKGSFLNGS 3384
TR|Q692E6|Q692E6_CVHSA            CLLRLKVDTSNPKTPKYKFVRIQPGQTFSVLACYNGSPSGVYQCAMRPNHTIKGSFLNGS 3384
TR|A0A0K1YZY7|A0A0K1YZY7_CVHSA    CLLRLKVDTSNPKTPKYKFVRIQPGQTFSVLACYNGSPSGVYQCAMRPNHTIKGSFLNGS 3384
SP|P0C6W2|R1AB_BCHK3              CLLRLKVDTSNPKTPKYKFVRIQPGQTFSVLACYNGSPSGVYQCAMRPNHTIKGSFLNGS 3378
SP|P0C6W6|R1AB_BCRP3              CLLRLKVDTSNPKTPKYKFVRIQPGQTFSVLACYNGSPSGVYQCAMRPNHTIKGSFLNGS 3382
SP|P0C6V9|R1AB_BC279              CLLRLKVDTSNPKTPKYKFVRIQPGQTFSVLACYNGSPSGVYQCAMRPNYTIKGSFLNGS 3390
TR|A0A0U1WHI4|A0A0U1WHI4_CVHSA    CLLRLKVDTSNPKTPKYKFVRIQPGQTFSVLACYNGSPSGVYQCAMRPNYTIKGSFLNGS 3379
TR|A0A0U1WHG0|A0A0U1WHG0_CVHSA    CLLRLKVDTSNPKTPKYKFVRIQPGQTFSVLACYNGSPSGVYQCAMRPNYTIKGSFLNGS 3379
TR|A0A166ZL34|A0A166ZL34_9NIDO    CLLRLKVDTSNPKTPKYKFVRIQPGQTFSVLACYNGSPSGVYQCAMRPNYTIKGSFLNGS 3186
TR|R9QTB2|R9QTB2_CVHSA            CLLRLKVDTSNPKTPKYKFVRIQPGQTFSVLACYNGSPSGVYQCAMRPNHTIKGSFLNGS 3376
TR|R9QTH2|R9QTH2_CVHSA            CLLRLKVDTSNPKTPKYKFVRIQPGQTFSVLACYNGSPSGVYQCAMRPNHTIKGSFLNGS 3385
SP|P0C6U8|R1A_CVHSA               CLLRLKVDTSNPKTPKYKFVRIQPGQTFSVLACYNGSPSGVYQCAMRPNHTIKGSFLNGS 3384
TR|Q6JH47|Q6JH47_CVHSA            CLLRLKVDTSNPKTPKYKFVRIQPGQTFSVLACYNGSPSGVYQCAMRPNHTIKGSFLNGS 3384
TR|Q692E5|Q692E5_CVHSA            CLLRLKVDTSNPKTPKYKFVRIQPGQTFSVLACYNGSPSGVYQCAMRPNHTIKGSFLNGS 3384
SP|P0C6F8|R1A_BCHK3               CLLRLKVDTSNPKTPKYKFVRIQPGQTFSVLACYNGSPSGVYQCAMRPNHTIKGSFLNGS 3378
TR|A0A0K1Z0N1|A0A0K1Z0N1_CVHSA    CLLRLKVDTSNPKTPKYKFVRIQPGQTFSVLACYNGSPSGVYQCAMRPNHTIKGSFLNGS 3384
SP|P0C6F5|R1A_BC279               CLLRLKVDTSNPKTPKYKFVRIQPGQTFSVLACYNGSPSGVYQCAMRPNYTIKGSFLNGS 3390
SP|P0C6T7|R1A_BCRP3               CLLRLKVDTSNPKTPKYKFVRIQPGQTFSVLACYNGSPSGVYQCAMRPNHTIKGSFLNGS 3382
                                  *:*:*****:****************************************.**********
```

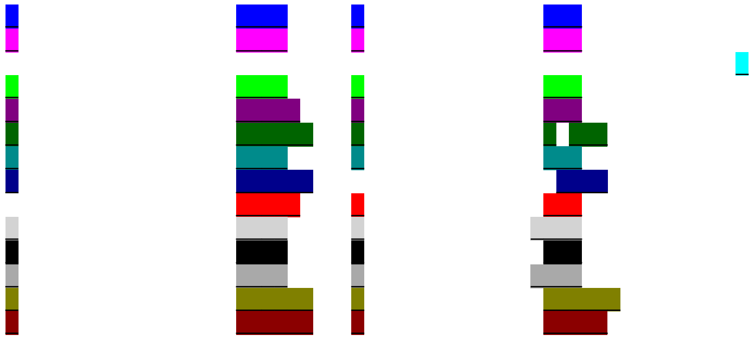

```
QHN73794                            CGSVGFNIDYDCVSFCYMHHMELPTGVHAGTDLEGNFYGPFVDRQTAQAAGTDTTITVNV 3467
SP|P0C6X7|R1AB_CVHSA                CGSVGFNIDYDCVSFCYMHHMELPTGVHAGTDLEGKFYGPFVDRQTAQAAGTDTTITLNV 3444
TR|Q6UZF5|Q6UZF5_CVHSA              CGSVGFNIDYDCVSFCYMHHMELPTGVHAGTDLEGKFYGPFVDRQTAQAAGTDTTITLNV 3444
TR|Q6UZF1|Q6UZF1_CVHSA              CGSVGFNIDYDCVSFCYMHHMELPTGVHAGTDLEGKFYGPFVDRQTAQAAGTDTTITLNV 3444
TR|Q6JH48|Q6JH48_CVHSA              CGSVGFNIDYDCVSFCYMHHMELPTGVHAGTDLEGKFYGPFVDRQTAQAAGTDTTITLNV 3444
TR|Q692E6|Q692E6_CVHSA              CGSVGFNIDYDCVSFCYMHHMELPTGVHAGTDLEGKFYGPFVDRQTAQAAGTDTTITLNV 3444
TR|A0A0K1YZY7|A0A0K1YZY7_CVHSA      CGSVGFNIDYDCVSFCYMHHMELPTGVHAGTDLEGKFYGPFVDRQTAQAAGTDTTITLNV 3444
SP|P0C6W2|R1AB_BCHK3                CGSVGFNIDYDCVSFCYMHHMELPTGVHAGTDLEGKFYGPFVDRQTAQAAGTDTTITLNV 3438
SP|P0C6W6|R1AB_BCRP3                CGSVGFNIDYDCVSFCYMHHMELPTEVHAGTDLEGKFYGPFVDRQTAQAAGTDTTITLNV 3442
SP|P0C6V9|R1AB_BC279                CGSVGFNIDYDCVSFCYMHHMELPTGVHAGTDLEGKFYGPFVDRQTAQAAGTDTTITLNV 3450
TR|A0A0U1WHI4|A0A0U1WHI4_CVHSA      CGSVGFNIDYDCVSFCYMHHMELPTGVHAGTDLEGKFYGPFVDRQTAQAAGTDTTITLNV 3439
TR|A0A0U1WHG0|A0A0U1WHG0_CVHSA      CGSVGFNIDYDCVSFCYMHHMELPTGVHAGTDLEGKFYGPFVDRQTAQAAGTDTTITLNV 3439
TR|A0A166ZL34|A0A166ZL34_9NIDO      CGSVGFNIDYDCVSFCYMHHMELPTGVHAGTDLEGKFYGPFVDRQTAQAAGTDTAITLNV 3246
TR|R9QTB2|R9QTB2_CVHSA              CGSVGFNIDYDCVSFCYMHHMELPTGVHAGTDLEGKFYGPFVDRQTAQAAGTDTTITLNV 3436
TR|R9QTH2|R9QTH2_CVHSA              CGSVGFNIDYDCVSFCYMHHMELPTGVHAGTDLEGKFYGPFVDRQTAQAAGTDTTITLNV 3445
SP|P0C6U8|R1A_CVHSA                 CGSVGFNIDYDCVSFCYMHHMELPTGVHAGTDLEGKFYGPFVDRQTAQAAGTDTTITLNV 3444
TR|Q6JH47|Q6JH47_CVHSA              CGSVGFNIDYDCVSFCYMHHMELPTGVHAGTDLEGKFYGPFVDRQTAQAAGTDTTITLNV 3444
TR|Q692E5|Q692E5_CVHSA              CGSVGFNIDYDCVSFCYMHHMELPTGVHAGTDLEGKFYGPFVDRQTAQAAGTDTTITLNV 3444
SP|P0C6F8|R1A_BCHK3                 CGSVGFNIDYDCVSFCYMHHMELPTGVHAGTDLEGKFYGPFVDRQTAQAAGTDTTITLNV 3438
TR|A0A0K1Z0N1|A0A0K1Z0N1_CVHSA      CGSVGFNIDYDCVSFCYMHHMELPTGVHAGTDLEGKFYGPFVDRQTAQAAGTDTTITLNV 3444
SP|P0C6F5|R1A_BC279                 CGSVGFNIDYDCVSFCYMHHMELPTGVHAGTDLEGKFYGPFVDRQTAQAAGTDTTITLNV 3450
SP|P0C6T7|R1A_BCRP3                 CGSVGFNIDYDCVSFCYMHHMELPTEVHAGTDLEGKFYGPFVDRQTAQAAGTDTTITLNV 3442
                                    ************************* *********:******************:**:**
```

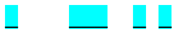

```
QHN73794                            LAWLYAAVINGDRWFLNRFTTTLNDFNLVAMKYNYEPLTQDHVDILGPLSAQTGIAVLDM 3527
SP|P0C6X7|R1AB_CVHSA                LAWLYAAVINGDRWFLNRFTTTLNDFNLVAMKYNYEPLTQDHVDILGPLSAQTGIAVLDM 3504
TR|Q6UZF5|Q6UZF5_CVHSA              LAWLYAAVINGDRWFLNRFTTTLNDFNLVAMKYNYEPLTQDHVDILGPLSAQTGIAVLDM 3504
TR|Q6UZF1|Q6UZF1_CVHSA              LAWLYAAVINGDRWFLNRFTTTLNDFNLVAMKYNYEPLTQDHVDILGPLSAQTGIAVLDM 3504
TR|Q6JH48|Q6JH48_CVHSA              LAWLYAAVINGDRWFLNRFTTTLNDFNLVAMKYNYEPLTQDHVDILGPLSAQTGIAVLDM 3504
TR|Q692E6|Q692E6_CVHSA              LAWLYAAVINGDRWFLNRFTTTLNDFNLVAMKYNYEPLTQDHVDILGPLSAQTGIAVLDM 3504
TR|A0A0K1YZY7|A0A0K1YZY7_CVHSA      LAWLYAAVINGDRWFLNRFTTTLNDFNLVAMKYNYEPLTQDHVDVLGPLSAQTGIAVLDM 3504
SP|P0C6W2|R1AB_BCHK3                LAWLYAAVINGDRWFLNRFTTTLNDFNLVAMKYNYEPLTQDHVDILGPLSAQTGIAVLDM 3498
SP|P0C6W6|R1AB_BCRP3                LAWLYAAVINGDRWFLNRFTTTLNDFNLVAMKYNYEPLTQDHVDILGPLSAQTGIAVLDM 3502
SP|P0C6V9|R1AB_BC279                LAWLYAAVINGDRWFLNRFTTTLNDFNLVAMKYNYEPLTQDHVDILGPLSAQTGIAVLDM 3510
TR|A0A0U1WHI4|A0A0U1WHI4_CVHSA      LAWLYAAVINGDRWFLNRFTTTLNDFNLVAMKYNYEPLTQDHVDILGPLSAQTGIAVLDM 3499
TR|A0A0U1WHG0|A0A0U1WHG0_CVHSA      LAWLYAAVINGDRWFLNRFTTTLNDFNLVAMKYNYEPLTQDHVDILGPLSAQTGIAVLDM 3499
TR|A0A166ZL34|A0A166ZL34_9NIDO      LAWLYAAVINGDRWFLNRFTTTLNDFNLVAMKYNYEPLTQDHVDILGPLSAQTGIAVLDM 3306
TR|R9QTB2|R9QTB2_CVHSA              LAWLYAAVINGDRWFLNRFTTTLNDFNLVAMKYNYEPLTQDHVDILGPLSAQTGIAVLDM 3496
TR|R9QTH2|R9QTH2_CVHSA              LAWLYAAVINGDRWFLNRFTTTLNDFNLVAMKYNYEPLTQDHVDILGPLSAQTGIAVLDM 3505
SP|P0C6U8|R1A_CVHSA                 LAWLYAAVINGDRWFLNRFTTTLNDFNLVAMKYNYEPLTQDHVDILGPLSAQTGIAVLDM 3504
TR|Q6JH47|Q6JH47_CVHSA              LAWLYAAVINGDRWFLNRFTTTLNDFNLVAMKYNYEPLTQDHVDILGPLSAQTGIAVLDM 3504
TR|Q692E5|Q692E5_CVHSA              LAWLYAAVINGDRWFLNRFTTTLNDFNLVAMKYNYEPLTQDHVDILGPLSAQTGIAVLDM 3504
SP|P0C6F8|R1A_BCHK3                 LAWLYAAVINGDRWFLNRFTTTLNDFNLVAMKYNYEPLTQDHVDILGPLSAQTGIAVLDM 3498
TR|A0A0K1Z0N1|A0A0K1Z0N1_CVHSA      LAWLYAAVINGDRWFLNRFTTTLNDFNLVAMKYNYEPLTQDHVDVLGPLSAQTGIAVLDM 3504
SP|P0C6F5|R1A_BC279                 LAWLYAAVINGDRWFLNRFTTTLNDFNLVAMKYNYEPLTQDHVDILGPLSAQTGIAVLDM 3510
SP|P0C6T7|R1A_BCRP3                 LAWLYAAVINGDRWFLNRFTTTLNDFNLVAMKYNYEPLTQDHVDILGPLSAQTGIAVLDM 3502
                                    ********************************************:***************
```

```
QHN73794                         CASLKELLQNGMNGRTILGSALLEDEFTPFDVVRQCSGVTFQSAVKRTIKGTHHWLLLTI 3587
SP|P0C6X7|R1AB_CVHSA             CAALKELLQNGMNGRTILGSTILEDEFTPFDVVRQCSGVTFQGKFKKIVKGTHHWMLLTF 3564
TR|Q6UZF5|Q6UZF5_CVHSA           CAALKELLQNGMNGRTILGSTILEDEFTPFDVVRQCSGVTFQGKFKKIVKGTHHWMLLTF 3564
TR|Q6UZF1|Q6UZF1_CVHSA           CAALKELLQNGMNGRTILGSTILEDEFTPFDVVRQCSGVTFQGKFKKIVKGTHHWMLLTF 3564
TR|Q6JH48|Q6JH48_CVHSA           CAALKELLQNGMNGRTILGSTILEDEFTPFDVVRQCSGVTFQGKFKKIVKGTHHWMLLTF 3564
TR|Q692E6|Q692E6_CVHSA           CAALKELLQNGMNGRTILGSTILEDEFTPFDVVRQCSGVTFQGKFKKIVKGTHHWMLLTF 3564
TR|A0A0K1YZY7|A0A0K1YZY7_CVHSA CAALKELLQNGMNGRTILGSTILEDEFTPFDVVRQCSGVTFQGKFKKIVKGTHHWMLLTF 3564
SP|P0C6W2|R1AB_BCHK3             CAALKELLQNGMNGRTILGSTILEDEFTPFDVVRQCSGVTFQGKFKKIVKGTHHWMLLTF 3558
SP|P0C6W6|R1AB_BCRP3             CAALKELLQNGMNGRTILGSTILEDEFTPFDVVRQCSGVTFQGKFKRIVKGTHHWMLLTF 3562
SP|P0C6V9|R1AB_BC279             CAALKELLQNGMNGRTILGSTILEDEFTPFDVVRQCSGVTFQGKFKKIVKGTHHWMLLTF 3570
TR|A0A0U1WHI4|A0A0U1WHI4_CVHSA CAALKELLQNGMNGRTILGSTILEDEFTPFDVVRQCSGVTFQGKFKKIVKGTHHWMLLTF 3559
TR|A0A0U1WHG0|A0A0U1WHG0_CVHSA CAALKELLQNGMNGRTILGSTILEDEFTPFDVVRQCSGVTFQGKFKKIVKGTHHWMLLTF 3559
TR|A0A166ZL34|A0A166ZL34_9NIDO CAALKELLQNGMNGRTILGSTILEDEFTPFDVVRQCSGVTFQGKFKKIVKGTHHWMLLTF 3366
TR|R9QTB2|R9QTB2_CVHSA           CAALKELLQNGMNGRTILGSTILEDEFTPFDVVRQCSGVTFQGKFKKIVKGTHHWMLLTF 3556
TR|R9QTH2|R9QTH2_CVHSA           CAALKELLQNGMNGRTILGSTILEDEFTPFDVVRQCSGVTFQGKFKKIVKGTHHWVLLTF 3565
SP|P0C6U8|R1A_CVHSA              CAALKELLQNGMNGRTILGSTILEDEFTPFDVVRQCSGVTFQGKFKKIVKGTHHWMLLTF 3564
TR|Q6JH47|Q6JH47_CVHSA           CAALKELLQNGMNGRTILGSTILEDEFTPFDVVRQCSGVTFQGKFKKIVKGTHHWMLLTF 3564
TR|Q692E5|Q692E5_CVHSA           CAALKELLQNGMNGRTILGSTILEDEFTPFDVVRQCSGVTFQGKFKKIVKGTHHWMLLTF 3564
SP|P0C6F8|R1A_BCHK3              CAALKELLQNGMNGRTILGSTILEDEFTPFDVVRQCSGVTFQGKFKKIVKGTHHWMLLTF 3558
TR|A0A0K1Z0N1|A0A0K1Z0N1_CVHSA CAALKELLQNGMNGRTILGSTILEDEFTPFDVVRQCSGVTFQGKFKKIVKGTHHWMLLTF 3564
SP|P0C6F5|R1A_BC279              CAALKELLQNGMNGRTILGSTILEDEFTPFDVVRQCSGVTFQGKFKKIVKGTHHWMLLTF 3570
SP|P0C6T7|R1A_BCRP3              CAALKELLQNGMNGRTILGSTILEDEFTPFDVVRQCSGVTFQGKFKRIVKGTHHWMLLTF 3562
                                 **:*****************::********************. .*: :******:***:
```
