## Supplementary material for "Structural genomics and interactomics of 2019 Wuhan novel coronavirus, 2019-nCoV, indicate evolutionary conserved functional regions of viral proteins": Ligand binding site mapping for wNsp5 (SARS Group 1)

Ligand BS:

benzyl (2-oxopropyl)carbamate (3D62-959, SARS)

N-[(1R)-2-(tert-butylamino)-2-oxo-1-(pyridin-3-yl)ethyl]-N-(4-tert-butylphenyl)furan-2-carboxamide (3V3M-0EN, SARS)

N-[4-(acetylamino)phenyl]-2-(1H-benzotriazol-1-yl)-N-[(1R)-2-[(2-methylbutan-2-yl)amino]-1-(1-methyl-1H-pyrrol-2-yl)-2-oxoethyl]acetamide (4MDS-23H, SARS)

(2S)-2-({[(3S,4aR,8aS)-2-(4-bromobenzoyl)decahydroisoquinolin-3-yl]methyl}amino)-3-(1H-imidazol-5-yl)propanal (4TWW-3A7, SARS)

(2S)-2-({[(3S,4aR,8aS)-2-(biphenyl-4-ylcarbonyl)decahydroisoquinolin-3-yl]methyl}amino)-3-(1H-imidazol-5-yl)propanal (4TWY-3BL, SARS)

(2S)-2-({[(3R,4aS,8aR)-2-(biphenyl-4-ylcarbonyl)decahydroisoquinolin-3-yl]methyl}amino)-3-(1H-imidazol-5-yl)propanal (4WY3-3X5, SARS)

N-(3-FUROYL)-D-VALYL-L-VALYL-N~1~-((1R,2Z)-4-ETHOXY-4-OXO-1-{[(3S)-2-OXOPYRROLIDIN-3-YL]METHYL}BUT-2-ENYL)-D-LEUCINAMIDE (2AMD-9IN, SARS)

(5S,8S,14R)-ETHYL 11-(3-AMINO-3-OXOPROPYL)-8-BENZYL-14-HYDROXY-5-ISOBUTYL-3,6,9,12-TETRAOXO-1-PHENYL-2-OXA-4,7,10,11-TETRAAZAPENTADECAN-15-OATE (2A5K-AZP, SARS)

N-((3S,6R)-6-((S,E)-4-ETHOXYCARBONYL-1-((S)-2-OXOPYRROLIDIN-3-YL)BUT-3-EN-2-YLCARBAMOYL)-2,9-DIMETHYL-4-OXODEC-8-EN-3-YL)-5-METHYLISOXAZOLE-3-CARBOXAMIDE (2ALV-CY6, SARS)

ETHYL (4R)-4-{[(2R,5S)-5-{[N-(TERT-BUTOXYCARBONYL)-L-SERYL]AMINO}-6-METHYL-2-(3-METHYLBUT-2-EN-1-YL)-4-OXOHEPTANOYL]AMINO}-5-[(3R)-2-OXOPYRROLIDIN-3-YL]PENTANOATE (2QIQ-CYV, SARS)

ETHYL (2E,4S)-4-[((2R)-2-{[N-(TERT-BUTOXYCARBONYL)-L-VALYL]AMINO}-2-PHENYLETHANOYL)AMINO]-5-[(3S)-2-OXOPYRROLIDIN-3-YL]PENT-2-ENOATE (2D2D-ENB, SARS)

ETHYL (4R)-4-({N-[(BENZYLOXY)CARBONYL]-L-PHENYLALANYL}AMINO)-5-[(3S)-2-OXOPYRROLIDIN-3-YL]PENTANOATE (3SZN-G75, SARS)

ETHYL (4R)-4-{[N-(TERT-BUTOXYCARBONYL)-L-PHENYLALANYL]AMINO}-5-[(3S)-2-OXOPYRROLIDIN-3-YL]PENTANOATE (3TIT-G81, SARS)

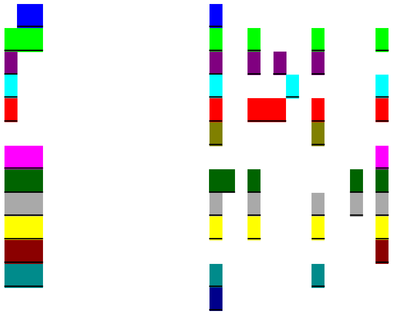

```
QHN73794                           TTLNGLWLDDVVYCPRHVICTSEDMLNPNYEDLLIRKSNHNFLVQAGNVQLRVIGHSMQN 3347
SP|P0C6X7|R1AB_CVHSA               TTLNGLWLDDTVYCPRHVICTAEDMLNPNYEDLLIRKSNHSFLVQAGNVQLRVIGHSMQN 3324
TR|Q6UZF5|Q6UZF5_CVHSA             TTLNGLWLDDTVYCPRHVICTAEDMLNPNYEDLLIRKSNHSFLVQAGNVQLRVIGHSMQN 3324
TR|Q6UZF1|Q6UZF1_CVHSA             TTLNGLWLDDTVYCPRHVICTAEDMLNPNYEDLLIRKSNHSFLVQAGNVQLRVIGHSMQN 3324
TR|Q6JH48|Q6JH48_CVHSA             TTLNGLWLDDTVYCPRHVICTAEDMLNPNYEDLLIRKANHSFLVQAGNVQLRVIGHSMQN 3324
TR|Q692E6|Q692E6_CVHSA             TTLNGLWLDDTVYCPRHVICTAEDMLNPNYEDLLIRKSNHSFLVQAGNVQLRVIGHSMQN 3324
TR|A0A0K1YZY7|A0A0K1YZY7_CVHSA     TTLNGLWLDDTVYCPRHVICTAEDMLNPNYEDLLIRKSNHSFLVQAGNVQLRVIGHSMQN 3324
SP|P0C6W2|R1AB_BCHK3               TTLNGLWLDDTVYCPRHVVCTAEDMLNPNYDDLLIRKSNHSFLVQAGNVQLRVIGHSMQN 3318
SP|P0C6W6|R1AB_BCRP3               TTLNGLWLDDTVYCPRHVICTAEDMLNPNYEDLLIRKSNHSFLVQAGNVQLRVIGHSMQN 3322
SP|P0C6V9|R1AB_BC279               TTLNGLWLDDTVYCPRHVICTAEDMLNPNYEDLLIRKSNHSFLVQAGNVQLRVIGHSMQN 3330
TR|A0A0U1WHI4|A0A0U1WHI4_CVHSA     TTLNGLWLDDTVYCPRHVICTAEDMLNPNYEDLLIRKSNHSFLVQAGNVQLRVIGHSMQN 3319
TR|A0A0U1WHG0|A0A0U1WHG0_CVHSA     TTLNGLWLDDTVYCPRHVVCTVEDMLNPNYEDLLIRKSNHSFLVQTGNVQLRVIGHSMQN 3319
TR|A0A166ZL34|A0A166ZL34_9NIDO     TTLNGLWLDDTVYCPRHVVCTVEDMLNPNYEDLLIRKSNHSFLVQTGNVQLRVIGHSMQN 3126
TR|R9QTB2|R9QTB2_CVHSA             TTLNGLWLDDTVYCPRHVICTAEDMLNPNYEDLLIRKSNHSFLVQAGNVQLRVIGHSMQN 3316
TR|R9QTH2|R9QTH2_CVHSA             TTLNGLWLDDTVYCPRHVICTAEDMLNPNYEDLLIRKSNHSFLVQAGNVQLRVIGHSMQN 3325
SP|P0C6U8|R1A_CVHSA                TTLNGLWLDDTVYCPRHVICTAEDMLNPNYEDLLIRKSNHSFLVQAGNVQLRVIGHSMQN 3324
TR|Q6JH47|Q6JH47_CVHSA             TTLNGLWLDDTVYCPRHVICTAEDMLNPNYEDLLIRKANHSFLVQAGNVQLRVIGHSMQN 3324
TR|Q692E5|Q692E5_CVHSA             TTLNGLWLDDTVYCPRHVICTAEDMLNPNYEDLLIRKSNHSFLVQAGNVQLRVIGHSMQN 3324
SP|P0C6F8|R1A_BCHK3                TTLNGLWLDDTVYCPRHVVCTAEDMLNPNYDDLLIRKSNHSFLVQAGNVQLRVIGHSMQN 3318
TR|A0A0K1Z0N1|A0A0K1Z0N1_CVHSA     TTLNGLWLDDTVYCPRHVICTAEDMLNPNYEDLLIRKSNHSFLVQAGNVQLRVIGHSMQN 3324
SP|P0C6F5|R1A_BC279                TTLNGLWLDDTVYCPRHVICTAEDMLNPNYEDLLIRKSNHSFLVQAGNVQLRVIGHSMQN 3330
SP|P0C6T7|R1A_BCRP3                TTLNGLWLDDTVYCPRHVICTAEDMLNPNYEDLLIRKSNHSFLVQAGNVQLRVIGHSMQN 3322
                                   **********.*******:** ********:******:**.****:**************
```

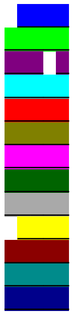

```
QHN73794                           CVLKLKVDTANPKTPKYKFVRIQPGQTFSVLACYNGSPSGVYQCAMRPNFTIKGSFLNGS 3407
SP|P0C6X7|R1AB_CVHSA               CLLRLKVDTSNPKTPKYKFVRIQPGQTFSVLACYNGSPSGVYQCAMRPNHTIKGSFLNGS 3384
TR|Q6UZF5|Q6UZF5_CVHSA             CLLRLKVDTSNPKTPKYKFVRIQPGQTFSVLACYNGSPSGVYQCAMRPNHTIKGSFLNGS 3384
TR|Q6UZF1|Q6UZF1_CVHSA             CLLRLKVDTSNPKTPKYKFVRIQPGQTFSVLACYNGSPSGVYQCAMRPNHTIKGSFLNGS 3384
TR|Q6JH48|Q6JH48_CVHSA             CLLRLKVDTSNPKTPKYKFVRIQPGQTFSVLACYNGSPSGVYQCAMRPNHTIKGSFLNGS 3384
TR|Q692E6|Q692E6_CVHSA             CLLRLKVDTSNPKTPKYKFVRIQPGQTFSVLACYNGSPSGVYQCAMRPNHTIKGSFLNGS 3384
TR|A0A0K1YZY7|A0A0K1YZY7_CVHSA     CLLRLKVDTSNPKTPKYKFVRIQPGQTFSVLACYNGSPSGVYQCAMRPNHTIKGSFLNGS 3384
SP|P0C6W2|R1AB_BCHK3               CLLRLKVDTSNPKTPKYKFVRIQPGQTFSVLACYNGSPSGVYQCAMRPNHTIKGSFLNGS 3378
SP|P0C6W6|R1AB_BCRP3               CLLRLKVDTSNPKTPKYKFVRIQPGQTFSVLACYNGSPSGVYQCAMRPNHTIKGSFLNGS 3382
SP|P0C6V9|R1AB_BC279               CLLRLKVDTSNPKTPKYKFVRIQPGQTFSVLACYNGSPSGVYQCAMRPNYTIKGSFLNGS 3390
TR|A0A0U1WHI4|A0A0U1WHI4_CVHSA     CLLRLKVDTSNPKTPKYKFVRIQPGQTFSVLACYNGSPSGVYQCAMRPNYTIKGSFLNGS 3379
TR|A0A0U1WHG0|A0A0U1WHG0_CVHSA     CLLRLKVDTSNPKTPKYKFVRIQPGQTFSVLACYNGSPSGVYQCAMRPNYTIKGSFLNGS 3379
TR|A0A166ZL34|A0A166ZL34_9NIDO     CLLRLKVDTSNPKTPKYKFVRIQPGQTFSVLACYNGSPSGVYQCAMRPNYTIKGSFLNGS 3186
TR|R9QTB2|R9QTB2_CVHSA             CLLRLKVDTSNPKTPKYKFVRIQPGQTFSVLACYNGSPSGVYQCAMRPNHTIKGSFLNGS 3376
TR|R9QTH2|R9QTH2_CVHSA             CLLRLKVDTSNPKTPKYKFVRIQPGQTFSVLACYNGSPSGVYQCAMRPNHTIKGSFLNGS 3385
SP|P0C6U8|R1A_CVHSA                CLLRLKVDTSNPKTPKYKFVRIQPGQTFSVLACYNGSPSGVYQCAMRPNHTIKGSFLNGS 3384
TR|Q6JH47|Q6JH47_CVHSA             CLLRLKVDTSNPKTPKYKFVRIQPGQTFSVLACYNGSPSGVYQCAMRPNHTIKGSFLNGS 3384
TR|Q692E5|Q692E5_CVHSA             CLLRLKVDTSNPKTPKYKFVRIQPGQTFSVLACYNGSPSGVYQCAMRPNHTIKGSFLNGS 3384
SP|P0C6F8|R1A_BCHK3                CLLRLKVDTSNPKTPKYKFVRIQPGQTFSVLACYNGSPSGVYQCAMRPNHTIKGSFLNGS 3378
TR|A0A0K1Z0N1|A0A0K1Z0N1_CVHSA     CLLRLKVDTSNPKTPKYKFVRIQPGQTFSVLACYNGSPSGVYQCAMRPNHTIKGSFLNGS 3384
SP|P0C6F5|R1A_BC279                CLLRLKVDTSNPKTPKYKFVRIQPGQTFSVLACYNGSPSGVYQCAMRPNYTIKGSFLNGS 3390
SP|P0C6T7|R1A_BCRP3                CLLRLKVDTSNPKTPKYKFVRIQPGQTFSVLACYNGSPSGVYQCAMRPNHTIKGSFLNGS 3382
                                   *:*:*****:****************************************.**********
```

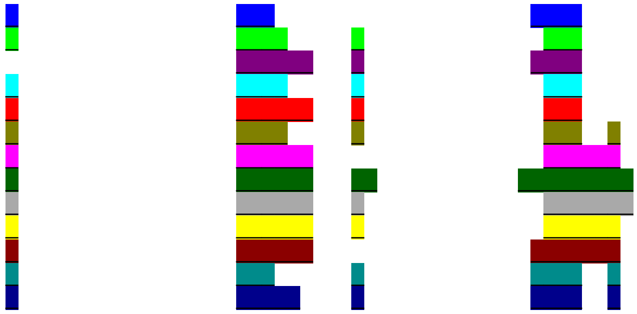

```
QHN73794                         CGSVGFNIDYDCVSFCYMHHMELPTGVHAGTDLEGNFYGPFVDRQTAQAAGTDTTITVNV 3467
SP|P0C6X7|R1AB_CVHSA             CGSVGFNIDYDCVSFCYMHHMELPTGVHAGTDLEGKFYGPFVDRQTAQAAGTDTTITLNV 3444
TR|Q6UZF5|Q6UZF5_CVHSA           CGSVGFNIDYDCVSFCYMHHMELPTGVHAGTDLEGKFYGPFVDRQTAQAAGTDTTITLNV 3444
TR|Q6UZF1|Q6UZF1_CVHSA           CGSVGFNIDYDCVSFCYMHHMELPTGVHAGTDLEGKFYGPFVDRQTAQAAGTDTTITLNV 3444
TR|Q6JH48|Q6JH48_CVHSA           CGSVGFNIDYDCVSFCYMHHMELPTGVHAGTDLEGKFYGPFVDRQTAQAAGTDTTITLNV 3444
TR|Q692E6|Q692E6_CVHSA           CGSVGFNIDYDCVSFCYMHHMELPTGVHAGTDLEGKFYGPFVDRQTAQAAGTDTTITLNV 3444
TR|A0A0K1YZY7|A0A0K1YZY7_CVHSA CGSVGFNIDYDCVSFCYMHHMELPTGVHAGTDLEGKFYGPFVDRQTAQAAGTDTTITLNV 3444
SP|P0C6W2|R1AB_BCHK3             CGSVGFNIDYDCVSFCYMHHMELPTGVHAGTDLEGKFYGPFVDRQTAQAAGTDTTITLNV 3438
SP|P0C6W6|R1AB_BCRP3             CGSVGFNIDYDCVSFCYMHHMELPTEVHAGTDLEGKFYGPFVDRQTAQAAGTDTTITLNV 3442
SP|P0C6V9|R1AB_BC279             CGSVGFNIDYDCVSFCYMHHMELPTGVHAGTDLEGKFYGPFVDRQTAQAAGTDTTITLNV 3450
TR|A0A0U1WHI4|A0A0U1WHI4_CVHSA CGSVGFNIDYDCVSFCYMHHMELPTGVHAGTDLEGKFYGPFVDRQTAQAAGTDTTITLNV 3439
TR|A0A0U1WHG0|A0A0U1WHG0_CVHSA CGSVGFNIDYDCVSFCYMHHMELPTGVHAGTDLEGKFYGPFVDRQTAQAAGTDTTITLNV 3439
TR|A0A166ZL34|A0A166ZL34_9NIDO CGSVGFNIDYDCVSFCYMHHMELPTGVHAGTDLEGKFYGPFVDRQTAQAAGTDTAITLNV 3246
TR|R9QTB2|R9QTB2_CVHSA           CGSVGFNIDYDCVSFCYMHHMELPTGVHAGTDLEGKFYGPFVDRQTAQAAGTDTTITLNV 3436
TR|R9QTH2|R9QTH2_CVHSA           CGSVGFNIDYDCVSFCYMHHMELPTGVHAGTDLEGKFYGPFVDRQTAQAAGTDTTITLNV 3445
SP|P0C6U8|R1A_CVHSA              CGSVGFNIDYDCVSFCYMHHMELPTGVHAGTDLEGKFYGPFVDRQTAQAAGTDTTITLNV 3444
TR|Q6JH47|Q6JH47_CVHSA           CGSVGFNIDYDCVSFCYMHHMELPTGVHAGTDLEGKFYGPFVDRQTAQAAGTDTTITLNV 3444
TR|Q692E5|Q692E5_CVHSA           CGSVGFNIDYDCVSFCYMHHMELPTGVHAGTDLEGKFYGPFVDRQTAQAAGTDTTITLNV 3444
SP|P0C6F8|R1A_BCHK3              CGSVGFNIDYDCVSFCYMHHMELPTGVHAGTDLEGKFYGPFVDRQTAQAAGTDTTITLNV 3438
TR|A0A0K1Z0N1|A0A0K1Z0N1_CVHSA CGSVGFNIDYDCVSFCYMHHMELPTGVHAGTDLEGKFYGPFVDRQTAQAAGTDTTITLNV 3444
SP|P0C6F5|R1A_BC279              CGSVGFNIDYDCVSFCYMHHMELPTGVHAGTDLEGKFYGPFVDRQTAQAAGTDTTITLNV 3450
SP|P0C6T7|R1A_BCRP3              CGSVGFNIDYDCVSFCYMHHMELPTEVHAGTDLEGKFYGPFVDRQTAQAAGTDTTITLNV 3442
                                 ************************* *********:******************:**:**

QHN73794                         LAWLYAAVINGDRWFLNRFTTTLNDFNLVAMKYNYEPLTQDHVDILGPLSAQTGIAVLDM 3527
SP|P0C6X7|R1AB_CVHSA             LAWLYAAVINGDRWFLNRFTTTLNDFNLVAMKYNYEPLTQDHVDILGPLSAQTGIAVLDM 3504
TR|Q6UZF5|Q6UZF5_CVHSA           LAWLYAAVINGDRWFLNRFTTTLNDFNLVAMKYNYEPLTQDHVDILGPLSAQTGIAVLDM 3504
TR|Q6UZF1|Q6UZF1_CVHSA           LAWLYAAVINGDRWFLNRFTTTLNDFNLVAMKYNYEPLTQDHVDILGPLSAQTGIAVLDM 3504
TR|Q6JH48|Q6JH48_CVHSA           LAWLYAAVINGDRWFLNRFTTTLNDFNLVAMKYNYEPLTQDHVDILGPLSAQTGIAVLDM 3504
TR|Q692E6|Q692E6_CVHSA           LAWLYAAVINGDRWFLNRFTTTLNDFNLVAMKYNYEPLTQDHVDILGPLSAQTGIAVLDM 3504
TR|A0A0K1YZY7|A0A0K1YZY7_CVHSA LAWLYAAVINGDRWFLNRFTTTLNDFNLVAMKYNYEPLTQDHVDVLGPLSAQTGIAVLDM 3504
SP|P0C6W2|R1AB_BCHK3             LAWLYAAVINGDRWFLNRFTTTLNDFNLVAMKYNYEPLTQDHVDILGPLSAQTGIAVLDM 3498
SP|P0C6W6|R1AB_BCRP3             LAWLYAAVINGDRWFLNRFTTTLNDFNLVAMKYNYEPLTQDHVDILGPLSAQTGIAVLDM 3502
SP|P0C6V9|R1AB_BC279             LAWLYAAVINGDRWFLNRFTTTLNDFNLVAMKYNYEPLTQDHVDILGPLSAQTGIAVLDM 3510
TR|A0A0U1WHI4|A0A0U1WHI4_CVHSA LAWLYAAVINGDRWFLNRFTTTLNDFNLVAMKYNYEPLTQDHVDILGPLSAQTGIAVLDM 3499
TR|A0A0U1WHG0|A0A0U1WHG0_CVHSA LAWLYAAVINGDRWFLNRFTTTLNDFNLVAMKYNYEPLTQDHVDILGPLSAQTGIAVLDM 3499
TR|A0A166ZL34|A0A166ZL34_9NIDO LAWLYAAVINGDRWFLNRFTTTLNDFNLVAMKYNYEPLTQDHVDILGPLSAQTGIAVLDM 3306
TR|R9QTB2|R9QTB2_CVHSA           LAWLYAAVINGDRWFLNRFTTTLNDFNLVAMKYNYEPLTQDHVDILGPLSAQTGIAVLDM 3496
TR|R9QTH2|R9QTH2_CVHSA           LAWLYAAVINGDRWFLNRFTTTLNDFNLVAMKYNYEPLTQDHVDILGPLSAQTGIAVLDM 3505
SP|P0C6U8|R1A_CVHSA              LAWLYAAVINGDRWFLNRFTTTLNDFNLVAMKYNYEPLTQDHVDILGPLSAQTGIAVLDM 3504
TR|Q6JH47|Q6JH47_CVHSA           LAWLYAAVINGDRWFLNRFTTTLNDFNLVAMKYNYEPLTQDHVDILGPLSAQTGIAVLDM 3504
TR|Q692E5|Q692E5_CVHSA           LAWLYAAVINGDRWFLNRFTTTLNDFNLVAMKYNYEPLTQDHVDILGPLSAQTGIAVLDM 3504
SP|P0C6F8|R1A_BCHK3              LAWLYAAVINGDRWFLNRFTTTLNDFNLVAMKYNYEPLTQDHVDILGPLSAQTGIAVLDM 3498
TR|A0A0K1Z0N1|A0A0K1Z0N1_CVHSA LAWLYAAVINGDRWFLNRFTTTLNDFNLVAMKYNYEPLTQDHVDVLGPLSAQTGIAVLDM 3504
SP|P0C6F5|R1A_BC279              LAWLYAAVINGDRWFLNRFTTTLNDFNLVAMKYNYEPLTQDHVDILGPLSAQTGIAVLDM 3510
SP|P0C6T7|R1A_BCRP3              LAWLYAAVINGDRWFLNRFTTTLNDFNLVAMKYNYEPLTQDHVDILGPLSAQTGIAVLDM 3502
                                 ********************************************:***************
```
