## Supplementary material for "Structural genomics and interactomics of 2019 Wuhan novel coronavirus, 2019-nCoV, indicate evolutionary conserved functional regions of viral proteins": Ligand binding site mapping for wNsp5 (SARS Group 2)

Ligand BS:

ETHYL (5S,8S,11R)-8-BENZYL-5-(3-TERT-BUTOXY-3-OXOPROPYL)-3,6,9-TRIOXO-11-{[(3S)-2-OXOPYRROLIDIN-3-YL]METHYL}-1-PHENYL-2-OXA-4,7,10-TRIAZATETRADECAN-14-OATE (3TIU-G82, SARS)

ETHYL (5S,8S,11R)-8-BENZYL-5-(2-TERT-BUTOXY-2-OXOETHYL)-3,6,9-TRIOXO-11-{[(3S)-2-OXOPYRROLIDIN-3-YL]METHYL}-1-PHENYL-2-OXA-4,7,10-TRIAZATETRADECAN-14-OATE (3TNS-G83, SARS)

N-[(benzyloxy)carbonyl]-O-tert-butyl-L-seryl-N-{(2R)-5-ethoxy-5-oxo-1-[(3S)-2-oxopyrrolidin-3-yl]pentan-2-yl}-L-phenylalaninamide (3TNT-G85, SARS)

N-[(5-METHYLISOXAZOL-3-YL)CARBONYL]-L-ALANYL-L-VALYL-N~1~-((1S)-4-ETHOXY-4-OXO-1-{[(3S)-2-OXOPYRROLIDIN-3-YL]METHYL}BUT-2-ENYL)-L-LEUCINAMIDE (1WOF-I12, SARS)

N-[(BENZYLOXY)CARBONYL]-O-(TERT-BUTYL)-L-THREONYL-3-CYCLOHEXYL-N-[(1S)-2-HYDROXY-1-{[(3S)-2-OXOPYRROLIDIN-3-YL]METHYL}ETHYL]-L-ALANINAMIDE (2GX4-NOL, SARS)

N-[(2S)-1-hydroxy-3-phenylpropan-2-yl]-Nalpha-[(2E)-3-phenylprop-2-enoyl]-L-phenylalaninamide (3SN8-S89, SARS)

(2S)-3-(1H-imidazol-5-yl)-2-({[(3S,4aR,8aS)-2-(N-phenyl-beta-alanyl)decahydroisoquinolin-3-yl]methyl}amino)propanal (5C5O-SDJ, SARS)

(2S)-3-(1H-imidazol-5-yl)-2-({[(3R,4aS,8aR)-2-(N-phenyl-beta-alanyl)decahydroisoquinolin-3-yl]methyl}amino)propanal (5C5N-SLH, SARS)

4-(DIMETHYLAMINO)BENZOIC ACID (2V6N-XP1, SARS)

N-[(benzyloxy)carbonyl]-3-[(2,2-dimethylpropanoyl)amino]-L-alanyl-N-[(1R)-4-oxo-1-{[(3S)-2-oxopyrrolidin-3-yl]methyl}pentyl]-L-leucinamide (2ZU4-ZU3, SARS)

N-[(benzyloxy)carbonyl]-O-tert-butyl-L-threonyl-N-[(1R)-4-cyclopropyl-4-oxo-1-{[(3S)-2-oxopyrrolidin-3-yl]methyl}butyl]-L-leucinamide (2ZU5-ZU5, SARS)

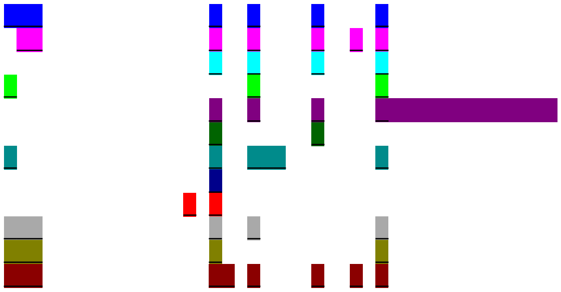

```
QHN73794                           TTLNGLWLDDVVYCPRHVICTSEDMLNPNYEDLLIRKSNHNFLVQAGNVQLRVIGHSMQN 3347
SP|P0C6X7|R1AB_CVHSA               TTLNGLWLDDTVYCPRHVICTAEDMLNPNYEDLLIRKSNHSFLVQAGNVQLRVIGHSMQN 3324
TR|Q6UZF5|Q6UZF5_CVHSA             TTLNGLWLDDTVYCPRHVICTAEDMLNPNYEDLLIRKSNHSFLVQAGNVQLRVIGHSMQN 3324
TR|Q6UZF1|Q6UZF1_CVHSA             TTLNGLWLDDTVYCPRHVICTAEDMLNPNYEDLLIRKSNHSFLVQAGNVQLRVIGHSMQN 3324
TR|Q6JH48|Q6JH48_CVHSA             TTLNGLWLDDTVYCPRHVICTAEDMLNPNYEDLLIRKANHSFLVQAGNVQLRVIGHSMQN 3324
TR|Q692E6|Q692E6_CVHSA             TTLNGLWLDDTVYCPRHVICTAEDMLNPNYEDLLIRKSNHSFLVQAGNVQLRVIGHSMQN 3324
TR|A0A0K1YZY7|A0A0K1YZY7_CVHSA     TTLNGLWLDDTVYCPRHVICTAEDMLNPNYEDLLIRKSNHSFLVQAGNVQLRVIGHSMQN 3324
SP|P0C6W2|R1AB_BCHK3               TTLNGLWLDDTVYCPRHVVCTAEDMLNPNYDDLLIRKSNHSFLVQAGNVQLRVIGHSMQN 3318
SP|P0C6W6|R1AB_BCRP3               TTLNGLWLDDTVYCPRHVICTAEDMLNPNYEDLLIRKSNHSFLVQAGNVQLRVIGHSMQN 3322
SP|P0C6V9|R1AB_BC279               TTLNGLWLDDTVYCPRHVICTAEDMLNPNYEDLLIRKSNHSFLVQAGNVQLRVIGHSMQN 3330
TR|A0A0U1WHI4|A0A0U1WHI4_CVHSA     TTLNGLWLDDTVYCPRHVICTAEDMLNPNYEDLLIRKSNHSFLVQAGNVQLRVIGHSMQN 3319
TR|A0A0U1WHG0|A0A0U1WHG0_CVHSA     TTLNGLWLDDTVYCPRHVVCTVEDMLNPNYEDLLIRKSNHSFLVQTGNVQLRVIGHSMQN 3319
TR|A0A166ZL34|A0A166ZL34_9NIDO     TTLNGLWLDDTVYCPRHVVCTVEDMLNPNYEDLLIRKSNHSFLVQTGNVQLRVIGHSMQN 3126
TR|R9QTB2|R9QTB2_CVHSA             TTLNGLWLDDTVYCPRHVICTAEDMLNPNYEDLLIRKSNHSFLVQAGNVQLRVIGHSMQN 3316
TR|R9QTH2|R9QTH2_CVHSA             TTLNGLWLDDTVYCPRHVICTAEDMLNPNYEDLLIRKSNHSFLVQAGNVQLRVIGHSMQN 3325
SP|P0C6U8|R1A_CVHSA                TTLNGLWLDDTVYCPRHVICTAEDMLNPNYEDLLIRKSNHSFLVQAGNVQLRVIGHSMQN 3324
TR|Q6JH47|Q6JH47_CVHSA             TTLNGLWLDDTVYCPRHVICTAEDMLNPNYEDLLIRKANHSFLVQAGNVQLRVIGHSMQN 3324
TR|Q692E5|Q692E5_CVHSA             TTLNGLWLDDTVYCPRHVICTAEDMLNPNYEDLLIRKSNHSFLVQAGNVQLRVIGHSMQN 3324
SP|P0C6F8|R1A_BCHK3                TTLNGLWLDDTVYCPRHVVCTAEDMLNPNYDDLLIRKSNHSFLVQAGNVQLRVIGHSMQN 3318
TR|A0A0K1Z0N1|A0A0K1Z0N1_CVHSA     TTLNGLWLDDTVYCPRHVICTAEDMLNPNYEDLLIRKSNHSFLVQAGNVQLRVIGHSMQN 3324
SP|P0C6F5|R1A_BC279                TTLNGLWLDDTVYCPRHVICTAEDMLNPNYEDLLIRKSNHSFLVQAGNVQLRVIGHSMQN 3330
SP|P0C6T7|R1A_BCRP3                TTLNGLWLDDTVYCPRHVICTAEDMLNPNYEDLLIRKSNHSFLVQAGNVQLRVIGHSMQN 3322
                                   **********.*******:** ********:******:**.****:**************
```

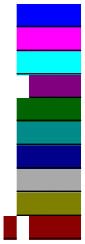

```
QHN73794                           CVLKLKVDTANPKTPKYKFVRIQPGQTFSVLACYNGSPSGVYQCAMRPNFTIKGSFLNGS 3407
SP|P0C6X7|R1AB_CVHSA               CLLRLKVDTSNPKTPKYKFVRIQPGQTFSVLACYNGSPSGVYQCAMRPNHTIKGSFLNGS 3384
TR|Q6UZF5|Q6UZF5_CVHSA             CLLRLKVDTSNPKTPKYKFVRIQPGQTFSVLACYNGSPSGVYQCAMRPNHTIKGSFLNGS 3384
TR|Q6UZF1|Q6UZF1_CVHSA             CLLRLKVDTSNPKTPKYKFVRIQPGQTFSVLACYNGSPSGVYQCAMRPNHTIKGSFLNGS 3384
TR|Q6JH48|Q6JH48_CVHSA             CLLRLKVDTSNPKTPKYKFVRIQPGQTFSVLACYNGSPSGVYQCAMRPNHTIKGSFLNGS 3384
TR|Q692E6|Q692E6_CVHSA             CLLRLKVDTSNPKTPKYKFVRIQPGQTFSVLACYNGSPSGVYQCAMRPNHTIKGSFLNGS 3384
TR|A0A0K1YZY7|A0A0K1YZY7_CVHSA     CLLRLKVDTSNPKTPKYKFVRIQPGQTFSVLACYNGSPSGVYQCAMRPNHTIKGSFLNGS 3384
SP|P0C6W2|R1AB_BCHK3               CLLRLKVDTSNPKTPKYKFVRIQPGQTFSVLACYNGSPSGVYQCAMRPNHTIKGSFLNGS 3378
SP|P0C6W6|R1AB_BCRP3               CLLRLKVDTSNPKTPKYKFVRIQPGQTFSVLACYNGSPSGVYQCAMRPNHTIKGSFLNGS 3382
SP|P0C6V9|R1AB_BC279               CLLRLKVDTSNPKTPKYKFVRIQPGQTFSVLACYNGSPSGVYQCAMRPNYTIKGSFLNGS 3390
TR|A0A0U1WHI4|A0A0U1WHI4_CVHSA     CLLRLKVDTSNPKTPKYKFVRIQPGQTFSVLACYNGSPSGVYQCAMRPNYTIKGSFLNGS 3379
TR|A0A0U1WHG0|A0A0U1WHG0_CVHSA     CLLRLKVDTSNPKTPKYKFVRIQPGQTFSVLACYNGSPSGVYQCAMRPNYTIKGSFLNGS 3379
TR|A0A166ZL34|A0A166ZL34_9NIDO     CLLRLKVDTSNPKTPKYKFVRIQPGQTFSVLACYNGSPSGVYQCAMRPNYTIKGSFLNGS 3186
TR|R9QTB2|R9QTB2_CVHSA             CLLRLKVDTSNPKTPKYKFVRIQPGQTFSVLACYNGSPSGVYQCAMRPNHTIKGSFLNGS 3376
TR|R9QTH2|R9QTH2_CVHSA             CLLRLKVDTSNPKTPKYKFVRIQPGQTFSVLACYNGSPSGVYQCAMRPNHTIKGSFLNGS 3385
SP|P0C6U8|R1A_CVHSA                CLLRLKVDTSNPKTPKYKFVRIQPGQTFSVLACYNGSPSGVYQCAMRPNHTIKGSFLNGS 3384
TR|Q6JH47|Q6JH47_CVHSA             CLLRLKVDTSNPKTPKYKFVRIQPGQTFSVLACYNGSPSGVYQCAMRPNHTIKGSFLNGS 3384
TR|Q692E5|Q692E5_CVHSA             CLLRLKVDTSNPKTPKYKFVRIQPGQTFSVLACYNGSPSGVYQCAMRPNHTIKGSFLNGS 3384
SP|P0C6F8|R1A_BCHK3                CLLRLKVDTSNPKTPKYKFVRIQPGQTFSVLACYNGSPSGVYQCAMRPNHTIKGSFLNGS 3378
TR|A0A0K1Z0N1|A0A0K1Z0N1_CVHSA     CLLRLKVDTSNPKTPKYKFVRIQPGQTFSVLACYNGSPSGVYQCAMRPNHTIKGSFLNGS 3384
SP|P0C6F5|R1A_BC279                CLLRLKVDTSNPKTPKYKFVRIQPGQTFSVLACYNGSPSGVYQCAMRPNYTIKGSFLNGS 3390
SP|P0C6T7|R1A_BCRP3                CLLRLKVDTSNPKTPKYKFVRIQPGQTFSVLACYNGSPSGVYQCAMRPNHTIKGSFLNGS 3382
                                   *:*:*****:*****************************************.**********
```

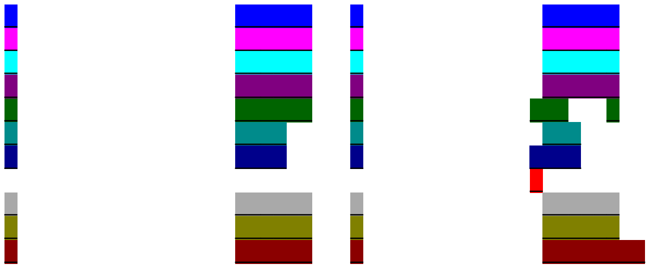

```
QHN73794                         CGSVGFNIDYDCVSFCYMHHMELPTGVHAGTDLEGNFYGPFVDRQTAQAAGTDTTITVNV 3467
SP|P0C6X7|R1AB_CVHSA             CGSVGFNIDYDCVSFCYMHHMELPTGVHAGTDLEGKFYGPFVDRQTAQAAGTDTTITLNV 3444
TR|Q6UZF5|Q6UZF5_CVHSA           CGSVGFNIDYDCVSFCYMHHMELPTGVHAGTDLEGKFYGPFVDRQTAQAAGTDTTITLNV 3444
TR|Q6UZF1|Q6UZF1_CVHSA           CGSVGFNIDYDCVSFCYMHHMELPTGVHAGTDLEGKFYGPFVDRQTAQAAGTDTTITLNV 3444
TR|Q6JH48|Q6JH48_CVHSA           CGSVGFNIDYDCVSFCYMHHMELPTGVHAGTDLEGKFYGPFVDRQTAQAAGTDTTITLNV 3444
TR|Q692E6|Q692E6_CVHSA           CGSVGFNIDYDCVSFCYMHHMELPTGVHAGTDLEGKFYGPFVDRQTAQAAGTDTTITLNV 3444
TR|A0A0K1YZY7|A0A0K1YZY7_CVHSA   CGSVGFNIDYDCVSFCYMHHMELPTGVHAGTDLEGKFYGPFVDRQTAQAAGTDTTITLNV 3444
SP|P0C6W2|R1AB_BCHK3             CGSVGFNIDYDCVSFCYMHHMELPTGVHAGTDLEGKFYGPFVDRQTAQAAGTDTTITLNV 3438
SP|P0C6W6|R1AB_BCRP3             CGSVGFNIDYDCVSFCYMHHMELPTEVHAGTDLEGKFYGPFVDRQTAQAAGTDTTITLNV 3442
SP|P0C6V9|R1AB_BC279             CGSVGFNIDYDCVSFCYMHHMELPTGVHAGTDLEGKFYGPFVDRQTAQAAGTDTTITLNV 3450
TR|A0A0U1WHI4|A0A0U1WHI4_CVHSA   CGSVGFNIDYDCVSFCYMHHMELPTGVHAGTDLEGKFYGPFVDRQTAQAAGTDTTITLNV 3439
TR|A0A0U1WHG0|A0A0U1WHG0_CVHSA   CGSVGFNIDYDCVSFCYMHHMELPTGVHAGTDLEGKFYGPFVDRQTAQAAGTDTTITLNV 3439
TR|A0A166ZL34|A0A166ZL34_9NIDO   CGSVGFNIDYDCVSFCYMHHMELPTGVHAGTDLEGKFYGPFVDRQTAQAAGTDTAITLNV 3246
TR|R9QTB2|R9QTB2_CVHSA           CGSVGFNIDYDCVSFCYMHHMELPTGVHAGTDLEGKFYGPFVDRQTAQAAGTDTTITLNV 3436
TR|R9QTH2|R9QTH2_CVHSA           CGSVGFNIDYDCVSFCYMHHMELPTGVHAGTDLEGKFYGPFVDRQTAQAAGTDTTITLNV 3445
SP|P0C6U8|R1A_CVHSA              CGSVGFNIDYDCVSFCYMHHMELPTGVHAGTDLEGKFYGPFVDRQTAQAAGTDTTITLNV 3444
TR|Q6JH47|Q6JH47_CVHSA           CGSVGFNIDYDCVSFCYMHHMELPTGVHAGTDLEGKFYGPFVDRQTAQAAGTDTTITLNV 3444
TR|Q692E5|Q692E5_CVHSA           CGSVGFNIDYDCVSFCYMHHMELPTGVHAGTDLEGKFYGPFVDRQTAQAAGTDTTITLNV 3444
SP|P0C6F8|R1A_BCHK3              CGSVGFNIDYDCVSFCYMHHMELPTGVHAGTDLEGKFYGPFVDRQTAQAAGTDTTITLNV 3438
TR|A0A0K1Z0N1|A0A0K1Z0N1_CVHSA   CGSVGFNIDYDCVSFCYMHHMELPTGVHAGTDLEGKFYGPFVDRQTAQAAGTDTTITLNV 3444
SP|P0C6F5|R1A_BC279              CGSVGFNIDYDCVSFCYMHHMELPTGVHAGTDLEGKFYGPFVDRQTAQAAGTDTTITLNV 3450
SP|P0C6T7|R1A_BCRP3              CGSVGFNIDYDCVSFCYMHHMELPTEVHAGTDLEGKFYGPFVDRQTAQAAGTDTTITLNV 3442
                                 ************************* *********:******************:**:**


QHN73794                         LAWLYAAVINGDRWFLNRFTTTLNDFNLVAMKYNYEPLTQDHVDILGPLSAQTGIAVLDM 3527
SP|P0C6X7|R1AB_CVHSA             LAWLYAAVINGDRWFLNRFTTTLNDFNLVAMKYNYEPLTQDHVDILGPLSAQTGIAVLDM 3504
TR|Q6UZF5|Q6UZF5_CVHSA           LAWLYAAVINGDRWFLNRFTTTLNDFNLVAMKYNYEPLTQDHVDILGPLSAQTGIAVLDM 3504
TR|Q6UZF1|Q6UZF1_CVHSA           LAWLYAAVINGDRWFLNRFTTTLNDFNLVAMKYNYEPLTQDHVDILGPLSAQTGIAVLDM 3504
TR|Q6JH48|Q6JH48_CVHSA           LAWLYAAVINGDRWFLNRFTTTLNDFNLVAMKYNYEPLTQDHVDILGPLSAQTGIAVLDM 3504
TR|Q692E6|Q692E6_CVHSA           LAWLYAAVINGDRWFLNRFTTTLNDFNLVAMKYNYEPLTQDHVDILGPLSAQTGIAVLDM 3504
TR|A0A0K1YZY7|A0A0K1YZY7_CVHSA   LAWLYAAVINGDRWFLNRFTTTLNDFNLVAMKYNYEPLTQDHVDVLGPLSAQTGIAVLDM 3504
SP|P0C6W2|R1AB_BCHK3             LAWLYAAVINGDRWFLNRFTTTLNDFNLVAMKYNYEPLTQDHVDILGPLSAQTGIAVLDM 3498
SP|P0C6W6|R1AB_BCRP3             LAWLYAAVINGDRWFLNRFTTTLNDFNLVAMKYNYEPLTQDHVDILGPLSAQTGIAVLDM 3502
SP|P0C6V9|R1AB_BC279             LAWLYAAVINGDRWFLNRFTTTLNDFNLVAMKYNYEPLTQDHVDILGPLSAQTGIAVLDM 3510
TR|A0A0U1WHI4|A0A0U1WHI4_CVHSA   LAWLYAAVINGDRWFLNRFTTTLNDFNLVAMKYNYEPLTQDHVDILGPLSAQTGIAVLDM 3499
TR|A0A0U1WHG0|A0A0U1WHG0_CVHSA   LAWLYAAVINGDRWFLNRFTTTLNDFNLVAMKYNYEPLTQDHVDILGPLSAQTGIAVLDM 3499
TR|A0A166ZL34|A0A166ZL34_9NIDO   LAWLYAAVINGDRWFLNRFTTTLNDFNLVAMKYNYEPLTQDHVDILGPLSAQTGIAVLDM 3306
TR|R9QTB2|R9QTB2_CVHSA           LAWLYAAVINGDRWFLNRFTTTLNDFNLVAMKYNYEPLTQDHVDILGPLSAQTGIAVLDM 3496
TR|R9QTH2|R9QTH2_CVHSA           LAWLYAAVINGDRWFLNRFTTTLNDFNLVAMKYNYEPLTQDHVDILGPLSAQTGIAVLDM 3505
SP|P0C6U8|R1A_CVHSA              LAWLYAAVINGDRWFLNRFTTTLNDFNLVAMKYNYEPLTQDHVDILGPLSAQTGIAVLDM 3504
TR|Q6JH47|Q6JH47_CVHSA           LAWLYAAVINGDRWFLNRFTTTLNDFNLVAMKYNYEPLTQDHVDILGPLSAQTGIAVLDM 3504
TR|Q692E5|Q692E5_CVHSA           LAWLYAAVINGDRWFLNRFTTTLNDFNLVAMKYNYEPLTQDHVDILGPLSAQTGIAVLDM 3504
SP|P0C6F8|R1A_BCHK3              LAWLYAAVINGDRWFLNRFTTTLNDFNLVAMKYNYEPLTQDHVDILGPLSAQTGIAVLDM 3498
TR|A0A0K1Z0N1|A0A0K1Z0N1_CVHSA   LAWLYAAVINGDRWFLNRFTTTLNDFNLVAMKYNYEPLTQDHVDVLGPLSAQTGIAVLDM 3504
SP|P0C6F5|R1A_BC279              LAWLYAAVINGDRWFLNRFTTTLNDFNLVAMKYNYEPLTQDHVDILGPLSAQTGIAVLDM 3510
SP|P0C6T7|R1A_BCRP3              LAWLYAAVINGDRWFLNRFTTTLNDFNLVAMKYNYEPLTQDHVDILGPLSAQTGIAVLDM 3502
                                 ********************************************:***************
```
