## Supplementary material for "Structural genomics and interactomics of 2019 Wuhan novel coronavirus, 2019-nCoV, indicate evolutionary conserved functional regions of viral proteins": Ligand binding site mapping for wNsp14

GUANOSINE-P3-ADENOSINE-5',5'-TRIPHOSPHATE (5C8S-G3A, SARS)

S-ADENOSYL-L-HOMOCYSTEINE (5C8S-SAH, SARS)

S-ADENOSYLMETHIONINE (5C8T-SAM, SARS)

```
QHN73794                      VIFTQTTETAHSCNVNRFNVAITRAKVGILCIMSDRDLYDKLQFTSLEIPRRNVATLQAE 5927
SP|P0C6X7|R1AB_CVHSA          VIFTQTTETAHSCNVNRFNVAITRAKIGILCIMSDRDLYDKLQFTSLEIPRRNVATLQAE 5904
TR|Q6UZF5|Q6UZF5_CVHSA        VIFTQTTETAHSCNVNRFNVAITRAKIGILCIMSDRDLYDKLQFTSLEIPRRNVATLQAE 5904
TR|Q6UZF1|Q6UZF1_CVHSA        VIFTQTTETAHSCNVNRFNVAITRAKIGILCIMSDRDLYDKLQFTSLEIPRRNVATLQAE 5904
TR|Q6JH48|Q6JH48_CVHSA        VIFTQTTETAHSCNVNRFNVAITRAKIGILCIMSDRDLYDKLQFTSLEIPRRNVATLQAE 5904
TR|Q692E6|Q692E6_CVHSA        VIFTQTTETAHSCNVNRFNVAITRAKIGILCIMSDRDLYDKLQFTSLEIPRRNVATLQAE 5904
TR|A0A0K1YZY7|A0A0K1YZY7_CVHSA VIFTQTTETAHSCNVNRFNVAITRAKIGILCIMSDRDLYDKLQFTSLEVPRRNVATLQAE 5904
SP|P0C6W2|R1AB_BCHK3          VIFTQTTETAHSCNVNRFNVAITRAKIGILCIMSDRDLYDKLQFTSLEVPRRNVATLQAE 5898
SP|P0C6W6|R1AB_BCRP3          VIFTQTTETAHSCNVNRFNVAITRAKIGILCIMSDRDLYDKLQFTSLEVPRRNVATLQAE 5902
SP|P0C6V9|R1AB_BC279          VIFTQTTETAHSCNVNRFNVAITRAKIGILCIMSDRDLYDKLQFTSLEVPRRNVATLQAE 5910
TR|A0A0U1WHI4|A0A0U1WHI4_CVHSA VIFTQTTETAHSCNVNRFNVAITRAKIGILCIMSDRDLYDKLQFTSLEVPRRNVATLQAE 5899
TR|A0A0U1WHG0|A0A0U1WHG0_CVHSA VIFTQTTETAHSCNVNRFNVAITRAKIGILCIMSDRDLYDKLQFTSLEVPRRNVATLQAE 5899
TR|A0A166ZL34|A0A166ZL34_9NIDO VIFTQTTETAHSCNVNRFNVAITRAKIGILCIMSDRDLYDKLQFTSLEVPRRNVATLQAE 5706
TR|R9QTB2|R9QTB2_CVHSA        ------------------------------------------------------------
TR|R9QTH2|R9QTH2_CVHSA        ------------------------------------------------------------
SP|P0C6U8|R1A_CVHSA           ------------------------------------------------------------
TR|Q6JH47|Q6JH47_CVHSA        ------------------------------------------------------------
TR|Q692E5|Q692E5_CVHSA        ------------------------------------------------------------
SP|P0C6F8|R1A_BCHK3           ------------------------------------------------------------
TR|A0A0K1Z0N1|A0A0K1Z0N1_CVHSA ------------------------------------------------------------
SP|P0C6F5|R1A_BC279           ------------------------------------------------------------
SP|P0C6T7|R1A_BCRP3           ------------------------------------------------------------


QHN73794                      NVTGLFKDCSKVITGLHPTQAPTHLSVDTKFKTEGLCVDIPGIPKDMTYRRLISMMGFKM 5987
SP|P0C6X7|R1AB_CVHSA          NVTGLFKDCSKIITGLHPTQAPTHLSVDIKFKTEGLCVDIPGIPKDMTYRRLISMMGFKM 5964
TR|Q6UZF5|Q6UZF5_CVHSA        NVTGLFKDCSKIITGLHPTQAPTHLSVDIKFKTEGLCVDIPGIPKDMTYRRLISMMGFKM 5964
TR|Q6UZF1|Q6UZF1_CVHSA        NVTGLFKDCSKIITGLHPTQAPTHLSVDIKFKTEGLCVDIPGIPKDMTYRRLISMMGFKM 5964
TR|Q6JH48|Q6JH48_CVHSA        NVTGLFKDCSKIITGLHPTQAPTHLSVDIKFKTEGLCVDIPGIPKDMTYRRLISMMGFKM 5964
TR|Q692E6|Q692E6_CVHSA        NVTGLFKDCSKIITGLHPTQAPTHLSVDIKFKTEGLCVDIPGIPKDMTYRRLISMMGFKM 5964
TR|A0A0K1YZY7|A0A0K1YZY7_CVHSA NVTGLFKDCSKIITGLHPTQAPTHLSVDTKFKTEGLCVDIPGIPKDMTYRRLISMMGFKM 5964
SP|P0C6W2|R1AB_BCHK3          NVTGLFKDCSKIITGLHPTQAPTHLSVDTKFKTEGLCVDIPGIPKDMTYRRLISMMGFKM 5958
SP|P0C6W6|R1AB_BCRP3          NVTGLFKDCSKIITGLHPTQAPTHLSVDTKFKTEGLCVDIPGIPKDMTYRRLISMMGFKM 5962
SP|P0C6V9|R1AB_BC279          NVTGLFKDCSKIITGLHPTQAPTHLSVDTKFKTEGLCVDIPGIPKDMTYRRLISMMGFKM 5970
TR|A0A0U1WHI4|A0A0U1WHI4_CVHSA NVTGLFKDCSKIITGLHPTQAPTHLSVDTKFKTEGLCVDIPGIPKDMTYRRLISMMGFKM 5959
TR|A0A0U1WHG0|A0A0U1WHG0_CVHSA NVTGLFKDCSKIITGLHPTQAPTHLSVDTKFKTEGLCVDIPGIPKDMTYRRLISMMGFKM 5959
TR|A0A166ZL34|A0A166ZL34_9NIDO NVTGLFKDCSKIITGLHPTQAPTHLSVDTKFKTEGLCVDIPGIPKDMTYRRLISMMGFKM 5766
TR|R9QTB2|R9QTB2_CVHSA        ------------------------------------------------------------
TR|R9QTH2|R9QTH2_CVHSA        ------------------------------------------------------------
SP|P0C6U8|R1A_CVHSA           ------------------------------------------------------------
TR|Q6JH47|Q6JH47_CVHSA        ------------------------------------------------------------
TR|Q692E5|Q692E5_CVHSA        ------------------------------------------------------------
SP|P0C6F8|R1A_BCHK3           ------------------------------------------------------------
TR|A0A0K1Z0N1|A0A0K1Z0N1_CVHSA ------------------------------------------------------------
SP|P0C6F5|R1A_BC279           ------------------------------------------------------------
SP|P0C6T7|R1A_BCRP3           ------------------------------------------------------------


QHN73794                      NYQVNGYPNMFITREEAIRHVRAWIGFDVEGCHATREAVGTNLPLQLGFSTGVNLVAVPT 6047
SP|P0C6X7|R1AB_CVHSA          NYQVNGYPNMFITREEAIRHVRAWIGFDVEGCHATRDAVGTNLPLQLGFSTGVNLVAVPT 6024
TR|Q6UZF5|Q6UZF5_CVHSA        NYQVNGYPNMFITREEAIRHVRAWIGFDVEGCHATRDAVGTNLPLQLGFSTGVNLVAVPT 6024
TR|Q6UZF1|Q6UZF1_CVHSA        NYQVNGYPNMFITREEAIRHVRAWIGFDVEGCHATRDAVGTNLPLQLGFSTGVNLVAVPT 6024
TR|Q6JH48|Q6JH48_CVHSA        NYQVNGYPNMFITREEAIRHVRAWIGFDVEGCHATRDAVGTNLPLQLGFSTGVNLVAVPT 6024
TR|Q692E6|Q692E6_CVHSA        NYQVNGYPNMFITREEAIRHVRAWIGFDVEGCHATRDAVGTNLPLQLGFSTGVNLVAVPT 6024
TR|A0A0K1YZY7|A0A0K1YZY7_CVHSA NYQVNGYPNMFITREEAIRHVRAWIGFDVEGCHATRDAVGTNLPLQLGFSTGVNLVAVPT 6024
SP|P0C6W2|R1AB_BCHK3          NYQVNGYPNMFITREEAIRHVRAWIGFDVEGCHATRDAVGTNLPLQLGFSTGVNLVAVPT 6018
SP|P0C6W6|R1AB_BCRP3          NYQVNGYPNMFITREEAIRHVRAWIGFDVEGCHATRDAVGTNLPLQLGFSTGVNLVAIPT 6022
SP|P0C6V9|R1AB_BC279          NYQVNGYPNMFITREEAIRHVRAWIGFDVEGCHATRDAVGTNLPLQLGFSTGVNLVAVPT 6030
TR|A0A0U1WHI4|A0A0U1WHI4_CVHSA NYQVNGYPNMFITREEAIRHVRAWIGFDVEGCHATRDAVGTNLPLQLGFSTGVNLVAVPT 6019
TR|A0A0U1WHG0|A0A0U1WHG0_CVHSA NYQVNGYPNMFITREEAIRHVRAWIGFDVEGCHATRDAVGTNLPLQLGFSTGVNLVAVPT 6019
TR|A0A166ZL34|A0A166ZL34_9NIDO NYQVNGYPNMFITREEAIRHVRAWIGFDVEGCHATRDAVGTNLPLQLGFSTGVNLVAVPT 5826
TR|R9QTB2|R9QTB2_CVHSA        ------------------------------------------------------------
TR|R9QTH2|R9QTH2_CVHSA        ------------------------------------------------------------
SP|P0C6U8|R1A_CVHSA           ------------------------------------------------------------
TR|Q6JH47|Q6JH47_CVHSA        ------------------------------------------------------------
TR|Q692E5|Q692E5_CVHSA        ------------------------------------------------------------
SP|P0C6F8|R1A_BCHK3           ------------------------------------------------------------
TR|A0A0K1Z0N1|A0A0K1Z0N1_CVHSA ------------------------------------------------------------
SP|P0C6F5|R1A_BC279           ------------------------------------------------------------
SP|P0C6T7|R1A_BCRP3           ------------------------------------------------------------
```

```
QHN73794                          GYVDTPNNTDFSRVSAKPPPGDQFKHLIPLMYKGLPWNVVRIKIVQMLSDTLKNLSDRVV 6107
SP|P0C6X7|R1AB_CVHSA              GYVDTENNTEFTRVNAKPPPGDQFKHLIPLMYKGLPWNVVRIKIVQMLSDTLKGLSDRVV 6084
TR|Q6UZF5|Q6UZF5_CVHSA            GYVDTENNTEFTRVNAKPPPGDQFKHLIPLMYKGLPWNVVRIKIVQMLSDTLKGLSDRVV 6084
TR|Q6UZF1|Q6UZF1_CVHSA            GYVDTENNTEFTRVNAKPPPGDQFKHLIPLMYKGLPWNVVRIKIVQMLSDTLKGLSDRVV 6084
TR|Q6JH48|Q6JH48_CVHSA            GYVDTENNTEFTRVNAKPPPGDQFKHLIPLMYKGLPWNVVRIKIVQMLSDTLKGLSDRVV 6084
TR|Q692E6|Q692E6_CVHSA            GYVDTENNTEFTRVNAKPPPGDQFKHLIPLMYKGLPWNVVRIKIVQMLSDTLKGLSDRVV 6084
TR|A0A0K1YZY7|A0A0K1YZY7_CVHSA    GYVDTENNTEFTRVNAKPPPGDQFKHLIPLMYKGLPWNVVRIKIVQMLSDTLKGLSDRVV 6084
SP|P0C6W2|R1AB_BCHK3              GYVDTENSTEFTRVNAKPPPGDQFKHLIPLMYKGLPWNVVRIKIVQMLSDTLKGLSDRVV 6078
SP|P0C6W6|R1AB_BCRP3              GYVDTENNTEFTRVNAKPPPGDQFKHLIPLMYKGLPWNVVRIKIVQMLSDTLKGLSDRVV 6082
SP|P0C6V9|R1AB_BC279              GYVDTENNTEFTRVNAKPPPGDQFKHLIPLMYKGLPWNVVRIKIVQMLSDTLKGLSDRVV 6090
TR|A0A0U1WHI4|A0A0U1WHI4_CVHSA    GYVDTENNTEFTRVNAKPPPGDQFKHLIPLMYKGLPWNVVRIKIVQMLSDTLKGLSDRVV 6079
TR|A0A0U1WHG0|A0A0U1WHG0_CVHSA    GYVDTENNTEFTRVNAKPPPGDQFKHLIPLMYKGLPWSVVRIKIVQMLSDTLKGLSDRVV 6079
TR|A0A166ZL34|A0A166ZL34_9NIDO    GYVDTENNTEFTRVNAKPPPGDQFKHLIPLMYKGLPWSVVRIKIVQMLSDTLKGLSDRVV 5886
TR|R9QTB2|R9QTB2_CVHSA            ------------------------------------------------------------
TR|R9QTH2|R9QTH2_CVHSA            ------------------------------------------------------------
SP|P0C6U8|R1A_CVHSA               ------------------------------------------------------------
TR|Q6JH47|Q6JH47_CVHSA            ------------------------------------------------------------
TR|Q692E5|Q692E5_CVHSA            ------------------------------------------------------------
SP|P0C6F8|R1A_BCHK3               ------------------------------------------------------------
TR|A0A0K1Z0N1|A0A0K1Z0N1_CVHSA    ------------------------------------------------------------
SP|P0C6F5|R1A_BC279               ------------------------------------------------------------
SP|P0C6T7|R1A_BCRP3               ------------------------------------------------------------


QHN73794                          FVLWAHGFELTSMKYFVKIGPERTCCLCDRRATCFSTASDTYACWHHSIGFDYVYNPFMI 6167
SP|P0C6X7|R1AB_CVHSA              FVLWAHGFELTSMKYFVKIGPERTCCLCDKRATCFSTSSDTYACWNHSVGFDYVYNPFMI 6144
TR|Q6UZF5|Q6UZF5_CVHSA            FVLWAHGFELTSMKYFVKIGPERTCCLCDKRATCFSTSSDTYACWNHSVGFDYVYNPFMI 6144
TR|Q6UZF1|Q6UZF1_CVHSA            FVLWAHGFELTSMKYFVKIGPERTCCLCDKRATCFSTSSDTYACWNHSVGFDYVYNPFMI 6144
TR|Q6JH48|Q6JH48_CVHSA            FVLWAHGFELTSMKYFVKIGPERTCCLCDKRATCFSTSSDTYACWNHSVGFDYVYNPFMI 6144
TR|Q692E6|Q692E6_CVHSA            FVLWAHGFELTSMKYFVKIGPERTCCLCDKRATCFSTSSDTYACWNHSVGFDYVYNPFMI 6144
TR|A0A0K1YZY7|A0A0K1YZY7_CVHSA    FVLWAHGFELTSMKYFVKIGPERTCCLCDKRATCFSTSSDTYACWNHSVGFDYVYNPFMI 6144
SP|P0C6W2|R1AB_BCHK3              FVLWAHGFELTSMKYFVKIGPERTCCLCDKRATCFSTSSDTYACWNHSVGFDYVYNPFMI 6138
SP|P0C6W6|R1AB_BCRP3              FVLWAHGFELTSMKYFVKIGPERTCCLCDKRATCFSTSSDTYACWNHSVGFDYVYNPFMI 6142
SP|P0C6V9|R1AB_BC279              FVLWAHGFELTSMKYFVKIGPERTCCLCDRRATCFSTSSDTYACWNHSVGFDYVYNPFMI 6150
TR|A0A0U1WHI4|A0A0U1WHI4_CVHSA    FVLWAHGFELTSMKYFVKIGPERTCCLCDKRATCFSTSSDTYACWNHSVGFDYVYNPFMI 6139
TR|A0A0U1WHG0|A0A0U1WHG0_CVHSA    FVLWAHGFELTSMKYFVKIGSERTCCLCDKRATCFSTSSDTYACWNHSVGFDYVYNPFMI 6139
TR|A0A166ZL34|A0A166ZL34_9NIDO    FVLWAHGFELTSMKYFVKIGSERTCCLCDKRATCFSTSSDTYACWNHSVGFDYVYNPFMI 5946
TR|R9QTB2|R9QTB2_CVHSA            ------------------------------------------------------------
TR|R9QTH2|R9QTH2_CVHSA            ------------------------------------------------------------
SP|P0C6U8|R1A_CVHSA               ------------------------------------------------------------
TR|Q6JH47|Q6JH47_CVHSA            ------------------------------------------------------------
TR|Q692E5|Q692E5_CVHSA            ------------------------------------------------------------
SP|P0C6F8|R1A_BCHK3               ------------------------------------------------------------
TR|A0A0K1Z0N1|A0A0K1Z0N1_CVHSA    ------------------------------------------------------------
SP|P0C6F5|R1A_BC279               ------------------------------------------------------------
SP|P0C6T7|R1A_BCRP3               ------------------------------------------------------------


QHN73794                          DVQQWGFTGNLQSNHDLYCQVHGNAHVASCDAIMTRCLAVHECFVKRVDWTIEYPIIGDE 6227
SP|P0C6X7|R1AB_CVHSA              DVQQWGFTGNLQSNHDQHCQVHGNAHVASCDAIMTRCLAVHECFVKRVDWSVEYPIIGDE 6204
TR|Q6UZF5|Q6UZF5_CVHSA            DVQQWGFTGNLQSNHDQHCQVHGNAHVASCDAIMTRCLAVHECFVKRVDWSVEYPIIGDE 6204
TR|Q6UZF1|Q6UZF1_CVHSA            DVQQWGFTGNLQSNHDQHCQVHGNAHVASCDAIMTRCLAVHECFVKRVDWSVEYPIIGDE 6204
TR|Q6JH48|Q6JH48_CVHSA            DVQQWGFTGNLQSNHDQHCQVHGNAHVASCDAIMTRCLAVHECFVKRVDWSVEYPIIGDE 6204
TR|Q692E6|Q692E6_CVHSA            DVQQWGFTGNLQSNHDQHCQVHGNAHVASCDAIMTRCLAVHECFVKRVDWSVEYPIIGDE 6204
TR|A0A0K1YZY7|A0A0K1YZY7_CVHSA    DVQQWGFTGNLQSNHDQHCQVHGNAHVASCDAIMTRCLAVHECFVKRVDWSVEYPIIGDE 6204
SP|P0C6W2|R1AB_BCHK3              DVQQWGFTGNLQSNHDQHCQVHGNAHVASCDAIMTRCLAVHECFVKRVDWSVEYPIIGDE 6198
SP|P0C6W6|R1AB_BCRP3              DVQQWGFTGNLQSNHDQHCQVHGNAHVASCDAIMTRCLAVHECFVKRVDWSVEYPIIGDE 6202
SP|P0C6V9|R1AB_BC279              DVQQWGFTGNLQSNHDQHCQVHGNAHVASCDAIMTRCLAVHECFVKRVDWSVEYPIIGDE 6210
TR|A0A0U1WHI4|A0A0U1WHI4_CVHSA    DVQQWGLTGNLQSNHDQHCQVHGNAHVASCDAIMTRCLAVHECFVKRVDWSVEYPIIGDE 6199
TR|A0A0U1WHG0|A0A0U1WHG0_CVHSA    DVQQWGFTGNLQSNHDQHCQVHGNAHVASCDAIMTRCLAVHECFVKRVDWSVEYPVIGDE 6199
TR|A0A166ZL34|A0A166ZL34_9NIDO    DVQQWGFTGNLQSNHDQHCQVHGNAHVASCDAIMTRCLAVHECFVKRVDWSVEYPVIGDE 6006
TR|R9QTB2|R9QTB2_CVHSA            ------------------------------------------------------------
TR|R9QTH2|R9QTH2_CVHSA            ------------------------------------------------------------
SP|P0C6U8|R1A_CVHSA               ------------------------------------------------------------
TR|Q6JH47|Q6JH47_CVHSA            ------------------------------------------------------------
TR|Q692E5|Q692E5_CVHSA            ------------------------------------------------------------
SP|P0C6F8|R1A_BCHK3               ------------------------------------------------------------
TR|A0A0K1Z0N1|A0A0K1Z0N1_CVHSA    ------------------------------------------------------------
SP|P0C6F5|R1A_BC279               ------------------------------------------------------------
SP|P0C6T7|R1A_BCRP3               ------------------------------------------------------------
```

```
QHN73794                              LKINAACRKVQHMVVKAALLADKFPVLHDIGNPKAIKCVPQADVEWKFYDAQPCSDKAYK 6287
SP|P0C6X7|R1AB_CVHSA                  LRVNSACRKVQHMVVKSALLADKFPVLHDIGNPKAIKCVPQAEVEWKFYDAQPCSDKAYK 6264
TR|Q6UZF5|Q6UZF5_CVHSA                LRVNSACRKVQHMVVKSALLADKFPVLHDIGNPKAIKCVPQAEVEWKFYDAQPCSDKAYK 6264
TR|Q6UZF1|Q6UZF1_CVHSA                LRVNSACRKVQHMVVKSALLADKFPVLHDIGNPKAIKCVPQAEVEWKFYDAQPCSDKAYK 6264
TR|Q6JH48|Q6JH48_CVHSA                LRVNSACRKVQHMVVKSALLADKFPVLHDIGNPKAIKCVPQAEVEWKFYDAQPCSDKAYK 6264
TR|Q692E6|Q692E6_CVHSA                LRVNSACRKVQHMVVKSALLADKFPVLHDIGNPKAIKCVPQAEVEWKFYDAQPCSDKAYK 6264
TR|A0A0K1YZY7|A0A0K1YZY7_CVHSA        LKINSACRKVQHMVVKSALLADKFPVLHDIGNPKAIKCVPQAEVEWKFYDAQPCSDKAYK 6264
SP|P0C6W2|R1AB_BCHK3                  LKINAACRKVQHMVVKSALLADKFTVLHDIGNPKAIRCVPQAEVDWKFYDAQPCSDKAYK 6258
SP|P0C6W6|R1AB_BCRP3                  LKINSACRKVQHMVVKSALLADKFPVLHDIGNPKAIKCVPQAEVEWKFYDAQPCSDKAYK 6262
SP|P0C6V9|R1AB_BC279                  LKINAACRKVQHMVVKSALLADKFSVLHDIGNPKAIKCVPQAEVDWKFYDAQPCSDKAYK 6270
TR|A0A0U1WHI4|A0A0U1WHI4_CVHSA        LKINAACRKVQHMVVKSALLADKFPVLHDIGNPKAIKCVPQADVEWKFYDAQPCSDKAYK 6259
TR|A0A0U1WHG0|A0A0U1WHG0_CVHSA        LKINAACRKVQHMVVKSALLADKFPVLHDIGNPKAIKCVPQADVEWKFYDVQPCSDKAYK 6259
TR|A0A166ZL34|A0A166ZL34_9NIDO        LKINAACRKVQHMVVKSALLADKFPVLHDIGNPKAIKCVPQADVEWKFYDVQPCSDKAYK 6066
TR|R9QTB2|R9QTB2_CVHSA                ------------------------------------------------------------
TR|R9QTH2|R9QTH2_CVHSA                ------------------------------------------------------------
SP|P0C6U8|R1A_CVHSA                   ------------------------------------------------------------
TR|Q6JH47|Q6JH47_CVHSA                ------------------------------------------------------------
TR|Q692E5|Q692E5_CVHSA                ------------------------------------------------------------
SP|P0C6F8|R1A_BCHK3                   ------------------------------------------------------------
TR|A0A0K1Z0N1|A0A0K1Z0N1_CVHSA        ------------------------------------------------------------
SP|P0C6F5|R1A_BC279                   ------------------------------------------------------------
SP|P0C6T7|R1A_BCRP3                   ------------------------------------------------------------

QHN73794                              IEELFYSYATHSDKFTDGVCLFWNCNVDRYPANSIVCRFDTRVLSNLNLPGCDGGSLYVN 6347
SP|P0C6X7|R1AB_CVHSA                  IEELFYSYATHHDKFTDGVCLFWNCNVDRYPANAIVCRFDTRVLSNLNLPGCDGGSLYVN 6324
TR|Q6UZF5|Q6UZF5_CVHSA                IEELFYSYATHHDKFTDGVCLFWNCNVDRYPANAIVCRFDTRVLSNLNLPGCDGGSLYVN 6324
TR|Q6UZF1|Q6UZF1_CVHSA                IEELFYSYATHHDKFTDGVCLFWNCNVDRYPANAIVCRFDTRVLSNLNLPGCDGGSLYVN 6324
TR|Q6JH48|Q6JH48_CVHSA                IEELFYSYATHHDKFTDGVCLFWNCNVDRYPANAIVCRFDTRVLSNLNLPGCDGGSLYVN 6324
TR|Q692E6|Q692E6_CVHSA                IEELFYSYATHHDKFTDGVCLFWNCNVDRYPANAIVCRFDTRVLSNLNLPGCDGGSLYVN 6324
TR|A0A0K1YZY7|A0A0K1YZY7_CVHSA        IEELFYSYATHHDKFTDGVCLFWNCNVDRYPANAIVCRFDTRVLSNLNLPGCDGGSLYVN 6324
SP|P0C6W2|R1AB_BCHK3                  IEELFYSYATHHDKFTDGVCLFWNCNVDRYPANAIVCRFDTRVLSNLNLPGCDGGSLYVN 6318
SP|P0C6W6|R1AB_BCRP3                  IEELFYSYATHHDKFTDGVCLFWNCNVDRYPANAIVCRFDTRVLSNLNLPGCDGGSLYVN 6322
SP|P0C6V9|R1AB_BC279                  IEELFYSYATHHDKFTDGVCLFWNCNVDRYPANAIVCRFDTRVLSNLNLPGCDGGSLYVN 6330
TR|A0A0U1WHI4|A0A0U1WHI4_CVHSA        IEELFYSYATHHDKFTDGVCLFWNCNVDRYPANAIVCRFDTRVLSNLNLPGCDGGSLYVN 6319
TR|A0A0U1WHG0|A0A0U1WHG0_CVHSA        IEELFYSYATHHDKFTDGVCLFWNCNVDRYPANAIVCRFDTRVLSNLNLPGCDGGSLYVN 6319
TR|A0A166ZL34|A0A166ZL34_9NIDO        IEELFYSYATHHDKFTDGVCLFWNCNVDRYPANAIVCRFDTRVLSNLNLPGCDGGSLYVN 6126
TR|R9QTB2|R9QTB2_CVHSA                ------------------------------------------------------------
TR|R9QTH2|R9QTH2_CVHSA                ------------------------------------------------------------
SP|P0C6U8|R1A_CVHSA                   ------------------------------------------------------------
TR|Q6JH47|Q6JH47_CVHSA                ------------------------------------------------------------
TR|Q692E5|Q692E5_CVHSA                ------------------------------------------------------------
SP|P0C6F8|R1A_BCHK3                   ------------------------------------------------------------
TR|A0A0K1Z0N1|A0A0K1Z0N1_CVHSA        ------------------------------------------------------------
SP|P0C6F5|R1A_BC279                   ------------------------------------------------------------
SP|P0C6T7|R1A_BCRP3                   ------------------------------------------------------------

QHN73794                              KHAFHTPAFDKSAFVNLKQLPFFYYSDSPCESHGKQVVSDIDYVPLKSATCITRCNLGGA 6407
SP|P0C6X7|R1AB_CVHSA                  KHAFHTPAFDKSAFTNLKQLPFFYYSDSPCESHGKQVVSDIDYVPLKSATCITRCNLGGA 6384
TR|Q6UZF5|Q6UZF5_CVHSA                KHAFHTPAFDKSAFTNLKQLPFFYYSDSPCESHGKQVVSDIDYVPLKSATCITRCNLGGA 6384
TR|Q6UZF1|Q6UZF1_CVHSA                KHAFHTPAFDKSAFTNLKQLPFFYYSDSPCESHGKQVVSDIDYVPLKSATCITRCNLGGA 6384
TR|Q6JH48|Q6JH48_CVHSA                KHAFHTPAFDKSAFTNLKQLPFFYYSDSPCESHGKQVVSDIDYVPLKSATCITRCNLGGA 6384
TR|Q692E6|Q692E6_CVHSA                KHAFHTPAFDKSAFTNLKQLPFFYYSDSPCESHGKQVVSDIDYVPLKSATCITRCNLGGA 6384
TR|A0A0K1YZY7|A0A0K1YZY7_CVHSA        KHAFHTPAFDKSAFTNLKQLPFFYYSDSPCESHGKQVVSDIDYVPLKSATCITRCNLGGA 6384
SP|P0C6W2|R1AB_BCHK3                  KHAFHTPAFDKSAFTHLKQLPFFYYSDSPCESHGKQVVSDIDYVPLKSATCITRCNLGGA 6378
SP|P0C6W6|R1AB_BCRP3                  KHAFHTPAFDKSAFTNLKQLPFFYYSDSPCESHGKQVVSDIDYVPLKSATCITRCNLGGA 6382
SP|P0C6V9|R1AB_BC279                  KHAFHTPAFDKSAFTYLKQLPFFYYSDSPCESHGKQVVSDIDYVPLKSATCITRCNLGGA 6390
TR|A0A0U1WHI4|A0A0U1WHI4_CVHSA        KHAFHTPAFDKSAFSNLKQLPFFYYSDSPCESHGKQVVSDIDYVPLKSATCITRCNLGGA 6379
TR|A0A0U1WHG0|A0A0U1WHG0_CVHSA        KHAFHTPAFDKSAFTHLKQLPFFYYSDSPCESHGKQVVSDIDYVPLKSATCITRCNLGGA 6379
TR|A0A166ZL34|A0A166ZL34_9NIDO        KHAFYTPAFDKSAFTHLKQLPFFYYSDSPCESHGKQVVSDIDYVPLKSATCITRCNLGGA 6186
TR|R9QTB2|R9QTB2_CVHSA                ------------------------------------------------------------
TR|R9QTH2|R9QTH2_CVHSA                ------------------------------------------------------------
SP|P0C6U8|R1A_CVHSA                   ------------------------------------------------------------
TR|Q6JH47|Q6JH47_CVHSA                ------------------------------------------------------------
TR|Q692E5|Q692E5_CVHSA                ------------------------------------------------------------
SP|P0C6F8|R1A_BCHK3                   ------------------------------------------------------------
TR|A0A0K1Z0N1|A0A0K1Z0N1_CVHSA        ------------------------------------------------------------
SP|P0C6F5|R1A_BC279                   ------------------------------------------------------------
SP|P0C6T7|R1A_BCRP3                   ------------------------------------------------------------
```

```
QHN73794                           VCRHHANEYRLYLDAYNMMISAGFSLWVYKQFDTYNLWNTFTRLQSLENVAFNVVNKGHF 6467
SP|P0C6X7|R1AB_CVHSA               VCRHHANEYRQYLDAYNMMISAGFSLWIYKQFDTYNLWNTFTRLQSLENVAYNVVNKGHF 6444
TR|Q6UZF5|Q6UZF5_CVHSA             VCRHHANEYRQYLDAYNMMISAGFSLWIYKQFDTYNLWNTFTRLQSLENVAYNVVNKGHF 6444
TR|Q6UZF1|Q6UZF1_CVHSA             VCRHHANEYRQYLDAYNMMISAGFSLWIYKQFDTYNLWNTFTRLQSLENVAYNVVNKGHF 6444
TR|Q6JH48|Q6JH48_CVHSA             VCRHHANEYRQYLDAYNMMISAGFSLWIYKQFDTYNLWNTFTRLQSLENVAYNVVNKGHF 6444
TR|Q692E6|Q692E6_CVHSA             VCRHHANEYRQYLDAYNMMISAGFSLWIYKQFDTYNLWNTFTRLQSLENVAYNVVNKGHF 6444
TR|A0A0K1YZY7|A0A0K1YZY7_CVHSA VCRHHANEYRQYLDAYNMMISAGFSLWIYKQFDTYNLWNTFTRLQSLENVAYNVVNKGHF 6444
SP|P0C6W2|R1AB_BCHK3               VCRHHANEYRQYLDAYNMMISAGFSLWIYKQFDTYNLWNTFTKLQSLENVAYNVVNKGHF 6438
SP|P0C6W6|R1AB_BCRP3               VCRHHANEYRQYLDAYNMMISAGFSLWIYKQFDTYNLWNTFTRLQSLENVAYNVVNKGHF 6442
SP|P0C6V9|R1AB_BC279               VCRRHANEYRQYLDAYNMMISAGFSLWIYKQFDTYNLWNTFTRLQSLENVAYNVVNKGHF 6450
TR|A0A0U1WHI4|A0A0U1WHI4_CVHSA VCRHHANEYRQYLDAYNMMISAGFSLWIYKQFDTYNLWNTFTRLQSLENVAYNVINKGHF 6439
TR|A0A0U1WHG0|A0A0U1WHG0_CVHSA VCRHHANEYRQYLDAYNMMISAGFSLWIYKQFDTYNLWNTFTRLQSLENVAYNVINKGHF 6439
TR|A0A166ZL34|A0A166ZL34_9NIDO VCRHHANEYRQYLDAYNMMISAGFSLWIYKQFDTYNLWNTFTRLQSLENVAYNVINKGHF 6246
TR|R9QTB2|R9QTB2_CVHSA             ------------------------------------------------------------
TR|R9QTH2|R9QTH2_CVHSA             ------------------------------------------------------------
SP|P0C6U8|R1A_CVHSA                ------------------------------------------------------------
TR|Q6JH47|Q6JH47_CVHSA             ------------------------------------------------------------
TR|Q692E5|Q692E5_CVHSA             ------------------------------------------------------------
SP|P0C6F8|R1A_BCHK3                ------------------------------------------------------------
TR|A0A0K1Z0N1|A0A0K1Z0N1_CVHSA ------------------------------------------------------------
SP|P0C6F5|R1A_BC279                ------------------------------------------------------------
SP|P0C6T7|R1A_BCRP3                ------------------------------------------------------------
```
