## Supplementary material for "Structural genomics and interactomics of 2019 Wuhan novel coronavirus, 2019-nCoV, indicate evolutionary conserved functional regions of viral proteins": Ligand binding site mapping for wNsp16

Ligand BS:

- S-ADENOSYLMETHIONINE (5YN6-SAM, MERS)
- S-ADENOSYL-L-HOMOCYSTEINE (5YN8-SAH, MERS)
- SINEFUNGIN (5YNB-SFG, MERS)
- P1-7-METHYLGUANOSINE-P3-ADENOSINE-5',5'-TRIPHOSPHATE (5YNF-GTA, MERS)
- 7-METHYL-GUANOSINE-5'-TRIPHOSPHATE-5'-GUANOSINE (5YNI-GTG, MERS)

```
QHN73794                           KVVKVTIDYTEISFMLWCKDGHVETFYPKLQSSQAWQPGVAMPNLYKMQRMLLEKCDLQN 6827
SP|P0C6X7|R1AB_CVHSA               KVVKVTIDYAEISFMLWCKDGHVETFYPKLQASQAWQPGVAMPNLYKMQRMLLEKCDLQN 6804
TR|Q6UZF5|Q6UZF5_CVHSA             KVVKVTIDYAEISFMLWCKDGHVETFYPKLQASQAWQPGVAMPNLYKMQRMLLEKCDLQN 6804
TR|Q6UZF1|Q6UZF1_CVHSA             KVVKVTIDYAEISFMLWCKDGHVETFYPKLQASQAWQPGVAMPNLYKMQRMLLEKCDLQN 6804
TR|Q6JH48|Q6JH48_CVHSA             KVVKVTIDYAEISFMLWCKDGHVETFYPKLQASQAWQPGVAMPNLYKMQRMLLEKCDLQN 6804
TR|Q692E6|Q692E6_CVHSA             KVVKVTIDYAEISFMLWCKDGHVETFYPKLQASQAWQPGVAMPNLYKMQRMLLEKCDLQN 6804
TR|A0A0K1YZY7|A0A0K1YZY7_CVHSA     KVVKVTIDYVEISFMLWCKDGHVETFYPKLQASQAWQPGVAMPNLYKMQRMLLEKCDLQN 6804
SP|P0C6W2|R1AB_BCHK3               KVVKVTIDYAEISFMLWCKDGHVETFYPKLQASQAWQPGVAMPNLYKMQRMLLEKCDLQN 6798
SP|P0C6W6|R1AB_BCRP3               KVVKVTIDYAEISFMLWCKDGHVETFYPKLQASQAWQPGVAMPNLYKMQRMLLEKCDLQN 6802
SP|P0C6V9|R1AB_BC279               KVVKVTIDYAEISFMLWCKDGHVETFYPKLQASQAWQPGVAMPNLYKMQRMLLEKCDLQN 6810
TR|A0A0U1WHI4|A0A0U1WHI4_CVHSA     KVVKVTIDYAEISFMLWCKDGHVETFYPKLQASQAWQPGVAMPNLYKMQRMLLEKCDLQN 6799
TR|A0A0U1WHG0|A0A0U1WHG0_CVHSA     KVVKVTIDYAEISFMLWCKDGYVETFYPKLQASQAWQPGVAMPNLYKMQRMLLEKCDLQN 6799
TR|A0A166ZL34|A0A166ZL34_9NIDO     KVVKVTIDYAEISFMLWCKDGYVETFYPKLQASQAWQPGVAMPNLYKMQRMLLEKCDLQN 6606
TR|R9QTB2|R9QTB2_CVHSA             ------------------------------------------------------------
TR|R9QTH2|R9QTH2_CVHSA             ------------------------------------------------------------
SP|P0C6U8|R1A_CVHSA                ------------------------------------------------------------
TR|Q6JH47|Q6JH47_CVHSA             ------------------------------------------------------------
TR|Q692E5|Q692E5_CVHSA             ------------------------------------------------------------
SP|P0C6F8|R1A_BCHK3                ------------------------------------------------------------
TR|A0A0K1Z0N1|A0A0K1Z0N1_CVHSA     ------------------------------------------------------------
SP|P0C6F5|R1A_BC279                ------------------------------------------------------------
SP|P0C6T7|R1A_BCRP3                ------------------------------------------------------------
```

```
QHN73794                           YGDSATLPKGIMMNVAKYTQLCQYLNTLTLAVPYNMRVIHFGAGSDKGVAPGTAVLRQWL 6887
SP|P0C6X7|R1AB_CVHSA               YGENAVIPKGIMMNVAKYTQLCQYLNTLTLAVPYNMRVIHFGAGSDKGVAPGTAVLRQWL 6864
TR|Q6UZF5|Q6UZF5_CVHSA             YGENAVIPKGIMMNVAKYTQLCQYLNTLTLAVPYNMRVIHFGAGSDKGVAPGTAVLRQWL 6864
TR|Q6UZF1|Q6UZF1_CVHSA             YGENAVIPKGIMMNVAKYTQLCQYLNTLTLAVPYNMRVIHFGAGSDKGVAPGTAVLRQWL 6864
TR|Q6JH48|Q6JH48_CVHSA             YGENAVIPKGIMMNVAKYTQLCQYLNTLTLAVPYNMRVIHFGAGSDKGVAPGTAVLRQWL 6864
TR|Q692E6|Q692E6_CVHSA             YGENAVIPKGIMMNVAKYTQLCQYLNTLTLAVPYNMRVIHFGAGSDKGVAPGTAVLRQWL 6864
TR|A0A0K1YZY7|A0A0K1YZY7_CVHSA     YGENAVIPKGIMMNVAKYTQLCQYLNTLTLAVPYNMRVIHFGAGSDKGVAPGTAVLRQWL 6864
SP|P0C6W2|R1AB_BCHK3               YGENAVIPKGIMMNVAKYTQLCQYLNTLTLAVPYNMRVIHFGAGSDKGVAPGTAVLRQWL 6858
SP|P0C6W6|R1AB_BCRP3               YGENAVIPKGIMMNVAKYTQLCQYLNTLTLAVPYNMRVIHFGAGSDKGVAPGTAVLRQWL 6862
SP|P0C6V9|R1AB_BC279               YGENAVIPKGIMMNVAKYTQLCQYLNTLTLAVPYNMRVIHFGAGSDKGVAPGTAVLRQWL 6870
TR|A0A0U1WHI4|A0A0U1WHI4_CVHSA     YGENAVIPKGIMMNVAKYTQLCQYLNTLTLAVPYNMRVIHFGAGSDKGVAPGTAVLRQWL 6859
TR|A0A0U1WHG0|A0A0U1WHG0_CVHSA     YGENAVIPKGIMMNVAKYTQLCQYLNTLTLAVPYNMRVIHFGAGSDKGVAPGTAVLRQWL 6859
TR|A0A166ZL34|A0A166ZL34_9NIDO     YGENAVIPKGIMMNVAKYTQLCQYLNTLTLAVPYNMRVIHFGAGSDKGVAPGTAVLRQWL 6666
TR|R9QTB2|R9QTB2_CVHSA             ------------------------------------------------------------
TR|R9QTH2|R9QTH2_CVHSA             ------------------------------------------------------------
SP|P0C6U8|R1A_CVHSA                ------------------------------------------------------------
TR|Q6JH47|Q6JH47_CVHSA             ------------------------------------------------------------
TR|Q692E5|Q692E5_CVHSA             ------------------------------------------------------------
SP|P0C6F8|R1A_BCHK3                ------------------------------------------------------------
TR|A0A0K1Z0N1|A0A0K1Z0N1_CVHSA     ------------------------------------------------------------
SP|P0C6F5|R1A_BC279                ------------------------------------------------------------
SP|P0C6T7|R1A_BCRP3                ------------------------------------------------------------
```

```
QHN73794                        PTGTLLVDSDLNDFVSDADSTLIGDCATVHTANKWDLIISDMYDPKTKNVTKENDSKEGF 6947
SP|P0C6X7|R1AB_CVHSA            PTGTLLVDSDLNDFVSDADSTLIGDCATVHTANKWDLIISDMYDPRTKHVTKENDSKEGF 6924
TR|Q6UZF5|Q6UZF5_CVHSA          PTGTLLVDSDLNDFVSDADSTLIGDCATVHTANKWDLIISDMYDPRTKHVTKENDSKEGF 6924
TR|Q6UZF1|Q6UZF1_CVHSA          PTGTLLVDSDLNDFVSDADSTLIGDCATVHTANKWDLIISDMYDPRTKHVTKENDSKEGF 6924
TR|Q6JH48|Q6JH48_CVHSA          PTGTLLVDSDLNDFVSDADSTLIGDCATVHTANKWDLIISDMYDPRTKHVTKENDSKEGF 6924
TR|Q692E6|Q692E6_CVHSA          PTGTLLVDSDLNDFVSDADSTLIGDCATVHTANKWDLIISDMYDPRTKHVTKENDSKEGF 6924
TR|A0A0K1YZY7|A0A0K1YZY7_CVHSA  PTGTLLVDSDLNDFVSDADSTLIGDCATVHTANKWDLIISDMYDPKTKHVTKENDSKEGF 6924
SP|P0C6W2|R1AB_BCHK3            PTGTLLVDSDLNDFVSDADSTLIGDCATVHTANKWDLIISDMYDPKTKHVLKDNDSKEGF 6918
SP|P0C6W6|R1AB_BCRP3            PTGTLLVDSDLNDFVSDADSTLIGDCATVHTANKWDLIVSDMYDPKAKHVTKENDSKEGF 6922
SP|P0C6V9|R1AB_BC279            PTGALLVDSDLNDFVSDADSTLIGDCATVHTANKWDLIISDMYDPKTKHVTKENDSKEGF 6930
TR|A0A0U1WHI4|A0A0U1WHI4_CVHSA  PIGTLLVDSDLNDFVSDADSTLIGDCATVHTANKWDLIVSDMYDPKTKHVTEENDSKEGF 6919
TR|A0A0U1WHG0|A0A0U1WHG0_CVHSA  PIGTLLVDSDLNDFVSDADSTLIGECATVHTANKWDLIVSDMYDPKTKHVTKENDSKEGF 6919
TR|A0A166ZL34|A0A166ZL34_9NIDO  PIGTLLVDSDLNDFVSDADSTLIGECATVHTANKWDLIVSDMYDPKTKHVTKENDSKEGF 6726
TR|R9QTB2|R9QTB2_CVHSA          ------------------------------------------------------------
TR|R9QTH2|R9QTH2_CVHSA          ------------------------------------------------------------
SP|P0C6U8|R1A_CVHSA             ------------------------------------------------------------
TR|Q6JH47|Q6JH47_CVHSA          ------------------------------------------------------------
TR|Q692E5|Q692E5_CVHSA          ------------------------------------------------------------
SP|P0C6F8|R1A_BCHK3             ------------------------------------------------------------
TR|A0A0K1Z0N1|A0A0K1Z0N1_CVHSA  ------------------------------------------------------------
SP|P0C6F5|R1A_BC279             ------------------------------------------------------------
SP|P0C6T7|R1A_BCRP3             ------------------------------------------------------------

QHN73794                        FTYICGFIQQKLALGGSVAIKITEHSWNADLYKLMGHFAWWTAFVTNVNASSSEAFLIGC 7007
SP|P0C6X7|R1AB_CVHSA            FTYLCGFIKQKLALGGSIAVKITEHSWNADLYKLMGHFSWWTAFVTNVNASSSEAFLIGA 6984
TR|Q6UZF5|Q6UZF5_CVHSA          FTYLCGFIKQKLALGGSIAVKITEHSWNADLYKLMGHFSWWTAFVTNVNASSSEAFLIGA 6984
TR|Q6UZF1|Q6UZF1_CVHSA          FTYLCGFIKQKLALGGSIAVKITEHSWNADLYKLMGHFSWWTAFVTNVNASSSEAFLIGA 6984
TR|Q6JH48|Q6JH48_CVHSA          FTYLCGFIKQKLALGGSIAVKITEHSWNADLYKLMGHFSWWTAFVTNVNASSSEAFLIGA 6984
TR|Q692E6|Q692E6_CVHSA          FTYLCGFIKQKLALGGSIAVKITEHSWNADLYKLMGHFSWWTAFVTNVNASSSEAFLIGA 6984
TR|A0A0K1YZY7|A0A0K1YZY7_CVHSA  FTYLCGFIKQKLALGGSAAVKITEHSWNADLYKLMGHFSWWTAFVTNVNASSSEAFLIGV 6984
SP|P0C6W2|R1AB_BCHK3            FTYLCGFIKQKLALGGSVAVKITEHSWNADLYKLMGHFSWWTAFVTNVNASSSEAFLIGV 6978
SP|P0C6W6|R1AB_BCRP3            FTYLCGFIKQKLALGGSVAVKITEHSWNADLYKLMGHFSWWTAFVTNVNASSSEAFLIGV 6982
SP|P0C6V9|R1AB_BC279            FTYLCGFIKQKLALGGSVAVKITEHSWNADLYKLMGHFSWWTAFVTNVNASSSEAFLIGV 6990
TR|A0A0U1WHI4|A0A0U1WHI4_CVHSA  FTYLCGFIKQKLALGGSVAVKITEHSWNADLYKLMGYFSWWTAFVTNVNASSSEAFLIGV 6979
TR|A0A0U1WHG0|A0A0U1WHG0_CVHSA  FTYLCGFIKQKLALGGSVAVKITEHSWNADLYKLMGHFSWWTAFVTNVNASSSEAFLIGV 6979
TR|A0A166ZL34|A0A166ZL34_9NIDO  FTYLCGFIKQKLALGGSVAVKITEHSWNADLYKLMGHFSWWTAFVTNVNASSSEAFLIGV 6786
TR|R9QTB2|R9QTB2_CVHSA          ------------------------------------------------------------
TR|R9QTH2|R9QTH2_CVHSA          ------------------------------------------------------------
SP|P0C6U8|R1A_CVHSA             ------------------------------------------------------------
TR|Q6JH47|Q6JH47_CVHSA          ------------------------------------------------------------
TR|Q692E5|Q692E5_CVHSA          ------------------------------------------------------------
SP|P0C6F8|R1A_BCHK3             ------------------------------------------------------------
TR|A0A0K1Z0N1|A0A0K1Z0N1_CVHSA  ------------------------------------------------------------
SP|P0C6F5|R1A_BC279             ------------------------------------------------------------
SP|P0C6T7|R1A_BCRP3             ------------------------------------------------------------

QHN73794                        NYLGKPREQIDGYVMHANYIFWRNTNPIQLSSYSLFDMSKFPLKLRGTAVMSLKEGQIND 7067
SP|P0C6X7|R1AB_CVHSA            NYLGKPKEQIDGYTMHANYIFWRNTNPIQLSSYSLFDMSKFPLKLRGTAVMSLKENQIND 7044
TR|Q6UZF5|Q6UZF5_CVHSA          NYLGKPKEQIDGYTMHANYIFWRNTNPIQLSSYSLFDMSKFPLKLRGTAVMSLKENQIND 7044
TR|Q6UZF1|Q6UZF1_CVHSA          NYLGKPKEQIDGYTMHANYIFWRNTNPIQLSSYSLFDMSKFPLKLRGTAVMSLKENQIND 7044
TR|Q6JH48|Q6JH48_CVHSA          NYLGKPKEQIDGYTMHANYIFWRNTNPIQLSSYSLFDMSKFPLKLRGTAVMSLKENQIND 7044
TR|Q692E6|Q692E6_CVHSA          NYLGKPKEQIDGYTMHANYIFWRNTNPIQLSSYSLFDMSKFPLKLRGTAVMSLKENQIND 7044
TR|A0A0K1YZY7|A0A0K1YZY7_CVHSA  NYLGKPKEQIDGYTMHANYIFWRNTNPIQLSSYSLFDMSKFPLKLRGTAVMFLKENQIND 7044
SP|P0C6W2|R1AB_BCHK3            NYLGKPKEQIDGYTMHANYIFWRNTNPIQLSSYSLFDMSKFPLKLRGTAVMSLKENQIND 7038
SP|P0C6W6|R1AB_BCRP3            NYLGKPKEQIDGYTMHANYIFWRNTNPIQLSSYSLFDMSKFPLKLRGTAVMSLKENQIND 7042
SP|P0C6V9|R1AB_BC279            NYLGKLREQIDGYTMHANYIFWRNTNPIQLSSYSLFDMSKFPLKLRGTAVMSLKENQIND 7050
TR|A0A0U1WHI4|A0A0U1WHI4_CVHSA  NYLGKPKEQIDGYTMHANYVFWRNTNPIQLSSYSLFDMSKFPLKLRGTAVMSLKENQIND 7039
TR|A0A0U1WHG0|A0A0U1WHG0_CVHSA  NYLGKPKEQIDGYTMHANYIFWRNTNPIQLSSYSLFDMSKFPLKLRGTAVMSLKENQIND 7039
TR|A0A166ZL34|A0A166ZL34_9NIDO  NYLGKPKEQIDGYTMHANYIFWRNTNPIQLSSYSLFDMSKFPLKLRGTAVMSLKENQIND 6846
TR|R9QTB2|R9QTB2_CVHSA          ------------------------------------------------------------
TR|R9QTH2|R9QTH2_CVHSA          ------------------------------------------------------------
SP|P0C6U8|R1A_CVHSA             ------------------------------------------------------------
TR|Q6JH47|Q6JH47_CVHSA          ------------------------------------------------------------
TR|Q692E5|Q692E5_CVHSA          ------------------------------------------------------------
SP|P0C6F8|R1A_BCHK3             ------------------------------------------------------------
TR|A0A0K1Z0N1|A0A0K1Z0N1_CVHSA  ------------------------------------------------------------
SP|P0C6F5|R1A_BC279             ------------------------------------------------------------
SP|P0C6T7|R1A_BCRP3             ------------------------------------------------------------
```

```
QHN73794                           MILSLLSKGRLIIRENNRVVISSDVLVNN 7096
SP|P0C6X7|R1AB_CVHSA               MIYSLLEKGRLIIRENNRVVVSSDILVNN 7073
TR|Q6UZF5|Q6UZF5_CVHSA             MIYSLLEKGRLIIRENNRVVVSSDILVNN 7073
TR|Q6UZF1|Q6UZF1_CVHSA             MIYSLLEKGRLIIRENNRVVVSSDILVNN 7073
TR|Q6JH48|Q6JH48_CVHSA             MIYSLLEKGRLIIRENNRVVVSSDILVNN 7073
TR|Q692E6|Q692E6_CVHSA             MIYSLLEKGRLIIRENNRVVVSSDILVNN 7073
TR|A0A0K1YZY7|A0A0K1YZY7_CVHSA     MIYSLLEKGRLIIRENNTVVVSSDVLVNH 7073
SP|P0C6W2|R1AB_BCHK3               MIYSLLEKGRLIIRENNRVVVSSDILVNN 7067
SP|P0C6W6|R1AB_BCRP3               MIYSLLEKGRLIIRENNRVVVSSDILVNN 7071
SP|P0C6V9|R1AB_BC279               MIYSLLENGRLIIRENNRVVVSSDILVNN 7079
TR|A0A0U1WHI4|A0A0U1WHI4_CVHSA     MIYSLLEKGRLIIRENNTVVVSSDVLVNH 7068
TR|A0A0U1WHG0|A0A0U1WHG0_CVHSA     MIYSLLEKGRLIVRENNRVIVSSDVLVNN 7068
TR|A0A166ZL34|A0A166ZL34_9NIDO     MIYSLLEKGRLIVRENNRVIVSSDVLVNN 6875
TR|R9QTB2|R9QTB2_CVHSA             -----------------------------
TR|R9QTH2|R9QTH2_CVHSA             -----------------------------
SP|P0C6U8|R1A_CVHSA                -----------------------------
TR|Q6JH47|Q6JH47_CVHSA             -----------------------------
TR|Q692E5|Q692E5_CVHSA             -----------------------------
SP|P0C6F8|R1A_BCHK3                -----------------------------
TR|A0A0K1Z0N1|A0A0K1Z0N1_CVHSA     -----------------------------
SP|P0C6F5|R1A_BC279                -----------------------------
SP|P0C6T7|R1A_BCRP3                -----------------------------
```
