## Supplementary material for "Structural genomics and interactomics of 2019 Wuhan novel coronavirus, 2019-nCoV, indicate evolutionary conserved functional regions of viral proteins": Protein binding site mapping for wNsp7, wNsp8, wNsp10, wNsp 12, wNsp14, wNsp16

Intra-Viral Heteromeric Interaction Protein Binding Sites:

- wNsp7
- wNsp8
- wNsp12
- wNsp10
- wNsp14
- wNsp16

```
QHN73794                          LLPPKNSIDAFKLNIKLLGVGGKPCIKVATVQSKMSDVKCTSVVLLSVLQQLRVESSSKL 3887
SP|P0C6X7|R1AB_CVHSA              LLPPKSSIDAFKLNIKLLGIGGKPCIKVATVQSKMSDVKCTSVVLLSVLQQLRVESSSKL 3864
TR|Q6UZF5|Q6UZF5_CVHSA            LLPPKSSIDAFKLNIKLLGIGGKPCIKVATVQSKMSDVKCTSVVLLSVLQQLRVESSSKL 3864
TR|Q6UZF1|Q6UZF1_CVHSA            LLPPKSSIDAFKLNIKLLGIGGKPCIKVATVQSKMSDVKCTSVVLLSVLQQLRVESSSKL 3864
TR|Q6JH48|Q6JH48_CVHSA            LLPPKSSIDAFKLNIKLLGIGGKPCIKVATVQSKMSDVKCTSVVLLSVLQQLRVESSSKL 3864
TR|Q692E6|Q692E6_CVHSA            LLPPKSSIDAFKLNIKLLGIGGKPCIKVATVQSKMSDVKCTSVVLLSVLQQLRVESSSKL 3864
TR|A0A0K1YZY7|A0A0K1YZY7_CVHSA    LLPPKSSIDAFRLNIKLLGIGGKPCIKVATVQSKMSDVKCTSVVLLSVLQQLRVESSSKL 3864
SP|P0C6W2|R1AB_BCHK3              LLPPKSSIDAFKLNIKLLGIGGKPCIKVATVQSKMSDVKCTSVVLLSVLQQLRVESSSKL 3858
SP|P0C6W6|R1AB_BCRP3              LLPPKSSIDAFKLNIKLLGIGGKPCIKVATVQSKMSDVKCTSVVLLSVLQQLRVESSSKL 3862
SP|P0C6V9|R1AB_BC279              LLPPKSSIDAFKLNIKLLGIGGKPCIKVATVQSKMSDVKCTSVVLLSVLQQLRVESSSKL 3870
TR|A0A0U1WHI4|A0A0U1WHI4_CVHSA    LLPPKSSIDAFKLNIKLLGIGGKPCIKVATVQSKMSDVKCTSVVLLSVLQQLRVESSSKL 3859
TR|A0A0U1WHG0|A0A0U1WHG0_CVHSA    LLPPKSSIDAFKLNIKLLGIGGKPCIKVATVQSKMSDVKCTSVVLLSVLQQLRVESSSKL 3859
TR|A0A166ZL34|A0A166ZL34_9NIDO    LLPPKSSIDAFKLNIKLLGIGGKPCIKVATVQSKMSDVKCTSVVLLSVLQQLRVESSSKL 3666
TR|R9QTB2|R9QTB2_CVHSA            LLPPKSSIDAFKLNIKLLGIGGKPCIKVATVQSKMSDVKCTSVVLLSVLQQLRVESSSKL 3856
TR|R9QTH2|R9QTH2_CVHSA            LLPPKSSIDAFKLNIKLLGIGGKPCIKVATVQSKMSDVKCTSVVLLSVLQQLRVESSSKL 3865
SP|P0C6U8|R1A_CVHSA               LLPPKSSIDAFKLNIKLLGIGGKPCIKVATVQSKMSDVKCTSVVLLSVLQQLRVESSSKL 3864
TR|Q6JH47|Q6JH47_CVHSA            LLPPKSSIDAFKLNIKLLGIGGKPCIKVATVQSKMSDVKCTSVVLLSVLQQLRVESSSKL 3864
TR|Q692E5|Q692E5_CVHSA            LLPPKSSIDAFKLNIKLLGIGGKPCIKVATVQSKMSDVKCTSVVLLSVLQQLRVESSSKL 3864
SP|P0C6F8|R1A_BCHK3               LLPPKSSIDAFKLNIKLLGIGGKPCIKVATVQSKMSDVKCTSVVLLSVLQQLRVESSSKL 3858
TR|A0A0K1Z0N1|A0A0K1Z0N1_CVHSA    LLPPKSSIDAFRLNIKLLGIGGKPCIKVATVQSKMSDVKCTSVVLLSVLQQLRVESSSKL 3864
SP|P0C6F5|R1A_BC279               LLPPKSSIDAFKLNIKLLGIGGKPCIKVATVQSKMSDVKCTSVVLLSVLQQLRVESSSKL 3870
SP|P0C6T7|R1A_BCRP3               LLPPKSSIDAFKLNIKLLGIGGKPCIKVATVQSKMSDVKCTSVVLLSVLQQLRVESSSKL 3862
                                  *****.*****:*******:****************************************

QHN73794                          WAQCVQLHNDILLAKDTTEAFEKMVSLLSVLLSMQGAVDINKLCEEMLDNRATLQAIASE 3947
SP|P0C6X7|R1AB_CVHSA              WAQCVQLHNDILLAKDTTEAFEKMVSLLSVLLSMQGAVDINRLCEEMLDNRATLQAIASE 3924
TR|Q6UZF5|Q6UZF5_CVHSA            WAQCVQLHNDILLAKDTTEAFEKMVSLLSVLLSMQGAVDINRLCEEMLDNRATLQAIASE 3924
TR|Q6UZF1|Q6UZF1_CVHSA            WAQCVQLHNDILLAKDTTEAFEKMVSLLSVLLSMQGAVDINRLCEEMLDNRATLQAIASE 3924
TR|Q6JH48|Q6JH48_CVHSA            WAQCVQLHNDILLAKDTTEAFEKMVSLLSVLLSMQGAVDINRLCEEMLDNRATLQAIASE 3924
TR|Q692E6|Q692E6_CVHSA            WAQCVQLHNDILLAKDTTEAFEKMVSLLSVLLSMQGAVDINRLCEEMLDNRATLQAIASE 3924
TR|A0A0K1YZY7|A0A0K1YZY7_CVHSA    WAQCVQLHNDILLAKDTTAAFEKMVSLLSVLLSMQGAVDISKLCEEMLDNRATLQAIASE 3924
SP|P0C6W2|R1AB_BCHK3              WAQCVQLHNDILLAKDTTEAFEKMVSLLSVLLSMQGAVDINKLCEEMLDNRATLQAIASE 3918
SP|P0C6W6|R1AB_BCRP3              WAQCVQLHNDILLAKDTTEAFEKMVSLLSVLLSMQGAVDINKLCEEMLDNRATLQAIASE 3922
SP|P0C6V9|R1AB_BC279              WAQCVQLHNDILLAKDTTEAFEKMVSLLSVLLSMQGAVDINKLCEEMLDNRATLQAIASE 3930
TR|A0A0U1WHI4|A0A0U1WHI4_CVHSA    WAQCVQLHNDILLAKDTTEAFEKMVSLLSVLLSMQGAVDINKLCEEMLDNRATLQAIASE 3919
TR|A0A0U1WHG0|A0A0U1WHG0_CVHSA    WARCVQLHNDILLAKDTTEAFEKMVSLLSVLLSMQGAVDINKLCEEMLDNRATLQAIASE 3919
TR|A0A166ZL34|A0A166ZL34_9NIDO    WARCVQLHNDILLAKDTTEAFEKMVSLLSVLLSMQGAVDINKLCEEMLDNRATLQAIASE 3726
TR|R9QTB2|R9QTB2_CVHSA            WAQCVQLHNDILLAKDTTEAFEKMVSLLSVLLSMQGAVDINKLCEEMLDNRATLQAIASE 3916
TR|R9QTH2|R9QTH2_CVHSA            WAQCVQLHNDILLAKDTTEAFEKMVSLLSVLLSMQGAVDINKLCEEMLDNRATLQAIASE 3925
SP|P0C6U8|R1A_CVHSA               WAQCVQLHNDILLAKDTTEAFEKMVSLLSVLLSMQGAVDINRLCEEMLDNRATLQAIASE 3924
TR|Q6JH47|Q6JH47_CVHSA            WAQCVQLHNDILLAKDTTEAFEKMVSLLSVLLSMQGAVDINRLCEEMLDNRATLQAIASE 3924
TR|Q692E5|Q692E5_CVHSA            WAQCVQLHNDILLAKDTTEAFEKMVSLLSVLLSMQGAVDINRLCEEMLDNRATLQAIASE 3924
SP|P0C6F8|R1A_BCHK3               WAQCVQLHNDILLAKDTTEAFEKMVSLLSVLLSMQGAVDINKLCEEMLDNRATLQAIASE 3918
TR|A0A0K1Z0N1|A0A0K1Z0N1_CVHSA    WAQCVQLHNDILLAKDTTAAFEKMVSLLSVLLSMQGAVDISKLCEEMLDNRATLQAIASE 3924
SP|P0C6F5|R1A_BC279               WAQCVQLHNDILLAKDTTEAFEKMVSLLSVLLSMQGAVDINKLCEEMLDNRATLQAIASE 3930
SP|P0C6T7|R1A_BCRP3               WAQCVQLHNDILLAKDTTEAFEKMVSLLSVLLSMQGAVDINKLCEEMLDNRATLQAIASE 3922
                                  **:*************** *********************.:******************

QHN73794                          FSSLPSYAAFATAQEAYEQAVANGDSEVVLKKLKKSLNVAKSEFDRDAAMQRKLEKMADQ 4007
SP|P0C6X7|R1AB_CVHSA              FSSLPSYAAYATAQEAYEQAVANGDSEVVLKKLKKSLNVAKSEFDRDAAMQRKLEKMADQ 3984
TR|Q6UZF5|Q6UZF5_CVHSA            FSSLPSYAAYATAQEAYEQAVANGDSEVVLKKLKKSLNVAKSEFDRDAAMQRKLEKMADQ 3984
TR|Q6UZF1|Q6UZF1_CVHSA            FSSLPSYAAYATAQEAYEQAVANGDSEVVLKKLKKSLNVAKSEFDRDAAMQRKLEKMADQ 3984
TR|Q6JH48|Q6JH48_CVHSA            FSSLPSYAAYATAQEAYEQAVANGDSEVVLKKLKKSLNVAKSEFDRDAAMQRKLEKMADQ 3984
TR|Q692E6|Q692E6_CVHSA            FSSLPSYAAYATAQEAYEQAVANGDSEVVLKKLKKSLNVAKSEFDRDAAMQRKLEKMADQ 3984
TR|A0A0K1YZY7|A0A0K1YZY7_CVHSA    FSSLPSYAAYATAQEAYEQAVANGDSEVVLKKLKKSLNVAKSEFDRDAAMQRKLEKMADQ 3984
SP|P0C6W2|R1AB_BCHK3              FSSLPSYAAYATAQEAYEQAVSNGDSEVVLKKLKKSLNVAKSEFDHDAAMQRKLEKMADQ 3978
SP|P0C6W6|R1AB_BCRP3              FSSLPSYAAYATAQEAYEQAVANGDSEVVLKKLKKSLNVAKSEFDRNAAMQRKLEKMADQ 3982
SP|P0C6V9|R1AB_BC279              FSSLPSYAAYATAQEAYEQAVANGDSEVVLKKLKKSLNVAKSEFDRDAAMQRKLEKMADQ 3990
TR|A0A0U1WHI4|A0A0U1WHI4_CVHSA    FSSLPSYAAYATAQEAYGQAVANGDSEVVLKKLKKSLNVAKSEFDRDAAMQRKLEKMADQ 3979
TR|A0A0U1WHG0|A0A0U1WHG0_CVHSA    FSSLPSYAAYATAQEAYEQAVANGDSEVILKKLKKSLNVAKSEFDRDAAMQRKLEKMADQ 3979
TR|A0A166ZL34|A0A166ZL34_9NIDO    FSSLPSYAAYATAQEAYEQAVANGDSEVILKKLKKSLNVAKSEFDRDAAMQRKLEKMADQ 3786
TR|R9QTB2|R9QTB2_CVHSA            FSSLPSYAAYATAQEAYEQAVANGDSEVVLKKLKKSLNVAKSEFDRDAAMQRKLEKMADQ 3976
TR|R9QTH2|R9QTH2_CVHSA            FSSLPSYAAYATAQEAYEQAVANGDSEVVLKKLKKCLNVAKSEFDRDAAMQRKLEKMADQ 3985
SP|P0C6U8|R1A_CVHSA               FSSLPSYAAYATAQEAYEQAVANGDSEVVLKKLKKSLNVAKSEFDRDAAMQRKLEKMADQ 3984
TR|Q6JH47|Q6JH47_CVHSA            FSSLPSYAAYATAQEAYEQAVANGDSEVVLKKLKKSLNVAKSEFDRDAAMQRKLEKMADQ 3984
TR|Q692E5|Q692E5_CVHSA            FSSLPSYAAYATAQEAYEQAVANGDSEVVLKKLKKSLNVAKSEFDRDAAMQRKLEKMADQ 3984
SP|P0C6F8|R1A_BCHK3               FSSLPSYAAYATAQEAYEQAVSNGDSEVVLKKLKKSLNVAKSEFDHDAAMQRKLEKMADQ 3978
TR|A0A0K1Z0N1|A0A0K1Z0N1_CVHSA    FSSLPSYAAYATAQEAYEQAVANGDSEVVLKKLKKSLNVAKSEFDRDAAMQRKLEKMADQ 3984
SP|P0C6F5|R1A_BC279               FSSLPSYAAYATAQEAYEQAVANGDSEVVLKKLKKSLNVAKSEFDRDAAMQRKLEKMADQ 3990
SP|P0C6T7|R1A_BCRP3               FSSLPSYAAYATAQEAYEQAVANGDSEVVLKKLKKSLNVAKSEFDRNAAMQRKLEKMADQ 3982
                                  *********:******* ***:******:******.*********::*************
```

```
QHN73794                           AMTQMYKQARSEDKRAKVTSAMQTMLFTMLRKLDNDALNNIINNARDGCVPLNIIPLTTA 4067
SP|P0C6X7|R1AB_CVHSA               AMTQMYKQARSEDKRAKVTSAMQTMLFTMLRKLDNDALNNIINNARDGCVPLNIIPLTTA 4044
TR|Q6UZF5|Q6UZF5_CVHSA             AMTQMYKQARSEDKRAKVTSAMQTMLFTMLRKLDNDALNNIINNARDGCVPLNIIPLTTA 4044
TR|Q6UZF1|Q6UZF1_CVHSA             AMTQMYKQARSEDKRAKVTSAMQTMLFTMLRKLDNDALNNIINNARDGCVPLNIIPLTTA 4044
TR|Q6JH48|Q6JH48_CVHSA             AMTQMYKQARSEDKRAKVTSAMQTMLFTMLRKLDNDALNNIINNARDGCVPLNIIPLTTA 4044
TR|Q692E6|Q692E6_CVHSA             AMTQMYKQARSEDKRAKVTSAMQTMLFTMLRKLDNDALNNIINNARDGCVPLNIIPLTTA 4044
TR|A0A0K1YZY7|A0A0K1YZY7_CVHSA     AMTQMYKQARSEDKRAKVTSAMQTMLFTMLRKLDNDALNNIINNARDGCVPLNIIPLTTA 4044
SP|P0C6W2|R1AB_BCHK3               AMTQMYKQARSEDKRAKVTSAMQTMLFTMLRKLDNDALNNIINNARDGCVPLNIIPLTTA 4038
SP|P0C6W6|R1AB_BCRP3               AMTQMYKQARSEDKRAKVTSAMQTMLFTMLRKLDNDALNNIINNARDGCVPLNIIPLTTA 4042
SP|P0C6V9|R1AB_BC279               AMTQMYKQARSEDKRAKVTSAMQTMLFTMLRKLDNDALNNIINNARDGCVPLNIIPLTTA 4050
TR|A0A0U1WHI4|A0A0U1WHI4_CVHSA     AMTQMYKQARSEDKRAKVTSAMQTMLFTMLRKLDNDALNNIINNARDGCVPLNIIPLTTA 4039
TR|A0A0U1WHG0|A0A0U1WHG0_CVHSA     AMTQMYKQARSEDKRAKVTSAMQTMLFTMLRKLDNDALNNIINNARDGCVPLNIIPLTTA 4039
TR|A0A166ZL34|A0A166ZL34_9NIDO     AMTQMYKQARSEDKRAKVTSAMQTMLFTMLRKLDNDALNNIINNARDGCVPLNIIPLTTA 3846
TR|R9QTB2|R9QTB2_CVHSA             AMTQMYKQARSEDKRAKVTSAMQTMLFTMLRKLDNDALNNIINNARDGCVPLNIIPLTTA 4036
TR|R9QTH2|R9QTH2_CVHSA             AMTQMYKQARSEDKRAKVTSAMQTMLFTMLRKLDNDALNNIINNARDGCVPLNIIPLTTA 4045
SP|P0C6U8|R1A_CVHSA                AMTQMYKQARSEDKRAKVTSAMQTMLFTMLRKLDNDALNNIINNARDGCVPLNIIPLTTA 4044
TR|Q6JH47|Q6JH47_CVHSA             AMTQMYKQARSEDKRAKVTSAMQTMLFTMLRKLDNDALNNIINNARDGCVPLNIIPLTTA 4044
TR|Q692E5|Q692E5_CVHSA             AMTQMYKQARSEDKRAKVTSAMQTMLFTMLRKLDNDALNNIINNARDGCVPLNIIPLTTA 4044
SP|P0C6F8|R1A_BCHK3                AMTQMYKQARSEDKRAKVTSAMQTMLFTMLRKLDNDALNNIINNARDGCVPLNIIPLTTA 4038
TR|A0A0K1Z0N1|A0A0K1Z0N1_CVHSA     AMTQMYKQARSEDKRAKVTSAMQTMLFTMLRKLDNDALNNIINNARDGCVPLNIIPLTTA 4044
SP|P0C6F5|R1A_BC279                AMTQMYKQARSEDKRAKVTSAMQTMLFTMLRKLDNDALNNIINNARDGCVPLNIIPLTTA 4050
SP|P0C6T7|R1A_BCRP3                AMTQMYKQARSEDKRAKVTSAMQTMLFTMLRKLDNDALNNIINNARDGCVPLNIIPLTTA 4042
                                   ************************************************************

QHN73794                           AKLMVVIPDYNTYKNTCDGTTFTYASALWEIQQVVDADSKIVQLSEISMDNSPNLAWPLI 4127
SP|P0C6X7|R1AB_CVHSA               AKLMVVVPDYGTYKNTCDGNTFTYASALWEIQQVVDADSKIVQLSEINMDNSPNLAWPLI 4104
TR|Q6UZF5|Q6UZF5_CVHSA             AKLMVVVPDYGTYKNTCDGNTFTYASALWEIQQVVDADSKIVQLSEINMDNSPNLAWPLI 4104
TR|Q6UZF1|Q6UZF1_CVHSA             AKLMVVVPDYGTYKNTCDGNTFTYASALWEIQQVVDADSKIVQLSEINMDNSPNLAWPLI 4104
TR|Q6JH48|Q6JH48_CVHSA             AKLMVVVPDYGTYKNTCDGNTFTYASALWEIQQVVDADSKIVQLSEINMDNSPNLAWPLI 4104
TR|Q692E6|Q692E6_CVHSA             AKLMVVVPDYGTYKNTCDGNTFTYASALWEIQQVVDADSKIVQLSEINMDNSPNLAWPLI 4104
TR|A0A0K1YZY7|A0A0K1YZY7_CVHSA     AKLMVVVPDYGTYKNTCDGNTFTYASALWEIQQVVDADSKIIQLSEINMDNSPNLAWPLI 4104
SP|P0C6W2|R1AB_BCHK3               AKLMVVVPDYGTYKNTCDGNTFTYASALWEIQQVVDADSKIVQLSEINMDNSPNLAWPLI 4098
SP|P0C6W6|R1AB_BCRP3               AKLMVVVPDYGTYKNTCDGNTFTYASALWEIQQVVDADSKIVQLSEINMENSSNLAWPLI 4102
SP|P0C6V9|R1AB_BC279               AKLMVVVPDYGTYKNTCDGNTFTYASALWEIQQVVDADSKIVQLSEINMDNSQNLAWPLI 4110
TR|A0A0U1WHI4|A0A0U1WHI4_CVHSA     AKLMVVVPDYGTYKNTCDGNTFTYASALWEIQQVVDADSKIVPLSEINMDNSPNLAWPLI 4099
TR|A0A0U1WHG0|A0A0U1WHG0_CVHSA     AKLMVVVPDYGTYKNTCDGNTFTYASALWEIQQVVDADSKIVPLSEINMDNSPNLAWPLI 4099
TR|A0A166ZL34|A0A166ZL34_9NIDO     AKLMVVVPDYGTYKNTCDGNTFTYASALWEIQQVVDADSKIVPLSEINMDNSPNLAWPLI 3906
TR|R9QTB2|R9QTB2_CVHSA             AKLMVVVPDYGTYKNTCDGNTFTYASALWEIQQVVDADSKIVQLSEINMDNSPNLAWPLI 4096
TR|R9QTH2|R9QTH2_CVHSA             AKLMVVVPDYGTYKNTCDGNTFTYASALWEIQQVVDADSKIVQLSEINMDNSPNLAWPLI 4105
SP|P0C6U8|R1A_CVHSA                AKLMVVVPDYGTYKNTCDGNTFTYASALWEIQQVVDADSKIVQLSEINMDNSPNLAWPLI 4104
TR|Q6JH47|Q6JH47_CVHSA             AKLMVVVPDYGTYKNTCDGNTFTYASALWEIQQVVDADSKIVQLSEINMDNSPNLAWPLI 4104
TR|Q692E5|Q692E5_CVHSA             AKLMVVVPDYGTYKNTCDGNTFTYASALWEIQQVVDADSKIVQLSEINMDNSPNLAWPLI 4104
SP|P0C6F8|R1A_BCHK3                AKLMVVVPDYGTYKNTCDGNTFTYASALWEIQQVVDADSKIVQLSEINMDNSPNLAWPLI 4098
TR|A0A0K1Z0N1|A0A0K1Z0N1_CVHSA     AKLMVVVPDYGTYKNTCDGNTFTYASALWEIQQVVDADSKIIQLSEINMDNSPNLAWPLI 4104
SP|P0C6F5|R1A_BC279                AKLMVVVPDYGTYKNTCDGNTFTYASALWEIQQVVDADSKIVQLSEINMDNSQNLAWPLI 4110
SP|P0C6T7|R1A_BCRP3                AKLMVVVPDYGTYKNTCDGNTFTYASALWEIQQVVDADSKIVQLSEINMENSSNLAWPLI 4102
                                   ******:***.********.*********************: ****.*:** *******

QHN73794                           VTALRANSAVKLQNNELSPVALRQMSCAAGTTQTACTDDNALAYYNTTKGGRFVLALLSD 4187
SP|P0C6X7|R1AB_CVHSA               VTALRANSAVKLQNNELSPVALRQMSCAAGTTQTACTDDNALAYYNNSKGGRFVLALLSD 4164
TR|Q6UZF5|Q6UZF5_CVHSA             VTALRANSAVKLQNNELSPVALRQMSCAAGTTQTACTDDNALAYYNNSKGGRFVLALLSD 4164
TR|Q6UZF1|Q6UZF1_CVHSA             VTALRANSAVKLQNNELSPVALRQMSCAAGTTQTACTDDNALAYYNNSKGGRFVLALLSD 4164
TR|Q6JH48|Q6JH48_CVHSA             VTALRANSAVKLQNNELSPVALRQMSCAAGTTQTACTDDNALAYYNNSKGGRFVLALLSD 4164
TR|Q692E6|Q692E6_CVHSA             VTALRANSAVKLQNNELSPVALRQMSCAAGTTQTACTDDNALAYYNNSKGGRFVLALLSD 4164
TR|A0A0K1YZY7|A0A0K1YZY7_CVHSA     VTALRANSAVKLQNNELSPVALRQMSCAAGTTQTACTDDNALAYYNNSKGGRFVLALLSD 4164
SP|P0C6W2|R1AB_BCHK3               VTALRANSAVKLQNNELSPVALRQMSCAAGTTQTACTDDNALAYYNNAKGGRFVLALLSD 4158
SP|P0C6W6|R1AB_BCRP3               VTALRANSAVKLQNNELSPVALRQMSCAAGTTQTACTDDNALAYYNNSKGGRFVLALLSD 4162
SP|P0C6V9|R1AB_BC279               VTALRANSAVKLQNNELSPVALRQMSCAAGTTQTACTDDNALAYYNNSKGGRFVLALLSD 4170
TR|A0A0U1WHI4|A0A0U1WHI4_CVHSA     VTALRANSAVKLQNNELSPVALRQMSCAAGTTQTACTDDNALAYYNNSKGGRFVLALLSD 4159
TR|A0A0U1WHG0|A0A0U1WHG0_CVHSA     VTALRANSAVKLQNNELSPVALRQMSCAAGTTQTACTDDNALAYYNNSKGGRFVLALLSD 4159
TR|A0A166ZL34|A0A166ZL34_9NIDO     VTALRANSAVKLQNNELSPVALRQMSCAAGTTQTACTDDNALAYYNNSKGGRFVLALLSD 3966
TR|R9QTB2|R9QTB2_CVHSA             VTALRANSAVKLQNNELSPVALRQMSCAAGTTQTACTDDNALAYYNNSKGGRFVLALLSD 4156
TR|R9QTH2|R9QTH2_CVHSA             VTALRANSAVKLQNNELSPVALRQMSCAAGTTQTACTDDNALAYYNNSKGGRFVLALLSD 4165
SP|P0C6U8|R1A_CVHSA                VTALRANSAVKLQNNELSPVALRQMSCAAGTTQTACTDDNALAYYNNSKGGRFVLALLSD 4164
TR|Q6JH47|Q6JH47_CVHSA             VTALRANSAVKLQNNELSPVALRQMSCAAGTTQTACTDDNALAYYNNSKGGRFVLALLSD 4164
TR|Q692E5|Q692E5_CVHSA             VTALRANSAVKLQNNELSPVALRQMSCAAGTTQTACTDDNALAYYNNSKGGRFVLALLSD 4164
SP|P0C6F8|R1A_BCHK3                VTALRANSAVKLQNNELSPVALRQMSCAAGTTQTACTDDNALAYYNNAKGGRFVLALLSD 4158
TR|A0A0K1Z0N1|A0A0K1Z0N1_CVHSA     VTALRANSAVKLQNNELSPVALRQMSCAAGTTQTACTDDNALAYYNNSKGGRFVLALLSD 4164
SP|P0C6F5|R1A_BC279                VTALRANSAVKLQNNELSPVALRQMSCAAGTTQTACTDDNALAYYNNSKGGRFVLALLSD 4170
SP|P0C6T7|R1A_BCRP3                VTALRANSAVKLQNNELSPVALRQMSCAAGTTQTACTDDNALAYYNNSKGGRFVLALLSD 4162
                                   **********************************************.:************
```

```
QHN73794                        LQDLKWARFPKSDGTGTIYTELEPPCRFVTDTPKGPKVKYLYFIKGLNNLNRGMVLGSLA 4247
SP|P0C6X7|R1AB_CVHSA            HQDLKWARFPKSDGTGTIYTELEPPCRFVTDTPKGPKVKYLYFIKGLNNLNRGMVLGSLA 4224
TR|Q6UZF5|Q6UZF5_CVHSA          HQDLKWARFPKSDGTGTIYTELEPPCRFVTDTPKGPKVKYLYFIKGLNNLNRGMVLGSLA 4224
TR|Q6UZF1|Q6UZF1_CVHSA          HQDLKWARFPKSDGTGTIYTELEPPCRFVTDTPKGPKVKYLYFIKGLNNLNRGMVLGSLA 4224
TR|Q6JH48|Q6JH48_CVHSA          HQDLKWARFPKSDGTGTIYTELEPPCRFVTDTPKGPKVKYLYFIKGLNNLNRGMVLGSLA 4224
TR|Q692E6|Q692E6_CVHSA          HQDLKWARFPKSDGTGTIYTELEPPCRFVTDTPKGPKVKYLYFIKGLNNLNRGMVLGSLA 4224
TR|A0A0K1YZY7|A0A0K1YZY7_CVHSA  HQDLKWARFPKSDGTGTIYTELEPPCRFVTDTPKGPKVKYLYFIKGLNNLNRGMVLGSLA 4224
SP|P0C6W2|R1AB_BCHK3            HQDLKWARFPKSDGTGTIYTELEPPCRFVTDTPKGPKVKYLYFIKGLNNLNRGMVLGSLA 4218
SP|P0C6W6|R1AB_BCRP3            HQDLKWARFPKSDGTGTIYTELEPPCRFVTDTPKGPKVKYLHFIKGLNNLNRGMVLGSLA 4222
SP|P0C6V9|R1AB_BC279            HQDLKWARFPKSDGTGTIYTELEPPCRFVTDTPRGPKVKYLYFIKGLNNLNRGMVLGSLA 4230
TR|A0A0U1WHI4|A0A0U1WHI4_CVHSA  HQDLKWARFPKSDGTGTIYTELEPPCRFVTDTPKGPKVKYLYFIKGLNNLNRGMVLGSLA 4219
TR|A0A0U1WHG0|A0A0U1WHG0_CVHSA  HQDLKWARFPKSDGTGTIYTELEPPCRFVTDTPKGPKVKYLYFIKGLNNLNRGMVLGSLA 4219
TR|A0A166ZL34|A0A166ZL34_9NIDO  HQDLKWARFPKSDGTGTIYTELEPPCRFVTDTPKGPKVKYLYFIKGLNNLNRGMVLGSLA 4026
TR|R9QTB2|R9QTB2_CVHSA          HQDLKWARFPKSDGTGTIYTELEPPCRFVTDTPKGPKVKYLYFIKGLNNLNRGMVLGSLA 4216
TR|R9QTH2|R9QTH2_CVHSA          HQDLKWARFPKSDGSGTIYTELEPPCRFVTDTPKGPKVKYLYFIKGLNNLNRGMVLGSLA 4225
SP|P0C6U8|R1A_CVHSA             HQDLKWARFPKSDGTGTIYTELEPPCRFVTDTPKGPKVKYLYFIKGLNNLNRGMVLGSLA 4224
TR|Q6JH47|Q6JH47_CVHSA          HQDLKWARFPKSDGTGTIYTELEPPCRFVTDTPKGPKVKYLYFIKGLNNLNRGMVLGSLA 4224
TR|Q692E5|Q692E5_CVHSA          HQDLKWARFPKSDGTGTIYTELEPPCRFVTDTPKGPKVKYLYFIKGLNNLNRGMVLGSLA 4224
SP|P0C6F8|R1A_BCHK3             HQDLKWARFPKSDGTGTIYTELEPPCRFVTDTPKGPKVKYLYFIKGLNNLNRGMVLGSLA 4218
TR|A0A0K1Z0N1|A0A0K1Z0N1_CVHSA  HQDLKWARFPKSDGTGTIYTELEPPCRFVTDTPKGPKVKYLYFIKGLNNLNRGMVLGSLA 4224
SP|P0C6F5|R1A_BC279             HQDLKWARFPKSDGTGTIYTELEPPCRFVTDTPRGPKVKYLYFIKGLNNLNRGMVLGSLA 4230
SP|P0C6T7|R1A_BCRP3             HQDLKWARFPKSDGTGTIYTELEPPCRFVTDTPKGPKVKYLHFIKGLNNLNRGMVLGSLA 4222
                                 *************:******************:*******:******************

QHN73794                        ATVRLQAGNATEVPANSTVLSFCAFAVDAAKAYKDYLASGGQPITNCVKMLCTHTGTGQA 4307
SP|P0C6X7|R1AB_CVHSA            ATVRLQAGNATEVPANSTVLSFCAFAVDPAKAYKDYLASGGQPITNCVKMLCTHTGTGQA 4284
TR|Q6UZF5|Q6UZF5_CVHSA          ATVRLQAGNATEVPANSTVLSFCAFAVDPAKAYKDYLASGGQPITNCVKMLCTHTGTGQA 4284
TR|Q6UZF1|Q6UZF1_CVHSA          ATVRLQAGNATEVPANSTVLSFCAFAVDPAKAYKDYLASGGQPITNCVKMLCTHTGTGQA 4284
TR|Q6JH48|Q6JH48_CVHSA          ATVRLQAGNATEVPANSTVLSFCAFAVDPAKAYKDYLASGGQPITNCVKMLCTHTGTGQA 4284
TR|Q692E6|Q692E6_CVHSA          ATVRLQAGNATEVPANSTVLSFCAFAVDPAKAYKDYLASGGQPITNCVKMLCTHTGTGQA 4284
TR|A0A0K1YZY7|A0A0K1YZY7_CVHSA  ATVRLQAGNATEVPANSTVLSFCAFAVDPAKAYKDYLASGGQPITNCVKMLCTHTGTGQA 4284
SP|P0C6W2|R1AB_BCHK3            ATVRLQAGNATEVPANSTVLSFCAFAVDPAKAYKDYLASGGQPITNCVKMLCTHTGTGQA 4278
SP|P0C6W6|R1AB_BCRP3            ATVRLQAGNATEVPANSTVLSFCAFAVDPAKAYKDYLASGGQPITNCVKMLCTHTGTGQA 4282
SP|P0C6V9|R1AB_BC279            ATVRLQAGNATEVPANSAVLSFCAFAVDPAKAYKDYLASGGQPITNCVKMLCTHTGTGQA 4290
TR|A0A0U1WHI4|A0A0U1WHI4_CVHSA  ATVRLQAGNATEVPANSTVLSFCAFAVDPAKAYKDYLSSGGQPITNCVKMLCTHTGTGQA 4279
TR|A0A0U1WHG0|A0A0U1WHG0_CVHSA  ATVRLQAGNATEVPANSTVLSFCAFAVDPAKAYKDYLSSGGQPITNCVKMLCTHTGTGQA 4279
TR|A0A166ZL34|A0A166ZL34_9NIDO  ATVRLQAGNATEVPANSTVLSFCAFAVDPAKAYKDYLSSGGQPITNCVKMLCTHTGTGQA 4086
TR|R9QTB2|R9QTB2_CVHSA          ATVRLQAGNATEVPANSTVLSFCAFAVDPAKAYKDYLASGGQPITNCVKMLCTHTGTGQA 4276
TR|R9QTH2|R9QTH2_CVHSA          ATVRLQAGNATEVPANSTVLSFCAFAVDPAKAYKDYLASGGQPITNCVKMLCTHTGTGQA 4285
SP|P0C6U8|R1A_CVHSA             ATVRLQAGNATEVPANSTVLSFCAFAVDPAKAYKDYLASGGQPITNCVKMLCTHTGTGQA 4284
TR|Q6JH47|Q6JH47_CVHSA          ATVRLQAGNATEVPANSTVLSFCAFAVDPAKAYKDYLASGGQPITNCVKMLCTHTGTGQA 4284
TR|Q692E5|Q692E5_CVHSA          ATVRLQAGNATEVPANSTVLSFCAFAVDPAKAYKDYLASGGQPITNCVKMLCTHTGTGQA 4284
SP|P0C6F8|R1A_BCHK3             ATVRLQAGNATEVPANSTVLSFCAFAVDPAKAYKDYLASGGQPITNCVKMLCTHTGTGQA 4278
TR|A0A0K1Z0N1|A0A0K1Z0N1_CVHSA  ATVRLQAGNATEVPANSTVLSFCAFAVDPAKAYKDYLASGGQPITNCVKMLCTHTGTGQA 4284
SP|P0C6F5|R1A_BC279             ATVRLQAGNATEVPANSAVLSFCAFAVDPAKAYKDYLASGGQPITNCVKMLCTHTGTGQA 4290
SP|P0C6T7|R1A_BCRP3             ATVRLQAGNATEVPANSTVLSFCAFAVDPAKAYKDYLASGGQPITNCVKMLCTHTGTGQA 4282
                                *****************:********** ********:**********************

QHN73794                        ITVTPEANMDQESFGGASCCLYCRCHIDHPNPKGFCDLKGKYVQIPTTCANDPVGFTLKN 4367
SP|P0C6X7|R1AB_CVHSA            ITVTPEANMDQESFGGASCCLYCRCHIDHPNPKGFCDLKGKYVQIPTTCANDPVGFTLRN 4344
TR|Q6UZF5|Q6UZF5_CVHSA          ITVTPEANMDQESFGGASCCLYCRCHIDHPNPKGFCDLKGKYVQIPTTCANDPVGFTLRN 4344
TR|Q6UZF1|Q6UZF1_CVHSA          ITVTPEANMDQESFGGASCCLYCRCHIDHPNPKGFCDLKGKYVQIPTTCANDPVGFTLRN 4344
TR|Q6JH48|Q6JH48_CVHSA          ITVTPEANMDQESFGGASCCLYCRCHIDHPNPKGFCDLKGKYVQIPTTCANDPVGFTLRN 4344
TR|Q692E6|Q692E6_CVHSA          ITVTPEANMDQESFGGASCCLYCRCHIDHPNPKGFCDLKGKYVQIPTTCANDPVGFTLRN 4344
TR|A0A0K1YZY7|A0A0K1YZY7_CVHSA  ITVTPEANMDQESFGGASCCLYCRCHIDHPNPKGFCDLKGKYIQIPTTCANDPVGFTLRN 4344
SP|P0C6W2|R1AB_BCHK3            ITVTPEANMDQESFGGASCCLYCRCHIDHPNPKGFCDLKGKYVQIPTTCANDPVGFTLRN 4338
SP|P0C6W6|R1AB_BCRP3            ITVTPEANMDQESFGGASCCLYCRCHIDHPNPKGFCDLKGKYVQIPTTCANDPVGFTLRN 4342
SP|P0C6V9|R1AB_BC279            ITVTPEANMDQESFGGASCCLYCRCHIDHPNPKGFCDLKGKYVQIPATCANDPVGFTLKN 4350
TR|A0A0U1WHI4|A0A0U1WHI4_CVHSA  ITVTPEANMDQESFGGASCCLYCRCHIDHPNPKGFCDLKGKYVQIPTTCANDPVGFTLRN 4339
TR|A0A0U1WHG0|A0A0U1WHG0_CVHSA  ITVTPEANMDQESFGGASCCLYCRCHIDHPNPKGFCDLKGKYVQIPTTCANDPVGFTLRN 4339
TR|A0A166ZL34|A0A166ZL34_9NIDO  ITVTPEANMDQESFGGASCCLYCRCHIDHPNPKGFCDLKGKYVQIPTTCANDPVGFTLRN 4146
TR|R9QTB2|R9QTB2_CVHSA          ITVTPEANMDQESFGGASCCLYCRCHIDHPNPKGFCDLKGKYVQIPTTCANDPVGFTLRN 4336
TR|R9QTH2|R9QTH2_CVHSA          ITVTPEANMDQESFGGASCCLYCRCHIDHPNPKGFCDLKGKYVQIPTTCANDPVGFTLRN 4345
SP|P0C6U8|R1A_CVHSA             ITVTPEANMDQESFGGASCCLYCRCHIDHPNPKGFCDLKGKYVQIPTTCANDPVGFTLRN 4344
TR|Q6JH47|Q6JH47_CVHSA          ITVTPEANMDQESFGGASCCLYCRCHIDHPNPKGFCDLKGKYVQIPTTCANDPVGFTLRN 4344
TR|Q692E5|Q692E5_CVHSA          ITVTPEANMDQESFGGASCCLYCRCHIDHPNPKGFCDLKGKYVQIPTTCANDPVGFTLRN 4344
SP|P0C6F8|R1A_BCHK3             ITVTPEANMDQESFGGASCCLYCRCHIDHPNPKGFCDLKGKYVQIPTTCANDPVGFTLRN 4338
TR|A0A0K1Z0N1|A0A0K1Z0N1_CVHSA  ITVTPEANMDQESFGGASCCLYCRCHIDHPNPKGFCDLKGKYIQIPTTCANDPVGFTLRN 4344
SP|P0C6F5|R1A_BC279             ITVTPEANMDQESFGGASCCLYCRCHIDHPNPKGFCDLKGKYVQIPATCANDPVGFTLKN 4350
SP|P0C6T7|R1A_BCRP3             ITVTPEANMDQESFGGASCCLYCRCHIDHPNPKGFCDLKGKYVQIPTTCANDPVGFTLRN 4342
                                ******************************************:***:***********:*
```

```
QHN73794                       TVCTVCGMWKGYGCSCDQLREPMLQSADAQSFLNRVCGVSAARLTPCGTGTSTDVVYRAF 4427
SP|P0C6X7|R1AB_CVHSA           TVCTVCGMWKGYGCSCDQLREPLMQSADASTFLNRVCGVSAARLTPCGTGTSTDVVYRAF 4404
TR|Q6UZF5|Q6UZF5_CVHSA         TVCTVCGMWKGYGCSCDQLREPLMQSADASTFLNRVCGVSAARLTPCGTGTSTDVVYRAF 4404
TR|Q6UZF1|Q6UZF1_CVHSA         TVCTVCGMWKGYGCSCDQLREPLMQSADASTFLNRVCGVSAARLTPCGTGTSTDVVYRAF 4404
TR|Q6JH48|Q6JH48_CVHSA         TVCTVCGMWKGYGCSCDQLREPLMQSADASTFLNRVCGVSAARLTPCGTGTSTDVVYRAF 4404
TR|Q692E6|Q692E6_CVHSA         TVCTVCGMWKGYGCSCDQLREPLMQSADASTFFKRVCGVSAARLTPCGTGTSTDVVYRAF 4404
TR|A0A0K1YZY7|A0A0K1YZY7_CVHSA TVCTVCGMWKGYGCSCDQLREPMMQSADASTFLNRVCGVSAARLTPCGTGISTDVVYRAF 4404
SP|P0C6W2|R1AB_BCHK3           TVCTVCGMWKGYGCSCDQLREPMMQSADASTFLNRVCGVSAARLTPCGTGTSTDVVYRAF 4398
SP|P0C6W6|R1AB_BCRP3           TVCTVCGMWKGYGCSCDQLREPMMQSADASTFLNRVCGVSAARLTPCGTGISTDVVYRAF 4402
SP|P0C6V9|R1AB_BC279           TVCTVCGTWKGYGCSCDQLREPMMQSADASTFLNRVCGVSAARLTPCGTGTSTDVVYRAF 4410
TR|A0A0U1WHI4|A0A0U1WHI4_CVHSA TVCTVCGMWKGYGCSCDQLREPMMQSADASAFLNRVCGVSAARLTPCGTGTSTDVVYRAF 4399
TR|A0A0U1WHG0|A0A0U1WHG0_CVHSA TVCTVCGMWKGYGCSCDQLREPMMQSADASPFLNRVCGVSAARLTPCGTGTSTDVVYRAF 4399
TR|A0A166ZL34|A0A166ZL34_9NIDO TVCTVCGMWKGYGCSCDQLREPMMQSADASPFLNRVCGVSAARLTPCGTGTSTDVVYRAF 4206
TR|R9QTB2|R9QTB2_CVHSA         TVCTVCGMWKGYGCSCDQLREPMMQSADASTFLNGFAV---------------------- 4374
TR|R9QTH2|R9QTH2_CVHSA         TVCTVCGMWKGYGCSCDQLREPMMQSADASTFLNGFAV---------------------- 4383
SP|P0C6U8|R1A_CVHSA            TVCTVCGMWKGYGCSCDQLREPLMQSADASTFLNGFAV---------------------- 4382
TR|Q6JH47|Q6JH47_CVHSA         TVCTVCGMWKGYGCSCDQLREPLMQSADASTFLNGFAV---------------------- 4382
TR|Q692E5|Q692E5_CVHSA         TVCTVCGMWKGYGCSCDQLREPLMQSADASTFLNGFAV---------------------- 4382
SP|P0C6F8|R1A_BCHK3            TVCTVCGMWKGYGCSCDQLREPMMQSADASTFLNGFAV---------------------- 4376
TR|A0A0K1Z0N1|A0A0K1Z0N1_CVHSA TVCTVCGMWKGYGCSCDQLREPMMQSADASTFLNGFAV---------------------- 4382
SP|P0C6F5|R1A_BC279            TVCTVCGTWKGYGCSCDQLREPMMQSADASTFLNGFAV---------------------- 4388
SP|P0C6T7|R1A_BCRP3            TVCTVCGMWKGYGCSCDQLREPMMQSADASTFLNGFAV---------------------- 4380
                               ******* **************::*****. *:: ..

QHN73794                       DIYNDKVAGFAKFLKTNCCRFQEKDEDDNLIDSYFVVKRHTFSNYQHEETIYNLLKDCPA 4487
SP|P0C6X7|R1AB_CVHSA           DIYNEKVAGFAKFLKTNCCRFQEKDEEGNLLDSYFVVKRHTMSNYQHEETIYNLVKDCPA 4464
TR|Q6UZF5|Q6UZF5_CVHSA         DIYNEKVAGFAKFLKTNCCRFQEKDEEGNLLDSYFVVKRHTMSNYQHEETIYNLVKDCPA 4464
TR|Q6UZF1|Q6UZF1_CVHSA         DIYNEKVAGFAKFLKTNCCRFQEKDEEGNLLDSYFVVKRHTMSNYQHEETIYNLVKDCPA 4464
TR|Q6JH48|Q6JH48_CVHSA         DIYNEKVAGFAKFLKTNCCRFQEKDEEGNLLDSYFVVKRHTMSNYQHEETIYNLVKDCPA 4464
TR|Q692E6|Q692E6_CVHSA         DIYNEKVAGFAKFLKTNCCRFQEKDEEGNLLDSYFVVKRHTMSNYQHEETIYNLVKDCPA 4464
TR|A0A0K1YZY7|A0A0K1YZY7_CVHSA DIYNEKVAGFAKFLKTNCCRFQEKDEEGNLLDSYFVVKRHTMSNYQHEETIYNLVKDCPA 4464
SP|P0C6W2|R1AB_BCHK3           DIYNEKVAGFAKFLKTNCCRFQEKDEEGNLLDSYFVVKRHTMSNYQHEETIYNLIKECPA 4458
SP|P0C6W6|R1AB_BCRP3           DIYNEKVAGFAKFLKTNCCRFQEKDEEGNLLDSYFVVKRHTMSNYQHEETIYNLVKDCPA 4462
SP|P0C6V9|R1AB_BC279           DIYNERVAGFAKFLKTNCCRFQEKDEEGNLLDSYFVVKRHTMSNYQHEETIYNLVKECPA 4470
TR|A0A0U1WHI4|A0A0U1WHI4_CVHSA DIYNEKVAGFAKFLKTNCCRFQEKDEEGNLIDSYFVVKRHTMSNYQHEEAIYNLLKECPA 4459
TR|A0A0U1WHG0|A0A0U1WHG0_CVHSA DIYNEKVAGFAKFLKTNCCRFQEMDEDGNLIDSYFVVKRHTMSNYQHEEAIYNLLKECPA 4459
TR|A0A166ZL34|A0A166ZL34_9NIDO DIYNEKVAGFAKFLKTNCCRFQEMDEDGNLIDSYFVVKRHTMSNYQHEEAIYNLLKECPA 4266
TR|R9QTB2|R9QTB2_CVHSA         ------------------------------------------------------------
TR|R9QTH2|R9QTH2_CVHSA         ------------------------------------------------------------
SP|P0C6U8|R1A_CVHSA            ------------------------------------------------------------
TR|Q6JH47|Q6JH47_CVHSA         ------------------------------------------------------------
TR|Q692E5|Q692E5_CVHSA         ------------------------------------------------------------
SP|P0C6F8|R1A_BCHK3            ------------------------------------------------------------
TR|A0A0K1Z0N1|A0A0K1Z0N1_CVHSA ------------------------------------------------------------
SP|P0C6F5|R1A_BC279            ------------------------------------------------------------
SP|P0C6T7|R1A_BCRP3            ------------------------------------------------------------


QHN73794                       VAKHDFFKFRIDGDMVPHISRQRLTKYTMADLVYALRHFDEGNCDTLKEILVTYNCCDDD 4547
SP|P0C6X7|R1AB_CVHSA           VAVHDFFKFRVDGDMVPHISRQRLTKYTMADLVYALRHFDEGNCDTLKEILVTYNCCDDD 4524
TR|Q6UZF5|Q6UZF5_CVHSA         VAVHDFFKFRVDGDMVPHISRQRLTKYTMADLVYALRHFDEGNCDTLKEILVTYNCCDDD 4524
TR|Q6UZF1|Q6UZF1_CVHSA         VAVHDFFKFRVDGDMVPHISRQRLTKYTMADLVYALRHFDEGNCDTLKEILVTYNCCDDD 4524
TR|Q6JH48|Q6JH48_CVHSA         VAVHDFFKFRVDGDMVPHISRQRLTKYTMADLVYALRHFDEGNCDTLKEILVTYNCCDDD 4524
TR|Q692E6|Q692E6_CVHSA         VAVHDFFKFRVDGDMVPHISRQRLTKYTMADLVYALRHFDEGNCDTLKEILVTYNCCDDD 4524
TR|A0A0K1YZY7|A0A0K1YZY7_CVHSA VAVHDFFKFRVDGDMVPHISRQRLTKYTMADLVYALRHFDEGNCDTLKEILVTYNCCDDD 4524
SP|P0C6W2|R1AB_BCHK3           VAVHDFFKFRVDGDMVPHISRQRLTKYTMADLVYALRHFDEGNCDTLKEILVTYNCCDDN 4518
SP|P0C6W6|R1AB_BCRP3           VAVHDFFKFRVDGDMVPHISRQRLTKYTMADLVYALRHFDEGNCDTLKEILVTYNCCDDD 4522
SP|P0C6V9|R1AB_BC279           VAVHDFFKFRVDGDMVPHISRQRLTKYTMADLVYALRHFDEGNCDTLKEILVTYNCCDDD 4530
TR|A0A0U1WHI4|A0A0U1WHI4_CVHSA VAVHDFFKFRVDGDMVPHISRQRLTKYTMADLVYALRHFDEGNCDTLKEILVTYNCCDDD 4519
TR|A0A0U1WHG0|A0A0U1WHG0_CVHSA VAVHDFFKFRVDGDMVPHISRQRLTKYTMADLVYALRHFDEGNCDTLKEILVTYNCCDDD 4519
TR|A0A166ZL34|A0A166ZL34_9NIDO VAVHDFFKFRVDGDMVPHISRQRLTKYTMADLVYALRHFDEGNCDTLKEILVTYNCCDDD 4326
TR|R9QTB2|R9QTB2_CVHSA         ------------------------------------------------------------
TR|R9QTH2|R9QTH2_CVHSA         ------------------------------------------------------------
SP|P0C6U8|R1A_CVHSA            ------------------------------------------------------------
TR|Q6JH47|Q6JH47_CVHSA         ------------------------------------------------------------
TR|Q692E5|Q692E5_CVHSA         ------------------------------------------------------------
SP|P0C6F8|R1A_BCHK3            ------------------------------------------------------------
TR|A0A0K1Z0N1|A0A0K1Z0N1_CVHSA ------------------------------------------------------------
SP|P0C6F5|R1A_BC279            ------------------------------------------------------------
SP|P0C6T7|R1A_BCRP3            ------------------------------------------------------------
```

```
QHN73794                              YFNKKDWYDFVENPDILRVYANLGERVRQALLKTVQFCDAMRNAGIVGVLTLDNQDLNGN 4607
SP|P0C6X7|R1AB_CVHSA                  YFNKKDWYDFVENPDILRVYANLGERVRQSLLKTVQFCDAMRDAGIVGVLTLDNQDLNGN 4584
TR|Q6UZF5|Q6UZF5_CVHSA                YFNKKDWYDFVENPDILRVYANLGERVRQSLLKTVQFCDAMRDAGIVGVLTLDNQDLNGN 4584
TR|Q6UZF1|Q6UZF1_CVHSA                YFNKKDWYDFVENPDILRVYANLGERVRQSLLKTVQFCDAMRDAGIVGVLTLDNQDLNGN 4584
TR|Q6JH48|Q6JH48_CVHSA                YFNKKDWYDFVENPDILRVYANLGERVRQSLLKTVQFCDAMRDAGIVGVLTLDNQDLNGN 4584
TR|Q692E6|Q692E6_CVHSA                YFNKKDWYDFVENPDILRVYANLGERVRQSLLKTVQFCDAMRDAGIVGVLTLDNQDLNGN 4584
TR|A0A0K1YZY7|A0A0K1YZY7_CVHSA YFNKKDWYDFVENPDILRVYANLGERVRQALLKTVQFCDAMRDAGIVGVLTLDNQDLNGN 4584
SP|P0C6W2|R1AB_BCHK3                  YFNKKDWYDFVENPDVLRVYANLGERVRRALLKTVQFCDAMRDAGIVGVLTLDNQDLNGN 4578
SP|P0C6W6|R1AB_BCRP3                  YFNKKDWYDFVENPDILRVYANLGERVRQALLKTVQFCDAMRDAGIVGVLTLDNQDLNGN 4582
SP|P0C6V9|R1AB_BC279                  YFNKKDWYDFVENPDILRVYANLGERVRQALLKTVQFCDAMRDAGIVGVLTLDNQDLNGN 4590
TR|A0A0U1WHI4|A0A0U1WHI4_CVHSA YFNKKDWYDFVENPDILRVYANLGERVRQALLKTVQFCDAMRDAGIVGVLTLDNQDLNGN 4579
TR|A0A0U1WHG0|A0A0U1WHG0_CVHSA YFNKKDWYDFVENPDILRVYANLGERVRQALLKTVQFCDAMRDAGIVGVLTLDNQDLNGN 4579
TR|A0A166ZL34|A0A166ZL34_9NIDO YFNKKDWYDFVENPDILRVYANLGERVRQALLKTVQFCDAMRDAGIVGVLTLDNQDLNGN 4386
TR|R9QTB2|R9QTB2_CVHSA                ------------------------------------------------------------
TR|R9QTH2|R9QTH2_CVHSA                ------------------------------------------------------------
SP|P0C6U8|R1A_CVHSA                   ------------------------------------------------------------
TR|Q6JH47|Q6JH47_CVHSA                ------------------------------------------------------------
TR|Q692E5|Q692E5_CVHSA                ------------------------------------------------------------
SP|P0C6F8|R1A_BCHK3                   ------------------------------------------------------------
TR|A0A0K1Z0N1|A0A0K1Z0N1_CVHSA ------------------------------------------------------------
SP|P0C6F5|R1A_BC279                   ------------------------------------------------------------
SP|P0C6T7|R1A_BCRP3                   ------------------------------------------------------------

QHN73794                              WYDFGDFIQTTPGSGVPVVDSYYSLLMPILTLTRALTAESHVDTDLTKPYIKWDLLKYDF 4667
SP|P0C6X7|R1AB_CVHSA                  WYDFGDFVQVAPGCGVPIVDSYYSLLMPILTLTRALAAESHMDADLAKPLIKWDLLKYDF 4644
TR|Q6UZF5|Q6UZF5_CVHSA                WYDFGDFVQVAPGCGVPIVDSYYSLLMPILTLTRALAAESHMDADLAKPLIKWDLLKYDF 4644
TR|Q6UZF1|Q6UZF1_CVHSA                WYDFGDFVQVAPGCGVPIVDSYYSLLMPILTLTRALAAESHMDADLAKPLIKWDLLKYDF 4644
TR|Q6JH48|Q6JH48_CVHSA                WYDFGDFVQVAPGCGVPIVDSYYSLLMPILTLTRALAAESHMDADLAKPLIKWDLLKYDF 4644
TR|Q692E6|Q692E6_CVHSA                WYDFGDFVQVAPGCGVPIVDSYYSLLMPILTLTRALAAESHMDADLAKPLIKWDLLKYDF 4644
TR|A0A0K1YZY7|A0A0K1YZY7_CVHSA WYDFGDFVQVAPGCGVPIVDSYYSLLMPILTMTRALAAESHMDADLAKPLIKWDLLKYDF 4644
SP|P0C6W2|R1AB_BCHK3                  WYDFGDFVQVAPGCGVPIVDSYYSLLMPILTLTKALAAESHMDADLAKPLVKWDLLKYDF 4638
SP|P0C6W6|R1AB_BCRP3                  WYDFGDFVQVAPGCGVPIVDSYYSLLMPILTLTRALAAESHMDADLAKPLIKWDLLKYDF 4642
SP|P0C6V9|R1AB_BC279                  WYDFGDFVQVAPGCGVPIVDSYYSLLMPILTLTKALAAESHMDADLAKPLIKWDLLKYDF 4650
TR|A0A0U1WHI4|A0A0U1WHI4_CVHSA WYDFGDFVQVAPGCGVPIVDSYYSLLMPILTLTRALAAESHMDADLTKPLIKWDLLKYDF 4639
TR|A0A0U1WHG0|A0A0U1WHG0_CVHSA WYDFGDFVQVTPGCGVPIVDSYYSLLMPILTLTRALAAESHMDTDLTKPLIKWDLLKYDF 4639
TR|A0A166ZL34|A0A166ZL34_9NIDO WYDFGDFVQVTPGCGVPIVDSYYSLLMPILTLTRALAAESHMDTDLTKPLIKWDLLKYDF 4446
TR|R9QTB2|R9QTB2_CVHSA                ------------------------------------------------------------
TR|R9QTH2|R9QTH2_CVHSA                ------------------------------------------------------------
SP|P0C6U8|R1A_CVHSA                   ------------------------------------------------------------
TR|Q6JH47|Q6JH47_CVHSA                ------------------------------------------------------------
TR|Q692E5|Q692E5_CVHSA                ------------------------------------------------------------
SP|P0C6F8|R1A_BCHK3                   ------------------------------------------------------------
TR|A0A0K1Z0N1|A0A0K1Z0N1_CVHSA ------------------------------------------------------------
SP|P0C6F5|R1A_BC279                   ------------------------------------------------------------
SP|P0C6T7|R1A_BCRP3                   ------------------------------------------------------------

QHN73794                              TEERLKLFDRYFKYWDQTYHPNCVNCLDDRCILHCANFNVLFSTVFPPTSFGPLVRKIFV 4727
SP|P0C6X7|R1AB_CVHSA                  TEERLCLFDRYFKYWDQTYHPNCINCLDDRCILHCANFNVLFSTVFPPTSFGPLVRKIFV 4704
TR|Q6UZF5|Q6UZF5_CVHSA                TEERLCLFDRYFKYWDQTYHPNCINCLDDRCILHCANFNVLFSTVFPPTSFGPLVRKIFV 4704
TR|Q6UZF1|Q6UZF1_CVHSA                TEERLCLFDRYFKYWDQTYHPNCINCLDDRCILHCANFNVLFSTVFPPTSFGPLVRKIFV 4704
TR|Q6JH48|Q6JH48_CVHSA                TEERLCLFDRYFKYWDQTYHPNCINCLDDRCILHCANFNVLFSTVFPPTSFGPLVRKIFV 4704
TR|Q692E6|Q692E6_CVHSA                TEERLCLFDRYFKYWDQTYHPNCINCLDDRCILHCANFNVLFSTVFPPTSFGPLVRKIFV 4704
TR|A0A0K1YZY7|A0A0K1YZY7_CVHSA TEERLCLFDRYFKYWDQTYHPNCINCLDDRCILHCANFNVLFSTVFPPTSFGPLVRKIFV 4704
SP|P0C6W2|R1AB_BCHK3                  TEERLCLFDRYFKYWDQTYHPNCINCLDDRCILHCANFNVLFSTVFPPTSFGPLVRKIFV 4698
SP|P0C6W6|R1AB_BCRP3                  TAERLCLFDRYFKYWDQTYHPNCINCLDDRCILHCANFNVLFSTVFPPTSFGPLVRKIFV 4702
SP|P0C6V9|R1AB_BC279                  TEERLCLFDRYFKYWDQTYHPNCINCLDDRCILHCANFNVLFSTVFPPTSFGPLVRKIFV 4710
TR|A0A0U1WHI4|A0A0U1WHI4_CVHSA TEERLCLFDRYFKYWDQTYHPNCVNCLDDRCILHCANFNVLFSTVFPPTSFGPLVRKIFV 4699
TR|A0A0U1WHG0|A0A0U1WHG0_CVHSA TEERLCLFDRYFKYWDQTYHPNCINCLDDRCILHCANFNVLFSTVFPPTSFGPLVRKMFV 4699
TR|A0A166ZL34|A0A166ZL34_9NIDO TEERLCLFDRYFKYWDQTYHPNCINCLDDRCILHCANFNVLFSTVFPPTSFGPLVRKMFV 4506
TR|R9QTB2|R9QTB2_CVHSA                ------------------------------------------------------------
TR|R9QTH2|R9QTH2_CVHSA                ------------------------------------------------------------
SP|P0C6U8|R1A_CVHSA                   ------------------------------------------------------------
TR|Q6JH47|Q6JH47_CVHSA                ------------------------------------------------------------
TR|Q692E5|Q692E5_CVHSA                ------------------------------------------------------------
SP|P0C6F8|R1A_BCHK3                   ------------------------------------------------------------
TR|A0A0K1Z0N1|A0A0K1Z0N1_CVHSA ------------------------------------------------------------
SP|P0C6F5|R1A_BC279                   ------------------------------------------------------------
SP|P0C6T7|R1A_BCRP3                   ------------------------------------------------------------
```

```
QHN73794                          DGVPFVVSTGYHFRELGVVHNQDVNLHSSRLSFKELLVYAADPAMHAASGNLLLDKRTTC 4787
SP|P0C6X7|R1AB_CVHSA              DGVPFVVSTGYHFRELGVVHNQDVNLHSSRLSFKELLVYAADPAMHAASGNLLLDKRTTC 4764
TR|Q6UZF5|Q6UZF5_CVHSA            DGVPFVVSTGYHFRELGVVHNQDVNLHSSRLSFKELLVYAADPAMHAASGNLLLDKRTTC 4764
TR|Q6UZF1|Q6UZF1_CVHSA            DGVPFVVSTGYHFRELGVVHNQDVNLHSSRLSFKELLVYAADPAMHAASGNLLLDKRTTC 4764
TR|Q6JH48|Q6JH48_CVHSA            DGVPFVVSTGYHFRELGVVHNQDVNLHSSRLSFKELLVYAADPAMHAASGNLLLDKRTTC 4764
TR|Q692E6|Q692E6_CVHSA            DGVPFVVSTGYHFRELGVVHNQDVNLHSSRLSFKELLVYAADPAMHAASGNLLLDKRTTC 4764
TR|A0A0K1YZY7|A0A0K1YZY7_CVHSA    DGVPFVVSTGYHFRELGVVHNQDVNLHSSRLSFKELLVYAADPAMHAASGNLLLDKRTTC 4764
SP|P0C6W2|R1AB_BCHK3              DGVPFVVSTGYHFRELGVVHNQDVNLHSSRLSFKELLVYAADPAMHAASGNLLLDKRTTC 4758
SP|P0C6W6|R1AB_BCRP3              DGVPFVVSTGYHFRELGVVHNQDVNLHSSRLSFKELLVYAADPAMHAASGNLLLDKRTTC 4762
SP|P0C6V9|R1AB_BC279              DGVPFVVSTGYHFRELGVVHNQDVNLHSSRLSFKELLVYAADPAMHAASGNLLLDKRTTC 4770
TR|A0A0U1WHI4|A0A0U1WHI4_CVHSA    DGVPFVVSTGYHFRELGVVHNQDVNLHSSRLSFKELLVYAADPAMHAASGNLLLDKRTTC 4759
TR|A0A0U1WHG0|A0A0U1WHG0_CVHSA    DGVPFVVSTGYHFRELGVVHNQDVNLHSSRLSFKELLVYAADPAMHAASGNLLLDKRTTC 4759
TR|A0A166ZL34|A0A166ZL34_9NIDO    DGVPFVVSTGYHFRELGVVHNQDVNLHSSRLSFKELLVYAADPAMHAASGNLLLDKRTTC 4566
TR|R9QTB2|R9QTB2_CVHSA            ------------------------------------------------------------
TR|R9QTH2|R9QTH2_CVHSA            ------------------------------------------------------------
SP|P0C6U8|R1A_CVHSA               ------------------------------------------------------------
TR|Q6JH47|Q6JH47_CVHSA            ------------------------------------------------------------
TR|Q692E5|Q692E5_CVHSA            ------------------------------------------------------------
SP|P0C6F8|R1A_BCHK3               ------------------------------------------------------------
TR|A0A0K1Z0N1|A0A0K1Z0N1_CVHSA    ------------------------------------------------------------
SP|P0C6F5|R1A_BC279               ------------------------------------------------------------
SP|P0C6T7|R1A_BCRP3               ------------------------------------------------------------

QHN73794                          FSVAALTNNVAFQTVKPGNFNKDFYDFAVSKGFFKEGSSVELKHFFFAQDGNAAISDYDY 4847
SP|P0C6X7|R1AB_CVHSA              FSVAALTNNVAFQTVKPGNFNKDFYDFAVSKGFFKEGSSVELKHFFFAQDGNAAISDYDY 4824
TR|Q6UZF5|Q6UZF5_CVHSA            FSVAALTNNVAFQTVKPGNFNKDFYDFAVSKGFFKEGSSVELKHFFFAQDGNAAISDYDY 4824
TR|Q6UZF1|Q6UZF1_CVHSA            FSVAALTNNVAFQTVKPGNFNKDFYDFAVSKGFFKEGSSVELKHFFFAQDGNAAISDYDY 4824
TR|Q6JH48|Q6JH48_CVHSA            FSVAALTNNVAFQTVKPGNFNKDFYDFAVSKGFFKEGSSVELKHFFFAQDGNAAISDYDY 4824
TR|Q692E6|Q692E6_CVHSA            FSVAALTNNVAFQTVKPGNFNKDFYDFAVSKGFFKEGSSVELKHFFFAQDGNAAISDYDY 4824
TR|A0A0K1YZY7|A0A0K1YZY7_CVHSA    FSVAALTNNVAFQTVKPGNFNKDFYDFAVSKGFFKEGSSVELKHFFFAQDGNAAISDYDY 4824
SP|P0C6W2|R1AB_BCHK3              FSVAALTNNVAFQTVKPGNFNKDFYDFAVSKGFFKEGSSVELKHFFFAQDGNAAISDYDY 4818
SP|P0C6W6|R1AB_BCRP3              FSVAALTNNVAFQTVKPGNFNKDFYDFAVSKGFFKEGSSVELKHFFFAQDGNAAISDYDY 4822
SP|P0C6V9|R1AB_BC279              FSVAALTNNVAFQTVKPGNFNKDFYDFAVSKGFFKEGSSVELKHFFFAQDGNAAISDYDY 4830
TR|A0A0U1WHI4|A0A0U1WHI4_CVHSA    FSVAALTNNVSFQTVKPGNFNKDFYDFAVSKGFFKEGSSVELKHFFFAQDGNAAISDYDY 4819
TR|A0A0U1WHG0|A0A0U1WHG0_CVHSA    FSVAALTNNVSFQTVKPGNFNKDFYDFAVSKGFFKEGSSVELKHFFFAQDGNAAISDYDY 4819
TR|A0A166ZL34|A0A166ZL34_9NIDO    FSVAALTNNVSFQTVKPGNFNKDFYDFAVSKGFFKEGSSVELKHFFFAQDGNAAISDYDY 4626
TR|R9QTB2|R9QTB2_CVHSA            ------------------------------------------------------------
TR|R9QTH2|R9QTH2_CVHSA            ------------------------------------------------------------
SP|P0C6U8|R1A_CVHSA               ------------------------------------------------------------
TR|Q6JH47|Q6JH47_CVHSA            ------------------------------------------------------------
TR|Q692E5|Q692E5_CVHSA            ------------------------------------------------------------
SP|P0C6F8|R1A_BCHK3               ------------------------------------------------------------
TR|A0A0K1Z0N1|A0A0K1Z0N1_CVHSA    ------------------------------------------------------------
SP|P0C6F5|R1A_BC279               ------------------------------------------------------------
SP|P0C6T7|R1A_BCRP3               ------------------------------------------------------------

QHN73794                          YRYNLPTMCDIRQLLFVVEVVDKYFDCYDGGCINANQVIVNNLDKSAGFPFNKWGKARLY 4907
SP|P0C6X7|R1AB_CVHSA              YRYNLPTMCDIRQLLFVVEVVDKYFDCYDGGCINANQVIVNNLDKSAGFPFNKWGKARLY 4884
TR|Q6UZF5|Q6UZF5_CVHSA            YRYNLPTMCDIRQLLFVVEVVDKYFDCYDGGCINANQVIVNNLDKSAGFPFNKWGKARLY 4884
TR|Q6UZF1|Q6UZF1_CVHSA            YRYNLPTMCDIRQLLFVVEVVDKYFDCYDGGCINANQVIVNNLDKSAGFPFNKWGKARLY 4884
TR|Q6JH48|Q6JH48_CVHSA            YRYNLPTMCDIRQLLFVVEVVDKYFDCYDGGCINANQVIVNNLDKSAGFPFNKWGKARLY 4884
TR|Q692E6|Q692E6_CVHSA            YRYNLPTMCDIRQLLFVVEVVDKYFDCYDGGCINANQVIVNNLDKSAGFPFNKWGKARLY 4884
TR|A0A0K1YZY7|A0A0K1YZY7_CVHSA    YRYNLPTMCDIRQLLFVVEVVDKYFDCYDGGCINANQVIVNNLDKSAGFPFNKWGKARLY 4884
SP|P0C6W2|R1AB_BCHK3              YRYNLPTMCDIRQLLFVVEVVDKYFDCYDGGCINANQVIVNNLDKSAGFPFNKWGKARLY 4878
SP|P0C6W6|R1AB_BCRP3              YRYNLPTMCDIRQLLFVVEVVDKYFDCYDGGCINANQVIVNNLDKSAGFPFNKWGKARLY 4882
SP|P0C6V9|R1AB_BC279              YRYNLPTMCDIRQLLFVVEVVDKYFDCYDGGCINANQVIVNNLDKSAGFPFNKWGKARLY 4890
TR|A0A0U1WHI4|A0A0U1WHI4_CVHSA    YRYNLPTMCDIRQLLFVVEVVDKYFDCYDGGCINANQVIVNNLDKSAGFPFNKWGKARLY 4879
TR|A0A0U1WHG0|A0A0U1WHG0_CVHSA    YRYNLPTMCDIRQLLFVVEVVDKYFDCYDGGCINANQVIVNNLDKSAGFPFNKWGKARLY 4879
TR|A0A166ZL34|A0A166ZL34_9NIDO    YRYNLPTMCDIRQLLFVVEVVDKYFDCYDGGCINANQVIVNNLDKSAGFPFNKWGKARLY 4686
TR|R9QTB2|R9QTB2_CVHSA            ------------------------------------------------------------
TR|R9QTH2|R9QTH2_CVHSA            ------------------------------------------------------------
SP|P0C6U8|R1A_CVHSA               ------------------------------------------------------------
TR|Q6JH47|Q6JH47_CVHSA            ------------------------------------------------------------
TR|Q692E5|Q692E5_CVHSA            ------------------------------------------------------------
SP|P0C6F8|R1A_BCHK3               ------------------------------------------------------------
TR|A0A0K1Z0N1|A0A0K1Z0N1_CVHSA    ------------------------------------------------------------
SP|P0C6F5|R1A_BC279               ------------------------------------------------------------
SP|P0C6T7|R1A_BCRP3               ------------------------------------------------------------
```

```
QHN73794                          YDSMSYEDQDALFAYTKRNVIPTITQMNLKYAISAKNRARTVAGVSICSTMTNRQFHQKL 4967
SP|P0C6X7|R1AB_CVHSA              YDSMSYEDQDALFAYTKRNVIPTITQMNLKYAISAKNRARTVAGVSICSTMTNRQFHQKL 4944
TR|Q6UZF5|Q6UZF5_CVHSA            YDSMSYEDQDALFAYTKRNVIPTITQMNLKYAISAKNRARTVAGVSICSTMTNRQFHQKL 4944
TR|Q6UZF1|Q6UZF1_CVHSA            YDSMSYEDQDALFAYTKRNVIPTITQMNLKYAISAKNRARTVAGVSICSTMTNRQFHQKL 4944
TR|Q6JH48|Q6JH48_CVHSA            YDSMSYEDQDALFAYTKRNVIPTITQMNLKYAISAKNRARTVAGVSICSTMTNRQFHQKL 4944
TR|Q692E6|Q692E6_CVHSA            YDSMSYEDQDALFAYTKRNVIPTITQMNLKYAISAKNRARTVAGVSICSTMTNRQFHQKL 4944
TR|A0A0K1YZY7|A0A0K1YZY7_CVHSA    YDSMSYEDQDALFAYTKRNVIPTITQMNLKYAISAKNRARTVAGVSICSTMTNRQFHQKL 4944
SP|P0C6W2|R1AB_BCHK3              YDSMSYEDQDALFAYTKRNVIPTITQMNLKYAISAKNRARTVAGVSICSTMTNRQFHQKL 4938
SP|P0C6W6|R1AB_BCRP3              YDSMSYEDQDALFAYTKRNVIPTITQMNLKYAISAKNRARTVAGVSICSTMTNRQFHQKL 4942
SP|P0C6V9|R1AB_BC279              YDSMSYEDQDVLFAYTKRNVIPTITQMNLKYAISAKNRARTVAGVSICSTMTNRQFHQKL 4950
TR|A0A0U1WHI4|A0A0U1WHI4_CVHSA    YDSMSYEDQDALFAYTKRNVLPTITQMNLKYAISAKNRARTVAGVSICSTMTNRQFHQKL 4939
TR|A0A0U1WHG0|A0A0U1WHG0_CVHSA    YDSMSYEDQDALFAYTKRNVLPTITQMNLKYAISAKNRARTVAGVSICSTMTNRQFHQKL 4939
TR|A0A166ZL34|A0A166ZL34_9NIDO    YDSMSYEDQDALFAYTKRNVLPTITQMNLKYAISAKNRARTVAGVSICSTMTNRQFHQKL 4746
TR|R9QTB2|R9QTB2_CVHSA            ------------------------------------------------------------
TR|R9QTH2|R9QTH2_CVHSA            ------------------------------------------------------------
SP|P0C6U8|R1A_CVHSA               ------------------------------------------------------------
TR|Q6JH47|Q6JH47_CVHSA            ------------------------------------------------------------
TR|Q692E5|Q692E5_CVHSA            ------------------------------------------------------------
SP|P0C6F8|R1A_BCHK3               ------------------------------------------------------------
TR|A0A0K1Z0N1|A0A0K1Z0N1_CVHSA    ------------------------------------------------------------
SP|P0C6F5|R1A_BC279               ------------------------------------------------------------
SP|P0C6T7|R1A_BCRP3               ------------------------------------------------------------

QHN73794                          LKSIAATRGATVVIGTSKFYGGWHNMLKTVYSDVENPHLMGWDYPKCDRAMPNMLRIMAS 5027
SP|P0C6X7|R1AB_CVHSA              LKSIAATRGATVVIGTSKFYGGWHNMLKTVYSDVETPHLMGWDYPKCDRAMPNMLRIMAS 5004
TR|Q6UZF5|Q6UZF5_CVHSA            LKSIAATRGATVVIGTSKFYGGWHNMLKTVYSDVETPHLMGWDYPKCDRAMPNMLRIMAS 5004
TR|Q6UZF1|Q6UZF1_CVHSA            LKSIAATRGATVVIGTSKFYGGWHNMLKTVYSDVETPHLMGWDYPKCDRAMPNMLRIMAS 5004
TR|Q6JH48|Q6JH48_CVHSA            LKSIAATRGATVVIGTSKFYGGWHNMLKTVYSDVETPHLMGWDYPKCDRAMPNMLRIMAS 5004
TR|Q692E6|Q692E6_CVHSA            LKSIAATRGATVVIGTSKFYGGWHNMLKTVYSDVETPHLMGWDYPKCDRAMPNMLRIMAS 5004
TR|A0A0K1YZY7|A0A0K1YZY7_CVHSA    LKSIAATRGATVVIGTSKFYGGWHNMLKTVYSDVETPHLMGWDYPKCDRAMPNMLRIMAS 5004
SP|P0C6W2|R1AB_BCHK3              LKSIAATRGATVVIGTSKFYGGWHNMLKTVYSDVESPHLMGWDYPKCDRAMPNMLRIMAS 4998
SP|P0C6W6|R1AB_BCRP3              LKSIAATRGATVVIGTSKFYGGWHNMLKTVYSDVETPHLMGWDYPKCDRAMPNMLRIMAS 5002
SP|P0C6V9|R1AB_BC279              LKSIAATRGATVVIGTSKFYGGWHNMLKTVYSDVETPHLMGWDYPKCDRAMPNMLRIMAS 5010
TR|A0A0U1WHI4|A0A0U1WHI4_CVHSA    LKSIAATRGATVVIGTSKFYGGWHNMLKTVYSDVETPYLMGWDYPKCDRAMPNMLRIMAF 4999
TR|A0A0U1WHG0|A0A0U1WHG0_CVHSA    LKSIAATRGATVVIGTSKFYGGWHNMLKTVYSDVETPHLMGWDYPKCDRAMPNMLRIMAS 4999
TR|A0A166ZL34|A0A166ZL34_9NIDO    LKSIAATRGATVVIGTSKFYGGWHNMLKTVYSDVETPHLMGWDYPKCDRAMPNMLRIMAS 4806
TR|R9QTB2|R9QTB2_CVHSA            ------------------------------------------------------------
TR|R9QTH2|R9QTH2_CVHSA            ------------------------------------------------------------
SP|P0C6U8|R1A_CVHSA               ------------------------------------------------------------
TR|Q6JH47|Q6JH47_CVHSA            ------------------------------------------------------------
TR|Q692E5|Q692E5_CVHSA            ------------------------------------------------------------
SP|P0C6F8|R1A_BCHK3               ------------------------------------------------------------
TR|A0A0K1Z0N1|A0A0K1Z0N1_CVHSA    ------------------------------------------------------------
SP|P0C6F5|R1A_BC279               ------------------------------------------------------------
SP|P0C6T7|R1A_BCRP3               ------------------------------------------------------------

QHN73794                          LVLARKHTTCCSLSHRFYRLANECAQVLSEMVMCGGSLYVKPGGTSSGDATTAYANSVFN 5087
SP|P0C6X7|R1AB_CVHSA              LVLARKHNTCCNLSHRFYRLANECAQVLSEMVMCGGSLYVKPGGTSSGDATTAYANSVFN 5064
TR|Q6UZF5|Q6UZF5_CVHSA            LVLARKHNTCCNLSHRFYRLANECAQVLSEMVMCGGSLYVKPGGTSSGDATTAYANSVFN 5064
TR|Q6UZF1|Q6UZF1_CVHSA            LVLARKHNTCCNLSHRFYRLANECAQVLSEMVMCGGSLYVKPGGTSSGDATTAYANSVFN 5064
TR|Q6JH48|Q6JH48_CVHSA            LVLARKHNTCCNLSHRFYRLANECAQVLSEMVMCGGSLYVKPGGTSSGDATTAYANSVFN 5064
TR|Q692E6|Q692E6_CVHSA            LVLARKHNTCCNLSHRFYRLANECAQVLSEMVMCGGSLYVKPGGTSSGDATTAYANSVFN 5064
TR|A0A0K1YZY7|A0A0K1YZY7_CVHSA    LVLARKHSTCCNLSHRFYRLANECAQVLSEMVMCGGSLYVKPGGTSSGDATTAYANSVFN 5064
SP|P0C6W2|R1AB_BCHK3              LILARKHSTCCNLSHRFYRLANECAQVLSEMVMCGGSLYVKPGGTSSGDATTAYANSVFN 5058
SP|P0C6W6|R1AB_BCRP3              LVLARKHSTCCNLSHRFYRLANECAQVLSEMVMCGGSLYVKPGGTSSGDATTAYANSVFN 5062
SP|P0C6V9|R1AB_BC279              LVLARKHSTCCNLSHRFYRLANECAQVLSEMVMCGGSLYVKPGGTSSGDATTAYANSVFN 5070
TR|A0A0U1WHI4|A0A0U1WHI4_CVHSA    LVFSRKHSTCCNLSHRFYRLANECAQVLSEMVMCGGSLYVKPGGTSSGDATTAYANSVFN 5059
TR|A0A0U1WHG0|A0A0U1WHG0_CVHSA    LVLARKHSTCCNLSHRFYRLANECAQVLSEMVMCGGSLYVKPGGTSSGDATTAYANSVFN 5059
TR|A0A166ZL34|A0A166ZL34_9NIDO    LVLARKHSTCCNLSHRFYRLANECAQVLSEMVMCGGSLYVKPGGTSSGDATTAYANSVFN 4866
TR|R9QTB2|R9QTB2_CVHSA            ------------------------------------------------------------
TR|R9QTH2|R9QTH2_CVHSA            ------------------------------------------------------------
SP|P0C6U8|R1A_CVHSA               ------------------------------------------------------------
TR|Q6JH47|Q6JH47_CVHSA            ------------------------------------------------------------
TR|Q692E5|Q692E5_CVHSA            ------------------------------------------------------------
SP|P0C6F8|R1A_BCHK3               ------------------------------------------------------------
TR|A0A0K1Z0N1|A0A0K1Z0N1_CVHSA    ------------------------------------------------------------
SP|P0C6F5|R1A_BC279               ------------------------------------------------------------
SP|P0C6T7|R1A_BCRP3               ------------------------------------------------------------
```

```
QHN73794                        ICQAVTANVNALLSTDGNKIADKYVRNLQHRLYECLYRNRDVDTDFVNEFYAYLRKHFSM 5147
SP|P0C6X7|R1AB_CVHSA            ICQAVTANVNALLSTDGNKIADKYVRNLQHRLYECLYRNRDVDHEFVDEFYAYLRKHFSM 5124
TR|Q6UZF5|Q6UZF5_CVHSA          ICQAVTANVNALLSTDGNKIADKYVRNLQHRLYECLYRNRDVDHEFVDEFYAYLRKHFSM 5124
TR|Q6UZF1|Q6UZF1_CVHSA          ICQAVTANVNALLSTDGNKIADKYVRNLQHRLYECLYRNRDVDHEFVDEFYAYLRKHFSM 5124
TR|Q6JH48|Q6JH48_CVHSA          ICQAVTANVNALLSTDGNKIADKYVRNLQHRLYECLYRNRDVDHEFVDEFYAYLRKHFSM 5124
TR|Q692E6|Q692E6_CVHSA          ICQAVTANVNALLSTDGNKIADKYVRNLQHRLYECLYRNRDVDHEFVDEFYAYLRKHFSM 5124
TR|A0A0K1YZY7|A0A0K1YZY7_CVHSA ICQAVTANVNALLSTDGNKIADKYVRNLQHRLYECLYRNRDVDHEFVDEFYAYLRKHFSM 5124
SP|P0C6W2|R1AB_BCHK3            ICQAVTANVNALLSTDGNKIADKYVRNLQHRLYECLYRNRDVDHEFVDEFYAYLRKHFSM 5118
SP|P0C6W6|R1AB_BCRP3            ICQAVTANVNALLSTDGNKIADKYVRNLQHRLYECLYRNRDVDHEFVGEFYAYLRKHFSM 5122
SP|P0C6V9|R1AB_BC279            ICQAVTANVNALLSTDGNKIADKYVRNLQHRLYECLYRNRDVDHEFVDEFYAYLRKHFSM 5130
TR|A0A0U1WHI4|A0A0U1WHI4_CVHSA ICQAVTANVNALLSTDGNKIADKYVRNLQHRLYECLYRNRDVDHEFVDEFYAYLRKHFSM 5119
TR|A0A0U1WHG0|A0A0U1WHG0_CVHSA ICQAVTANVNALLSTDGNKIADKYVRNIQHRLYECLYRNRDVDHEFVDEFYAYLRKHFSM 5119
TR|A0A166ZL34|A0A166ZL34_9NIDO ICQAVTANVNALLSTDGNKIADKYVRNIQHRLYECLYRNRDVDHEFVDEFYAYLRKHFSM 4926
TR|R9QTB2|R9QTB2_CVHSA          ------------------------------------------------------------
TR|R9QTH2|R9QTH2_CVHSA          ------------------------------------------------------------
SP|P0C6U8|R1A_CVHSA             ------------------------------------------------------------
TR|Q6JH47|Q6JH47_CVHSA          ------------------------------------------------------------
TR|Q692E5|Q692E5_CVHSA          ------------------------------------------------------------
SP|P0C6F8|R1A_BCHK3             ------------------------------------------------------------
TR|A0A0K1Z0N1|A0A0K1Z0N1_CVHSA ------------------------------------------------------------
SP|P0C6F5|R1A_BC279             ------------------------------------------------------------
SP|P0C6T7|R1A_BCRP3             ------------------------------------------------------------


QHN73794                        MILSDDAVVCFNSTYASQGLVASIKNFKSVLYYQNNVFMSEAKCWTETDLTKGPHEFCSQ 5207
SP|P0C6X7|R1AB_CVHSA            MILSDDAVVCYNSNYAAQGLVASIKNFKAVLYYQNNVFMSEAKCWTETDLTKGPHEFCSQ 5184
TR|Q6UZF5|Q6UZF5_CVHSA          MILSDDAVVCYNSNYAAQGLVASIKNFKAVLYYQNNVFMSEAKCWTETDLTKGPHEFCSQ 5184
TR|Q6UZF1|Q6UZF1_CVHSA          MILSDDAVVCYNSNYAAQGLVASIKNFKAVLYYQNNVFMSEAKCWTETDLTKGPHEFCSQ 5184
TR|Q6JH48|Q6JH48_CVHSA          MILSDDAVVCYNSNYAAQGLVASIKNFKAVLYYQNNVFMSEAKCWTETDLTKGPHEFCSQ 5184
TR|Q692E6|Q692E6_CVHSA          MILSDDAVVCYNSNYAAQGLVASIKNFKAVLYYQNNVFMSEAKCWTETDLTKGPHEFCSQ 5184
TR|A0A0K1YZY7|A0A0K1YZY7_CVHSA MILSDDAVVCYNSNYAAQGLVASIKNFKAVLYYQNNVFMSEAKCWTETDLTKGPHEFCSQ 5184
SP|P0C6W2|R1AB_BCHK3            MILSDDAVVCYNSNYAAQGLVASIKNFKAVLYYQNNVFMSEAKCWTETDLTRGPHEFCSQ 5178
SP|P0C6W6|R1AB_BCRP3            MILSDDAVVCYNSNYAAQGLVASIKNFKAVLYYQNNVFMSEAKCWTETDLTKGPHEFCSQ 5182
SP|P0C6V9|R1AB_BC279            MILSDDAVVCYNSNYAAQGLVASIKNFKAVHYYQNNVFMSEAKCWTETDLTKGPHEFCSQ 5190
TR|A0A0U1WHI4|A0A0U1WHI4_CVHSA MILSDDAVVCYNSNYAAQGLVASIKNFKAVLYYQNNVFMSEAKCWTETDLTKGPHEFCSQ 5179
TR|A0A0U1WHG0|A0A0U1WHG0_CVHSA MILSDDAVVCYNSNYAAQGLVASIKNFKAVLYYQNNVFMSEAKCWTETDLTKGPHEFCSQ 5179
TR|A0A166ZL34|A0A166ZL34_9NIDO MILSDDAVVCYNSNYAAQGLVASIKNFKAVLYYQNNVFMSEAKCWTETDLTKGPHEFCSQ 4986
TR|R9QTB2|R9QTB2_CVHSA          ------------------------------------------------------------
TR|R9QTH2|R9QTH2_CVHSA          ------------------------------------------------------------
SP|P0C6U8|R1A_CVHSA             ------------------------------------------------------------
TR|Q6JH47|Q6JH47_CVHSA          ------------------------------------------------------------
TR|Q692E5|Q692E5_CVHSA          ------------------------------------------------------------
SP|P0C6F8|R1A_BCHK3             ------------------------------------------------------------
TR|A0A0K1Z0N1|A0A0K1Z0N1_CVHSA ------------------------------------------------------------
SP|P0C6F5|R1A_BC279             ------------------------------------------------------------
SP|P0C6T7|R1A_BCRP3             ------------------------------------------------------------

                                                           _
QHN73794                        HTMLVKQGDDYVYLPYPDPSRILGAGCFVDDIVKTDGTLMIERFVSLAIDAYPLTKHPNQ 5267
SP|P0C6X7|R1AB_CVHSA            HTMLVKQGDDYVYLPYPDPSRILGAGCFVDDIVKTDGTLMIERFVSLAIDAYPLTKHPNQ 5244
TR|Q6UZF5|Q6UZF5_CVHSA          HTMLVKQGDDYVYLPYPDPSRILGAGCFVDDIVKTDGTLMIERFVSLAIDAYPLTKHPNQ 5244
TR|Q6UZF1|Q6UZF1_CVHSA          HTMLVKQGDDYVYLPYPDPSRILGAGCFVDDIVKTDGTLMIERFVSLAIDAYPLTKHPNQ 5244
TR|Q6JH48|Q6JH48_CVHSA          HTMLVKQGDDYVYLPYPDPSRILGAGCFVDDIVKTDGTLMIERFVSLAIDAYPLTKHPNQ 5244
TR|Q692E6|Q692E6_CVHSA          HTMLVKQGDDYVYLPYPDPSRILGAGCFVDDIVKTDGTLMIERFVSLAIDAYPLTKHPNQ 5244
TR|A0A0K1YZY7|A0A0K1YZY7_CVHSA HTMLVKQGDDYVYLPYPDPSRILGAGCFVDDIVKTDGTLMIERFVSLAIDAYPLTKHPNQ 5244
SP|P0C6W2|R1AB_BCHK3            HTMLVKQGDDYVYLPYPDPSRILGAGCFVDDIVKTDGTLMIERFVSLAIDAYPLTKHPNQ 5238
SP|P0C6W6|R1AB_BCRP3            HTMLVKQGDDYVYLPYPDPSRILGAGCFVDDIVKTDGTLMIERFVSLAIDAYPLTKHPNQ 5242
SP|P0C6V9|R1AB_BC279            HTMLVKQGDDYVYLPYPDPSRILGAGCFVDDIVKTDGTLMIERFVSLAIDAYPLTKHPNQ 5250
TR|A0A0U1WHI4|A0A0U1WHI4_CVHSA HTMLVKQGDDYVYLPYPDPSRILGAGCFVDDIVKTDGTLMIERFVSLAIDAYPLTKHPNQ 5239
TR|A0A0U1WHG0|A0A0U1WHG0_CVHSA HTMLVKQGDDYVYLPYPDPSRILGAGCFVDDIVKTDGTLMIERFVSLAIDAYPLTKHPNQ 5239
TR|A0A166ZL34|A0A166ZL34_9NIDO HTMLVKQGDDYVYLPYPDPSRILGAGCFVDDIVKTDGTLMIERFVSLAIDAYPLTKHPNQ 5046
TR|R9QTB2|R9QTB2_CVHSA          ------------------------------------------------------------
TR|R9QTH2|R9QTH2_CVHSA          ------------------------------------------------------------
SP|P0C6U8|R1A_CVHSA             ------------------------------------------------------------
TR|Q6JH47|Q6JH47_CVHSA          ------------------------------------------------------------
TR|Q692E5|Q692E5_CVHSA          ------------------------------------------------------------
SP|P0C6F8|R1A_BCHK3             ------------------------------------------------------------
TR|A0A0K1Z0N1|A0A0K1Z0N1_CVHSA ------------------------------------------------------------
SP|P0C6F5|R1A_BC279             ------------------------------------------------------------
SP|P0C6T7|R1A_BCRP3             ------------------------------------------------------------
```

```
QHN73794                       EYADVFHLYLQYIRKLHDELTGHMLDMYSVMLTNDNTSRYWEPEFYEAMYTPHTVLQAVG 5327
SP|P0C6X7|R1AB_CVHSA           EYADVFHLYLQYIRKLHDELTGHMLDMYSVMLTNDNTSRYWEPEFYEAMYTPHTVLQAVG 5304
TR|Q6UZF5|Q6UZF5_CVHSA         EYADVFHLYLQYIRKLHDELTGHMLDMYSVMLTNDNTSRYWEPEFYEAMYTPHTVLQAVG 5304
TR|Q6UZF1|Q6UZF1_CVHSA         EYADVFHLYLQYIRKLHDELTGHMLDMYSVMLTNDNTSRYWEPEFYEAMYTPHTVLQAVG 5304
TR|Q6JH48|Q6JH48_CVHSA         EYAAVFHLYLQYIRKLHDELTGHMLDMYSVMLTNDNTSRYWEPEFYEAMYTPHTVLQAVG 5304
TR|Q692E6|Q692E6_CVHSA         EYADVFHLYLQYIRKLHDELTGHMLDMYSVMLTNDNTSRYWEPEFYEAMYTPHTVLQAVG 5304
TR|A0A0K1YZY7|A0A0K1YZY7_CVHSA EYADVFHLYLQYIRKLHDELTGHMLDMYSVMLTNDNTSRYWEPEFYEAMYTPHTVLQAVG 5304
SP|P0C6W2|R1AB_BCHK3           EYADVFHLYLQYIRKLHDELTGHMLDMYSVMLTNDNTSRYWEPEFYEAMYTPHTVLQAVG 5298
SP|P0C6W6|R1AB_BCRP3           EYADVFHLYLQYIRKLHDELTGHMLDMYSVMLTNDNTSRYWEPEFYEAMYTPHTVLQAVG 5302
SP|P0C6V9|R1AB_BC279           EYADVFHLYLQYIRKLHDELTGHMLDMYSVMLTNDNTSRYWEPEFYEAMYTPHTVLQAVG 5310
TR|A0A0U1WHI4|A0A0U1WHI4_CVHSA EYADVFHLYLQYIRKLHDELTGHMLDMYSVMLTNDNTSRYWEPEFYEAMYTPHTILQAVG 5299
TR|A0A0U1WHG0|A0A0U1WHG0_CVHSA EYADVFHLYLQYIRKLHDELTGHMLDMYSVMLTNDNTSRYWEPEFYEAMYTPHTILQAVG 5299
TR|A0A166ZL34|A0A166ZL34_9NIDO EYADVFHLYLQYIRKLHDELTGHMLDMYSVMLTNDNTSRYWEPEFYEAMYTPHTILQAVG 5106
TR|R9QTB2|R9QTB2_CVHSA         ------------------------------------------------------------
TR|R9QTH2|R9QTH2_CVHSA         ------------------------------------------------------------
SP|P0C6U8|R1A_CVHSA            ------------------------------------------------------------
TR|Q6JH47|Q6JH47_CVHSA         ------------------------------------------------------------
TR|Q692E5|Q692E5_CVHSA         ------------------------------------------------------------
SP|P0C6F8|R1A_BCHK3            ------------------------------------------------------------
TR|A0A0K1Z0N1|A0A0K1Z0N1_CVHSA ------------------------------------------------------------
SP|P0C6F5|R1A_BC279            ------------------------------------------------------------
SP|P0C6T7|R1A_BCRP3            ------------------------------------------------------------

QHN73794                       ACVLCNSQTSLRCGACIRRPFLCCKCCYDHVISTSHKLVLSVNPYVCNAPGCDVTDVTQL 5387
SP|P0C6X7|R1AB_CVHSA           ACVLCNSQTSLRCGACIRRPFLCCKCCYDHVISTSHKLVLSVNPYVCNAPGCDVTDVTQL 5364
TR|Q6UZF5|Q6UZF5_CVHSA         ACVLCNSQTSLRCGACIRRPFLCCKCCYDHVISTSHKLVLSVNPYVCNAPGCDVTDVTQL 5364
TR|Q6UZF1|Q6UZF1_CVHSA         ACVLCNSQTSLRCGACIRRPFLCCKCCYDHVISTSHKLVLSVNPYVCNAPGCDVTDVTQL 5364
TR|Q6JH48|Q6JH48_CVHSA         ACVLCNSQTSLRCGACIRRPFLCCKCCYDHVISTSHKLVLSVNPYVCNAPGCDVTDVTQL 5364
TR|Q692E6|Q692E6_CVHSA         ACVLCNSQTSLRCGACIRRPFLCCKCCYDHVISTSHKLVLSVNPYVCNAPGCDVTDVTQL 5364
TR|A0A0K1YZY7|A0A0K1YZY7_CVHSA ACVLCNSQTSLRCGACIRRPFLCCKCCYDHVISTSHKLVLSVNPYVCNAPGCDVTDVTQL 5364
SP|P0C6W2|R1AB_BCHK3           ACVLCNSQTSLRCGACIRRPFLCCKCCYDHVISTSHKLVLSVNPYVCNAPGCDVTDVTQL 5358
SP|P0C6W6|R1AB_BCRP3           ACVLCNSQTSLRCGACIRRPFLCCKCCYDHVISTSHKLVLSVNPYVCNAPGCDVTDVTQL 5362
SP|P0C6V9|R1AB_BC279           ACVLCNSQTSLRCGACIRRPFLCCKCCYDHVISTSHKLVLSVNPYVCNAPGCDVTDVTQL 5370
TR|A0A0U1WHI4|A0A0U1WHI4_CVHSA ACVLCNSQTSLRCGACIRRPFLCCKCCYDHVISTSHKLVLSVNPYVCNAPGCDVTDVTQL 5359
TR|A0A0U1WHG0|A0A0U1WHG0_CVHSA ACVLCNSQTSLRCGACIRRPFLCCKCCYDHVISTSHKLVLSVNPYVCNAPGCDVTDVTQL 5359
TR|A0A166ZL34|A0A166ZL34_9NIDO ACVLCNSQTSLRCGACIRRPFLCCKCCYDHVISTSHKLVLSVNPYVCNAPGCDVTDVTQL 5166
TR|R9QTB2|R9QTB2_CVHSA         ------------------------------------------------------------
TR|R9QTH2|R9QTH2_CVHSA         ------------------------------------------------------------
SP|P0C6U8|R1A_CVHSA            ------------------------------------------------------------
TR|Q6JH47|Q6JH47_CVHSA         ------------------------------------------------------------
TR|Q692E5|Q692E5_CVHSA         ------------------------------------------------------------
SP|P0C6F8|R1A_BCHK3            ------------------------------------------------------------
TR|A0A0K1Z0N1|A0A0K1Z0N1_CVHSA ------------------------------------------------------------
SP|P0C6F5|R1A_BC279            ------------------------------------------------------------
SP|P0C6T7|R1A_BCRP3            ------------------------------------------------------------

QHN73794                       YLGGMSYYCKSHKPPISFPLCANGQVFGLYKNTCVGSDNVTDFNAIATCDWTNAGDYILA 5447
SP|P0C6X7|R1AB_CVHSA           YLGGMSYYCKSHKPPISFPLCANGQVFGLYKNTCVGSDNVTDFNAIATCDWTNAGDYILA 5424
TR|Q6UZF5|Q6UZF5_CVHSA         YLGGMSYYCKSHKPPISFPLCANGQVFGLYKNTCVGSDNVTDFNAIATCDWTNAGDYILA 5424
TR|Q6UZF1|Q6UZF1_CVHSA         YLGGMSYYCKSHKPPISFPLCANGQVFGLYKNTCVGSDNVTDFNAIATCDWTNAGDYILA 5424
TR|Q6JH48|Q6JH48_CVHSA         YLGGMSYYCKSHKPPISFPLCANGQVFGLYKNTCVGSDNVTDFNAIATCDWTNAGDYILA 5424
TR|Q692E6|Q692E6_CVHSA         YLGGMSYYCKSHKPPISFPLCANGQVFGLYKNTCVGSDNVTDFNAIATCDWTNAGDYILA 5424
TR|A0A0K1YZY7|A0A0K1YZY7_CVHSA YLGGMSYYCKSHKPPISFPLCANGQVFGLYKNTCVGSDNVTDFNAIATCDWTNAGDYILA 5424
SP|P0C6W2|R1AB_BCHK3           YLGGMSYYCKSHKPPISFPLCANGQVFGLYKNTCVGSDNVTDFNAIATCDWTNAGDYILA 5418
SP|P0C6W6|R1AB_BCRP3           YLGGMSYYCKSHKPPISFPLCANGQVFGLYKNTCVGSDNVTDFNAIATCDWTNAGDYILA 5422
SP|P0C6V9|R1AB_BC279           YLGGMSYYCKLHKPPISFPLCANGQVFGLYKNTCVGSDNVTDFNAIATCDWTNAGDYILA 5430
TR|A0A0U1WHI4|A0A0U1WHI4_CVHSA YLGGMSYYCKSHKPPISFPLCANGQVFGLYKNTCVGSDNVTDFNAIATCDWTNAGDYILA 5419
TR|A0A0U1WHG0|A0A0U1WHG0_CVHSA YLGGMSYYCKSHKPSISFPLCANGQVFGLYKNTCVGSDNVTDFNAIATCDWTNAGDYILA 5419
TR|A0A166ZL34|A0A166ZL34_9NIDO YLGGMSYYCKSHKPSISFPLCANGQVFGLYKNTCVGSDNVTDFNAIATCDWTNAGDYILA 5226
TR|R9QTB2|R9QTB2_CVHSA         ------------------------------------------------------------
TR|R9QTH2|R9QTH2_CVHSA         ------------------------------------------------------------
SP|P0C6U8|R1A_CVHSA            ------------------------------------------------------------
TR|Q6JH47|Q6JH47_CVHSA         ------------------------------------------------------------
TR|Q692E5|Q692E5_CVHSA         ------------------------------------------------------------
SP|P0C6F8|R1A_BCHK3            ------------------------------------------------------------
TR|A0A0K1Z0N1|A0A0K1Z0N1_CVHSA ------------------------------------------------------------
SP|P0C6F5|R1A_BC279            ------------------------------------------------------------
SP|P0C6T7|R1A_BCRP3            ------------------------------------------------------------
```

```
QHN73794                           NTCTERLKLFAAETLKATEETFKLSYGIATVREVLSDRELHLSWEVGKPRPPLNRNYVFT 5507
SP|P0C6X7|R1AB_CVHSA               NTCTERLKLFAAETLKATEETFKLSYGIATVREVLSDRELHLSWEVGKPRPPLNRNYVFT 5484
TR|Q6UZF5|Q6UZF5_CVHSA             NTCTERLKLFAAETLKATEETFKLSYGIATVREVLSDRELHLSWEVGKPRPPLNRNYVFT 5484
TR|Q6UZF1|Q6UZF1_CVHSA             NTCTERLKLFAAETLKATEETFKLSYGIATVREVLSDRELHLSWEVGKPRPPLNRNYVFT 5484
TR|Q6JH48|Q6JH48_CVHSA             NTCTERLKLFAAETLKATEETFKLSYGIATVREVLSDRELHLSWEVGKPRPPLNRNYVFT 5484
TR|Q692E6|Q692E6_CVHSA             NTCTERLKLFAAETLKATEETFKLSYGIATVREVLSDRELHLSWEVGKPRPPLNRNYVFT 5484
TR|A0A0K1YZY7|A0A0K1YZY7_CVHSA NTCTERLKLFAAETLKATEETFKLSYGIATVREVLSDRELHLSWEVGKPRPPLNRNYVFT 5484
SP|P0C6W2|R1AB_BCHK3               NTCTERLKLFAAETLKATEETFKLSYGIATVREVLSDRELYLSWEVGKPRPPLNRNYVFT 5478
SP|P0C6W6|R1AB_BCRP3               NTCTERLKLFAAETLKATEETFKLSYGIATVREVLSDRELHLSWEVGKPRPPLNRNYVFT 5482
SP|P0C6V9|R1AB_BC279               NTCTERLKLFAAETLKATEETFKLSYGIATVREVLSDRELHLSWEVGKPRPPLNRNYVFT 5490
TR|A0A0U1WHI4|A0A0U1WHI4_CVHSA NTCTERLKLFAAETLKATEETFKLSYGIATVREVLSDKELHLSWEVGKPRPPLNRNYVFT 5479
TR|A0A0U1WHG0|A0A0U1WHG0_CVHSA NTCTERLKLFAAETLKATEETFKLSYGIATVREVLSDKELYLSWEVGKPRPPLNRNYVFT 5479
TR|A0A166ZL34|A0A166ZL34_9NIDO NTCTERLKLFAAETLKATEETFKLSYGIATVREVLSDKELYLSWEVGKPRPPLNRNYVFT 5286
TR|R9QTB2|R9QTB2_CVHSA             ------------------------------------------------------------
TR|R9QTH2|R9QTH2_CVHSA             ------------------------------------------------------------
SP|P0C6U8|R1A_CVHSA                ------------------------------------------------------------
TR|Q6JH47|Q6JH47_CVHSA             ------------------------------------------------------------
TR|Q692E5|Q692E5_CVHSA             ------------------------------------------------------------
SP|P0C6F8|R1A_BCHK3                ------------------------------------------------------------
TR|A0A0K1Z0N1|A0A0K1Z0N1_CVHSA ------------------------------------------------------------
SP|P0C6F5|R1A_BC279                ------------------------------------------------------------
SP|P0C6T7|R1A_BCRP3                ------------------------------------------------------------

QHN73794                           GYRVTKNSKVQIGEYTFEKGDYGDAVVYRGTTTYKLNVGDYFVLTSHTVMPLSAPTLVPQ 5567
SP|P0C6X7|R1AB_CVHSA               GYRVTKNSKVQIGEYTFEKGDYGDAVVYRGTTTYKLNVGDYFVLTSHTVMPLSAPTLVPQ 5544
TR|Q6UZF5|Q6UZF5_CVHSA             GYRVTKNSKVQIGEYTFEKGDYGDAVVYRGTTTYKLNVGDYFVLTSHTVMPLSAPTLVPQ 5544
TR|Q6UZF1|Q6UZF1_CVHSA             GYRVTKNSKVQIGEYTFEKGDYGDAVVYRGTTTYKLNVGDYFVLTSHTVMPLSAPTLVSQ 5544
TR|Q6JH48|Q6JH48_CVHSA             GYRVTKNSKVQIGEYTFEKGDYGDAVVYRGTTTYKLNVGDYFVLTSHTVMPLSAPTLVSQ 5544
TR|Q692E6|Q692E6_CVHSA             GYRVTKNSKVQIGEYTFEKGDYGDAVVYRGTTTYKLNVGDYFVLTSHTVMPLSAPTLVPQ 5544
TR|A0A0K1YZY7|A0A0K1YZY7_CVHSA GYRVTKNSKVQIGEYTFEKGDYGDAVVYRGTTTYKLNVGDYFVLTSHTVMPLSAPTLVPQ 5544
SP|P0C6W2|R1AB_BCHK3               GYRVTKNSKVQIGEYTFEKGDYGDAVVYRGTTTYKLNVGDYFVLTSHTVMPLSAPTLVPQ 5538
SP|P0C6W6|R1AB_BCRP3               GYRVTKNSKVQIGEYTFEKGDYGDAVVYRGTTTYKLNIGDYFVLTSHTVMPLSAPTLVPQ 5542
SP|P0C6V9|R1AB_BC279               GYRVTKNSKVQIGEYTFEKGDYGDAVVYRGTTTYKLNVGDYFVLTSHTVMPLSAPTLVPQ 5550
TR|A0A0U1WHI4|A0A0U1WHI4_CVHSA GYRVTKNSKVQIGEYTFEKGDYGDAVVYRGTTTYKLNVGDYFVLTSHTVMPLSAPTLVPQ 5539
TR|A0A0U1WHG0|A0A0U1WHG0_CVHSA GYRVTKNSKVQIGEYTFEKGDYGDAVVYRGTTTYKLNVGDYFVLTSHTVMPLSAPTLVPQ 5539
TR|A0A166ZL34|A0A166ZL34_9NIDO GYRVTKNSKVQIGEYTFEKGDYGDAVVYRGTTTYKLNVGDYFVLTSHTVMPLSAPTLVPQ 5346
TR|R9QTB2|R9QTB2_CVHSA             ------------------------------------------------------------
TR|R9QTH2|R9QTH2_CVHSA             ------------------------------------------------------------
SP|P0C6U8|R1A_CVHSA                ------------------------------------------------------------
TR|Q6JH47|Q6JH47_CVHSA             ------------------------------------------------------------
TR|Q692E5|Q692E5_CVHSA             ------------------------------------------------------------
SP|P0C6F8|R1A_BCHK3                ------------------------------------------------------------
TR|A0A0K1Z0N1|A0A0K1Z0N1_CVHSA ------------------------------------------------------------
SP|P0C6F5|R1A_BC279                ------------------------------------------------------------
SP|P0C6T7|R1A_BCRP3                ------------------------------------------------------------

QHN73794                           EHYVRITGLYPTLNISDEFSSNVANYQKVGMQKYSTLQGPPGTGKSHFAIGLALYYPSAR 5627
SP|P0C6X7|R1AB_CVHSA               EHYVRITGLYPTLNISDEFSSNVANYQKVGMQKYSTLQGPPGTGKSHFAIGLALYYPSAR 5604
TR|Q6UZF5|Q6UZF5_CVHSA             EHYVRITGLYPTLNISDEFSSNVANYQKVGMQKYSTLQGPPGTGKSHFAIGLALYYPSAR 5604
TR|Q6UZF1|Q6UZF1_CVHSA             EHYVRITGLYPTLNISDEFSSNVANYQKVGMQKYSTLQGPPGTGKSHFAIGLALYYPSAR 5604
TR|Q6JH48|Q6JH48_CVHSA             EHYVRITGLYPTLNISDEFSSNVANYQKVGMQKYSTLQGPPGTGKSHFAIGLALYYPSAR 5604
TR|Q692E6|Q692E6_CVHSA             EHYVRITGLYPTLNISDEFSSNVANYQKVGMQKYSTLQGPPGTGKSHFAIGLALYYPSAR 5604
TR|A0A0K1YZY7|A0A0K1YZY7_CVHSA EYYVRITGLYPTLNISDEFSSNVANYQKVGMQKYSTLQGPPGTGKSHFAIGLALYYPSAR 5604
SP|P0C6W2|R1AB_BCHK3               EHYVRITGLYPTLNISNEFSSNVANYQKIGMQKYSTLQGPPGTGKSHFAIGLALYYPSAR 5598
SP|P0C6W6|R1AB_BCRP3               EHYVRITGLYPTLNISDEFSSNVANYQKVGMQKYSTLQGPPGTGKSHFAIGLALYYPSAR 5602
SP|P0C6V9|R1AB_BC279               EHYVRITGLYPTLNISDEFSSNVANYQKVGMQKYSTLQGPPGTGKSHFAIGLALYYPSAR 5610
TR|A0A0U1WHI4|A0A0U1WHI4_CVHSA EHYVRITGLYPTLNISEEFSSNVANYQKVGMQKYSTLQGPPGTGKSHFAIGLALYYPSAR 5599
TR|A0A0U1WHG0|A0A0U1WHG0_CVHSA EHYVRITGLYPTLNISEEFSSNVANYQKVGMQKYSTLQGPPGTGKSHFAIGLALYYPSAR 5599
TR|A0A166ZL34|A0A166ZL34_9NIDO EHYVRITGLYPTLNISEEFSSNVANYQKVGMQKYSTLQGPPGTGKSHFAIGLALYYPSAR 5406
TR|R9QTB2|R9QTB2_CVHSA             ------------------------------------------------------------
TR|R9QTH2|R9QTH2_CVHSA             ------------------------------------------------------------
SP|P0C6U8|R1A_CVHSA                ------------------------------------------------------------
TR|Q6JH47|Q6JH47_CVHSA             ------------------------------------------------------------
TR|Q692E5|Q692E5_CVHSA             ------------------------------------------------------------
SP|P0C6F8|R1A_BCHK3                ------------------------------------------------------------
TR|A0A0K1Z0N1|A0A0K1Z0N1_CVHSA ------------------------------------------------------------
SP|P0C6F5|R1A_BC279                ------------------------------------------------------------
SP|P0C6T7|R1A_BCRP3                ------------------------------------------------------------
```

```
QHN73794                        IVYTACSHAAVDALCEKALKYLPIDKCSRIIPARARVECFDKFKVNSTLEQYVFCTVNAL 5687
SP|P0C6X7|R1AB_CVHSA            IVYTACSHAAVDALCEKALKYLPIDKCSRIIPARARVECFDKFKVNSTLEQYVFCTVNAL 5664
TR|Q6UZF5|Q6UZF5_CVHSA          IVYTACSHAAVDALCEKALKYLPIDKCSRIIPARARVECFDKFKVNSTLEQYVFCTVNAL 5664
TR|Q6UZF1|Q6UZF1_CVHSA          IVYTACSHAAVDALCEKALKYLPIDKCSRIIPARARVECFDKFKVNSTLEQYVFCTVNAL 5664
TR|Q6JH48|Q6JH48_CVHSA          IVYTACSHAAVDALCEKALKYLPIDKCSRIIPARARVECFDKFKVNSTLEQYVFCTVNAL 5664
TR|Q692E6|Q692E6_CVHSA          IVYTACSHAAVDALCEKALKYLPIDKCSRIIPARARVECFDKFKVNSTLEQYVFCTVNAL 5664
TR|A0A0K1YZY7|A0A0K1YZY7_CVHSA  IVYTACSHAAVDALCEKALKYLPIDKCSRIIPARARVECFDKFKVNSTLEQYVFCTVNAL 5664
SP|P0C6W2|R1AB_BCHK3            IVYTACSHAAVDALCEKALKYLPIDKCSRIIPARARVECFDKFKVNSTLEQYVFCTVNAL 5658
SP|P0C6W6|R1AB_BCRP3            IVYTACSHAAVDALCEKALKYLPIDKCSRIIPARARVECFDKFKVNSTLEQYVFCTVNAL 5662
SP|P0C6V9|R1AB_BC279            IVYTACSHAAVDALCEKALKYLPIDKCSRIIPARARVECFDKFKVNSTLEQYVFCTVNAL 5670
TR|A0A0U1WHI4|A0A0U1WHI4_CVHSA  IVYTACSHAAVDALCEKALKYLPIDKCSRIIPARARVECFDKFKVNSTLEQYVFCTVNAL 5659
TR|A0A0U1WHG0|A0A0U1WHG0_CVHSA  IVYTACSHAAVDALCEKALKYLPIDKCSRIIPARARVECFAKFKVNSTLEQYVFCTVNAL 5659
TR|A0A166ZL34|A0A166ZL34_9NIDO  IVYTACSHAAVDALCEKALKYLPIDKCSRIIPARARVECFAKFKVNSTLEQYVFCTVNAL 5466
TR|R9QTB2|R9QTB2_CVHSA          ------------------------------------------------------------
TR|R9QTH2|R9QTH2_CVHSA          ------------------------------------------------------------
SP|P0C6U8|R1A_CVHSA             ------------------------------------------------------------
TR|Q6JH47|Q6JH47_CVHSA          ------------------------------------------------------------
TR|Q692E5|Q692E5_CVHSA          ------------------------------------------------------------
SP|P0C6F8|R1A_BCHK3             ------------------------------------------------------------
TR|A0A0K1Z0N1|A0A0K1Z0N1_CVHSA  ------------------------------------------------------------
SP|P0C6F5|R1A_BC279             ------------------------------------------------------------
SP|P0C6T7|R1A_BCRP3             ------------------------------------------------------------

QHN73794                        PETTADIVVFDEISMATNYDLSVVNARLRAKHYVYIGDPAQLPAPRTLLTKGTLEPEYFN 5747
SP|P0C6X7|R1AB_CVHSA            PETTADIVVFDEISMATNYDLSVVNARLRAKHYVYIGDPAQLPAPRTLLTKGTLEPEYFN 5724
TR|Q6UZF5|Q6UZF5_CVHSA          PETTADIVVFDEISMATNYDLSVVNARLRAKHYVYIGDPAQLPAPRTLLTKGTLEPEYFN 5724
TR|Q6UZF1|Q6UZF1_CVHSA          PETTADIVVFDEISMATNYDLSVVNARLRAKHYVYIGDPAQLPAPRTLLTKGTLEPEYFN 5724
TR|Q6JH48|Q6JH48_CVHSA          PETTADIVVFDEISMATNYDLSVVNARLRAKHYVYIGDPAQLPAPRTLLTKGTLEPEYFN 5724
TR|Q692E6|Q692E6_CVHSA          PETTADIVVFDEISMATNYDLSVVNARLRAKHYVYIGDPAQLPAPRTLLTKGTLEPEYFN 5724
TR|A0A0K1YZY7|A0A0K1YZY7_CVHSA  PETTADIVVFDEISMATNYDLSVVNARLRAKHYVYIGDPAQLPAPRTLLTKGTLEPEYFN 5724
SP|P0C6W2|R1AB_BCHK3            PETTADIVVFDEISMATNYDLSVVNARLRAKHYVYIGDPAQLPAPRTLLTKGTLEPEYFN 5718
SP|P0C6W6|R1AB_BCRP3            PETTADIVVFDEISMATNYDLSVVNARLRAKHYVYIGDPAQLPAPRTLLTKGTLEPEYFN 5722
SP|P0C6V9|R1AB_BC279            PETTADIVVFDEISMATNYDLSVVNARLRAKHYVYIGDPAQLPAPRTLLTKGTLEPEYFN 5730
TR|A0A0U1WHI4|A0A0U1WHI4_CVHSA  PETTADIVVFDEISMATNYDLSVVNARLRAKHYVYIGDPAQLPAPRTLLTKGTLEPEYFN 5719
TR|A0A0U1WHG0|A0A0U1WHG0_CVHSA  PETTADIVVFDEISMATNYDLSVVNARLRAKHYVYIGDPAQLPAPRTLLTKGTLEPEYFN 5719
TR|A0A166ZL34|A0A166ZL34_9NIDO  PETTADIVVFDEISMATNYDLSVVNARLRAKHYVYIGDPAQLPAPRTLLTKGTLEPEYFN 5526
TR|R9QTB2|R9QTB2_CVHSA          ------------------------------------------------------------
TR|R9QTH2|R9QTH2_CVHSA          ------------------------------------------------------------
SP|P0C6U8|R1A_CVHSA             ------------------------------------------------------------
TR|Q6JH47|Q6JH47_CVHSA          ------------------------------------------------------------
TR|Q692E5|Q692E5_CVHSA          ------------------------------------------------------------
SP|P0C6F8|R1A_BCHK3             ------------------------------------------------------------
TR|A0A0K1Z0N1|A0A0K1Z0N1_CVHSA  ------------------------------------------------------------
SP|P0C6F5|R1A_BC279             ------------------------------------------------------------
SP|P0C6T7|R1A_BCRP3             ------------------------------------------------------------

QHN73794                        SVCRLMKTIGPDMFLGTCRRCPAEIVDTVSALVYDNKLKAHKDKSAQCFKMFYKGVITHD 5807
SP|P0C6X7|R1AB_CVHSA            SVCRLMKTIGPDMFLGTCRRCPAEIVDTVSALVYDNKLKAHKDKSAQCFKMFYKGVITHD 5784
TR|Q6UZF5|Q6UZF5_CVHSA          SVCRLMKTIGPDMFLGTCRRCPAEIVDTVSALVYDNKLKAHKDKSAQCFKMFYKGVITHD 5784
TR|Q6UZF1|Q6UZF1_CVHSA          SVCRLMKTIGPDMFLGTCRRCPAEIVDTVSALVYDNKLKAHKDKSAQCFKMFYKGVITHD 5784
TR|Q6JH48|Q6JH48_CVHSA          SVCRLMKTIGPDMFLGTCRRCPAEIVDTVSALVYDNKLKAHKDKSAQCFKMFYKGVITHD 5784
TR|Q692E6|Q692E6_CVHSA          SVCRLMKTIGPDMFLGTCRRCPAEIVDTVSALVYDNKLKAHKEKSAQCFKMFYKGVITHD 5784
TR|A0A0K1YZY7|A0A0K1YZY7_CVHSA  SVCRLMKTIGPDMFLGTCRRCPAEIVDTVSALVYDNKLKAHKEKSAQCFKMFYKGVITHD 5784
SP|P0C6W2|R1AB_BCHK3            SVCRLMKTIGPDMFLGTCRRCPAEIVDTVSALVYDNKLKAHKEKSAQCFKMYYKGVITHD 5778
SP|P0C6W6|R1AB_BCRP3            SVCRLMKTIGPDMFLGTCRRCPAEIVDTVSALVYDNKLKAHKEKSAQCFKMFYKGVITHD 5782
SP|P0C6V9|R1AB_BC279            SVCRLMKTIGPDMFLGTCRRCPAEIVDTVSALVYDNKLKAHKEKSAQCFKMFYKGVITHD 5790
TR|A0A0U1WHI4|A0A0U1WHI4_CVHSA  SVCRLMKTIGPDMFLGTCRRCPAEIVDTVSALVYDNKLKAHKEKSAQCFKMFYKGVITHD 5779
TR|A0A0U1WHG0|A0A0U1WHG0_CVHSA  SVCRLMKTIGPDMFLGTCRRCPAEIVDTVSALVYDNKLKAHKEKSAQCFKMFYKGVITHD 5779
TR|A0A166ZL34|A0A166ZL34_9NIDO  SVCRLMKTIGPDMFLGTCRRCPAEIVDTVSALVYDNKLKAHKEKSAQCFKMFYKGVITHD 5586
TR|R9QTB2|R9QTB2_CVHSA          ------------------------------------------------------------
TR|R9QTH2|R9QTH2_CVHSA          ------------------------------------------------------------
SP|P0C6U8|R1A_CVHSA             ------------------------------------------------------------
TR|Q6JH47|Q6JH47_CVHSA          ------------------------------------------------------------
TR|Q692E5|Q692E5_CVHSA          ------------------------------------------------------------
SP|P0C6F8|R1A_BCHK3             ------------------------------------------------------------
TR|A0A0K1Z0N1|A0A0K1Z0N1_CVHSA  ------------------------------------------------------------
SP|P0C6F5|R1A_BC279             ------------------------------------------------------------
SP|P0C6T7|R1A_BCRP3             ------------------------------------------------------------
```

```
QHN73794                          VSSAINRPQIGVVREFLTRNPAWRKAVFISPYNSQNAVASKILGLPTQTVDSSQGSEYDY 5867
SP|P0C6X7|R1AB_CVHSA              VSSAINRPQIGVVREFLTRNPAWRKAVFISPYNSQNAVASKILGLPTQTVDSSQGSEYDY 5844
TR|Q6UZF5|Q6UZF5_CVHSA            VSSAINRPQIGVVREFLTRNPAWRKAVFISPYNSQNAVASKILGLPTQTVDSSQGSEYDY 5844
TR|Q6UZF1|Q6UZF1_CVHSA            VSSAINRPQIGVVREFLTRNPAWRKAVFISPYNSQNAVASKILGLPTQTVDSSQGSEYDY 5844
TR|Q6JH48|Q6JH48_CVHSA            VSSAINRPQIGVVREFLTRNPAWRKAVFISPYNSQNAVASKILGLPTQTVDSSQGSEYDY 5844
TR|Q692E6|Q692E6_CVHSA            VSSAINRPQIGVVREFLTRNPAWRKAVFISPYNSQNAVASKILGLPTQTVDSSQGSEYDY 5844
TR|A0A0K1YZY7|A0A0K1YZY7_CVHSA    VSSAINRPQIGVVREFLTRNPAWRKAVFISPYNSQNAVASKILGLPTQTVDSSQGSEYDY 5844
SP|P0C6W2|R1AB_BCHK3              VSSAINRPQIGVVREFLTRNPAWRKAVFISPYNSQNAVASKILGLPTQTVDSSQGSEYDY 5838
SP|P0C6W6|R1AB_BCRP3              VSSAINRPQIGVVREFLTRNPAWRKAVFISPYNSQNAVASKILGLPTQTVDSSQGSEYDY 5842
SP|P0C6V9|R1AB_BC279              VSSAINRPQIGVVREFLTRNPAWRKAVFISPYNSQNAVASKILGLPTQTVDSSQGSEYDY 5850
TR|A0A0U1WHI4|A0A0U1WHI4_CVHSA    VSSAINRPQIGVVREFLTRNPAWRKAVFISPYNSQNAVASKILGLPTQTVDSSQGSEYDY 5839
TR|A0A0U1WHG0|A0A0U1WHG0_CVHSA    VSSAINRPQIGVVREFLTRNPVWRKAVFISPYNSQNAVASKILGLPTQTVDSSQGSEYDY 5839
TR|A0A166ZL34|A0A166ZL34_9NIDO    VSSAINRPQIGVVREFLTRNPVWRKAVFISPYNSQNAVASKILGLPTQTVDSSQGSEYDY 5646
TR|R9QTB2|R9QTB2_CVHSA            ------------------------------------------------------------
TR|R9QTH2|R9QTH2_CVHSA            ------------------------------------------------------------
SP|P0C6U8|R1A_CVHSA               ------------------------------------------------------------
TR|Q6JH47|Q6JH47_CVHSA            ------------------------------------------------------------
TR|Q692E5|Q692E5_CVHSA            ------------------------------------------------------------
SP|P0C6F8|R1A_BCHK3               ------------------------------------------------------------
TR|A0A0K1Z0N1|A0A0K1Z0N1_CVHSA    ------------------------------------------------------------
SP|P0C6F5|R1A_BC279               ------------------------------------------------------------
SP|P0C6T7|R1A_BCRP3               ------------------------------------------------------------

QHN73794                          VIFTQTTETAHSCNVNRFNVAITRAKVGILCIMSDRDLYDKLQFTSLEIPRRNVATLQAE 5927
SP|P0C6X7|R1AB_CVHSA              VIFTQTTETAHSCNVNRFNVAITRAKIGILCIMSDRDLYDKLQFTSLEIPRRNVATLQAE 5904
TR|Q6UZF5|Q6UZF5_CVHSA            VIFTQTTETAHSCNVNRFNVAITRAKIGILCIMSDRDLYDKLQFTSLEIPRRNVATLQAE 5904
TR|Q6UZF1|Q6UZF1_CVHSA            VIFTQTTETAHSCNVNRFNVAITRAKIGILCIMSDRDLYDKLQFTSLEIPRRNVATLQAE 5904
TR|Q6JH48|Q6JH48_CVHSA            VIFTQTTETAHSCNVNRFNVAITRAKIGILCIMSDRDLYDKLQFTSLEIPRRNVATLQAE 5904
TR|Q692E6|Q692E6_CVHSA            VIFTQTTETAHSCNVNRFNVAITRAKIGILCIMSDRDLYDKLQFTSLEIPRRNVATLQAE 5904
TR|A0A0K1YZY7|A0A0K1YZY7_CVHSA    VIFTQTTETAHSCNVNRFNVAITRAKIGILCIMSDRDLYDKLQFTSLEVPRRNVATLQAE 5904
SP|P0C6W2|R1AB_BCHK3              VIFTQTTETAHSCNVNRFNVAITRAKIGILCIMSDRDLYDKLQFTSLEVPRRNVATLQAE 5898
SP|P0C6W6|R1AB_BCRP3              VIFTQTTETAHSCNVNRFNVAITRAKIGILCIMSDRDLYDKLQFTSLEVPRRNVATLQAE 5902
SP|P0C6V9|R1AB_BC279              VIFTQTTETAHSCNVNRFNVAITRAKIGILCIMSDRDLYDKLQFTSLEVPRRNVATLQAE 5910
TR|A0A0U1WHI4|A0A0U1WHI4_CVHSA    VIFTQTTETAHSCNVNRFNVAITRAKIGILCIMSDRDLYDKLQFTSLEVPRRNVATLQAE 5899
TR|A0A0U1WHG0|A0A0U1WHG0_CVHSA    VIFTQTTETAHSCNVNRFNVAITRAKIGILCIMSDRDLYDKLQFTSLEVPRRNVATLQAE 5899
TR|A0A166ZL34|A0A166ZL34_9NIDO    VIFTQTTETAHSCNVNRFNVAITRAKIGILCIMSDRDLYDKLQFTSLEVPRRNVATLQAE 5706
TR|R9QTB2|R9QTB2_CVHSA            ------------------------------------------------------------
TR|R9QTH2|R9QTH2_CVHSA            ------------------------------------------------------------
SP|P0C6U8|R1A_CVHSA               ------------------------------------------------------------
TR|Q6JH47|Q6JH47_CVHSA            ------------------------------------------------------------
TR|Q692E5|Q692E5_CVHSA            ------------------------------------------------------------
SP|P0C6F8|R1A_BCHK3               ------------------------------------------------------------
TR|A0A0K1Z0N1|A0A0K1Z0N1_CVHSA    ------------------------------------------------------------
SP|P0C6F5|R1A_BC279               ------------------------------------------------------------
SP|P0C6T7|R1A_BCRP3               ------------------------------------------------------------

QHN73794                          NVTGLFKDCSKVITGLHPTQAPTHLSVDTKFKTEGLCVDIPGIPKDMTYRRLISMMGFKM 5987
SP|P0C6X7|R1AB_CVHSA              NVTGLFKDCSKIITGLHPTQAPTHLSVDIKFKTEGLCVDIPGIPKDMTYRRLISMMGFKM 5964
TR|Q6UZF5|Q6UZF5_CVHSA            NVTGLFKDCSKIITGLHPTQAPTHLSVDIKFKTEGLCVDIPGIPKDMTYRRLISMMGFKM 5964
TR|Q6UZF1|Q6UZF1_CVHSA            NVTGLFKDCSKIITGLHPTQAPTHLSVDIKFKTEGLCVDIPGIPKDMTYRRLISMMGFKM 5964
TR|Q6JH48|Q6JH48_CVHSA            NVTGLFKDCSKIITGLHPTQAPTHLSVDIKFKTEGLCVDIPGIPKDMTYRRLISMMGFKM 5964
TR|Q692E6|Q692E6_CVHSA            NVTGLFKDCSKIITGLHPTQAPTHLSVDIKFKTEGLCVDIPGIPKDMTYRRLISMMGFKM 5964
TR|A0A0K1YZY7|A0A0K1YZY7_CVHSA    NVTGLFKDCSKIITGLHPTQAPTHLSVDTKFKTEGLCVDIPGIPKDMTYRRLISMMGFKM 5964
SP|P0C6W2|R1AB_BCHK3              NVTGLFKDCSKIITGLHPTQAPTHLSVDTKFKTEGLCVDIPGIPKDMTYRRLISMMGFKM 5958
SP|P0C6W6|R1AB_BCRP3              NVTGLFKDCSKIITGLHPTQAPTHLSVDTKFKTEGLCVDIPGIPKDMTYRRLISMMGFKM 5962
SP|P0C6V9|R1AB_BC279              NVTGLFKDCSKIITGLHPTQAPTHLSVDTKFKTEGLCVDIPGIPKDMTYRRLISMMGFKM 5970
TR|A0A0U1WHI4|A0A0U1WHI4_CVHSA    NVTGLFKDCSKIITGLHPTQAPTHLSVDTKFKTEGLCVDIPGIPKDMTYRRLISMMGFKM 5959
TR|A0A0U1WHG0|A0A0U1WHG0_CVHSA    NVTGLFKDCSKIITGLHPTQAPTHLSVDTKFKTEGLCVDIPGIPKDMTYRRLISMMGFKM 5959
TR|A0A166ZL34|A0A166ZL34_9NIDO    NVTGLFKDCSKIITGLHPTQAPTHLSVDTKFKTEGLCVDIPGIPKDMTYRRLISMMGFKM 5766
TR|R9QTB2|R9QTB2_CVHSA            ------------------------------------------------------------
TR|R9QTH2|R9QTH2_CVHSA            ------------------------------------------------------------
SP|P0C6U8|R1A_CVHSA               ------------------------------------------------------------
TR|Q6JH47|Q6JH47_CVHSA            ------------------------------------------------------------
TR|Q692E5|Q692E5_CVHSA            ------------------------------------------------------------
SP|P0C6F8|R1A_BCHK3               ------------------------------------------------------------
TR|A0A0K1Z0N1|A0A0K1Z0N1_CVHSA    ------------------------------------------------------------
SP|P0C6F5|R1A_BC279               ------------------------------------------------------------
SP|P0C6T7|R1A_BCRP3               ------------------------------------------------------------
```

```
QHN73794                       KHAFHTPAFDKSAFVNLKQLPFFYYSDSPCESHGKQVVSDIDYVPLKSATCITRCNLGGA 6407
SP|P0C6X7|R1AB_CVHSA           KHAFHTPAFDKSAFTNLKQLPFFYYSDSPCESHGKQVVSDIDYVPLKSATCITRCNLGGA 6384
TR|Q6UZF5|Q6UZF5_CVHSA         KHAFHTPAFDKSAFTNLKQLPFFYYSDSPCESHGKQVVSDIDYVPLKSATCITRCNLGGA 6384
TR|Q6UZF1|Q6UZF1_CVHSA         KHAFHTPAFDKSAFTNLKQLPFFYYSDSPCESHGKQVVSDIDYVPLKSATCITRCNLGGA 6384
TR|Q6JH48|Q6JH48_CVHSA         KHAFHTPAFDKSAFTNLKQLPFFYYSDSPCESHGKQVVSDIDYVPLKSATCITRCNLGGA 6384
TR|Q692E6|Q692E6_CVHSA         KHAFHTPAFDKSAFTNLKQLPFFYYSDSPCESHGKQVVSDIDYVPLKSATCITRCNLGGA 6384
TR|A0A0K1YZY7|A0A0K1YZY7_CVHSA KHAFHTPAFDKSAFTNLKQLPFFYYSDSPCESHGKQVVSDIDYVPLKSATCITRCNLGGA 6384
SP|P0C6W2|R1AB_BCHK3           KHAFHTPAFDKSAFTHLKQLPFFYYSDSPCESHGKQVVSDIDYVPLKSATCITRCNLGGA 6378
SP|P0C6W6|R1AB_BCRP3           KHAFHTPAFDKSAFTNLKQLPFFYYSDSPCESHGKQVVSDIDYVPLKSATCITRCNLGGA 6382
SP|P0C6V9|R1AB_BC279           KHAFHTPAFDKSAFTYLKQLPFFYYSDSPCESHGKQVVSDIDYVPLKSATCITRCNLGGA 6390
TR|A0A0U1WHI4|A0A0U1WHI4_CVHSA KHAFHTPAFDKSAFSNLKQLPFFYYSDSPCESHGKQVVSDIDYVPLKSATCITRCNLGGA 6379
TR|A0A0U1WHG0|A0A0U1WHG0_CVHSA KHAFHTPAFDKSAFTHLKQLPFFYYSDSPCESHGKQVVSDIDYVPLKSATCITRCNLGGA 6379
TR|A0A166ZL34|A0A166ZL34_9NIDO KHAFYTPAFDKSAFTHLKQLPFFYYSDSPCESHGKQVVSDIDYVPLKSATCITRCNLGGA 6186
TR|R9QTB2|R9QTB2_CVHSA         ------------------------------------------------------------
TR|R9QTH2|R9QTH2_CVHSA         ------------------------------------------------------------
SP|P0C6U8|R1A_CVHSA            ------------------------------------------------------------
TR|Q6JH47|Q6JH47_CVHSA         ------------------------------------------------------------
TR|Q692E5|Q692E5_CVHSA         ------------------------------------------------------------
SP|P0C6F8|R1A_BCHK3            ------------------------------------------------------------
TR|A0A0K1Z0N1|A0A0K1Z0N1_CVHSA ------------------------------------------------------------
SP|P0C6F5|R1A_BC279            ------------------------------------------------------------
SP|P0C6T7|R1A_BCRP3            ------------------------------------------------------------

QHN73794                       VCRHHANEYRLYLDAYNMMISAGFSLWVYKQFDTYNLWNTFTRLQSLENVAFNVVNKGHF 6467
SP|P0C6X7|R1AB_CVHSA           VCRHHANEYRQYLDAYNMMISAGFSLWIYKQFDTYNLWNTFTRLQSLENVAYNVVNKGHF 6444
TR|Q6UZF5|Q6UZF5_CVHSA         VCRHHANEYRQYLDAYNMMISAGFSLWIYKQFDTYNLWNTFTRLQSLENVAYNVVNKGHF 6444
TR|Q6UZF1|Q6UZF1_CVHSA         VCRHHANEYRQYLDAYNMMISAGFSLWIYKQFDTYNLWNTFTRLQSLENVAYNVVNKGHF 6444
TR|Q6JH48|Q6JH48_CVHSA         VCRHHANEYRQYLDAYNMMISAGFSLWIYKQFDTYNLWNTFTRLQSLENVAYNVVNKGHF 6444
TR|Q692E6|Q692E6_CVHSA         VCRHHANEYRQYLDAYNMMISAGFSLWIYKQFDTYNLWNTFTRLQSLENVAYNVVNKGHF 6444
TR|A0A0K1YZY7|A0A0K1YZY7_CVHSA VCRHHANEYRQYLDAYNMMISAGFSLWIYKQFDTYNLWNTFTRLQSLENVAYNVVNKGHF 6444
SP|P0C6W2|R1AB_BCHK3           VCRHHANEYRQYLDAYNMMISAGFSLWIYKQFDTYNLWNTFTKLQSLENVAYNVVNKGHF 6438
SP|P0C6W6|R1AB_BCRP3           VCRHHANEYRQYLDAYNMMISAGFSLWIYKQFDTYNLWNTFTRLQSLENVAYNVVNKGHF 6442
SP|P0C6V9|R1AB_BC279           VCRRHANEYRQYLDAYNMMISAGFSLWIYKQFDTYNLWNTFTRLQSLENVAYNVVNKGHF 6450
TR|A0A0U1WHI4|A0A0U1WHI4_CVHSA VCRHHANEYRQYLDAYNMMISAGFSLWIYKQFDTYNLWNTFTRLQSLENVAYNVINKGHF 6439
TR|A0A0U1WHG0|A0A0U1WHG0_CVHSA VCRHHANEYRQYLDAYNMMISAGFSLWIYKQFDTYNLWNTFTRLQSLENVAYNVINKGHF 6439
TR|A0A166ZL34|A0A166ZL34_9NIDO VCRHHANEYRQYLDAYNMMISAGFSLWIYKQFDTYNLWNTFTRLQSLENVAYNVINKGHF 6246
TR|R9QTB2|R9QTB2_CVHSA         ------------------------------------------------------------
TR|R9QTH2|R9QTH2_CVHSA         ------------------------------------------------------------
SP|P0C6U8|R1A_CVHSA            ------------------------------------------------------------
TR|Q6JH47|Q6JH47_CVHSA         ------------------------------------------------------------
TR|Q692E5|Q692E5_CVHSA         ------------------------------------------------------------
SP|P0C6F8|R1A_BCHK3            ------------------------------------------------------------
TR|A0A0K1Z0N1|A0A0K1Z0N1_CVHSA ------------------------------------------------------------
SP|P0C6F5|R1A_BC279            ------------------------------------------------------------
SP|P0C6T7|R1A_BCRP3            ------------------------------------------------------------

QHN73794                       DGQQGEVPVSIINNTVYTKVDGVDVELFENKTTLPVNVAFELWAKRNIKPVPEVKILNNL 6527
SP|P0C6X7|R1AB_CVHSA           DGHAGEAPVSIINNAVYTKVDGIDVEIFENKTTLPVNVAFELWAKRNIKPVPEIKILNNL 6504
TR|Q6UZF5|Q6UZF5_CVHSA         DGHAGEAPVSIINNAVYTKVDGIDVEIFENKTTLPVNVAFELWAKRNIKPVPEIKILNNL 6504
TR|Q6UZF1|Q6UZF1_CVHSA         DGHAGEAPVSIINNAVYTKVDGIDVEIFENKTTLPVNVAFELWAKRNIKPVPEIKILNNL 6504
TR|Q6JH48|Q6JH48_CVHSA         DGHAGEAPVSIINNAVYTKVDGIDVEIFENKTTLPVNVAFELWAKRNIKPVPEIKILNNL 6504
TR|Q692E6|Q692E6_CVHSA         DGHAGEAPVSIINNAVYTKVDGIDVEIFENKTTLPVNVAFELWAKRNIKPVPEIKILNNL 6504
TR|A0A0K1YZY7|A0A0K1YZY7_CVHSA DGQAGEAPVSIINNAVYTKVDGVDVEIFENKTTLPVNVAFELWAKRNIKPVPEIKILNNL 6504
SP|P0C6W2|R1AB_BCHK3           DGQSGEAPVSIINNAVYTKVDGIDVEIFENKTTLPVNVAFELWAKRNIKPVPEIKILNNL 6498
SP|P0C6W6|R1AB_BCRP3           DGQAGETPVSIINNAVYTKVDGFDVEIFENKTTLPVNVAFELWAKRNIKSVPEIKILNNL 6502
SP|P0C6V9|R1AB_BC279           DGQIGEAPVSIINNAVYTKVDGNDVEIFENKTTLPVNVAFELWAKRNIKPVPEIKILNNL 6510
TR|A0A0U1WHI4|A0A0U1WHI4_CVHSA DGQIGEVPVSIINNAVYTKVDGVDVEIFENKTILPVNVAFELWAKRNIKPVPEIKILNNL 6499
TR|A0A0U1WHG0|A0A0U1WHG0_CVHSA DGQIGEAPVSIINNAVYTKVDGVDVEIFENKTTLPVNVAFELWAKRNIKPVPEIKILNNL 6499
TR|A0A166ZL34|A0A166ZL34_9NIDO DGQIGEAPVSIINNAVYTKVDGVDVEIFENKTTLPVNVAFELWAKRNIKPVPEIKILNNL 6306
TR|R9QTB2|R9QTB2_CVHSA         ------------------------------------------------------------
TR|R9QTH2|R9QTH2_CVHSA         ------------------------------------------------------------
SP|P0C6U8|R1A_CVHSA            ------------------------------------------------------------
TR|Q6JH47|Q6JH47_CVHSA         ------------------------------------------------------------
TR|Q692E5|Q692E5_CVHSA         ------------------------------------------------------------
SP|P0C6F8|R1A_BCHK3            ------------------------------------------------------------
TR|A0A0K1Z0N1|A0A0K1Z0N1_CVHSA ------------------------------------------------------------
SP|P0C6F5|R1A_BC279            ------------------------------------------------------------
SP|P0C6T7|R1A_BCRP3            ------------------------------------------------------------
```

```
QHN73794                        GVDIAANTVIWDYKRDAPAHISTIGVCSMTDIAKKPTETICAPLTVFFDGRVDGQVDLFR 6587
SP|P0C6X7|R1AB_CVHSA            GVDIAANTVIWDYKREAPAHVSTIGVCTMTDIAKKPTESACSSLTVLFDGRVEGQVDLFR 6564
TR|Q6UZF5|Q6UZF5_CVHSA          GVDIAANTVIWDYKREAPAHVSTIGVCTMTDIAKKPTESACSSLTVLFDGRVEGQVDLFR 6564
TR|Q6UZF1|Q6UZF1_CVHSA          GVDIAANTVIWDYKREAPAHVSTIGVCTMTDIAKKPTESACSSLTVLFDGRVEGQVDLFR 6564
TR|Q6JH48|Q6JH48_CVHSA          GVDIAANTVIWDYKREAPAHVSTIGVCTMTDIAKKPTESACSSLTVLFDGRVEGQVDLFR 6564
TR|Q692E6|Q692E6_CVHSA          GVDIAANTVIWDYKREAPAHVSTIGVCTMTDIAKKPTESACSSLTVLFDGRVEGQVDLFR 6564
TR|A0A0K1YZY7|A0A0K1YZY7_CVHSA GVDIAANTVIWDYKREAPAHMSTIGVSTMTDIAKKPTESACSSLTVLFDGRVEGQVDLFR 6564
SP|P0C6W2|R1AB_BCHK3            GVDIAANNVIWDYKREAPAHVSTIGVCTMTDIAKKPTESACSSLIVLFDGRVEGQVDFFR 6558
SP|P0C6W6|R1AB_BCRP3            GVDIAANTVIWDHKREAPVHMSTIGVCTMTDIAKKPTESACSSLTVLFDGRVEGQVDLFR 6562
SP|P0C6V9|R1AB_BC279            GVDIAANTVIWDYKREAPAHVSTIGVCTMTDIAKKPTESACSSLTVLFDGRVEGQVDLFR 6570
TR|A0A0U1WHI4|A0A0U1WHI4_CVHSA GVDIAANTVIWDYKREAPAHVSTIGICTMTDIAKKPTESACSSLTVLFDGRVEGQVDLFR 6559
TR|A0A0U1WHG0|A0A0U1WHG0_CVHSA GVDIAANTVIWDYKREAPAHVSTIGICTMTDIAKKPTESTCSSLTVLFDGRVEGQVDLFR 6559
TR|A0A166ZL34|A0A166ZL34_9NIDO GVDIAANTVIWDYKREAPAHVSTIGICTMTDIAKKPTESTCSSLTVLFDGRVEGQVDLFR 6366
TR|R9QTB2|R9QTB2_CVHSA          ------------------------------------------------------------
TR|R9QTH2|R9QTH2_CVHSA          ------------------------------------------------------------
SP|P0C6U8|R1A_CVHSA             ------------------------------------------------------------
TR|Q6JH47|Q6JH47_CVHSA          ------------------------------------------------------------
TR|Q692E5|Q692E5_CVHSA          ------------------------------------------------------------
SP|P0C6F8|R1A_BCHK3             ------------------------------------------------------------
TR|A0A0K1Z0N1|A0A0K1Z0N1_CVHSA ------------------------------------------------------------
SP|P0C6F5|R1A_BC279             ------------------------------------------------------------
SP|P0C6T7|R1A_BCRP3             ------------------------------------------------------------

QHN73794                        NARNGVLITEGSVKGLQPSVGPKQASLNGVTLIGEAVKTQFNYYKKVDGVVQQLPETYFT 6647
SP|P0C6X7|R1AB_CVHSA            NARNGVLITEGSVKGLTPSKGPAQASVNGVTLIGESVKTQFNYFKKVDGIIQQLPETYFT 6624
TR|Q6UZF5|Q6UZF5_CVHSA          NARNGVLITEGSVKGLTPSKGPAQASVNGVTLIGESVKTQFNYFKKVDGIIQQLPETYFT 6624
TR|Q6UZF1|Q6UZF1_CVHSA          NARNGVLITEGSVKGLTPSKGPAQASVNGVTLIGESVKTQFNYFKKVDGIIQQLPETYFT 6624
TR|Q6JH48|Q6JH48_CVHSA          NARNGVLITEGSVKGLTPSKGPAQASVNGVTLIGESVKTQFNYFKKVDGIIQQLPETYFT 6624
TR|Q692E6|Q692E6_CVHSA          NARNGVLITEGSVKGLTPSKGPAQASVNGVTLIGESVKTQFNYFKKVDGIIQQLPETYFT 6624
TR|A0A0K1YZY7|A0A0K1YZY7_CVHSA NARNGVLITEGSVKGLTPSKGPAQASVNGVTLIGESVKTQFNYFKKVDGIIQQLPETYFT 6624
SP|P0C6W2|R1AB_BCHK3            NARNGVLITEGSVKGLTPSKGPAQASVNGVTLIGESVKTQFNYFKKVDGIIQQLPETYFT 6618
SP|P0C6W6|R1AB_BCRP3            NARNGVLITEGSVKGLTPSKGPAQASVNGVTLIGESVKTQFNYFKKVDGIIQQLPETYFT 6622
SP|P0C6V9|R1AB_BC279            NARNGVLITEGSVKGLTPSKGPAQASVNGVTLIGESVKTQFNYFKKVDGIIQQLPETYFT 6630
TR|A0A0U1WHI4|A0A0U1WHI4_CVHSA NARNGVLITEGSVKGLTPSKGPVQASVNGVTLIGESVKTQFNYFKKVDGIIQQLPETYFT 6619
TR|A0A0U1WHG0|A0A0U1WHG0_CVHSA NARNGVLITEGSVKGLTPSKGPVQASVNGVTLIGESVKTQFNYFKKVDGIIQQLPETYFT 6619
TR|A0A166ZL34|A0A166ZL34_9NIDO NARNGVLITEGSVKGLTPSKGPVQASVNGVTLIGESVKTQFNYFKKVDGIIQQLPETYFT 6426
TR|R9QTB2|R9QTB2_CVHSA          ------------------------------------------------------------
TR|R9QTH2|R9QTH2_CVHSA          ------------------------------------------------------------
SP|P0C6U8|R1A_CVHSA             ------------------------------------------------------------
TR|Q6JH47|Q6JH47_CVHSA          ------------------------------------------------------------
TR|Q692E5|Q692E5_CVHSA          ------------------------------------------------------------
SP|P0C6F8|R1A_BCHK3             ------------------------------------------------------------
TR|A0A0K1Z0N1|A0A0K1Z0N1_CVHSA ------------------------------------------------------------
SP|P0C6F5|R1A_BC279             ------------------------------------------------------------
SP|P0C6T7|R1A_BCRP3             ------------------------------------------------------------

QHN73794                        QSRNLQEFKPRSQMEIDFLELAMDEFIERYKLEGYAFEHIVYGDFSHSQLGGLHLLIGLA 6707
SP|P0C6X7|R1AB_CVHSA            QSRDLEDFKPRSQMETDFLELAMDEFIQRYKLEGYAFEHIVYGDFSHGQLGGLHLMIGLA 6684
TR|Q6UZF5|Q6UZF5_CVHSA          QSRDLEDFKPRSQMETDFLELAMDEFIQRYKLEGYAFEHIVYGDFSHGQLGGLHLMIGLA 6684
TR|Q6UZF1|Q6UZF1_CVHSA          QSRDLEDFKPRSQMETDFLELAMDEFIQRYKLEGYAFEHIVYGDFSHGQLGGLHLMIGLA 6684
TR|Q6JH48|Q6JH48_CVHSA          QSRDLEDFKPRSQMETDFLELAMDEFIQRYKLEGYAFEHIVYGDFSHGQLGGLHLMIGLA 6684
TR|Q692E6|Q692E6_CVHSA          QSRDLEDFKPRSQMETDFLELAMDEFIQRYKLEGYAFEHIVYGDFSHGQLGGLHLMIGLA 6684
TR|A0A0K1YZY7|A0A0K1YZY7_CVHSA QSRDLEDFKPRSQMETDFLELVMDEFIQRYKLEGYAFEHIVYGDFSHGQLGGLHLMIGLA 6684
SP|P0C6W2|R1AB_BCHK3            QSRDLEDFKPRSQMETDFLELAMDEFIQRYKLEGYAFEHIVYGDFSHGQLGGLHLMIGLA 6678
SP|P0C6W6|R1AB_BCRP3            QSRDLEDFKPRSQMETDFLELAMDEFIQRYKLEGYAFEHIVYGDFSHGQLGGLHLMIGLA 6682
SP|P0C6V9|R1AB_BC279            QSRDLEDFKPRSKMETDFLELAMDEFIQRYKLEGYAFEHIVYGDFSHGQLGGLHLMIGLA 6690
TR|A0A0U1WHI4|A0A0U1WHI4_CVHSA QSRDLEDFKPRSQMETDFLELAMDEFIQRYKLEGYAFEHIVYGDFSHGQLGGLHLMIGLA 6679
TR|A0A0U1WHG0|A0A0U1WHG0_CVHSA QSRDLEDFKPRSQMETDFLELAMDEFIQRYKLEGYAFDHIVYGDFSHGQLGGLHLMIGLA 6679
TR|A0A166ZL34|A0A166ZL34_9NIDO QSRDLEDFKPRSQMETDFLELAMDEFIQRYKLEGYAFDHIVYGDFSHGQLGGLHLMIGLA 6486
TR|R9QTB2|R9QTB2_CVHSA          ------------------------------------------------------------
TR|R9QTH2|R9QTH2_CVHSA          ------------------------------------------------------------
SP|P0C6U8|R1A_CVHSA             ------------------------------------------------------------
TR|Q6JH47|Q6JH47_CVHSA          ------------------------------------------------------------
TR|Q692E5|Q692E5_CVHSA          ------------------------------------------------------------
SP|P0C6F8|R1A_BCHK3             ------------------------------------------------------------
TR|A0A0K1Z0N1|A0A0K1Z0N1_CVHSA ------------------------------------------------------------
SP|P0C6F5|R1A_BC279             ------------------------------------------------------------
SP|P0C6T7|R1A_BCRP3             ------------------------------------------------------------
```

```
QHN73794                          KRFKESPFELEDFIPMDSTVKNYFITDAQTGSSKCVCSVIDLLLDDFVEIIKSQDLSVVS 6767
SP|P0C6X7|R1AB_CVHSA              KRSQDSPLKLEDFIPMDSTVKNYFITDAQTGSSKCVCSVIDLLLDDFVEIIKSQDLSVIS 6744
TR|Q6UZF5|Q6UZF5_CVHSA            KRSQDSPLKLEDFIPMDSTVKNYFITDAQTGSSKCVCSVIDLLLDDFVEIIKSQDLSVIS 6744
TR|Q6UZF1|Q6UZF1_CVHSA            KRSQDSPLKLEDFIPMDSTVKNYFITDAQTGSSKCVCSVIDLLLDDFVEIIKSQDLSVIS 6744
TR|Q6JH48|Q6JH48_CVHSA            KRSQDSPLKLEDFIPMDSTVKNYFITDAQTGSSKCVCSVIDLLLDDFVEIIKSQDLSVIS 6744
TR|Q692E6|Q692E6_CVHSA            KRSQDSPLKLEDFIPMDSTVKNYFITDAQTGSSKCVCSVIDLLLDDFVEIIKSQDLSVIS 6744
TR|A0A0K1YZY7|A0A0K1YZY7_CVHSA    KRSQDSPLKLEDFIPMDSTVKNYFITDAQTGSSKCVCSVIDLLLDDFVEIIKSQDLSVIS 6744
SP|P0C6W2|R1AB_BCHK3              KRSQDSLLKLEDFIPMDSTVKNYFITDAQTGSSKCVCSVIDLLLDDFVEIIKSQDLSVVS 6738
SP|P0C6W6|R1AB_BCRP3              KRSRDSPLKLEDFIPMDSTVKNYFITDAQTGSSKCVCSVIDLLLDDFVEIIKSQDLSVVS 6742
SP|P0C6V9|R1AB_BC279              KRSQDSPLKLEDFIPTDSTVKNYFITDAQTGSSKCVCSVIDLLLDDFVEIIKSQDLSVIS 6750
TR|A0A0U1WHI4|A0A0U1WHI4_CVHSA    KRSQDSPLKLEDFIPMDSTVKNYFITDAQTGSSKCVCSVIDLLLDDFVEIIKSQDLSVVS 6739
TR|A0A0U1WHG0|A0A0U1WHG0_CVHSA    KRSQDSPLKLEDFIPMDSTVKNYFITDAKTGSSKCVCSVIDLLLDDFVEIIKSQDLSVIS 6739
TR|A0A166ZL34|A0A166ZL34_9NIDO    KRSQDSPLKLEDFIPMDSTVKNYFITDAQTGSSKCVCSVIDLLLDDFVEIIKSQDLSVIS 6546
TR|R9QTB2|R9QTB2_CVHSA            ------------------------------------------------------------
TR|R9QTH2|R9QTH2_CVHSA            ------------------------------------------------------------
SP|P0C6U8|R1A_CVHSA               ------------------------------------------------------------
TR|Q6JH47|Q6JH47_CVHSA            ------------------------------------------------------------
TR|Q692E5|Q692E5_CVHSA            ------------------------------------------------------------
SP|P0C6F8|R1A_BCHK3               ------------------------------------------------------------
TR|A0A0K1Z0N1|A0A0K1Z0N1_CVHSA    ------------------------------------------------------------
SP|P0C6F5|R1A_BC279               ------------------------------------------------------------
SP|P0C6T7|R1A_BCRP3               ------------------------------------------------------------

QHN73794                          KVVKVTIDYTEISFMLWCKDGHVETFYPKLQSSQAWQPGVAMPNLYKMQRMLLEKCDLQN 6827
SP|P0C6X7|R1AB_CVHSA              KVVKVTIDYAEISFMLWCKDGHVETFYPKLQASQAWQPGVAMPNLYKMQRMLLEKCDLQN 6804
TR|Q6UZF5|Q6UZF5_CVHSA            KVVKVTIDYAEISFMLWCKDGHVETFYPKLQASQAWQPGVAMPNLYKMQRMLLEKCDLQN 6804
TR|Q6UZF1|Q6UZF1_CVHSA            KVVKVTIDYAEISFMLWCKDGHVETFYPKLQASQAWQPGVAMPNLYKMQRMLLEKCDLQN 6804
TR|Q6JH48|Q6JH48_CVHSA            KVVKVTIDYAEISFMLWCKDGHVETFYPKLQASQAWQPGVAMPNLYKMQRMLLEKCDLQN 6804
TR|Q692E6|Q692E6_CVHSA            KVVKVTIDYAEISFMLWCKDGHVETFYPKLQASQAWQPGVAMPNLYKMQRMLLEKCDLQN 6804
TR|A0A0K1YZY7|A0A0K1YZY7_CVHSA    KVVKVTIDYVEISFMLWCKDGHVETFYPKLQASQAWQPGVAMPNLYKMQRMLLEKCDLQN 6804
SP|P0C6W2|R1AB_BCHK3              KVVKVTIDYAEISFMLWCKDGHVETFYPKLQASQAWQPGVAMPNLYKMQRMLLEKCDLQN 6798
SP|P0C6W6|R1AB_BCRP3              KVVKVTIDYAEISFMLWCKDGHVETFYPKLQASQAWQPGVAMPNLYKMQRMLLEKCDLQN 6802
SP|P0C6V9|R1AB_BC279              KVVKVTIDYAEISFMLWCKDGHVETFYPKLQASQAWQPGVAMPNLYKMQRMLLEKCDLQN 6810
TR|A0A0U1WHI4|A0A0U1WHI4_CVHSA    KVVKVTIDYAEISFMLWCKDGHVETFYPKLQASQAWQPGVAMPNLYKMQRMLLEKCDLQN 6799
TR|A0A0U1WHG0|A0A0U1WHG0_CVHSA    KVVKVTIDYAEISFMLWCKDGYVETFYPKLQASQAWQPGVAMPNLYKMQRMLLEKCDLQN 6799
TR|A0A166ZL34|A0A166ZL34_9NIDO    KVVKVTIDYAEISFMLWCKDGYVETFYPKLQASQAWQPGVAMPNLYKMQRMLLEKCDLQN 6606
TR|R9QTB2|R9QTB2_CVHSA            ------------------------------------------------------------
TR|R9QTH2|R9QTH2_CVHSA            ------------------------------------------------------------
SP|P0C6U8|R1A_CVHSA               ------------------------------------------------------------
TR|Q6JH47|Q6JH47_CVHSA            ------------------------------------------------------------
TR|Q692E5|Q692E5_CVHSA            ------------------------------------------------------------
SP|P0C6F8|R1A_BCHK3               ------------------------------------------------------------
TR|A0A0K1Z0N1|A0A0K1Z0N1_CVHSA    ------------------------------------------------------------
SP|P0C6F5|R1A_BC279               ------------------------------------------------------------
SP|P0C6T7|R1A_BCRP3               ------------------------------------------------------------

QHN73794                          YGDSATLPKGIMMNVAKYTQLCQYLNTLTLAVPYNMRVIHFGAGSDKGVAPGTAVLRQWL 6887
SP|P0C6X7|R1AB_CVHSA              YGENAVIPKGIMMNVAKYTQLCQYLNTLTLAVPYNMRVIHFGAGSDKGVAPGTAVLRQWL 6864
TR|Q6UZF5|Q6UZF5_CVHSA            YGENAVIPKGIMMNVAKYTQLCQYLNTLTLAVPYNMRVIHFGAGSDKGVAPGTAVLRQWL 6864
TR|Q6UZF1|Q6UZF1_CVHSA            YGENAVIPKGIMMNVAKYTQLCQYLNTLTLAVPYNMRVIHFGAGSDKGVAPGTAVLRQWL 6864
TR|Q6JH48|Q6JH48_CVHSA            YGENAVIPKGIMMNVAKYTQLCQYLNTLTLAVPYNMRVIHFGAGSDKGVAPGTAVLRQWL 6864
TR|Q692E6|Q692E6_CVHSA            YGENAVIPKGIMMNVAKYTQLCQYLNTLTLAVPYNMRVIHFGAGSDKGVAPGTAVLRQWL 6864
TR|A0A0K1YZY7|A0A0K1YZY7_CVHSA    YGENAVIPKGIMMNVAKYTQLCQYLNTLTLAVPYNMRVIHFGAGSDKGVAPGTAVLRQWL 6864
SP|P0C6W2|R1AB_BCHK3              YGENAVIPKGIMMNVAKYTQLCQYLNTLTLAVPYNMRVIHFGAGSDKGVAPGTAVLRQWL 6858
SP|P0C6W6|R1AB_BCRP3              YGENAVIPKGIMMNVAKYTQLCQYLNTLTLAVPYNMRVIHFGAGSDKGVAPGTAVLRQWL 6862
SP|P0C6V9|R1AB_BC279              YGENAVIPKGIMMNVAKYTQLCQYLNTLTLAVPYNMRVIHFGAGSDKGVAPGTAVLRQWL 6870
TR|A0A0U1WHI4|A0A0U1WHI4_CVHSA    YGENAVIPKGIMMNVAKYTQLCQYLNTLTLAVPYNMRVIHFGAGSDKGVAPGTAVLRQWL 6859
TR|A0A0U1WHG0|A0A0U1WHG0_CVHSA    YGENAVIPKGIMMNVAKYTQLCQYLNTLTLAVPYNMRVIHFGAGSDKGVAPGTAVLRQWL 6859
TR|A0A166ZL34|A0A166ZL34_9NIDO    YGENAVIPKGIMMNVAKYTQLCQYLNTLTLAVPYNMRVIHFGAGSDKGVAPGTAVLRQWL 6666
TR|R9QTB2|R9QTB2_CVHSA            ------------------------------------------------------------
TR|R9QTH2|R9QTH2_CVHSA            ------------------------------------------------------------
SP|P0C6U8|R1A_CVHSA               ------------------------------------------------------------
TR|Q6JH47|Q6JH47_CVHSA            ------------------------------------------------------------
TR|Q692E5|Q692E5_CVHSA            ------------------------------------------------------------
SP|P0C6F8|R1A_BCHK3               ------------------------------------------------------------
TR|A0A0K1Z0N1|A0A0K1Z0N1_CVHSA    ------------------------------------------------------------
SP|P0C6F5|R1A_BC279               ------------------------------------------------------------
SP|P0C6T7|R1A_BCRP3               ------------------------------------------------------------
```
