## Supplementary material for "Structural genomics and interactomics of 2019 Wuhan novel coronavirus, 2019-nCoV, indicate evolutionary conserved functional regions of viral proteins": Protein binding site mapping for wS

Homo-oligomer Protein Binding Sites:

wS_homotrimer-Conf1-5X58

Host-Virus Interaction Protein Binding Sites:

wS_heterodimer-IGHV3-30-3-2GHW
wS_trimer-ACE2-Conf1-6ACG
wS_trimer-ACE2-Conf2-6ACJ
wS_trimer-ACE2-Conf3-6ACK
wS_trimer-IGHV3-30-3-Conf1-6NB6
wS_trimer-IGHV3-30-3-Conf2-6NB7

```
YP_009724390                   MFV-------FLVLLPLVSSQCVNLTTRTQLPPAYTNSFTRGVYYPDKVFRSSVLHSTQD 53
SP|P59594|SPIKE_CVHSA          MFI--FLL-FLTLTSGSDLDRCTTFDDVQAPNYTQHTSSMRGVYYPDEIFRSDTLYLTQD 57
TR|Q6UZF4|Q6UZF4_CVHSA         MFI--FLL-FLTLTSGSDLDRCTTFDDVQAPNYTQHTSSMRGVYYPDEIFRSDTLYLTQD 57
TR|Q6UZF0|Q6UZF0_CVHSA         MFI--FLL-FLTLTSGSDLDRCTTFDDVQAPNYTQHTSSMRGVYYPDEIFRSDTLYLTQD 57
TR|Q6JH46|Q6JH46_CVHSA         MFI--FLL-FLTLTSGSDLDRCTTFDDVQAPNYTQHTSSMRGVYYPDEIFRSDTLYLTQD 57
TR|Q692E4|Q692E4_CVHSA         MFI--FLL-FLTLTSGSDLDRCTTFDDVQAPNYTQHTSSMRGVYYPDEIFRSDTLYLTQD 57
SP|Q3LZX1|SPIKE_BCHK3          MKILIF---AFLANLAKAQEGCGIISRKPQPKMAQVSSSRRGVYYNDDIFRSDVLHLTQD 57
SP|Q3I5J5|SPIKE_BCRP3          MKILIL---AFLASLAKAQEGCGIISRKPQPKMAQVSSSRRGVYYNDDIFRSNVLHLTQD 57
SP|Q0Q475|SPIKE_BC279          MKVLIF---ALLFSLAKAQEGCGIISRKPQPKMEKVSSSRRGVYYNDDIFRSDVLHLTQD 57
TR|R9QTA0|R9QTA0_CVHSA         --MILL---LLFLSSAKAQEGCGVISNKPQRTFDQYSSTFRGVYYNDDIFRSDVLHLTQD 55
TR|R9QTH3|R9QTH3_CVHSA         MKIFLF---SLLFSAALAQEGCGLLSFKPQPKLAQFSSSKRGVYYNDDIFRSDVLHLTQD 57
TR|A0A0K1Z074|A0A0K1Z074_CVHSA MKILIF---AFLVTLVKAQEGCGVINLRTQPKLSQVSSSRRGVYYNDDIFRSDVLHLTQD 57
TR|A0A0U1WHI6|A0A0U1WHI6_CVHSA MKILIF---AFLVTLVKAQEGCGVINLKTQPTLTQVSSSRRGVYYNDDIFRSDVLHLTQD 57
TR|A0A166ZL64|A0A166ZL64_9NIDO MFIVFLISYTTFLIPYTATTTC--FSGPTIENKLNISSGSRGVYYPDDIFRSNVSVLVTG 58
TR|A0A0U1UYX4|A0A0U1UYX4_CVHSA MFIVFLISYTTFLIPYTATTTC--FKGPTTENKLNISSGSRGVYYPDDIFRSNVSVLVTG 58
                                 :                  *  :           .*  ***** *.:***..   . .

YP_009724390                   LFLPFFSNVTWFHAIHVSGTNGTKRFDNPVLPFNDGVYFASTEKSNIIRGWIFGTTLDSK 113
SP|P59594|SPIKE_CVHSA          LFLPFYSNVTGFHTINH-------TFGNPVIPFKDGIYFAATEKSNVVRGWVFGSTMNNK 110
TR|Q6UZF4|Q6UZF4_CVHSA         LFLPFYSNVTGFHTINH-------TFGNPVIPFKDGIYFAATEKSNVVRGWVFGSTMNNK 110
TR|Q6UZF0|Q6UZF0_CVHSA         LFLPFYSNVTGFHTINH-------TFGNPVIPFKDGIYFAATEKSNVVRGWVFGSTMNNK 110
TR|Q6JH46|Q6JH46_CVHSA         LFLPFYSNVTGFHTINH-------TFGNPVIPFKDGIYFAATEKSNVVRGWVFGSTMNNK 110
TR|Q692E4|Q692E4_CVHSA         LFLPFYSNVTGFHTINH-------TFGNPVIPFKDGIYFAATEKSNVVRGWVFGSTMNNK 110
SP|Q3LZX1|SPIKE_BCHK3          YFLPFDSNLTQYFSLNVDS-DRYTYFDNPILDFGDGVYFAATEKSNVIRGWIFGSSFDNT 116
SP|Q3I5J5|SPIKE_BCRP3          YFLPFDSNLTQYFSLNVDS-DRFTYFDNPILDFGDGVYFAATEKSNVIRGWIFGSTFDNT 116
SP|Q0Q475|SPIKE_BC279          YFLPFDSNLTQYFSLNIDS-NKYTYFDNPILDFGDGVYFAATEKSNVIRGWIFGSSFDNT 116
TR|R9QTA0|R9QTA0_CVHSA         YFLPFNTNVTRYLSLNAAQ-NTIVYFDNHVIPFYDGIYFAATERSNVIRGWIFGSTFDNR 114
TR|R9QTH3|R9QTH3_CVHSA         YFLPFHSNLTQYFSLNVDS-DRQVYFDNPTLNFGDGVYFAATEKSNVIRGWIFGSTMDNS 116
TR|A0A0K1Z074|A0A0K1Z074_CVHSA YFLPFHSNLTQYFSLSIQS-DKIVYFDNPILKFGDGIYFAATEKSNVIRGWVFGSTFDNT 116
TR|A0A0U1WHI6|A0A0U1WHI6_CVHSA YFLPFHSNLTQYFSLNIES-D--IYFDNPILKFGDGVYFAATEKSNVIRGWVFGSTFDNT 114
TR|A0A166ZL64|A0A166ZL64_9NIDO RFLRFNTTLTWYNSWNQ-------AYSSPVLPFGHGVYFSTIDKSNVVRGWIFGTTLDNT 111
TR|A0A0U1UYX4|A0A0U1UYX4_CVHSA RFLRFNTTLTWYNSWNQ-------AYSSPVLPFGHGVYFSTIDKSNVVRGWIFGTTLDNT 111
                                ** * :.:* : :           :..  : * .*:**:: ::**::***:**::::.

YP_009724390                   TQSLLIVNNATNVVIKVCEFQFCNDPFLGVYYHKNNKSWMESEFRVYSSANNCTFEYVSQ 173
SP|P59594|SPIKE_CVHSA          SQSVIIINNSTNVVIRACNFELCDNPFFAVSKPMG----TQTHTMIFDNAFNCTFEYISD 166
TR|Q6UZF4|Q6UZF4_CVHSA         SQSVIIINNSTNVVIRACNFELCDNPFFAVSKPMG----TQTHTMIFDNAFNCTFEYISD 166
TR|Q6UZF0|Q6UZF0_CVHSA         SQSVIIINNSTNVVIRACNFELCDNPFFAVSKPMG----TQTHTMIFDNAFNCTFEYISD 166
TR|Q6JH46|Q6JH46_CVHSA         SQSVIIINNSTNVVIRACNFELCDNPFFAVSKPMG----TQTHTMIFDNAFNCTFEYISD 166
TR|Q692E4|Q692E4_CVHSA         SQSVIIINNSTNVVIRACNFELCDNPFFAVSKPMG----TQTHTMIFDNAFNCTFEYISD 166
SP|Q3LZX1|SPIKE_BCHK3          TQSAVIVNNSTHIIIRVCNFNLCKEPMYTVSR--G----TQQNAWVYQSAFNCTYDRVEK 170
SP|Q3I5J5|SPIKE_BCRP3          TQSAVIVNNSTHIIIRVCNFNLCKEPMYTVSR--G----AQQSSWVYQSAFNCTYDRVEK 170
SP|Q0Q475|SPIKE_BC279          TQSAIIVNNSTHIIIRVCNFNLCKEPMYTVSK--G----TQQSSWVYQSAFNCTYDRVEK 170
TR|R9QTA0|R9QTA0_CVHSA         SQSAIIVNNSTHILVKVCNFVLCTEPMFTVSR--N----QHYKSWVYQHARNCTYDVAYP 168
TR|R9QTH3|R9QTH3_CVHSA         TQSAIIVNNSTHIIIRVCNFNLCKEPMFTVSR--G----VHFSSWVYQSAFNCTYDRVEK 170
TR|A0A0K1Z074|A0A0K1Z074_CVHSA TQSAIIVNNSTHIIIRVCYFNLCKDPMYTVSA--G----TQISSWVYQNAFNCTYDRVEK 170
TR|A0A0U1WHI6|A0A0U1WHI6_CVHSA TQSAIIVNNSTHIIIRVCYFNLCKDPMYTVSA--G----TQVSSWVYQSAFNCTYDRVEK 168
TR|A0A166ZL64|A0A166ZL64_9NIDO TQSALLVNNGSAITIEVCYFQFCDNPAFIISG--G----AQTNTAIYINLRNCTYVDTLR 165
TR|A0A0U1UYX4|A0A0U1UYX4_CVHSA TQSALLVNNGSAITIEVCYFQFCDNPAFIIRD--G----AQINTAIYINLRNCTYVDTLR 165
                               :** :::**.: : :..* * :* :*   :    .     .    ::    ***:
```

```
YP_009724390                   PFLMDLEGK-QGNFKNLREFVFKNIDGYFKIYSKHTPINLVR----DLPQGFSALEPLVD 228
SP|P59594|SPIKE_CVHSA          AFSLDVSEK-SGNFKHLREFVFKNKDGFLYVYKGYQPIDVVR----DLPSGFNTLKPIFK 221
TR|Q6UZF4|Q6UZF4_CVHSA         AFSLDVSEK-SGNFKHLREFVFKNKDGFLYVYKGYQPIDVVR----DLPSGFNTLKPIFK 221
TR|Q6UZF0|Q6UZF0_CVHSA         AFSLDVSEK-SGNFKHLREFVFKNKDGFLYVYKGYQPIDVVR----DLPSGFNTLKPIFK 221
TR|Q6JH46|Q6JH46_CVHSA         AFSLDVSEK-SGNFKHLREFVFKNKDGFLYVYKGYQPIDVVR----DLPSGFNTLKPIFK 221
TR|Q692E4|Q692E4_CVHSA         AFSLDVSEK-SGNFKHLREFVFKNKDGFLYVYKGYQPIDVVR----DLPSGFNTLKPIFK 221
SP|Q3LZX1|SPIKE_BCHK3          SFQLDTTPK-TGNFKDLREYVFKNRDGFLSVYQTYTAVNLPR----GLPTGFSVLKPILK 225
SP|Q3I5J5|SPIKE_BCRP3          SFQLDTAPK-TGNFKDLREYVFKNRDGFLSVYQTYTAVNLPR----GLPIGFSVLRPILK 225
SP|Q0Q475|SPIKE_BC279          SFQLDTAPK-TGNFKDLREYVFKNRDGFLSVYQTYTAVNLPR----GFPAGFSVLRPILK 225
TR|R9QTA0|R9QTA0_CVHSA         SFQLDVSLKNNVNFQHLREFIFKNVDGFLKIYSSYEPINVVS----GIPSGFSVLKPVMS 224
TR|R9QTH3|R9QTH3_CVHSA         SFQLDTAPK-TGNFKDLREYVFKNRDGFLSVYHSYTPVDIIR----GIPVGFSVLKPILK 225
TR|A0A0K1Z074|A0A0K1Z074_CVHSA SFQLDTAPK-SGNFIALREFVFKNRDGFFTVYQDYTPVNLLR----GLPAGLSVLKPILK 225
TR|A0A0U1WHI6|A0A0U1WHI6_CVHSA SFQLDTSPK-TGNFTDLREFVFKNRDGFFTVYQTYTPVNLLR----GLPSGLSVLKPILK 223
TR|A0A166ZL64|A0A166ZL64_9NIDO DLPLSFED-VDGGFKHLREFVFKNSDGFLHIYGAYQPYDLAIGATAALPAQFLPLKPLWK 224
TR|A0A0U1UYX4|A0A0U1UYX4_CVHSA DLPLSFAE-VDGGFKHLREFVFKNSDGFLHIYGAYQPYDLAIGATAALPAQFLPLKPLWK 224
                                : :.       .*  ***::*** **:: :*  :   ::       :*  :  *.*: .

YP_009724390                   LPIGINITRFQTLLALHRSYLTPGDSSSGWTAGAAAYYVGYLQPRTFLLKYNENGTITDA 288
SP|P59594|SPIKE_CVHSA          LPLGINITNFRAILTAFS------PAQDIWGTSAAAYFVGYLKPTTFMLKYDENGTITDA 275
TR|Q6UZF4|Q6UZF4_CVHSA         LPLGINITNFRAILTAFS------PAQDIWGTSAAAYFVGYLKPTTFMLKYDENGTITDA 275
TR|Q6UZF0|Q6UZF0_CVHSA         LPLGINITNFRAILTAFS------PAQDIWGTSAAAYFVGYLKPTTFMLKYDENGTITDA 275
TR|Q6JH46|Q6JH46_CVHSA         LPLGINITNFRAILTAFS------PAQDIWGTSAAAYFVGYLKPTTFMLKYDENGTITDA 275
TR|Q692E4|Q692E4_CVHSA         LPLGINITNFRAILTAFS------PAQDTWGTSAAAYFVGYLKPTTFMLKYDENGTITDA 275
SP|Q3LZX1|SPIKE_BCHK3          LPFGINITSYRVVMAMFS------QTTSNFLPESAAYYVGNLKYSTFMLRFNENGTITDA 279
SP|Q3I5J5|SPIKE_BCRP3          LPFGINITSYRVVMAMFS------QTTSNFLPESAAYYVGNLKYTTFMLSFNENGTITNA 279
SP|Q0Q475|SPIKE_BC279          LPFGINITSYRVVMTMFS------QFNSNFLPESAAYYVGNLKYTTFMLSFNENGTITDA 279
TR|R9QTA0|R9QTA0_CVHSA         LPLGINITGMRVVMTMFS------NTQANFLTENAAYYVGYLKPRTFMLQFNTNGTIVNA 278
TR|R9QTH3|R9QTH3_CVHSA         LPIGINITSFKVVMTMYS------QTTSNFLSESAAYYVGNLKYVTFMFQFNENGTIADA 279
TR|A0A0K1Z074|A0A0K1Z074_CVHSA LPFGINITSFRVVMAMFS------KTTSNYVPESAAYYVGNLKQSTFMLSFNQNGTITDA 279
TR|A0A0U1WHI6|A0A0U1WHI6_CVHSA LPFGINITSFRVVMAMFS------KTTSNYVPESAAYYVGNLKQSTFMLSFNQNGTIVDA 277
TR|A0A166ZL64|A0A166ZL64_9NIDO LPLGLNITNYKVVTTLKP------TNQ----AFQAAYIVGNLKHTTMMLSFNENGTMSNA 274
TR|A0A0U1UYX4|A0A0U1UYX4_CVHSA LPLGLNITNYKVVTTLKP------TNQ----AFQAAYIVGNLKHTTMMLSFNENGTMSNA 274
                               **:*:***  :.: :                   *** ** *:  *::: :: ***: :*

YP_009724390                   VDCALDPLSETKCTLKSFTVEKGIYQTSNFRVQPTESIVRFPNITNLCPFGEVFNATRFA 348
SP|P59594|SPIKE_CVHSA          VDCSQNPLAELKCSVKSFEIDKGIYQTSNFRVVPSGDVVRFPNITNLCPFGEVFNATKFP 335
TR|Q6UZF4|Q6UZF4_CVHSA         VDCSQNPLAELKCSVKSFEIDKGIYQTSNFRVVPSGDVVRFPNITNLCPFGEVFNATKFP 335
TR|Q6UZF0|Q6UZF0_CVHSA         VDCSQNPLAELKCSVKSFEIDKGIYQTSNFRVVPSGDVVRFPNITNLCPFGEVFNATKFP 335
TR|Q6JH46|Q6JH46_CVHSA         VDCSQNPLAELKCSVKSFEIDKGIYQTSNFRVVPSGDVVRFPNITNLCPFGEVFNATKFP 335
TR|Q692E4|Q692E4_CVHSA         VDCSQNPLAELKCSVKSFEIDKGIYQTSNFRVVPSGDVVRFPNITNLCPFGEVFNATKFP 335
SP|Q3LZX1|SPIKE_BCHK3          VDCSQNPLAELKCTIKNFNVDKGIYQTSNFRVSPTQEVIRFPNITNRCPFDKVFNATRFP 339
SP|Q3I5J5|SPIKE_BCRP3          IDCAQNPLAELKCTIKNFNVSKGIYQTSNFRVSPTQEVIRFPNITNRCPFDKVFNATRFP 339
SP|Q0Q475|SPIKE_BC279          VDCSQNPLAELKCTIKNFNVSKGIYQTSNFRVTPTQEVVRFPNITNRCPFDKVFNASRFP 339
TR|R9QTA0|R9QTA0_CVHSA         VDCSQDPLSELKCTLKNFNITKGIYQTSNFRVSPTQEVVRFPNITNRCPFDKVFNATRFP 338
TR|R9QTH3|R9QTH3_CVHSA         VDCSQNPLAELKCTLKNFNVSKGIYQTSNFRVSPSTEVIRFPNITNRCPFDRVFNASRFP 339
TR|A0A0K1Z074|A0A0K1Z074_CVHSA VDCSQNPLAELKCTTKSFNVSKGIYQTSNFRVAPVTEVVRFPNITNLCPFDKVFNATRFP 339
TR|A0A0U1WHI6|A0A0U1WHI6_CVHSA VDCSQDPLAELKCTTKSFNVSKGIYQTSNFRVSPVTEVVRFPNITNLCPFDKVFNATRFP 337
TR|A0A166ZL64|A0A166ZL64_9NIDO IDCSQDPLAELKCTLKQFDVGKGIYQTSNFRVQPTVDVARFPNITNVCPFDKVFNATRFP 334
TR|A0A0U1UYX4|A0A0U1UYX4_CVHSA IDCSQDPLAELKCTLKQFDVGKGIYQTSNFRVQPTVDVARFPNITNVCPFDKVFNATRFP 334
                               :**: :**:* **: *.* : *********** *  .: ******* ***..****::*

YP_009724390                   SVYAWNRKRISNCVADYSVLYNSASFSTFKCYGVSPTKLNDLCFTNVYADSFVIRGDEVR 408
SP|P59594|SPIKE_CVHSA          SVYAWERKKISNCVADYSVLYNSTFFSTFKCYGVSATKLNDLCFSNVYADSFVVKGDDVR 395
TR|Q6UZF4|Q6UZF4_CVHSA         SVYAWERKKISNCVADYSVLYNSTFFSTFKCYGVSATKLNDLCFSNVYADSFVVKGDDVR 395
TR|Q6UZF0|Q6UZF0_CVHSA         SVYAWERKKISNCVADYSVLYNSTFFSTFKCYGVSATKLNDLCFSNVYADSFVVKGDDVR 395
TR|Q6JH46|Q6JH46_CVHSA         SVYAWERKKISNCVADYSVLYNSTFFSTFKCYGVSATKLNDLCFSNVYADSFVVKGDDVR 395
TR|Q692E4|Q692E4_CVHSA         SVYAWERKKISNCVADYSVLYNSTFFSTFKCYGVSATKLNDLCFSNVYADSFVVKGDDVR 395
SP|Q3LZX1|SPIKE_BCHK3          NVYAWERTKISDCVADYTVLYNSTSFSTFKCYGVSPSKLIDLCFTSVYADTFLIRSSEVR 399
SP|Q3I5J5|SPIKE_BCRP3          NVYAWERTKISDCVADYTVLYNSTSFSTFKCYGVSPSKLIDLCFTSVYADTFLIRSSEVR 399
SP|Q0Q475|SPIKE_BC279          NVYAWERTKISDCVADYTVLYNSTSFSTFKCYGVSPSKLIDLCFTSVYADTFLIRSSEVR 399
TR|R9QTA0|R9QTA0_CVHSA         SVYAWERTKISDCVADYTVLYNSTSFSTFKCYGVSPSKLIDLCFTSVYADTFLIRSSEVR 398
TR|R9QTH3|R9QTH3_CVHSA         SVYAWERTKISDCVADYTVLYNSTSFSTFKCYGVSPSKLIDLCFTSVYADTFLIRFSEVR 399
TR|A0A0K1Z074|A0A0K1Z074_CVHSA SVYAWERTKISDCVADYTVFYNSTSFSTFNCYGVSPSKLIDLCFTSVYADTFLIRFSEVR 399
TR|A0A0U1WHI6|A0A0U1WHI6_CVHSA SVYAWERTKISDCVADYTVFYNSTSFSTFNCYGVSPSKLIDLCFTSVYADTFLIRFSEVR 397
TR|A0A166ZL64|A0A166ZL64_9NIDO SVYAWERTKISDCVADYTVFYNSTSFSTFNCYGVSPSKLIDLCFTSVYADTFLIRFSEVR 394
TR|A0A0U1UYX4|A0A0U1UYX4_CVHSA SVYAWERTKISDCVADYTVFYNSTSFSTFNCYGVSPSKLIDLCFTSVYADTFLIRFSEVR 394
                               .****:*.:**:*****:*:***: ****:***** :** ****:.****:*::: .:**
```

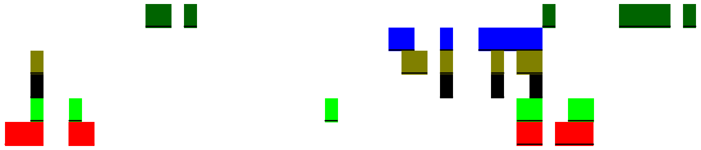

```
YP_009724390                   QIAPGQTGKIADYNYKLPDDFTGCVIAWNSNNLDSKVGGNYNYLYRLFRKSNLKPFERDI 468
SP|P59594|SPIKE_CVHSA          QIAPGQTGVIADYNYKLPDDFMGCVLAWNTRNIDATSTGNYNYKYRYLRHGKLRPFERDI 455
TR|Q6UZF4|Q6UZF4_CVHSA         QIAPGQTGVIADYNYKLPDDFMGCVLAWNTRNIDATSTGNYNYKYRYLRHGKLRPFERDI 455
TR|Q6UZF0|Q6UZF0_CVHSA         QIAPGQTGVIADYNYKLPDDFMGCVLAWNTRNIDATSTGNYNYKYRYLRHGKLRPFERDI 455
TR|Q6JH46|Q6JH46_CVHSA         QIAPGQTGVIADYNYKLPDDFMGCVLAWNTRNIDATSTGNYNYKYRSLRHGKLRPFERDI 455
TR|Q692E4|Q692E4_CVHSA         QIAPGQTGVIADYNYKLPDDFMGCVLAWNTRNIDATSTGNYNYKYRYLRHGKLRPFERDI 455
SP|Q3LZX1|SPIKE_BCHK3          QVAPGETGVIADYNYKLPDDFTGCVIAWNTAKHDTG-----NYYYRSHRKTKLKPFERDL 454
SP|Q3I5J5|SPIKE_BCRP3          QVAPGETGVIADYNYKLPDDFTGCVIAWNTAKQDQG-----QYYYRSHRKTKLKPFERDL 454
SP|Q0Q475|SPIKE_BC279          QVAPGETGVIADYNYKLPDDFTGCVIAWNTAQQDQG-----QYYYRSYRKEKLKPFERDL 454
TR|R9QTA0|R9QTA0_CVHSA         QVAPGETGVIADYNYKLPDDFTGCVIAWNTANQDQG-----QYYYRSSRKEKLKPFERDL 453
TR|R9QTH3|R9QTH3_CVHSA         QIAPGETGVIADYNYKLPDEFTGCVIAWNTANQDRG-----QYYYRSSRKTKLKPFERDL 454
TR|A0A0K1Z074|A0A0K1Z074_CVHSA QVAPGQTGVIADYNYKLPDDFTGCVIAWNTAKYDVG-----SYFYRSHRSSKLKPFERDL 454
TR|A0A0U1WHI6|A0A0U1WHI6_CVHSA QVAPGQTGVIADYNYKLPDDFTGCVIAWNTAKQDVG-----SYFYRSHRSSKLKPFERDL 452
TR|A0A166ZL64|A0A166ZL64_9NIDO QVAPGQTGVIADYNYKLPDDFIGCVIAWNTAKQDVG-----SYFYRSHRSSKLKPFERDL 449
TR|A0A0U1UYX4|A0A0U1UYX4_CVHSA QVAPGQTGVIADYNYKLPDDFIGCVIAWNTAKQDVG-----SYFYRSHRSSKLKPFERDL 449
                               *:***:** **********:* ***:***: : *       .* **  *  :*:*****:
```

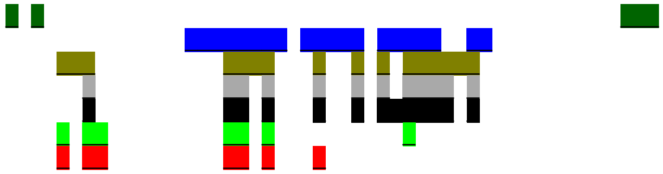

```
YP_009724390                   STEIYQAGSTPCNGVEGFNCYFPLQSYGFQPTNGVGYQPYRVVVLSFELLHAPATVCGPK 528
SP|P59594|SPIKE_CVHSA          SNVPFSPDGKPCTPP-ALNCYWPLNDYGFYTTTGIGYQPYRVVVLSFELLNAPATVCGPK 514
TR|Q6UZF4|Q6UZF4_CVHSA         SNVPFSPDGKPCTPP-ALNCYWPLNDYGFYTTTGIGYQPYRVVVLSFELLNAPATVCGPK 514
TR|Q6UZF0|Q6UZF0_CVHSA         SNVPFSPDGKPCTPP-ALNCYWPLNDYGFYTTTGIGYQPYRVVVLSFELLNAPATVCGPK 514
TR|Q6JH46|Q6JH46_CVHSA         SNVPFSPDGKPCTPP-ALNCYWPLNDYGFYTTTGIGYQPYRVVVLSFELLNAPATVCGPK 514
TR|Q692E4|Q692E4_CVHSA         SNVPFSPDGKPCTPP-ALNCYWPLNDYGFYTTTGIGYQPYRVVVLSFELLNAPATVCGPK 514
SP|Q3LZX1|SPIKE_BCHK3          SSDD-------------GNGVYTLSTYDFNPNVPVAYQATRVVVLSFELLNAPATVCGPK 501
SP|Q3I5J5|SPIKE_BCRP3          SSD--------------ENGVRTLSTYDFYPSVPVAYQATRVVVLSFELLNAPATVCGPK 500
SP|Q0Q475|SPIKE_BC279          SSD--------------ENGVYTLSTYDFYPSIPVEYQATRVVVLSFELLNAPATVCGPK 500
TR|R9QTA0|R9QTA0_CVHSA         SSD--------------ENGVYTLSTYDFYPSVPLDYQATRVVVLSFELLNAPATVCGPK 499
TR|R9QTH3|R9QTH3_CVHSA         SSD--------------ENGVRTLSTYDFYPSVPLEYQATRVVVLSFELLNAPATVCGPK 500
TR|A0A0K1Z074|A0A0K1Z074_CVHSA SSE--------------ENGARTLSTYDFNQNVPLEYQATRVVVLSFELLNAPATVCGPK 500
TR|A0A0U1WHI6|A0A0U1WHI6_CVHSA SSE--------------ENGVRTLSTYDFNQYVPLEYQATRVVVLSFELLNAPATVCGPK 498
TR|A0A166ZL64|A0A166ZL64_9NIDO SSE--------------ENGVLTLSTYDFNQNVPLEYQATRVVVLSFELLNAPATVCGPK 495
TR|A0A0U1UYX4|A0A0U1UYX4_CVHSA SSE--------------ENGVLTLSTYDFNQNVPLEYQATRVVVLSFELLNAPATVCGPK 495
                               *.                *    *. *.*     : **  **********:*********
```

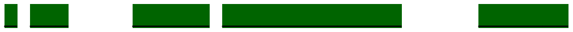

```
YP_009724390                   KSTNLVKNKCVNFNFNGLTGTGVLTESNKKFLPFQQFGRDIADTTDAVRDPQTLEILDIT 588
SP|P59594|SPIKE_CVHSA          LSTDLIKNQCVNFNFNGLTGTGVLTPSSKRFQPFQQFGRDVSDFTDSVRDPKTSEILDIS 574
TR|Q6UZF4|Q6UZF4_CVHSA         LSTDLIKNQCVNFNFNGLTGTGVLTPSSKRFQPFQQFGRDVSDFTDSVRDPKTSEILDIS 574
TR|Q6UZF0|Q6UZF0_CVHSA         LSTDLIKNQCVNFNFNGLTGTGVLTPSSKRFQPFQQFGRDVSDFTDSVRDPKTSEILDIS 574
TR|Q6JH46|Q6JH46_CVHSA         LSTDLIKNQCVNFNFNGLTGTGVLTPSSKRFQPFQQFGRDVSDFTDSVRDPKTSEILDIS 574
TR|Q692E4|Q692E4_CVHSA         LSTDLIKNQCVNFNFNGLTGTGVLTPSSKRFQPFQQFGRDVSDFTDSVRDPKTSEILDIS 574
SP|Q3LZX1|SPIKE_BCHK3          LSTELVKNQCVNFNFNGLKGTGVLTSSSKRFQSFQQFGRDTSDFTDSVRDPQTLEILDIS 561
SP|Q3I5J5|SPIKE_BCRP3          LSTQLVKNQCVNFNFNGLKGTGVLTESSKRFQSFQQFGRDTSDFTDSVRDPQTLEILDIS 560
SP|Q0Q475|SPIKE_BC279          LSTQLVKNQCVNFNFNGLRGTGVLTTSSKRFQSFQQFGRDTSDFTDSVRDPQTLEILDIS 560
TR|R9QTA0|R9QTA0_CVHSA         LSTTLVKNQCVNFNFNGLKGTGVLTASSKKFQSFQQFGRDASDFTDSVRDPQTLEILDIS 559
TR|R9QTH3|R9QTH3_CVHSA         LSTSLIKNQCVNFNFNGLKGTGVLTDSSKKFQSFQQFGRDASDFTDSVRDPQTLQILDIS 560
TR|A0A0K1Z074|A0A0K1Z074_CVHSA LSTSLVKNQCVNFNFNGFKGTGVLTDSSKTFQSFQQFGRDASDFTDSVRDPKTLQILDIS 560
TR|A0A0U1WHI6|A0A0U1WHI6_CVHSA LSTSLVKNQCVNFNFNGFKGTGVLTDSSKTFQSFQQFGRDASDFTDSVRDPQTLRILDIS 558
TR|A0A166ZL64|A0A166ZL64_9NIDO LSTPLVKNQCVNFNFNGLKGTGVLTDSSKTFQSFQQFGRDASDFTDSVRDPQTLQILDIS 555
TR|A0A0U1UYX4|A0A0U1UYX4_CVHSA LSTPLVKNQCVNFNFNGLKGTGVLTDSSKTFQSFQQFGRDASDFTDSVRDPQTLQILDIS 555
                                ** *:**:********: ****** *.* *  ******* :* **:****:* .****:
```

```
YP_009724390                          PCSFGGVSVITPGTNTSNQVAVLYQDVNCTEVPVAIHADQLTPTWRVYSTGSNVFQTRAG 648
SP|P59594|SPIKE_CVHSA                 PCSFGGVSVITPGTNASSEVAVLYQDVNCTDVSTAIHADQLTPAWRIYSTGNNVFQTQAG 634
TR|Q6UZF4|Q6UZF4_CVHSA                PCSFGGVSVITPGTNASSEVAVLYQDVNCTDVSTAIHADQLTPAWRIYSTGNNVFQTQAG 634
TR|Q6UZF0|Q6UZF0_CVHSA                PCSFGGVSVITPGTNASSEVAVLYQDVNCTDVSTAIHADQLTPAWRIYSTGNNVFQTQAG 634
TR|Q6JH46|Q6JH46_CVHSA                PCSFGGVSVITPGTNASSEVAVLYQDVNCTDVSTAIHADQLTPAWRIYSTGNNVFQTQAG 634
TR|Q692E4|Q692E4_CVHSA                PCSFGGVSVITPGTNASSEVAVLYQDVNCTDVSTAIHADQLTPAWRIYSTGNNVFQTQAG 634
SP|Q3LZX1|SPIKE_BCHK3                 PCSFGGVSVITPGTNASSEVAVLYQDVNCTDVPTAIRADQLTPAWRVYSTGVNVFQTQAG 621
SP|Q3I5J5|SPIKE_BCRP3                 PCSFGGVSVITPGTNASSEVAVLYQDVNCTDVPAAIHADQLTPAWRVYSTGTNVFQTQAG 620
SP|Q0Q475|SPIKE_BC279                 PCSFGGVSVITPGTNASSEVAVLYQDVNCTDVPTSIHADQLTPAWRVYSTGVNVFQTQAG 620
TR|R9QTA0|R9QTA0_CVHSA                PCSFGGVSVITPGTNASTEVAVLYQDVNCTDVPTAINADQLTPAWRVYSTGINVFQTQAG 619
TR|R9QTH3|R9QTH3_CVHSA                PCSFGGVSVITPGTNASSEVAVLYQDVNCTDVPTAIRADQLTPAWRVYSAGVNVFQTQAG 620
TR|A0A0K1Z074|A0A0K1Z074_CVHSA        PCSFGGVSVITPGTNTSSAVAVLYQDVNCTDVPKTIHADQLAPSWRVYTSGPFVFQTQAG 620
TR|A0A0U1WHI6|A0A0U1WHI6_CVHSA        PCSFGGVSVITPGTNTSSAVAVLYQDVNCTDVPTTLHADQLAPSWRVYTTGPYVFQTQAG 618
TR|A0A166ZL64|A0A166ZL64_9NIDO        PCSFGGVSVITPGTNTSSAVAVLYQDVNCTDVPTTIHADHLTHSWRVYTTGPYVFQTQAG 615
TR|A0A0U1UYX4|A0A0U1UYX4_CVHSA        PCSFGGVSVITPGTNTSSAVAVLYQDVNCTDVPTTIHADHLTHSWRVYTTGPYVFQTQAG 615
                                      ***************:*. ***********:*  ::.**:*: :**:*::*  ****:**

YP_009724390                          CLIGAEHVNNSYECDIPIGAGICASYQTQTNSPRRARSVASQSIIAYTMSLGAENSVAYS 708
SP|P59594|SPIKE_CVHSA                 CLIGAEHVDTSYECDIPIGAGICASYHTVS----LLRSTSQKSIVAYTMSLGADSSIAYS 690
TR|Q6UZF4|Q6UZF4_CVHSA                CLIGAEHVDTSYECDIPIGAGICASYHTVS----LLRSTSQKSIVAYTMSLGADSSIAYS 690
TR|Q6UZF0|Q6UZF0_CVHSA                CLIGAEHVDTSYECDIPIGAGICASYHTVS----LLRSTSQKSIVAYTMSLGADSSIAYS 690
TR|Q6JH46|Q6JH46_CVHSA                CLIGAEHVDTSYECDIPIGAGICASYHTVS----LLRSTSQKSIVAYTMSLGADSSIAYS 690
TR|Q692E4|Q692E4_CVHSA                CLIGAEHVDTSYECDIPIGAGICASYHTVS----LLRNTSQKSIVAYTMSLGADSSIAYS 690
SP|Q3LZX1|SPIKE_BCHK3                 CLIGAEHVNASYECDIPIGAGICASYHTAS----VLRSTGQKSIVAYTMSLGAENSIAYA 677
SP|Q3I5J5|SPIKE_BCRP3                 CLIGAEHVNASYECDIPIGAGICASYHTAS----TLRSVGQKSIVAYTMSLGAENSIAYA 676
SP|Q0Q475|SPIKE_BC279                 CLIGAEHVNASYECDIPIGAGICASYHTAS----VLRSTGQKSIVAYTMSLGAENSIAYA 676
TR|R9QTA0|R9QTA0_CVHSA                CLIGAEHVNASYECDIPIGAGICASYHTAS----VLRSTGQKSIVAYTMSLGAENSIAYA 675
TR|R9QTH3|R9QTH3_CVHSA                CLIGAEHVNASYECDIPIGAGICASYHTAS----LLRNTGQKSIVAYTMSLGAENSIAYA 676
TR|A0A0K1Z074|A0A0K1Z074_CVHSA        CLIGAEHVNSSYECDIPIGAGICASYHTAS----VLRSTGQKSIVAYTMSLGAENSVAYA 676
TR|A0A0U1WHI6|A0A0U1WHI6_CVHSA        CLIGAEHVNASYQCDIPIGAGICASYHTAS----LLRSTGQKSIVAYTMSLGAENSVAYA 674
TR|A0A166ZL64|A0A166ZL64_9NIDO        CLIGAEHVNASYQCDIPIGAGICASYHTAS----LLRSTGQKSIVAYTMSLGAENSVAYA 671
TR|A0A0U1UYX4|A0A0U1UYX4_CVHSA        CLIGAEHVNASYQCDIPIGAGICASYHTAS----LLRSTGQKSIVAYTMSLGAENSVAYA 671
                                      ********: **:*************:* :      *....:**:********:.*:**:

YP_009724390                          NNSIAIPTNFTISVTTEILPVSMTKTSVDCTMYICGDSTECSNLLLQYGSFCTQLNRALT 768
SP|P59594|SPIKE_CVHSA                 NNTIAIPTNFSISITTEVMPVSMAKTSVDCNMYICGDSTECANLLLQYGSFCTQLNRALS 750
TR|Q6UZF4|Q6UZF4_CVHSA                NNTIAIPTNFSISITTEVMPVSMAKTSVDCNMYICGDSTECANLLLQYGSFCTQLNRALS 750
TR|Q6UZF0|Q6UZF0_CVHSA                NNTIAIPTNFSISITTEVMPVSMAKTSVDCNMYICGDSTECANLLLQYGSFCTQLNRALS 750
TR|Q6JH46|Q6JH46_CVHSA                NNTIAIPTNFSISITTEVMPVSMAKTSVDCNMYICGDSTECANLLLQYGSFCTQLNRALS 750
TR|Q692E4|Q692E4_CVHSA                NNTIAIPTNFSISITTEVMPVSMAKTSVDCNMYICGDSTECANLLLQYGSFCTQLNRALS 750
SP|Q3LZX1|SPIKE_BCHK3                 NNSIAIPTNFSISVTTEVMPVSMAKTAVDCTMYICGDSLECSNLLLQYGSFCTQLNRALT 737
SP|Q3I5J5|SPIKE_BCRP3                 NNSIAIPTNFSISVTTEVMPVSMAKTSVDCTMYICGDSLECSNLLLQYGSFCTQLNRALS 736
SP|Q0Q475|SPIKE_BC279                 NNSIAIPTNFSISVTTEVMPVSIAKTSVDCTMYICGDSLECSNLLLQYGSFCTQLNRALT 736
TR|R9QTA0|R9QTA0_CVHSA                NNSIAIPTNFSISVTTEVMPVSMAKTSVDCTMYICGDSLECSNLLLQYGSFCTQLNRALT 735
TR|R9QTH3|R9QTH3_CVHSA                NNSIAIPTNFSISVTTEVMPVSMAKTSVDCTMYICGDSQECSNLLLQYGSFCTQLNRALS 736
TR|A0A0K1Z074|A0A0K1Z074_CVHSA        NNSIAIPTNFSISVTTEVMPVSMAKTSVDCTMYICGDSAECSNLLLQYGSFCTQLNRALS 736
TR|A0A0U1WHI6|A0A0U1WHI6_CVHSA        NNSIAIPTNFSISVTTEVMPVSMAKTSVDCTMYICGDSLECSNLLLQYGSFCTQLNRALS 734
TR|A0A166ZL64|A0A166ZL64_9NIDO        NNSIAIPTNFSISVTTEVMPVSMAKTSVDCTMYICGDSLECSNLLLQYGSFCTQLNRALS 731
TR|A0A0U1UYX4|A0A0U1UYX4_CVHSA        NNSIAIPTNFSISVTTEVMPVSMAKTSVDCTMYICGDSLECSNLLLQYGSFCTQLNRALS 731
                                      **:*******:**:***::***::**:***.******* **:*****************:

YP_009724390                          GIAVEQDKNTQEVFAQVKQIYKTPPIKDFGGFNFSQILPDPSKPSKRSFIEDLLFNKVTL 828
SP|P59594|SPIKE_CVHSA                 GIAAEQDRNTREVFAQVKQMYKTPTLKYFGGFNFSQILPDPLKPTKRSFIEDLLFNKVTL 810
TR|Q6UZF4|Q6UZF4_CVHSA                GIAAEQDRNTREVFAQVKQMYKTPTLKYFGGFNFSQILPDPLKPTKRSFIEDLLFNKVTL 810
TR|Q6UZF0|Q6UZF0_CVHSA                GIAAEQDRNTREVFAQVKQMYKTPTLKYFGGFNFSQILPDPLKPTKRSFIEDLLFNKVTL 810
TR|Q6JH46|Q6JH46_CVHSA                GIAAEQDRNTREVFAQVKQMYKTPTLKYFGGFNFSQILPDPLKPTKRSFIEDLLFNKVTL 810
TR|Q692E4|Q692E4_CVHSA                GIAAEQDRNTREVFAQVKQMYKTPTLKYFGGFNFSQILPDPLKPTKRSFIEDLLFNKVTL 810
SP|Q3LZX1|SPIKE_BCHK3                 GIAIEQDKNTQEVFAQVKQMYKTPAIKDFGGFNFSQILPDPSKPTKRSFIEDLLFNKVTL 797
SP|Q3I5J5|SPIKE_BCRP3                 GIAIEQDKNTQEVFAQVKQMYKTPAIKDFGGFNFSQILPDPSKPTKRSFIEDLLFNKVTL 796
SP|Q0Q475|SPIKE_BC279                 GIAIEQDKNTQEVFAQVKQMYKTPAIKDFGGFNFSQILPDPSKPTKRSFIEDLLFNKVTL 796
TR|R9QTA0|R9QTA0_CVHSA                GIAIEQDKNTQEVFAQVKQMYKTPAIKDFGGFNFSQILPDPSKPTKRSFIEDLLFNKVTL 795
TR|R9QTH3|R9QTH3_CVHSA                GIAVEQDKNTQEVFAQVKQMYKTPAIKDFGGFNFSQILPDPSKPTKRSFIEDLLFNKVTL 796
TR|A0A0K1Z074|A0A0K1Z074_CVHSA        GIAVEQDKNTQEVFAQVKQMYKTPTLRDFGGFNFSQILPDPLKPTKRSFIEDLLYNKVTL 796
TR|A0A0U1WHI6|A0A0U1WHI6_CVHSA        GIAVEQDKNTQEVFAQVKQMYKTPTIRDFGGFNFSQILPDPLKPTKRSFIEDLLYNKVTL 794
TR|A0A166ZL64|A0A166ZL64_9NIDO        GIAVEQDKNTQEVFAQVKQMYKTPTIRDFGGFNFSQILPDPLKPTKRSFIEDLLYNKVTL 791
TR|A0A0U1UYX4|A0A0U1UYX4_CVHSA        GIAVEQDKNTQEVFAQVKQMYKTPTIRDFGGFNFSQILPDPLKPTKRSFIEDLLYNKVTL 791
                                      *** ***:**:********:**** :: ************* **:*********:*****
```

```
YP_009724390                   ADAGFIKQYGDCLGDIAARDLICAQKFNGLTVLPPLLTDEMIAQYTSALLAGTITSGWTF 888
SP|P59594|SPIKE_CVHSA          ADAGFMKQYGECLGDINARDLICAQKFNGLTVLPPLLTDDMIAAYTAALVSGTATAGWTF 870
TR|Q6UZF4|Q6UZF4_CVHSA         ADAGFMKQYGECLGDINARDLICAQKFNGLTVLPPLLTDDMIAAYTAALVSGTATAGWTF 870
TR|Q6UZF0|Q6UZF0_CVHSA         ADAGFMKQYGECLGDINARDLICAQKFNGLTVLPPLLTDDMIAAYTAALVSGTATAGWTF 870
TR|Q6JH46|Q6JH46_CVHSA         ADAGFMKQYGECLGDINARDLICAQKFNGLTVLPPLLTDDMIAAYTAALVSGTATAGWTF 870
TR|Q692E4|Q692E4_CVHSA         ADAGFMKQYGECLGDINARDLICAQKFNGLTVLPPLLTDDMIAAYTAALVSGTATAGWTF 870
SP|Q3LZX1|SPIKE_BCHK3          ADAGFMKQYGDCLGDVSARDLICAQKFNGLTVLPPLLTDEMVAAYTAALVSGTATAGWTF 857
SP|Q3I5J5|SPIKE_BCRP3          ADAGFMKQYGECLGDISARDLICAQKFNGLTVLPPLLTDEMIAAYTAALVSGTATAGWTF 856
SP|Q0Q475|SPIKE_BC279          ADAGFMKQYGECLGDISARDLICAQKFNGLTVLPPLLTDEMIAAYTAALVSGTATAGWTF 856
TR|R9QTA0|R9QTA0_CVHSA         ADAGFMKQYGECLGDISARDLICAQKFNGLTVLPPLLTDEMIAAYTAALVSGTATAGWTF 855
TR|R9QTH3|R9QTH3_CVHSA         ADAGFMKQYGECLGDISARDLICAQKFNGLTVLPPLLTDEMIAAYTAALVSGTATAGWTF 856
TR|A0A0K1Z074|A0A0K1Z074_CVHSA ADAGFMKQYADCLGGINARDLICAQKFNGLTVLPPLLTDDMIAAYTAALISGTATAGWTF 856
TR|A0A0U1WHI6|A0A0U1WHI6_CVHSA ADAGFMKQYADCLGGINARDLICAQKFNGLTVLPPLLTDDMIAAYTAALISGTATAGWTF 854
TR|A0A166ZL64|A0A166ZL64_9NIDO ADAGFMKQYADCLGGINARDLICAQKFNGLTVLPPLLTDDMIAAYTAALISGTATAGWTF 851
TR|A0A0U1UYX4|A0A0U1UYX4_CVHSA ADAGFMKQYADCLGGINARDLICAQKFNGLTVLPPLLTDDMIAAYTAALISGTATAGWTF 851
                               *****:***.:***.: **********************:*:* **:**::** *:****

YP_009724390                   GAGAALQIPFAMQMAYRFNGIGVTQNVLYENQKLIANQFNSAIGKIQDSLSSTASALGKL 948
SP|P59594|SPIKE_CVHSA          GAGAALQIPFAMQMAYRFNGIGVTQNVLYENQKQIANQFNKAISQIQESLTTTSTALGKL 930
TR|Q6UZF4|Q6UZF4_CVHSA         GAGAALQIPFAMQMAYRFNGIGVTQNVLYENQKQIANQFNKAISQIQESLTTTSTALGKL 930
TR|Q6UZF0|Q6UZF0_CVHSA         GAGAALQIPFAMQMAYRFNGIGVTQNVLYENQKQIANQFNKAISQIQESLTTTSTALGKL 930
TR|Q6JH46|Q6JH46_CVHSA         GAGAALQIPFAMQMAYRFNGIGVTQNVLYENQKQIANQFNKAISQIQESLTTTSTALGKL 930
TR|Q692E4|Q692E4_CVHSA         GAGAALQIPFAMQMAYRFNGIGVTQNVLYENQKQIANQFNKAISQIQESLTTTSTALGKL 930
SP|Q3LZX1|SPIKE_BCHK3          GAGAALQIPFAMQMAYRFNGIGVTQNVLYENQKLIANQFNSAIGKIQESLSSTASALGKL 917
SP|Q3I5J5|SPIKE_BCRP3          GAGSALQIPFAMQMAYRFNGIGVTQNVLYENQKQIANQFNKAISQIQESLTTTSTALGKL 916
SP|Q0Q475|SPIKE_BC279          GAGSALQIPFAMQMAYRFNGIGVTQNVLYENQKQIANQFNKAISQIQESLTTTSTALGKL 916
TR|R9QTA0|R9QTA0_CVHSA         GAGAALQIPFAMQMAYRFNGIGVTQNVLYENQKQIANQFNKAISQIQESLTTTSTALGKL 915
TR|R9QTH3|R9QTH3_CVHSA         GAGAALQIPFAMQMAYRFNGIGVTQNVLYENQKQIANQFNKAISQIQESLTTTSTALGKL 916
TR|A0A0K1Z074|A0A0K1Z074_CVHSA GAGAALQIPFAMQMAYRFNGIGVTQNVLYENQKQIANQFNKAITQIQESLTTTSTALGKL 916
TR|A0A0U1WHI6|A0A0U1WHI6_CVHSA GAGAALQIPFAMQMAYRFNGIGVTQNVLYENQKQIANQFNKAITQIQESLTTTSTALGKL 914
TR|A0A166ZL64|A0A166ZL64_9NIDO GAGAALQIPFAMQMAYRFNGIGVTQNVLYENQKQIANQFNKAITQIQESLTTTATALGKL 911
TR|A0A0U1UYX4|A0A0U1UYX4_CVHSA GAGAALQIPFAMQMAYRFNGIGVTQNVLYENQKQIANQFNKAITQIQESLTTTATALGKL 911
                               ***:***************************** ******.** :**:**::*::*****

YP_009724390                   QDVVNQNAQALNTLVKQLSSNFGAISSVLNDILSRLDKVEAEVQIDRLITGRLQSLQTYV 1008
SP|P59594|SPIKE_CVHSA          QDVVNQNAQALNTLVKQLSSNFGAISSVLNDILSRLDKVEAEVQIDRLITGRLQSLQTYV 990
TR|Q6UZF4|Q6UZF4_CVHSA         QDVVNQNAQALNTLVKQLSSNFGAISSVLNDILSRLDKVEAEVQIDRLITGRLQSLQTYV 990
TR|Q6UZF0|Q6UZF0_CVHSA         QDVVNQNAQALNTLVKQLSSNFGAISSVLNDILSRLDKVEAEVQIDRLITGRLQSLQTYV 990
TR|Q6JH46|Q6JH46_CVHSA         QDVVNQNAQALNTLVKQLSSNFGAISSVLNDILSRLDKVEAEVQIDRLITGRLQSLQTYV 990
TR|Q692E4|Q692E4_CVHSA         QDVVNQNAQALNTLVKQLSSNFGAISSVLNDILSRLDKVEAEVQIDRLITGRLQSLQTYV 990
SP|Q3LZX1|SPIKE_BCHK3          QDVVNQNAQALNTLVKQLSSNFGAISSVLNDILSRLDKVEAEVQIDRLITGRLQSLQTYV 977
SP|Q3I5J5|SPIKE_BCRP3          QDVVNQNAQALNTLVKQLSSNFGAISSVLNDILSRLDKVEAEVQIDRLITGRLQSLQTYV 976
SP|Q0Q475|SPIKE_BC279          QDVVNDNAQALNTLVKQLSSNFGAISSVLNDILSRLDKVEAEVQIDRLITGRLQSLQTYV 976
TR|R9QTA0|R9QTA0_CVHSA         QDVVNQNAQALNTLVKQLSSNFGAISSVLNDILSRLDKVEAEVQIDRLITGRLQSLQTYV 975
TR|R9QTH3|R9QTH3_CVHSA         QDVVNQNAQALNTLVKQLSSNFGAISSVLNDILSRLDKVEAEVQIDRLITGRLQSLQTYV 976
TR|A0A0K1Z074|A0A0K1Z074_CVHSA QDVVNQNAQALNTLVKQLSSNFGAISSVLNDILSRLDKVEAEVQIDRLITGRLQSLQTYV 976
TR|A0A0U1WHI6|A0A0U1WHI6_CVHSA QDVVNQNAQALNTLVKQLSSNFGAISSALNDILSRLDKVEAEVQIDRLITGRLQSLQTYV 974
TR|A0A166ZL64|A0A166ZL64_9NIDO QDVVNQNAQALNTLVKQLSSNFGAISSALNDILSRLDKVEAEVQIDRLITGRLQSLQTYV 971
TR|A0A0U1UYX4|A0A0U1UYX4_CVHSA QDVVNQNAQALNTLVKQLSSNFGAISSALNDILSRLDKVEAEVQIDRLITGRLQSLQTYV 971
                               *****:*********************.********************************

YP_009724390                   TQQLIRAAEIRASANLAATKMSECVLGQSKRVDFCGKGYHLMSFPQSAPHGVVFLHVTYV 1068
SP|P59594|SPIKE_CVHSA          TQQLIRAAEIRASANLAATKMSECVLGQSKRVDFCGKGYHLMSFPQAAPHGVVFLHVTYV 1050
TR|Q6UZF4|Q6UZF4_CVHSA         TQQLIRAAEIRASANLAATKMSECVLGQSKRVDFCGKGYHLMSFPQAAPHGVVFLHVTYV 1050
TR|Q6UZF0|Q6UZF0_CVHSA         TQQLIRAAEIRASANLAATKMSECVLGQSKRVDFCGKGYHLMSFPQAAPHGVVFLHVTYV 1050
TR|Q6JH46|Q6JH46_CVHSA         TQQLIRAAEIRASANLAATKMSECVLGQSKRVDFCGKGYHLMSFPQAAPHGVVFLHVTYV 1050
TR|Q692E4|Q692E4_CVHSA         TQQLIRAAEIRASANLAATKMSECVLGQSKRVDFCGKGYHLMSFPQAAPHGVVFLHVTYV 1050
SP|Q3LZX1|SPIKE_BCHK3          TQQLIRAAEIRASANLAATKMSECVLGQSKRVDFCGKGYHLMSFPQSAPHGVVFLHVTYV 1037
SP|Q3I5J5|SPIKE_BCRP3          TQQLIRAAEIRASANLAATKMSECVLGQSKRVDFCGKGYHLMSFPQAAPHGVVFLHVTYV 1036
SP|Q0Q475|SPIKE_BC279          TQQLIRAAEIRASANLAATKMSECVLGQSKRVDFCGKGYHLMSFPQAAPHGVVFLHVTYV 1036
TR|R9QTA0|R9QTA0_CVHSA         TQQLIRAAEIRASANLAATKMSECVLGQSKRVDFCGKGYHLMSFPQAAPHGVVFLHVTYV 1035
TR|R9QTH3|R9QTH3_CVHSA         TQQLIRAAEIRASANLAATKMSECVLGQSKRVDFCGKGYHLMSFPQAAPHGVVFLHVTYV 1036
TR|A0A0K1Z074|A0A0K1Z074_CVHSA TQQLIRAAEIRASANLAATKMSECVLGQSKRVDFCGKGYHLMSFPQAAPHGVVFLHVTYV 1036
TR|A0A0U1WHI6|A0A0U1WHI6_CVHSA TQQLIRAAEIRTSANLAATKMSECVLGQSKRVDFCGKGYHLMSFPQSAPHGVVFLHVTYV 1034
TR|A0A166ZL64|A0A166ZL64_9NIDO TQQLIRAAEIRASANLAATKMSECVLGQSKRVDFCGKGYHLMSFPQSAPHGVVFLHVTYV 1031
TR|A0A0U1UYX4|A0A0U1UYX4_CVHSA TQQLIRAAEIRASANLAATKMSECVLGQSKRVDFCGKGYHLMSFPQSAPHGVVFLHVTYV 1031
                               ***********:**********************************:*************
```

```
YP_009724390                       PAQEKNFTTAPAICHDGKAHFPREGVFVSNGTHWFVTQRNFYEPQIITTDNTFVSGNCDV 1128
SP|P59594|SPIKE_CVHSA              PSQERNFTTAPAICHEGKAYFPREGVFVFNGTSWFITQRNFFSPQIITTDNTFVSGNCDV 1110
TR|Q6UZF4|Q6UZF4_CVHSA             PSQERNFTTAPAICHEGKAYFPREGVFVFNGTSWFITQRNFFSPQIITTDNTFVSGNCDV 1110
TR|Q6UZF0|Q6UZF0_CVHSA             PSQERNFTTAPAICHEGKAYFPREGVFVFNGTSWFITQRNFFSPQIITTDNTFVSGNCDV 1110
TR|Q6JH46|Q6JH46_CVHSA             PSQERNFTTAPAICHEGKAYFPREGVFVFNGTSWFITQRNFFSPQIITTDNTFVSGNCDV 1110
TR|Q692E4|Q692E4_CVHSA             PSQERNFTTAPAICHEGKAYFPREGVFVFNGTSWFITQRNFFSPQIITTDNTFVSGKCDV 1110
SP|Q3LZX1|SPIKE_BCHK3              PSQEKNFTTAPAICHEGKAYFPREGVFVSNGTSWFITQRNFYSPQLITTDNTFVSGNCDV 1097
SP|Q3I5J5|SPIKE_BCRP3              PSQERNFTTAPAICHEGKAYFPREGVFVSNGTSWFITQRNFYSPQIITTDNTFVAGSCDV 1096
SP|Q0Q475|SPIKE_BC279              PSQERNFTTAPAICHEGKAYFPREGVFVSNGTSWFITQRNFYSPQIITTDNTFVAGNCDV 1096
TR|R9QTA0|R9QTA0_CVHSA             PSQERNFTTAPAICHEGKAYFPREGVFVSNGTSWFITQRNFYSPQIITTDNTFVAGSCNV 1095
TR|R9QTH3|R9QTH3_CVHSA             PSQERNFTTAPAICHQGKAYFPREGVFVSNGTSWFITQRNFFSPQIITTDNTFVSGNCDV 1096
TR|A0A0K1Z074|A0A0K1Z074_CVHSA     PSQEKNFTTAPAICHEGKAYFPREGVFVFNGTSWFITQRNFYSPQTITTDNTFVAGNCDV 1096
TR|A0A0U1WHI6|A0A0U1WHI6_CVHSA     PSQERNFTTAPAICHEGKAYFPREGVFVSNGSSWFITQRNFYSPQIITTDNTFVAGSCDV 1094
TR|A0A166ZL64|A0A166ZL64_9NIDO     PAQEKNFTTAPAICHEGKAYFPREGVFVSNGSSWFITQRNFYSPQIITTDNTFVAGSCDV 1091
TR|A0A0U1UYX4|A0A0U1UYX4_CVHSA     PAQEKNFTTAPAICHEGKAYFPREGVFVSNGSSWFITQRNFYSPQIITTDNTFVAGSCDV 1091
                                   *:**:**********:***:******** **: **:*****:.** ********:*.*:*

YP_009724390                       VIGIVNNTVYDPLQPELDSFKEELDKYFKNHTSPDVDLGDISGINASVVNIQKEIDRLNE 1188
SP|P59594|SPIKE_CVHSA              VIGIINNTVYDPLQPELDSFKEELDKYFKNHTSPDVDLGDISGINASVVNIQKEIDRLNE 1170
TR|Q6UZF4|Q6UZF4_CVHSA             VIGIINNTVYDPLQPELDSFKEELDKYFKNHTSPDVDLGDISGINASVVNIQKEIDRLNE 1170
TR|Q6UZF0|Q6UZF0_CVHSA             VIGIINNTVYDPLQPELDSFKEELDKYFKNHTSPDVDLGDISGINASVVNIQKEIDRLNE 1170
TR|Q6JH46|Q6JH46_CVHSA             VIGIINNTVYDPLQPELDSFKEELDKYFKNHTSPDVDLGDISGINASVVNIQKEIDRLNE 1170
TR|Q692E4|Q692E4_CVHSA             VIGIINNTVYDPLQPELDSFKEELDKYFKNHTSPDVDLGDISGINASVVNIQKEIDRLNE 1170
SP|Q3LZX1|SPIKE_BCHK3              VIGIINNTVYDPLQPELDSFKEELDKYFKNHTSPDVDLGDISGINASVVNIQKEIDRLNE 1157
SP|Q3I5J5|SPIKE_BCRP3              VIGIINNTVYDPLQPELDSFKEELDKYFKNHTSPDVDLGDISGINASVVNIQKEIDRLNE 1156
SP|Q0Q475|SPIKE_BC279              VIGIINNTVYDPLQPELDSFKEELDKYFKNHTSPDVDLGDISGINASVVNIQKEIDRLNE 1156
TR|R9QTA0|R9QTA0_CVHSA             VIGIINNTVYDPLQPELDSFKEELDKYFKNHTSPDVDLGDISGINASVVNIQKEIDRLNE 1155
TR|R9QTH3|R9QTH3_CVHSA             VIGIINNTVYDPLQPELDSFKEELDKYFKNHTSPDVDLGDISGINASVVNIQKEIDRLNE 1156
TR|A0A0K1Z074|A0A0K1Z074_CVHSA     VIGIINNTVYDPLQPELDSFKEELDKYFKNHTSPDVDLGDISGINASVVNIQKEIDRLNE 1156
TR|A0A0U1WHI6|A0A0U1WHI6_CVHSA     VIGIINNTVYDPLQPELDSFKQELDKYFKNHTSPDVDFGDISGINASVVDIQKEIDRLNE 1154
TR|A0A166ZL64|A0A166ZL64_9NIDO     VIGIINNTVYDPLQPELDSFKQELDKYFKNHTSPDVDLGDISGINASVVDIQKEIDRLNE 1151
TR|A0A0U1UYX4|A0A0U1UYX4_CVHSA     VIGIINNTVYDPLQPELDSFKQELDKYFKNHTSPDVDLGDISGINASVVDIQKEIDRLNE 1151
                                   ****:****************:***************:***********:**********

YP_009724390                       VAKNLNESLIDLQELGKYEQYIKWPWYIWLGFIAGLIAIVMVTIMLCCMTSCCSCLKGCC 1248
SP|P59594|SPIKE_CVHSA              VAKNLNESLIDLQELGKYEQYIKWPWYVWLGFIAGLIAIVMVTILLCCMTSCCSCLKGAC 1230
TR|Q6UZF4|Q6UZF4_CVHSA             VAKNLNESLIDLQELGKYEQYIKWPWYVWLGFIAGLIAIVMVTILLCCMTSCCSCLKGAC 1230
TR|Q6UZF0|Q6UZF0_CVHSA             VAKNLNESLIDLQELGKYEQYIKWPWYVWLGFIAGLIAIVMVTILLCCMTSCCSCLKGAC 1230
TR|Q6JH46|Q6JH46_CVHSA             VAKNLNESLIDLQELGKYEQYIKWPWYVWLGFIAGLIAIVMVTILLCCMTSCCSCLKGAC 1230
TR|Q692E4|Q692E4_CVHSA             VAKNLNESLIDLQELGKYEQYIKWPWYVWLGFIAGLIAIVMVTILLCCMTSCCSCLKGAC 1230
SP|Q3LZX1|SPIKE_BCHK3              VAKNLNESLIDLQELGKYEQYIKWPWYVWLGFIAGLIAIVMVTILLCCMTSCCSCLKGAC 1217
SP|Q3I5J5|SPIKE_BCRP3              VAKNLNESLIDLQELGKYEQYIKWPWYVWLGFIAGLIAIVMVTILLCCMTSCCSCLKGAC 1216
SP|Q0Q475|SPIKE_BC279              VAKNLNESLIDLQELGKYEQYIKWPWYVWLGFIAGLIAIVMVTILLCCMTSCCSCLKGAC 1216
TR|R9QTA0|R9QTA0_CVHSA             VAKNLNESLIDLQELGKYEQYIKWPWYVWLGFIAGLIAIVMVTILLCCMTSCCSCLKGAC 1215
TR|R9QTH3|R9QTH3_CVHSA             VAKNLNESLIDLQELGKYEQYIKWPWYVWLGFIAGLIAIVMATILLCCMTSCCSCLKGAC 1216
TR|A0A0K1Z074|A0A0K1Z074_CVHSA     VAKNLNDSLIDLQELGKYEQYIKWPWYVWLGFIAGLVGLFMAIILLCYFTSCCSCCKGMC 1216
TR|A0A0U1WHI6|A0A0U1WHI6_CVHSA     VAKNLNESLIDLQELGKYEQYIKWPWYVWLGFIAGLVGLFMAIILLCYFTSCCSCCKGMC 1214
TR|A0A166ZL64|A0A166ZL64_9NIDO     VAKNLNESLIDLQELGKYEQYIKWPWYVWLGFIAGLVGLFMAIILLCYFTSCCSCCKGMC 1211
TR|A0A0U1UYX4|A0A0U1UYX4_CVHSA     VAKNLNESLIDLQELGKYEQYIKWPWYVWLGFIAGLVGLFMAIILLCYFTSCCSCCKGMC 1211
                                   ******:********************:********:.:.*. *:** :****** ** *

YP_009724390                       SCGSCCKFDEDDSEPVLKGVKLHYT 1273
SP|P59594|SPIKE_CVHSA              SCGSCCKFDEDDSEPVLKGVKLHYT 1255
TR|Q6UZF4|Q6UZF4_CVHSA             SCGSCCKFDEDDSEPVLKGVKLHYT 1255
TR|Q6UZF0|Q6UZF0_CVHSA             SCGSCCKFDEDDSEPVLKGVKLHYT 1255
TR|Q6JH46|Q6JH46_CVHSA             SCGSCCKFDEDDSEPVLKGVKLHYT 1255
TR|Q692E4|Q692E4_CVHSA             SCGSCCKFDEDDSEPVLKGVKLHYT 1255
SP|Q3LZX1|SPIKE_BCHK3              SCGSCCKFDEDDSEPVLKGVKLHYT 1242
SP|Q3I5J5|SPIKE_BCRP3              SCGSCCKFDEDDSEPVLKGVKLHYT 1241
SP|Q0Q475|SPIKE_BC279              SCGSCCKFDEDDSEPVLKGVKLHYT 1241
TR|R9QTA0|R9QTA0_CVHSA             SCGSCCKFDEDDSEPVLKGVKLHYT 1240
TR|R9QTH3|R9QTH3_CVHSA             SCGSCCKFDEDDSEPVLKGVKLHYT 1241
TR|A0A0K1Z074|A0A0K1Z074_CVHSA     SCGSCCRFDEDDSEPVLKGVKLHYT 1241
TR|A0A0U1WHI6|A0A0U1WHI6_CVHSA     SCGSCCRFDEDDSEPVLKGVKLHYT 1239
TR|A0A166ZL64|A0A166ZL64_9NIDO     SCGSCCRFDEDDSEPVLKGVKLHYT 1236
TR|A0A0U1UYX4|A0A0U1UYX4_CVHSA     SCGSCCRFDEDDSEPVLKGVKLHYT 1236
                                   ******:******************
```
